# H3N2 influenza virus tropism shifts to glycan receptors on tracheal ciliated cells

**DOI:** 10.1101/2025.09.12.675939

**Authors:** Chika Kikuchi, Aristotelis Antonopoulos, Shengyang Wang, Assel Biyasheva, Yu-Chun Chien, Ankit Bharat, Ryan McBride, Corwin M. Nycholat, Andrew J. Thompson, Anne Dell, Kay-Hooi Khoo, Robert Schleimer, Stuart M. Haslam, James C. Paulson

## Abstract

Human H3N2 influenza viruses, introduced during the 1968 pandemic, have evolved to recognize human-type sialic acid-containing receptors (Neu5Acα2-6Gal) extended with at least three LacNAc (Galβ1-4GlcNAc) repeats. To investigate this restriction in the context of virus attachment to the airway epithelium, we comprehensively analyzed the glycome of human nasal and tracheal epithelial cells. Using a synthetic N-glycan library that reflects the structural diversity of the human airway glycome, we found that only bi-antennary N-glycans with extended human-type receptors on at least one branch serve as receptors for the recent H3 hemagglutinins (HAs). Such receptors are found on tracheal epithelium but are deficient in nasal epithelium. Immunofluorescence analysis on human trachea reveals that recent H3 HAs preferentially attach to ciliated cells, consistent with single-cell RNA sequencing analysis indicating that these cells express glycosyltransferases that produce extended glycan chains. These findings suggest that H3N2 viruses have developed a tropism for tracheal ciliated cells. (146 words)

## Introduction

Influenza A viruses are a major pathogen in mammals and birds, causing annual human epidemics with moderate to severe respiratory disease and, in some cases, death.^1^ They are membrane-enveloped viruses with two surface glycoproteins, the hemagglutinin (H, HA) that attaches the virus to sialic acid-containing receptors on the surface of airway epithelial cells, and the neuraminidase (N, NA) or ‘receptor-destroying’ enzyme that prevents virus entrapment by mucins, and releases the newly formed virus from the surface of the infected cell. Immunologically distinct subtypes of the HA (19) and NA (11) are coded for by genes on separate segments of the viral RNA genome that can independently sort, creating numerous viruses circulating in animal species that can be distinguished by the HA and NA subtypes they carry (e.g. H1N1, H3N2, H5N1, H7N2).^1,2^ In the last century, three influenza A virus subtypes have been introduced into the human population from an animal virus progenitor, each time causing a major pandemic in an immunologically naïve human population; in 1918 (H1N1), 1958 (H2N2), 1968 (H3N2), and in 2009 with the reintroduction of a novel H1N1 swine reassortant virus. Following the pandemic, each virus continued to cause annual epidemics with constant mutation of the HA and NA to avoid immune recognition.^1^

Adaptation of H1N1, H2N2, and H3N2 pandemic viruses to humans coincided with a major shift in the receptor binding specificity of the hemagglutinins.^1–4^ In each case, the avian virus progenitor bound preferentially to ‘avian-type’ receptors, α2-3-linked sialic acids (Neu5Acα2-3Gal), while the corresponding pandemic viruses bound preferentially or exclusively to ‘human-type’ receptors, α2-6-linked sialic acids (Neu5Acα2-6Gal). In each case, the shift in receptor specificity was the result of two mutations in the receptor binding pocket of the HA, in particular, E190D and G225D for the 1918 H1N1 viruses, and Q226L and G228S for the H2N2 and H3N2 viruses.^5–7^ It is widely believed that these alterations are required for the binding of the virus to Neu5Acα2-6Gal receptors that are strongly expressed on the surface of epithelial cells lining the human upper airway, enabling infection and transmission.^8^ Indeed, using the ferret model of human influenza virus transmission, Tumpey and co-workers showed that two mutations in the hemagglutinin of the 1918 pandemic virus (D190E/D225G) were sufficient to revert its receptor specificity to that of the avian virus progenitor and prevent aerosol transmission between ferrets.^9^ Thus, acquisition of human-type receptor specificity by an avian/animal virus is now widely considered a prerequisite for human transmission, and a risk factor for causing a pandemic.^10^

Upon infection, influenza virus replicates in epithelial cells along the entire upper airway (mouth, nose, sinus, and larynx), the trachea, and the bronchial tree, causing symptomatic tracheobronchitis.^11,12^ The surface of the epithelium comprises mainly ciliated cells and mucin-producing goblet cells. Immunohistochemical analysis with H3N2 virus shows intense staining of ciliated cells in the epithelium of the upper airway, the trachea, and large bronchi, but not epithelial cells of bronchioles or alveoli,^13–15^ which coincides with the presence of human-type (Neu5Acα2-6Gal) receptors detected with *Sambucus nigra* agglutinin (SNA).^8^ Conversely, avian influenza viruses do not bind to ciliated cells of tissues from the upper airway, trachea, and bronchi.^14,15^ In contrast, with air-liquid interface (ALI) cultured tracheal/bronchial epithelial cells, H3N2 viruses primarily bind to and infect non-ciliated cells, while avian influenza viruses bind to and infect ciliated cells.^16,17^ On the surface, the differences in the tropism of H3N2 viruses for ciliated and non-ciliated cells in primary and cultured epithelial cells appear contradictory. Additional information is needed to identify what cells are involved in the initiation of influenza virus infections in the human airway.

Following the pandemic year, influenza viruses persist in the human population under constant antigenic selection, with gradual changes in their receptor binding properties to maintain fitness for infection and transmission in humans.^1,2,18,19^ Notable changes include more stringent recognition of human-type over avian-type receptors, increased avidity for human-type receptors, and a narrowing of receptor specificity to a subset of human-type receptors with extended glycan chains.^5,18–23^ Both H1N1 and H3N2 viruses have acquired a strong preference for human-type receptors extended by at least two LacNAc (Galβ1-4GlcNAc) repeat units (e.g. Neu5Acα2-6Galβ1-4GlcNAcβ1-3Galβ1-4GlcNAc).^19–21,23–25^ This restriction has become particularly stringent for the H3N2 viruses which require extensions of three or more LacNAc repeats.^19,21,24,25^ In recent years, this has become problematic for isolating H3N2 by growth in eggs or Madin-Darby canine kidney (MDCK) cells and for serotyping using hemagglutination because eggs, MDCK cells, and erythrocytes are deficient in such extended glycan receptors.^25–29^ This mismatch of the laboratory host glycome and virus receptor specificity has also impacted vaccine efficacy due to the inadvertent selection of antigenic variants when selecting for viruses that recognize ‘short’ receptors and replicate better in eggs.^30,31^

The restricted specificity of recent H3N2 viruses for extended human-type receptors documented using synthetic glycans,^19–21,24^ implies that such glycans are also present on human airway epithelial cells. Extended N-linked glycans (N-glycans) have been detected in a human epithelial cell line,^20^ in human lung,^32^ and in human nasopharyngeal, bronchiole and lung tissue explants that support infection of both human and avian influenza viruses.^33^ However, what is lacking is a direct analysis of the glycome of primary human airway epithelial cells to determine if human-type extended glycans are present and to determine if other sialylated glycans might serve as alternative receptors of H3N2 viruses.

Here, to understand the restricted receptor specificity of recent H3N2 viruses in the context of the HA maintaining fitness for transmission in humans, we have analyzed the glycome of primary human nasal and trachea epithelial cells. Human-type extended N-glycans are confirmed to be present in expectedly small quantities, particularly in tracheal epithelial cells. Using a library of synthetic N-glycans representing the structural diversity of human-type receptors in the epithelial cell glycome, we found that only a subset of N-glycans with branches containing Neu5Acα2-6Gal extended with three LacNAc repeats can serve as functional receptors. Immunofluorescence staining of trachea epithelium with exemplary HAs from early (1979) and recent (2018) H3N2 strains reveals a shift from binding preferentially to mucin-producing goblet cells to binding primarily to ciliated cells, suggesting that the restricted receptor specificity of recent isolates also changes the host cell tropism in airway epithelia. The results, combined with analysis of the expression of biosynthetic enzymes that make human-type extended glycans, suggest that ciliated cells on the tracheal epithelia provide optimal receptors for recent H3N2 viruses.

## Results

### H3N2 influenza HAs gradually acquired specificity for extended glycan receptors

The gradual adaptation of the receptor specificity of H3N2 viruses to extended glycan receptors is illustrated in **Figure 1**, evaluating recombinant HAs from representative strains from 1968-2020 using a quantitative enzyme-linked immunosorbent assay (ELISA) employing a panel of biotinylated synthetic glycans^29^. Glycans include symmetrical biantennary N-glycans containing mono-, di-, and tri-LacNAc repeats with a terminal human-type α2-6-linked sialic acid, and a linear asialo-glycan as a negative control (**Figure 1A**). The immobilization efficiency was confirmed by the binding of SNA lectin which recognizes terminal human-type NeuAcα2-6Gal motif regardless of the underlying structures (**Figure 1B**). The panel of recombinant H3 HAs includes: A/Hong Kong/1/68 (HK/68), A/Victoria/3/75 (Vic/75), A/Bangkok/1/79 (BK/79), A/Victoria/361/2011 (Vic/11), A/Wisconsin/4/2018 (WI/18), and A/Cambodia/e826360/2020 (Camb/20) (**Figure 1C**). The results show that the HA from the pandemic HK/68 virus binds to human-type α2-6 sialylated N-glycans with 1-3 LacNAc repeats. Over time there is a gradual increase in affinity for the extended glycan with three LacNAc repeats, and gradual loss of avidity to shorter glycans with mono- and di-LacNAcs. For HAs from viruses from 2011 and beyond, this culminates in strict specificity for extended glycans with three LacNAc repeats. The results extend previous observations that the H3N2 influenza viruses have acquired and maintained specificity for α2-6 sialylated N-glycans extended with three LacNAc repeats.

**Figure 1.**
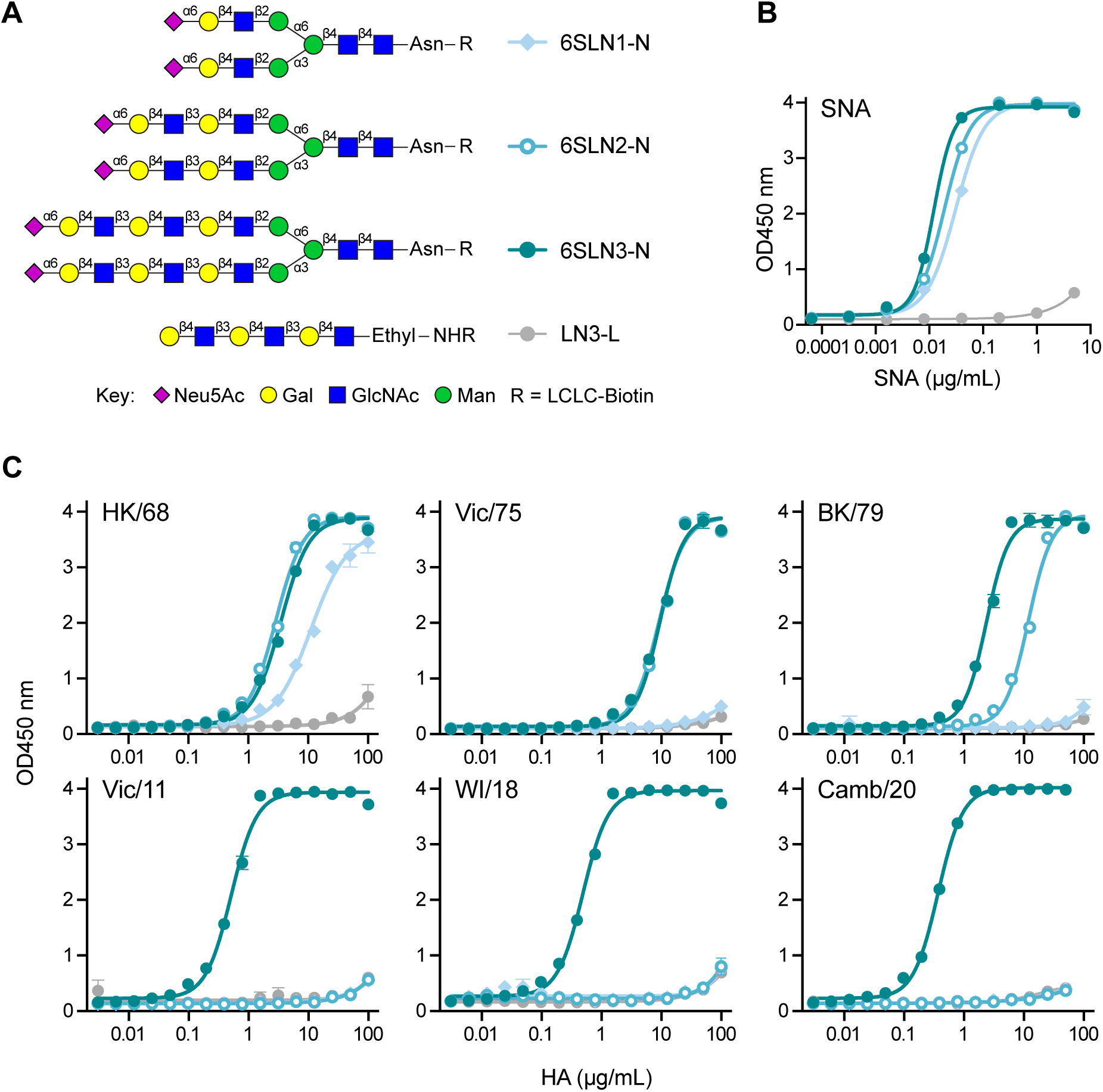
Adaptation of H3N2 viruses to extended human-type receptors. Binding of recombinant HAs from H3N2 viruses from 1968-2020 to biotinylated human-type glycan receptors bound to streptavidin coated ELISA plates. **(A)** Synthetic α2-6 sialylated bi-antennary N-glycans extended with one (6SLN1-N), two (6SLN2-N), or three (6SLN3-N) LacNAc repeats. An asialo tri-LacNAc linear glycan (LN3-L) is used as a negative control. (**B**) Loading of glycans on the ELISA plate is assessed with SNA lectin, specific for α2-6-sialosides. (**C**) Binding of recombinant H3 HAs from human H3N2 influenza strains show gradual restriction of specificity to α2-6 sialylated N-glycans extended with three LacNAc repeats. Data represent the mean of three replicate experiments. Error bars indicate the standard error of the mean (SEM).

### Glycome analysis of human airway epithelial cells reveals extended human-type receptors

We conducted analysis of the glycome of human airway epithelial cells to gain insights into the restriction of recent H3N2 viruses for extended human-type receptors required for virus attachment and initiation of infection. To this end, we collected tracheal and nasal epithelial cells from human donors under informed consent with approval of the Institutional Review Board of the Northwestern Feinberg School of Medicine. Trachea epithelial cells were scraped from discarded portions of eleven human donor transplant lungs, yielding samples consisting of >90% epithelial cells (samples designated TTr01-TTr11). For each trachea, one portion of the cells was frozen immediately as ‘primary’ epithelial cells in numbers (>2x10^6^) sufficient for subsequent glycomics analysis. Another portion was amplified through 2-3 passages in tissue culture. A portion of the cultured cells was further cultured in trans-well plates, to induce air-liquid differentiation with formation of beating cilia, while the remainder were frozen down. Nasal epithelial cells were collected from the inferior turbinate from healthy donors using Rhino-Pro curettes. Samples from 11-18 donors were pooled to achieve a sufficient number of cells (∼2x10^6^) for glycomics analysis.

Analysis of the epithelial cell glycome was conducted as previously described (see Methods).^29,34,35^ Briefly, glycoproteins were detergent extracted from cells and digested with trypsin, followed by treatment with PNGaseF to release N-glycans. After N-glycan release samples were subjected to β-elimination to release O-linked glycans (O-glycans). Glycan samples were then permethylated before further analysis. The N-glycan and O-glycan samples were separately analyzed by matrix-assisted laser desorption ionization-time of flight mass spectrometry (MALDI-TOF MS).^34,35^

For tracheal epithelial cells, N-glycan preparations from nine donor trachea were analyzed (**Figure 2, Figure S1**). Portions of the samples from three donors were also analyzed after treatment with sialidase S (Sial-S), a bacterial sialidase that removes only the α2-3-linked NeuAc residues (**Figure S2**). This treatment leaves glycans with α2-6-linked sialic acids that are candidate receptors for human influenza viruses. An example of the MS spectra for N-glycans from tracheal epithelial cells from one donor is illustrated in **Figure 2A**, where peak intensity is normalized to that of the most abundant oligomannose glycan, Man_9_GlcNAc_2_ (m/z 2396). In the inset, an expanded scale is shown to better visualize the region of larger complex-type glycans that have sialic acids (m/z 2415-7000), and the red box indicates the zoomed region in **Figure 2C**. Molecular ion abundances in this mass range were between 6.1% to 0.1% of the peak intensity of the most abundant Man_9_GlcNAc_2_ glycan (**Figure S1**). Despite these low abundances, signal-to-noise was excellent (**Figure S1**). This allowed MS/MS experiments to be conducted on the majority of peaks to obtain glycan compositions of the fragments of multi-antennary glycans (**Figures S3, S4**). Remarkably, the annotated N-glycans from the mass spectra of tracheal epithelial cells were highly complex, with more than 600 distinct N-glycan species representing glycans ranging in molecular weight from m/z 1579 to 6547 (see **Figure S5).** We found that the N-glycans detected were mostly core-fucosylated, bi- to tetra-antennary structures, with the antennae further extended with LacNAc increments, and richly decorated by fucose, blood group structures, and sialic acids.

**Figure 2.**
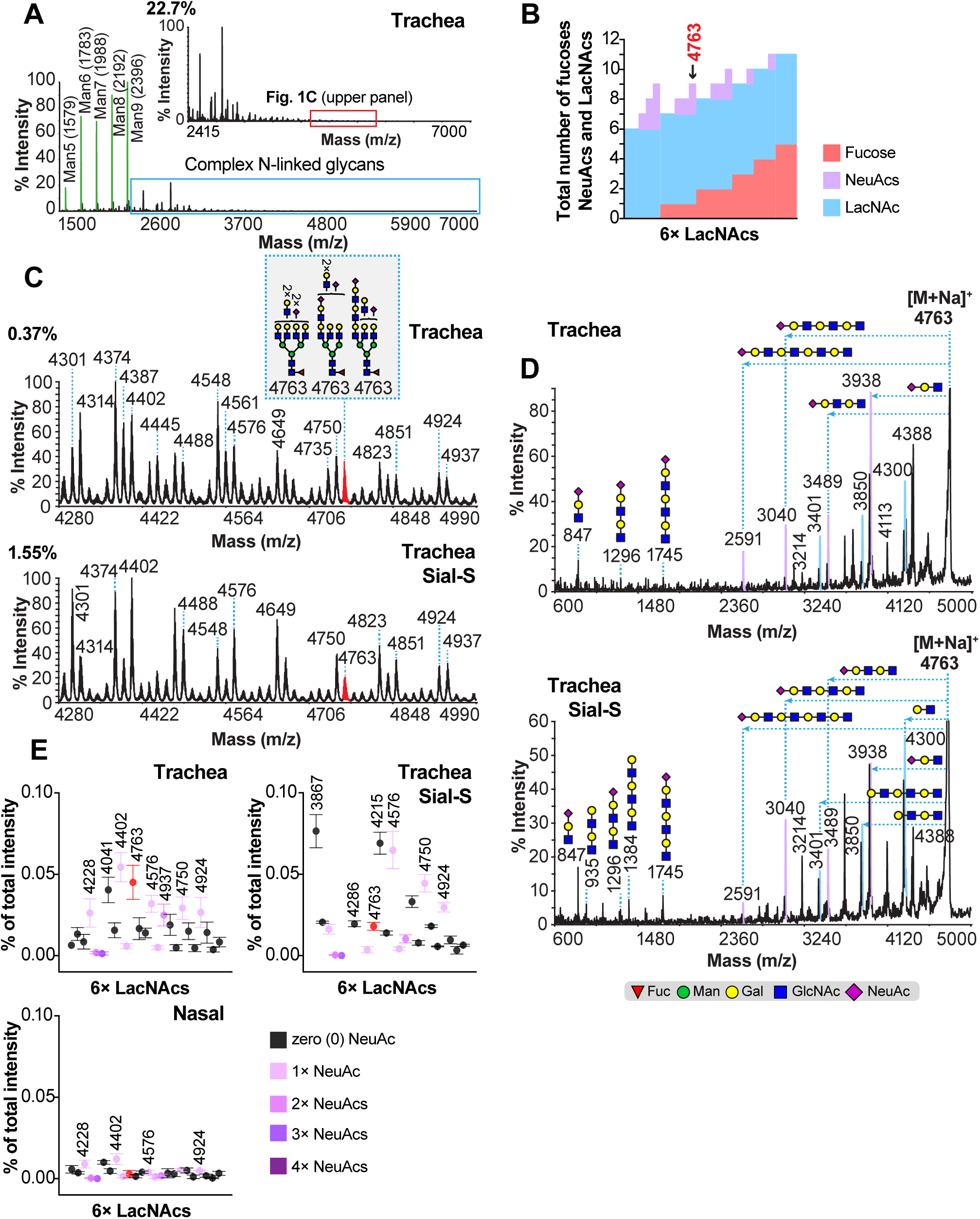
MS analysis reveals extended human type receptors in the glycome of human airway epithelial cells. **(A)** MALDI-TOF mass spectrum of permethylated N-glycans of human trachea epithelial cells. Green-colored peaks correspond to high mannose N-glycans (Man_5_GlcNAc_2_ to Man_9_GlcNAc_2_). Inset depicts the expanded section of the spectrum (blue box) area where complex N-glycans are detected and the red box in the inset is further zoomed in Figure 2C. (**B**) Annotations of MS peaks up to m/z 5000 with N-glycan compositions corresponding to a Man_3_GlcNAc_2_ core with six (6) LacNAc repeats, zero-three (0-3) NeuAc residues and zero-five (0-5) fucose residues. (**C**) Expanded region of MALDI-TOF mass spectra of N-glycans from tracheal epithelial cells corresponding to red box in panel **A** inset, before (top) and after (bottom) digestion with Sial-S. The red-colored peak in both panels corresponds to the molecular ion at m/z 4763, consistent with the annotated structures. (**D**) MALDI-TOF/TOF MS/MS spectra of the molecular ion at m/z 4763 before or after a Sial-S digestion (**upper** and **lower panel** respectively). Spectra are annotated for loss of sialylated antennae from the molecular ion with one to four LacNAc repeats (purple peaks), and the appearance of corresponding fragments at lower m/z. (**E**) Relative abundance of the molecular ions corresponding to N-glycans with six (6) LacNAc repeats for samples from trachea before (n=9) or after (n=3) treatment with Sial-S and nasal samples (n=3). As described in Methods, the value for each m/z represents a percentage (Avg ± SEM) of the sum of all peak intensities except those for the high mannose glycans (e.g. green peaks in panel **A**). Symbols are color coded based on the number of NeuAc residues found on the corresponding molecular ions. The red symbol is m/z 4763 analyzed in panel **D**.

To identify candidate receptors of recent H3N2 influenza viruses, we focused on the mass range above m/z 4000 for glycans that are large enough to have sialylated antennae with at least three LacNAc repeat units. Since the sialyltransferase responsible for attachment of sialic acid in α2-6 linkage cannot use LacNAc substituted with fucose as an acceptor substrate,^36,37^ we initially searched for molecular ions with compositions compatible with sialylated antennae with 6 LacNAc units and without terminal fucose substitution. As illustrated in **Figure 2B**, glycans with 6 LacNAc units were variably substituted with sialic acids (NeuAc) and fucose residues. One candidate that met the criteria is the molecular ion at m/z 4763, corresponding to six LacNAcs, two NeuAc residues, and a single fucose residue (e.g. a core fucose). This molecular ion is found in the spectra of all the tracheal samples (**Figure 2C upper panel** red peak, and **Figure S1**). Importantly, it is also found after Sial-S digestion (**Figure 2C lower panel**, red peak, and **Figure S2**), indicating the presence of two α2-6-linked NeuAc residues. Because this composition is consistent with multiple structural isomers, MALDI-TOF/TOF MS/MS analysis was performed on m/z 4763 before and after digestion with Sial-S (**Figure 2D**, **Figure S4**). This analysis revealed that the composition at m/z 4763 was comprised of core-fucosylated structural isomers with NeuAc-terminated poly-LacNAc branches of various lengths. Specifically, the fragment ions detected at m/z 3938, 3489, 3040, and 2591 corresponded to losses of antennae with a NeuAc residue attached to one, two, three and four LacNAc repeats, respectively, accompanied by complementary fragments in the low mass range of the spectrum corresponding to NeuAc with one, two or three LacNAc repeats (m/z 847, 1296 and 1745). Thus, since the same fragmentation pattern was detected before and after Sial-S digestion, we conclude that the molecular ion at m/z 4763 includes structural isomers of N-glycans carrying two α2-6-sialylated antennae with one to four LacNAc repeats (see inset for examples).

Following the same steps as above, additional molecular ions at m/z 3503, 3864, 3952, 4314, and 4402 are representative of glycans with similarly extended NeuAcα2-6-linked poly-LacNAcs (**Figure S4**). Moreover, MS/MS analysis of molecular ions at m/z 3952 and 4402 from spectra before and after Sial-S digestion (compare left and right panels, **Figure S4C** and **S4E**) showed an increase in the relative abundance of fragment ions at 3040 and 3489 after Sial-S digestion. These fragment ions correspond to losses of two (2) and three (3) LacNAc repeats, which match with increases in lower molecular weight fragments of m/z 935 and 1384 consistent with neutral branches of glycans previously capped with α2-3-linked sialic acid removed by Sial-S. Taken all together, the results indicated the presence of glycans with antennae containing three to four LacNAc units capped with α2-6-linked sialic acid.

Nasal epithelial cell samples were analyzed in parallel to the trachea samples. The spectra revealed that the N-glycan structures were similar to those in the human trachea samples in terms of fucosylation, sialylation, and blood group antigens (**Figures S6, S7**). Accordingly, the glycan peaks are annotated together with those from trachea epithelial cells (**Figure S5**). A major difference between the spectra of N-glycans from nasal epithelial cells relative to tracheal cells was the low relative abundance of the molecular ions that corresponded to larger glycans with poly-LacNAc extensions (**Figure 2E, Figure S8**). This is illustrated in **Figure 2E** that compares the abundance of the large glycans with six LacNAc units relative to the sum of the peak intensity of all complex type glycans (e.g. excluding the peaks corresponding to oligomannose glycans). Using the molecular ion at m/z 4763 as an example, the relative abundance in trachea was 0.045% (n=9) and 0.018% (n=3) before and after Sial-S digestion, respectively (**Figure 2E**). By contrast, for nasal spectra, the relative abundance of m/z 4763 had a mean value of 0.003% (n=3), which is approximately 15 times less than trachea. Similarly, the relative abundance of other large glycans with four to six LacNAc units (**Figure S8**) were dramatically reduced in the nasal N-glycans (**Figure 2E**, **Figures S8**). The data suggest that N-glycans with α2-6 sialylated poly-LacNAc branches comprising the preferred receptors of H3N2 viruses are much more abundant on tracheal epithelial cells than nasal epithelial cells.

To further identify candidate receptors for influenza virus in the trachea samples, we fractionated N-glycans on an SNA lectin column that retains glycans terminated with the NeuAcα2-6Gal linkage (**Figure S9**). Relative to the MS spectra of the total glycans (**Figure S9A**), the MS spectra of glycans that flowed through the column were enriched in non-sialylated glycans (**Figure S9B)**, consistent with the majority of the sialylated glycans containing at least one sialic acid terminated in the NeuAcα2-6Gal linkage. Glycans eluted from the column were highly enriched in glycans containing sialic acid (**Figure S9C**), including glycans annotated to contain sialylated poly-LacNAc antennae (e.g. m/z 3503, 3952, 4402, see **Figure S4**). Also prominent are multi-antennary sialylated glycans that are highly fucosylated (e.g. m/z 4026, 4200, 4548). These are notable because α2-6 sialylation of LacNAc substituted with fucose on either the GlcNAc or Gal is biosynthetically disfavored^37^, requiring the sialic acid to be on a non-fucosylated antennae (**Figure S10**).

Analysis was also performed on O-linked glycan fractions from primary human trachea and nasal epithelial cells (**Figures S11** and **S12**). Glycans with linear Galβ1-3GalNAc (Core-1) and branched Galβ1-3(GlcNAcβ1-6)GalNAc (Core-2) core structures were detected in all samples. Overall trachea O-glycans exhibited structures of much higher molecular weight (up to m/z 4000) compared to the nasal O-linked glycans. The O-linked glycans exhibited high levels of fucosylation and blood group antigens. Sialic acids were observed primarily on highly fucosylated antenna (**Figure S12**), representing sialyl Lewis X (NeuAcα2-3Galβ1-4(Fucα1-3)GlcNAc) or sialyl Lewis A (NeuAcα2-3Galβ1-3(Fucα1-4)GlcNAc) epitopes. We conclude that O-glycans have little or no α2-6 sialylated glycans as human-type receptors for influenza virus due to the high levels of fucosylation that prevent sialylation by α2-6 sialyltransferases.

Finally, we evaluated sulfated N-glycans and O-glycans, since sialylated/sulfated glycans have been implicated as functional receptors of avian influenza viruses viruses.^38–40^ Analysis of sulfated glycans required a different procedure since sulfates are lost in the standard processing (see Methods). For N-glycans there was little evidence for both sialic acid and sulfate on the same LacNAc moiety (**Figure S13**). While there was evidence for sialylated/sulfated glycans on O-glycans with a composition consistent with 6-sulfo-sialyl-Lewis-X (6S-NeuAcα2-3Galβ1-4(Fucα1-3)GlcNAc, these structures have previously only been identified as a potential receptor for avian viruses. Since no data have implicated sulfated α2-6 sialylated glycans as human influenza virus receptors to date, we did not further pursue the analysis of sulfated glycans in the glycome of trachea epithelial cells.^38^

### Synthetic glycan library reflecting the diversity of human-type receptors in the airway glycome

Having concluded that N-glycans with extended antennae are the primary candidate receptors for recent H3N2 viruses, we assembled a library of glycans that represented the structural diversity of α2-6 sialylated N-glycans in the glycome of tracheal and nasal epithelial cells (**Figure 3A**). A primary consideration in selecting the glycans was to determine the extent to which extended human-type receptors were required for recognition and what other dominant structural features might impact recognition. Accordingly, we selected 21 bi-, tri- and tetra-antennary core-fucosylated N-glycans, with compositions defined by m/z values (**Figure S5**), and antennae compositions identified by MS/MS experiments (**Figures S3, S4, S7, and S10**).

**Figure 3.**
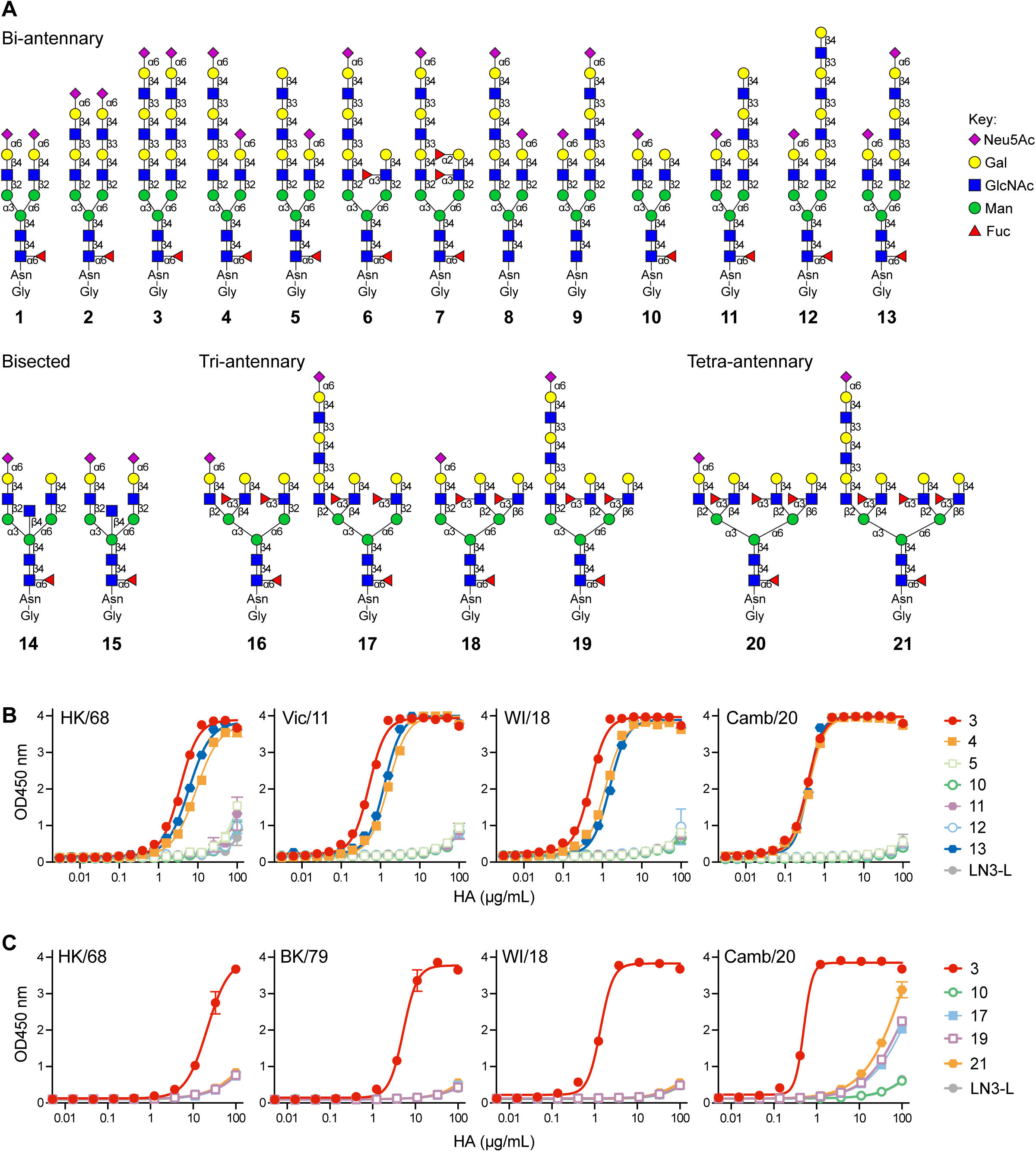
Specificity of H3 HAs for asymmetric human type N-glycan receptors. (**A**) Library of synthetic N-glycans representing candidate extended human-type receptors in the glycome of human tracheal epithelial cells. (**B,C**) Binding of recombinant H3 HAs to representative asymmetric (**B**) bi-antennary and (**C**) tri- and tetra-antennary glycans. All N-glycans are biotinylated and adsorbed to streptavidin coated ELISA plates. Loading of glycans was assessed using the SNA lectin (**Figure S14A,B**). They symmetrical tri-LacNAc biantennary N-glycan (3) is used as a positive control. LN3-L, a synthetic asialo linear tri-LacNAc glycan, was used as a negative control. Data represent the mean and SEM of three replicates and are representative of two independent experiments.

The library includes three di-sialylated, symmetrical core fucosylated glycans with one to three LacNAc repeats (**1-3**) analogous to the non-core-fucosylated glycans used in **Figure 1A**. For asymmetric biantennary glycans, variations included di-sialylated glycans with one extended (tri-LacNAc) antenna and one short (mono-LacNAc) antenna (**4**,**8,9**,**13**); mono-sialylated glycans with sialic acid on the extended (tri-LacNAc) antenna and no sialic acid on a short mono-LacNAc antenna substituted with fucose (**6**,**7**); and mono-sialylated glycans with the sialic acid on a short (mono-LacNAc) antenna and a non-sialylated antenna with one or three LacNAc repeats (**5**,**10**,**11**,**12**). Also included were two sialylated bisected biantennary glycans (**14**,**15**). Finally, there are mono-sialylated tri-antennary and tetra-antennary glycans with sialic acid on an extended (tri-LacNAc) or short (mono-LacNAc) antenna, with the other mono-LacNAc antenna substituted with fucose. These highly fucosylated glycans were a prominent feature of α2-6-linked mono-sialylated glycans (**Figure S3, S5, S9, S10**). For the tri-and tetra-antennary glycans sialic acid was placed on the GlcNAcβ1-2Manα1-3Man branch due to the strong branch specificity of the ST6Gal1 sialyltransferase.^36^ This glycan library with an Asn-Gly linker was synthesized employing a chemoenzymatic synthesis strategy.^41^ The amine function of the Asn-Gly linker facilitated conjugation to biotin and NHS-activated glass slides for analysis of H3 HA receptor specificity.

### Specificity of H3 HAs for human-type N-glycan receptors on airway epithelial cells

HAs from H3N2 viruses from 1968 to 2020 were compared for their specificity towards the library of biotinylated glycans using the ELISA type assay (**Figures 3A, 3B, S14**), and using a printed glycan microarray (**Figure S15, Table S1, Table S2**). On the glycan microarray, all the H3 HAs bound to the symmetrical core fucosylated di-sialylated N-glycans (**1-3**), with the WI/18 and Camb/20 HAs showing strict specificity for the extended glycan with three LacNAc repeats (**3**) and the earlier HK/68, Vic/75 and BK/79 HAs also binding to the glycans with one and/or two LacNAc repeats (**1**,**2**), as seen previously in glycan arrays for the di-sialylated non-core-fucosylated symmetrical glycans.^21,24^ Overall, however, binding in the glycan microarray assay was modest, and information for comparing the non-symmetrical glycans was limited. For this reason, we focused on the ELISA based assay for the asymmetric glycans.

Binding of H3 HAs to glycans in the ELISA assay revealed that di-sialylated biantennary N-glycans with only one extended arm (**4**,**13**) were bound by HK/68, Vic/11, WI/18 and Camb/20 with slightly reduced avidity relative to that of the di-sialylated symmetrical tri-LacNAc N-glycan used for reference (**3**), except for Camb/20 which bound with equal avidity (**Figure 3B**). Binding was nearly equivalent when the extended arm was on the Manα1-3Man branch, or the Manα1-6Man branch (**4**,**13**). Substituting the sialic acid on the short branch, with Fucα1-3GlcNAc reduced avidity (**Figure S14C**, compare **4** and **6**). Binding was dependent on the sialic acid present on the extended branch, since N-glycans with sialic acid only on the short branch (**5**,**11**,**12**) were not recognized. The di-sialylated bisected N-glycan (**18**) was not bound by Vic/11 or WI/18 HAs (**Figure S14C**).

We analyzed several tri- and tetra-antennary glycans with α2-6 sialic acid on one antenna, and LacNAc substituted with fucose on the other antenna (**Figure 3C**, **S14**). Surprisingly, the tri- and tetra-antennary glycans with an α2-6 sialic acid on a single tri-LacNAc extended antenna (**17**, **19**, **21**) were not recognized by HK/68, BK/79 and WI/18 and only weakly recognized by Camb/20 which exhibits the highest avidity for human-type receptors. As expected, a representative tri-antennary glycan with the sialic acid on a short mono-LacNAc antenna (**18**) and fucosylated LacNAc on the other antenna is also not recognized (**Figure S14**). We propose that the lack of recognition of the glycans with NeuAcα2-6-tri-LacNAc is due to the fucose substitution of the other branches.

Previously we found that symmetrical α2-6-sialylated tri-antennary glycans with tri-LacNAc extensions were avidly bound by H3 HAs from 1968-2018.^21^ Thus, while tri- and tetra-antennary glycans with multiple α2-6-sialylated extended antennae could serve as receptors for H3 viruses, we do not detect such glycans at detectable levels in the trachea airway glycome. Although compositions of m/z peaks seen in MS spectra of several trachea samples (e.g. m/z 5661 and 6023) are consistent with tri-antennary glycans with multiple sialylated extended branches, they were not detected following treatment of the samples with Sial-S, suggesting that one or more of the sialic acids were in α2-3 linkage (**Figures S1**, **S2**). Thus, since we have no evidence for tri- and tetra-antennary glycans with two or more α2-6-sialylated tri-LacNAc antennae, and since the representative glycans with a single α2-6-sialylated LacNAc extended antennae are not recognized (**Figure 3C**), we conclude that bi-antennary N-glycans with one or two α2-6-sialylated extended antennae are the predominant receptors of recent H3N2 viruses.

### Differential glycosylation of primary and ALI-cultured epithelial cells

We considered the possibility that primary epithelial cells differed in their glycosylation from ALI cultured epithelial cells based on seemingly contradictory reports that human influenza binds to ciliated cells in trachea tissue sections, but binds to and infects non-ciliated cells in ALI culture.^16,17^ Qualitatively, there is a notable difference in the staining of trachea sections and ALI cultured bronchial epithelial cells with NeuAcα2-6Gal specific SNA lectin used for detection of human-type receptors (**Figure 4A**).^8,20^ As shown in **Figure 4A**, while there is staining of ciliated cells and non-ciliated secretory cells in both sections, staining of the non-ciliated cells is more pronounced in ALI cultured cells, consistent with reports that H3N2 viruses preferentially infect non-ciliated cells in ALI-culture.^16,17^ To directly compare the glycome of primary and ALI cultured cells, we performed MS analysis on two sets of tracheal epithelial cells (TTr02 and TTr03) and the same cells grown in ALI culture (TTr02-ALI and TTr03-ALI). As evident from the portions of the MALDI-TOF mass spectra in **Figure 4B**, the spectra from the native (primary) cells and ALI cultured cells are visually quite different. However, TOF/TOF MS/MS on select molecular ions from the native and cultured cells showed that the N-glycans have similar compositions and structures, differing mainly in relative abundance (**Figure 4B**, and **Figures S1C/D, S16** and **S17**). We applied a de-isotoping and quantification algorithm to the peaks in both spectra to investigate the effect of ALI culture on the relative abundance of the peak areas of each molecular ion.^42^ Although the molecular ion abundances varied between the two sets of primary samples (e.g. TTr02 and TTr03; **Figure S1C/D**), comparison of the spectra from the primary (Native) corresponding cultured (ALI) samples revealed increases or decreases for numerous molecular ions in both sample sets (**Figure 4B**, **Figure S18**). Of the 274 identified N-glycan molecular ions between m/z 1590-4402, 49% showed a change in their abundance, of which 56% were sialylated. In general, there appeared to be reduced fucosylation in the ALI cultured cells, as evidenced by a reduction in peaks with high fucose (e.g., m/z 3737, 4361, 4535, 4984) and an increase in corresponding peaks with lower fucose (e.g., m/z 3664, 4026, 4649 and 4911).

**Figure 4.**
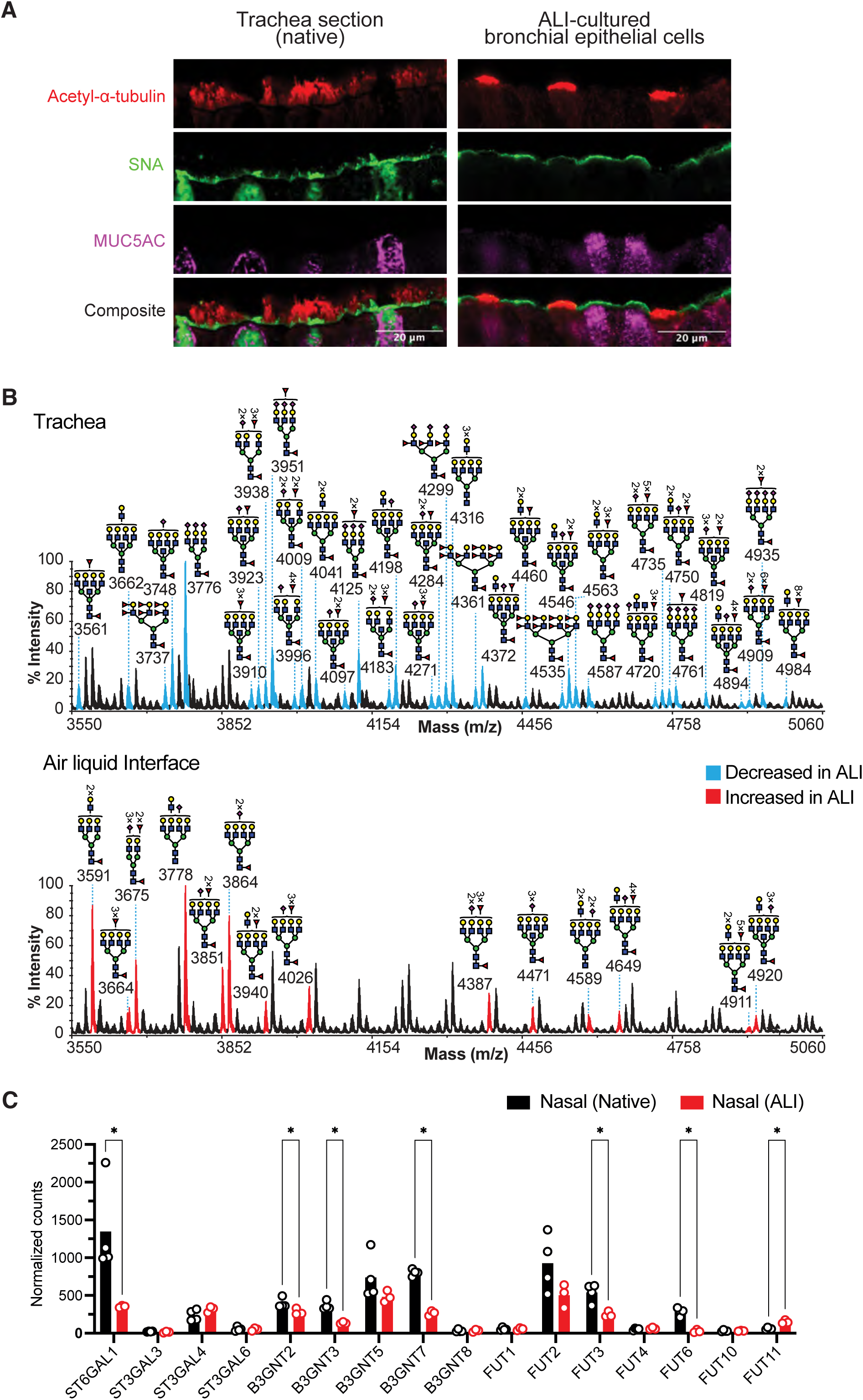
Differential glycosylation of primary and air liquid interface (ALI) cultured human airway epithelial cells. (**A**) Immunofluorescent images of a section of human trachea (left) and cross-section of ALI-cultured bronchial-epithelial cells (right) stained with anti-acetyl-α-tubulin (*red*) for ciliated cells, SNA (*green*) specific for α2-6 sialylated glycans, and anti-MUC5C (*purple*) for goblet cells. (**B**) MALDI-TOF MS spectra (m/z 3550-m/z 5060) of permethylated N-glycans from Native (top; TTr02) and ALI-cultured (bottom; TTr02-ALI) tracheal epithelial cells. Selected peaks are annotated for glycan structures as described in methods. Highlighted peaks are those found to be reduced following ALI culture (*blue*, top panel), or found to be increased after ALI culture (*red*, bottom panel). (**C**) Comparing expression levels of selected glycosyltransferases detected by RNAseq for primary (black bars) or ALI-cultured (red bars) nasal epithelial cells. Bars represent the mean of values from biological replicates of 4 native and 3 ALI-cultured samples of nasal epithelial cells. * p < 0.05.

Because glycosyltransferase gene expression is fundamental to the glycome produced by a cell, we compared the expression of glycosyltransferases in primary and ALI-cultured nasal epithelium samples using bulk RNAseq analysis. As shown in **Figure 4C**, several sialyltransferases, fucosyltransferases, and β3GNT enzymes show significant changes when the cells are grown in ALI-culture. Notably, ST6Gal1, the sialyltransferase responsible for the synthesis of human-type influenza virus receptors (NeuAcα2-6Gal) is significantly decreased in the ALI-cultured cells, as are several of the enzymes in the β3GNT family (e.g. β3GNT2, β3GNT3, β3GNT7) responsible for bi- and tri-LacNAc extended glycan sequences. Several fucosyltransferases are also reduced (e.g. FUT3 and FUT6), consistent with evidence of reduced fucosylation in ALI-cultured tracheal epithelial cells (**Figure 4B**) as discussed above. Taken together, the results in **Figure 4** provide evidence for altered glycosylation in ALI-cultured epithelial cells that could contribute to the differences seen in H3N2 virus binding to ciliated vs non-ciliated epithelial cells in sections of primary airway tissue and ALI-cultured epithelial cells.^14–17^

### Recent H3N2 viruses exhibit preferred tropism for ciliated cells

To assess the impact of the restricted receptor specificity of recent H3N2 viruses on binding to airway epithelial cells, we compared the binding of the HAs from BK/79 and WI/18 to sections of human tracheal epithelium (**Figure 5**). Trachea sections were stained with acetyl-α-tubulin for cilia, FOXJ1, a cytoplasmic marker of ciliated cells, and the mucin MUC5AC, a marker for mucin-producing goblet/secretory cells (**Figure 5A, B, Figure S19A, B**). SNA staining reveals broad staining across the entire apical surface, including ciliated cells and goblet cells (**Figure 5B, Figure S19A)** as also seen by others.^8,20^ The BK/79 HA, an older strain with less stringent receptor specificity (**Figure 1,3**), showed staining of the apical surface of both goblet (secretory) cells and ciliated cells (**Figure 5B, Figure S19B)**. In contrast, the WI/18 HA that exhibits strict specificity for extended N-glycan receptors predominantly stained the apical surface of ciliated cells with negligible staining of goblet cells (**Figure 5B, Figure S19B**). To assess this pronounced shift in tropism more quantitatively, we calculated the shortest distances between each HA punctum and the surface of FOXJ1^+^ ciliated cells or MUC5AC^+^ goblet cells by analyzing z-stacked images (**Figure 5C**, see Methods). The results showed that WI/18 HA is associated with FOXJ1^+^ ciliated cells, with 98.0% of HA staining pixels being closer to the FOXJ1^+^ marker of ciliated cells, while for BK/79 HA, one-third of the puncta were closer to MUC5AC^+^ staining cells (**Figure 5D**). Overall, the results show that the early H3 HA (BK/79) binds to both secretory and ciliated cells, while the more recent H3 HA exhibits a more restricted tropism to ciliated cells.

**Figure 5.**
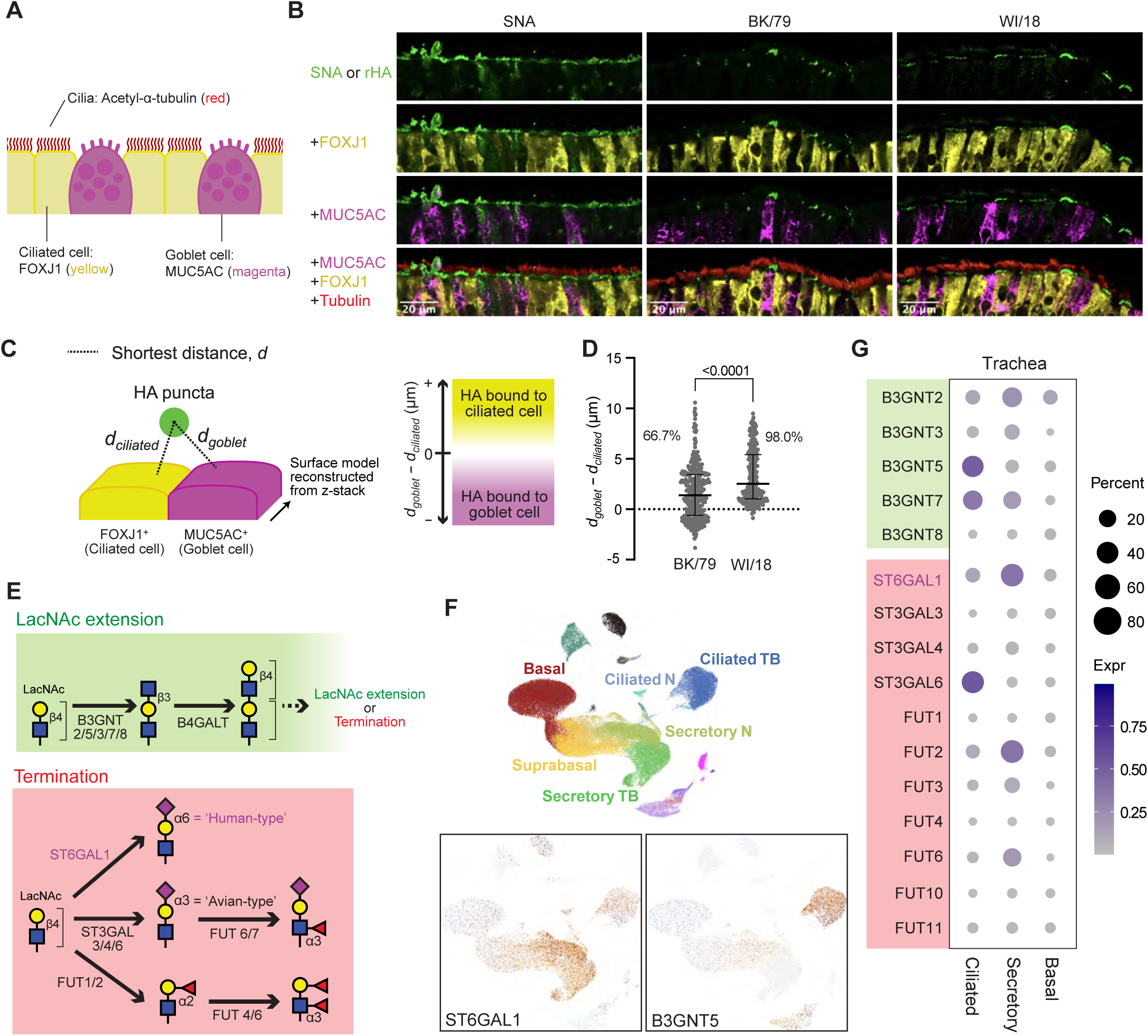
Preferential binding of WI/18 HA to ciliated epithelial cells coincides with biosynthetic capacity for synthesis of extended human-type receptors. (**A**) Schematic of the airway epithelial layer and cell type-specific markers for ciliated cells, acetyl-α-tubulin (*red*) and FOXJ1 (yellow), and for secretory/goblet cells, MUC5AC (*magenta*). (**B**) Human trachea sections were stained with cell type markers and either SNA lectin (*left)*, BK/79 recombinant HA (rHA) (*middle*), and WI/18 HA (*right)* (*green*). See Figure S18 for additional images. (**C**) Diagram of the analyses employed to determine whether the HA staining is closer to ciliated or goblet cells, based on a 3D model constructed from confocal images. (**D**) Proximity of BK/79 and WI/18 HA staining to markers of secretory/goblet cells and epithelial cells. (**E**) Glycan biosynthetic pathways for LacNAc extension (*top*) or terminating a LacNAc-terminated glycan substrate (*bottom*). (**F**) Uniform manifold approximation and projection (UMAP) representation of cell types from human nasal (N) and tracheobronchial (TB) samples (*upper panel*) and expression of ST6GAL1 (*lower left*) and B3GNT2 (*lower right*). (**G**) Expression of glycosyltransferase genes in ciliated, secretory/goblet, and basal cell populations in trachea. Data was reanalyzed from the publicly available scRNAseq data of Deprez et al (2020).

### Ciliated cells can express extended human-type receptors

To assess the possibility that the WI/18 HA tropism to ciliated cells was related to its restricted specificity for extended α2-6-sialylated N-glycan receptors, we analyzed the expression of the biosynthetic enzymes needed to produce these sequences, using publicly available scRNAseq datasets from healthy human nasal, tracheal, and bronchial epithelia.^43^ The enzymes involved in the biosynthesis of extended α2-6 sialylated branches extending from the core structure of N-glycans (Man_3_GlcNAc_2_) are illustrated in **Figure 5E**. These glycan sequences are synthesized in a non-template fashion based on the specificities of the glycosyltransferases present in the Golgi apparatus.^8,20,44^ A single LacNAc extension, produced by all cells, can be further extended by the addition of a β1-3 GlcNAc by members of the β3-N-acetyl-glucosaminyltransferase family (B3GNTs) and a ‘housekeeping’ β4-galactosyltransferase (B4GALT) to complete a second LacNAc repeat, which can then be further extended (**Figure 5E**, top). Extension is blocked by the addition of terminal sugars such as sialic acid or fucose by sialyl- or fucosyltransferases, respectively (**Figure 5E**, bottom).^37^ Notably, the extension and termination of extension are competitive reactions with the β3GNTs and the sialyl-, and fucosyl-transferases competing for a common acceptor substrate.^37,45^ Therefore, higher expression of the β3GNTs favors extension, while higher expression of the sialyl- or fucosyltransferases result in shorter antennae due to the termination of LacNAc extension by capping with sialic acid or fucose.

The key enzymes involved in the synthesis of extended α2-6-sialylated glycans comprising human type receptors are differentially expressed in ciliated cells and mucin-producing secretory cells (**Figure 5F,G, Figure S20)**. Both cell types express the enzyme ST6GAL1 that terminates extension with the NeuAcα2-6Gal sequence, creating human-type receptors (**Figure 5F**), with even greater expression seen in secretory cells. This is consistent with the binding of SNA to both secretory (goblet) cells and ciliated cells. ST3GAL6, a sialyltransferase that transfers sialic acid in α2-3 linkage (**Figure 5G**), is also highly expressed in ciliated cells. However, it appears to be biosynthetically of little consequence since trachea epithelium expresses predominately human-type receptors with sialic acids in α2-6-linkage.^8^ Relevant to the extension of glycan chains, several members of the β3GNT family are expressed in both cell types, T2, T3, T5, T7, and T8, all of which can use LacNAc on N-glycans as substrates and contribute to poly-LacNAc extension.^46–48^ Notably, T5 and T7 are highly expressed in ciliated cells relative to secretory cells (**Figure 5F,G**, **Figure S20**). Another difference between the two cell types is a higher expression of fucosyltransferases in secretory cells (**Figure 5G**), which block extension by transferring fucose to the Gal (FUT2; Fucα1-2Galβ1-4GlcNAc) or GlcNAc (FUT3, FUT6; Galβ1-4(Fucα1-3)GlcNAc). The combination of increased expression of the β3GNT enzymes and reduced expression of the terminal fucosyltransferases would favor the synthesis of extended poly-LacNAc sequences in ciliated cells. Thus, we propose that HAs from recent H3N2 viruses exhibit tropism to ciliated cells due to their biosynthetic capacity to produce extended α2-6-sialylated human-type receptors.

## Discussion

Human influenza virus pandemics in 1918 (H1N1), 1957 (H2N2), 1968 (H3N2), and 2009 (H1N1) resulted from the introduction of novel virus strains with HAs from influenza viruses of avian or swine origin.^2,23,49–53^ In all cases, only one or two mutations were needed for their hemagglutinin to acquire the ability to bind human-type, α2-6-sialylated receptors,^49–52^ a property now widely regarded as required for efficient transmission in the human population.^2,9^ Following the pandemic years, the H1N1 and H3N2 viruses gradually developed a restricted specificity and higher avidity for a subset of human-type receptors.^19–25^ In particular, based on the analysis of HA binding to libraries of synthetic glycans, H1N1^20,22,23,25,41^ and H3N2^2,19,21,25,54^ viruses have acquired strict specificity for α2-6-sialylated receptors extended with at least two or three LacNAc repeats, respectively.

How this restricted receptor specificity relates to receptors on the human airway is less clear. Glycomics analysis has been previously conducted on cultured human bronchial epithelial cells,^20^ human bronchial and lung tissue explants,^33^ and whole human lung,^32^ providing evidence for the presence of α2-6- and α2-3-sialylated LacNAc extended N-glycans in human lung. In this report, we have conducted glycomics analysis directly on primary human trachea and nasal epithelial cells to gain further insights into the restricted specificity of human H3N2 influenza viruses.

Our results show that human nasal and tracheal epithelial cells express among the most complex and diverse glycomes yet to be reported. Although the main focus of this study is receptors of influenza virus, the airway glycome will also be of high relevance for interactions of the diverse array of pathogenic and commensal microbes that interact with airway epithelial cells. For recent H3N2 influenza viruses, we find that N-glycans of tracheal epithelial cells meet the required specificity of the H3 HAs for α2-6-sialylated antennae extended with three or more LacNAc repeats (**Figure 1, Figure S4**). Notably, in total, they represent a minor fraction (<1%) of the sialylated N-glycans in the trachea glycome. Although nasal epithelial cells contain N-glycans of the same composition, they are much less abundant than those on tracheal epithelial cells (**Figure 1, Figure S8**). Tracheal O-glycans also had sialylated, extended glycan chains, but they were highly fucosylated, and as such are unlikely to be α2-6-sialylated since fucose addition to either the Gal or GlcNAc of LacNAc blocks transfer of sialic acid in α2-6-linkage by ST6Gal I (**Figure S12**). We conclude that the candidate receptors of recent H3N2 viruses are N-glycans enriched in tracheal epithelial cells.

Analysis of the binding of H3N2 HAs to synthetic N-glycans representing the structural diversity of the epithelial cell glycome revealed further insights into their specificity for human type receptors (**Figure 3**). Importantly, H3 HAs bind well to sialylated bi-antennary glycans extended with three LacNAc repeats on a single antenna (**4**, **13**) or both antennae (**3**) (**Figure 3**), consistent with the conclusions of Broszeit *et. al*.^25^ Equivalent avidity was seen if the single extended antenna was on the Manα1-3Man (**4**) or Manα1-6Man (**13**) branch (**Figure 3B**). In contrast, tri-antennary and tetra-antennary glycans with a single sialylated extended antenna on the GlcNAcβ1-2Manα1-3Man branch and the other branches with a single LacNAc substituted with fucose (**17**,**19**,**21**) bound weakly or not at all (**Figure 3C**). We previously found that H3 HAs and H3N2 viruses bind equally well to α2-6-sialylated bi-antennary and tri-antennary glycans symmetrically extended with three LacNAc repeats.^21,24^ Thus, while two or more extended branches are required for binding, the fucose substitutions on the other branches impact the binding to the extended branch or both. Since tri- and tetra-antennary N-glycans large enough to have multiple extended antennae were of very low relative abundance (**Figure S8**), we did not further investigate these nuances of specificity. In summary, because α2-6-sialylated biantennary N-glycans with one or two extended antennae are the most abundant of the high avidity N-glycans for recent H3 HAs, we propose that they are the primary receptors of recent H3N2 influenza viruses in the human airway.

Our results also suggest that the restricted specificity of recent H3N2 viruses promotes tropism to ciliated cells due to their expression of biosynthetic enzymes that favor the synthesis of α2-6-sialylated extended N-glycans (**Figure 5**, **S19, S20**). Previous reports have shown that earlier human H3N2 and H3N1 viruses bind strongly to ciliated cells of the tracheal and bronchial epithelium, whereas avian viruses bind poorly or not at all to these cells.^12,14,15,55^ On the surface, this seems contradictory to reports using ALI-cultured bronchial epithelial cells, where human H3N2 viruses preferentially infected non-ciliated cells, and avian viruses primarily infect ciliated cells.^16,17,56^ This apparent inconsistency might be explained by the fact that the ALI cultured cells have altered glycosylation, as seen in a comparison of the MS analysis of primary and ALI cultured epithelial cells (**Figure 4B**). Such changes in glycosylation can be manifested in small changes in the expression of glycosyltransferases (**Figure 4C**), which compete for the extension of glycan chains (β3GNT enzymes), or for termination of glycan chains with sialic acid (α2-6 and α2-3 specific sialyltransferases) or fucose (α1-2 and α1-3 specific fucosyltransferases). Further work is needed to understand the extent of glycosylation changes in ALI cultured tracheal/bronchial epithelial cells.

The restricted receptor specificity of H3N2 viruses has persisted for several decades, suggesting that preferred binding to extended human-type receptors has conferred fitness for the function of the HA in the initiation of infection and transmission in humans. One striking finding is that α2-6-sialylated extended N-glycans are more abundantly expressed in tracheal epithelium than nasal epithelium (**Figure 2E**, **Figure S8**). How is this relevant to the initiation of infection by influenza viruses? Influenza virus causes transient tracheobronchitis with virus shedding in the upper airway (mouth, nasal cavity, throat). For aerosol infection, the upper airway would intuitively be the initial site of infection. However, numerous studies suggest this may not be the case. The infectious dose by aerosol is only 2-3x10^3^ virus particles or 3 tissue culture infectious dose (TCID).^57–59^ In contrast, with direct nasal inoculation, the infectious dose is 100-1000 TCID.^59–61^ Following aerosol administration of 3 TCID of influenza virus to volunteers, only 1 TCID was deposited to the nasal cavity, far below the infectious dose by nasal inoculation.^59^ Moreover, the amount of influenza virus for an individual breathing for one hour in a room/airplane during flu season is about 30 TCID, also far below the TCID required for infection by nasal inoculation.^58,62^ Finally, inoculating influenza in the nasal cavity results in infection primarily in the upper respiratory tract, producing milder disease than aerosol administration.^59,60^ For all these reasons, Tellier has suggested that influenza infection by aerosol is likely initiated in the lower airway (e.g., trachea and bronchi).^59^ As we show here, the preferred receptors of H3N2 viruses, α2-6-sialylated extended N-glycans, are far more abundant on tracheal epithelial cells than nasal epithelial cells. A similar argument can be made for H1N1 viruses since they are similarly specific for extended human-type receptors, just not as strict as the H3N2 viruses.^20,25,41^ Thus, we suggest that the restricted specificity of these viruses for extended human-type receptors results in their efficient capture and initiation of infection in the lower airway (e.g. trachea).

## Resource Availability

**Lead contact.** Further information and requests for materials should be directed to the lead contact: James C. Paulson

## Materials availability

Any unique reagents generated in this study are available from the lead contact with a completed materials transfer agreement.

## Data and code availability

Datasets for glycomics analyses generated in this manuscript are provided in supplementary figures. Data for glycan microarray experiments are attached in the supplementary tables according to the established guidelines for the minimum information required for a glycomics experiment (MIRAGE). The code for analyses of nasal RNAseq data and tre-analyses of scRNAseq data and any additional information is available from the lead contact upon request.

## Acknowledgments

This work was partially funded by the National Institute of Allergy and Infectious Diseases, National Institutes of Health, Department of Health and Human Services, under grant AI 114730 (to JCP) and Contract No. 75N93021C00015 (to J.C.P. and I.A.W.), and Grant 082098 from The Wellcome Trust (to S.M.H). We thank the Academia Sinica Common Mass Spectrometry Facilities (grant AS-CFII-111-209) for sulfoglycomic LC-MS/MS data acquisition. We thank Roderick G. Carter for preparing the sections of human tracheal samples, the Microscopy and Histology Core Facilities at La Jolla Institute for preparing the sections of ALI-cultured samples, Dr. Pedro Avila for facilitating the collaboration at the Northwestern Feinberg School of Medicine, and Dr. Kathryn Spencer at Core Microscopy facility at Scripps Research Institute for assistance with confocal microscopy and image analyses.

## Author contributions

C.K., A.A., S.W., A.Bi., and J.C.P. wrote the first draft of the MS, and all authors reviewed and edited the manuscript. S.M.H., J.C.P., and R. S. conceived the project. C.K. conducted HA protein purification, SNA affinity column, glycan microarray analyses, immunofluorescent microscopy imaging and analyses, scRNAseq and bulk RNAseq analyses. A.A. performed the N-glycan and O-glycan MS analyses. C.K., S.W., and C.N. synthesized glycans. S.W. performed glycan ELISA. A.Bi. and A.Bh. collected human airway tissues and isolated epithelial cells. A.Bi. performed ALI-culture of airway epithelial cells and bulk RNAseq experiment. Y.C. and K-H.K. performed sulfoglycomics analyses. R.M. manufactured glycan microarray slides. J.C.P., S.M.H., R.S., A.D., and A.J.T. supervised the research.

## Supplemental figure titles

**Figure S1** MALDI-TOF mass spectra of permethylated N-glycans derived from primary tracheal epithelial cells from human donors.

**Figure S2** MALDI-TOF mass spectra of permethylated N-glycans derived from primary tracheal epithelial cells from human donors after Sial-S digestion.

**Figure S3** MALDI TOF/TOF MS/MS analysis of permethylated N-glycans derived from primary tracheal epithelial cells from human donors.

**Figure S4** MALDI TOF/TOF MS/MS analysis of permethylated N-glycans derived from primary tracheal epithelial cells from human donors before or after Sial-S digestion.

**Figure S5** N-glycan structure library derived from primary tracheal and nasal epithelial cells from human donors.

**Figure S6** MALDI-TOF mass spectra of permethylated N-glycans derived from primary nasal epithelial cells from human donors.

**Figure S7** MALDI-TOF/TOF MS/MS analysis of permethylated N-linked glycans from primary nasal epithelial cells from human donors.

**Figure S8** Human influenza H3 receptor N-linked glycan has low relative abundance on primary tracheal epithelial cells from human donors and almost absent in nasal epithelial cells.

**Figure S9** MALDI-TOF mass spectra of N-linked glycans enriched with an SNA column.

**Figure S10** MALDI-TOF/TOF MS/MS analysis of permethylated N-linked glycans from SNA column.

**Figure S11** O-linked glycans analysis.

**Figure S12** O-linked glycans structures library derived from primary tracheal and nasal epithelial cells from human donors.

**Figure S13** Sulfated glycan analyses.

**Figure S14** Glycan ELISA.

**Figure S15** Glycan array.

**Figure S16** MALDI-TOF mass spectra of permethylated N-linked glycans of primary tracheal epithelial cells from human donors cultured in air-liquid interface.

**Figure S17** MALDI-TOF/TOF MS/MS analysis of permethylated N-linked glycans of primary tracheal epithelial cells from human donors or after culture in air-liquid interface.

**Figure S18** Comparison of the relative abundances of the N-linked glycans of primary tracheal epithelial cells from human donors against the same cells cultured in air-liquid interface.

**Figure S19** IF of human trachea sections. **Figure S20** Additional scRNAseq analyses.

**Table S1** Glycan array compound list.

**Table S2** Glycan array MIRAGE data table.

## STAR Methods text EXPERIMENTAL MODEL

### Collection of human airway epithelial cells

This study was conducted with Institutional Review Board approval at the Northwestern Medicine Allergy-Immunology and Otolaryngology clinic in Chicago, Illinois, USA (STU00202145) for nasal samples and at Northwestern Medicine Lung Transplant Program, Northwestern University (Chicago, IL, USA) for tracheal samples (STU00056197). Informed consent was obtained from each participant.

Nasal epithelial scrapings of the inferior turbinate were collected from healthy subjects using Rhino-Pro curettes (Rhino-pro; Arlington Scientific, Springville, UT) and kept on ice in RPMI for no more than an hour before ready to proceeding with culturing, or harvesting for subsequent glycomics analyses, or lysis for RNA isolation. Every procedure is described in detail in the following sections.

Tracheal epithelium was gently scraped from discarded portions of human trachea from donor transplant lungs with Rhino-Pro and kept on ice in PBS for no longer than 30 minutes. Typically, the very bottom part of the trachea was used, within 4-5 cm of the carina. Curettage ensures high purity of collected samples, containing >90% epithelial cells^63^. Tracheal biopsies were obtained from the same portions of the trachea.

For glycomics analyses, nasal and trachea epithelial scrapings were washed by gentle centrifugation (0.5g x 5 min) with ice-cold PBS three times, then pelleted at 1.0g x 10 min and frozen at -80°C as a pellet of primary epithelial cells for subsequent glycomics analyses. Before pelleting, a 10 ul aliquot was taken for microscopic observation and cell counting. To obtain large numbers of cells per sample required for glycomics analyses (10^6^-10^7^ cells per each sample), it was necessary to pool nasal scrapings from multiple donors (11-18). Care was taken to select blood contamination-free scrapings. In contrast, each tracheal epithelial sample was derived from a single donor. For tracheal epithelial scrapings, pooling was not necessary because substantial portions of the tracheae were available for cell collection.

For cell culture, nasal and tracheal scrapings were slightly dispersed by gentle pipetting and cultured on collagen-coated plates in Bronchial Epithelial Cell Growth Medium, BEGM^TM^ (Lonza CC-3170) following the manufacturer’s recommendations. After 2-3 passages, the cells were either harvested for downstream analyses (RNA isolation or glycomics for submerged cultures) or were transferred to transwell inserts for re-differentiation of an experimental pseudo-stratified airway epithelium at air-liquid interface (ALI), according to a previously published protocol (Lonza),^64^ or frozen in 10% DMSO and 90% FBS and preserved in liquid nitrogen.

### Air-liquid interface culture of human airway cells

We employed the same ALI protocol for cultured primary cells derived from nasal or tracheal scrapings, and for commercially available Normal Broncho-Epithelial Cells (NBEC) from Lonza. To passage cells during an expansion phase we followed the instructions supplied by Lonza for NBEC with a few modifications: we used phosphate-buffered saline (PBS) to rinse the cultures before trypsinization and substituted trypsin with TripLE^TM^ Express. Usually, cells rounded after 3 minutes incubation at 37°C. Cell lifters were used to complete the detachment.

After expansion in BEGM, cells were plated on 12 mm Transwell inserts at 0.4x10^6^ cells per insert and allowed to form a monolayer for up to 72 hours, at which point the medium was removed from the top chamber, exposing the monolayer to air (airlift). To encourage differentiation of ALI cultures into model airway epithelium, retinoic acid, at a final concentration of 50 nM, was added to BEGM. Then cultures were maintained for three weeks at ALI conditions to mature; the medium was replaced three times a week, following the Lonza protocol. Each aliquot of BEGM supplemented with retinoic acid was used within two weeks. We did not use any cells beyond passage four (4).

For microscopy, the ALI cultures were rinsed with PBS twice and fixed in 10% neutral buffered formalin for 2 hours at room temperature. Formalin-fixed ALI-cultured HBEC were embedded in paraffin and sectioned at the Microscopy and Histology Core at the La Jolla Institute for Immunology.

### Cell culture

HEK293F cells (Freestyle 293F, Thermo Scientific) were maintained in FreeStyle™ 293 Expression Medium without supplementation. Cells were cultured in an orbital shaker incubator at 37°C with a humidified atmosphere of 8% CO2, rotating at 125 rpm.

## METHOD DETAILS

### Expression and purification of recombinant HA

Recombinant HA was expressed as previously described^24^. Briefly, HA ectodomain (residues 11-521; H3 numbering) were cloned under a CMV promoter for expression in a mammalian cell culture system using the NEBuilder HiFi DNA Assembly Master Mix (New England Biolabs). The construct also contains an N-terminal CD5 signal peptide for secretion, a C-terminal leucine zipper (GCN4) motif for HA trimerization, and His8-tag in the C-terminal of the construct. Final expression constructs were transfected into HEK293F cells. Typically, 150 mL of 1x10^6^ cells/mL culture was transfected with 150 μg plasmid with 600 μg PEI (polyethylenimine). After 6 days, the cell culture supernatant was collected, and proteins were purified using a 1 mL HisTrapFF column. HAs were eluted in a gradient of Tris-HCl buffer containing 0.5 M (final) imidazole, washed, and concentrated typically to 800 μg/mL for experiments. HAs were stored at 4 °C until used for experiments and experiments were done within 5 days post-purification of the protein.

### Analysis of H3 HA binding to glycans in an ELISA assay

To quantitatively analyze the binding of H3 HA to glycans in an ELISA assay, streptavidin-coated high binding capacity 384-well plates (Pierce) were rinsed with PBS and each well was incubated with 50 μL of a 2.4 μM solution of biotinylated glycans (6SLN1-N, 6SLN2-N, 6SLN3-N, LN3-L) in PBS overnight at 4 °C. The plate was washed with PBS-T (0.05% Tween 20 in PBS) to remove the excess glycan, and each well was incubated with 100 μL of blocking buffer (1% BSA and 0.6 μM desthiobiotin in PBS) at room temperature for 1 hour. The plate was subsequently washed with PBS-T and used without further processing. Purified His-tagged HA trimers were precomplexed with anti-His antibody (Biolegend) and HRP-conjugated anti-mouse IgG (H+L) (Thermo Fisher Scientific) in the ratio 4:2:1 (w/w/w) with 1 % BSA and incubated on ice for 20 minutes. The highest HA concentration was typically 100 μg/mL; samples were then diluted in twofold steps. 50 μL of precomplexed HA was added to each glycan-coated well and incubated at room temperature for 2 hours. The wells were washed with PBS-T completely and 50 μL of TMB substrate was added. The plate was incubated at room temperature for development and then quenched by adding 50 μL of 2 M sulfuric acid. The absorbance at 450 nm was measured with a BioTek Synergy H1 microplate reader and data were analyzed using GraphPad Prism software.

### Processing human trachea and nasal epithelial cells from tissues, and air-liquid interface cells to obtain N- and O-linked glycans

1.5-3×10^6^ cells per sample were collected. All samples were treated as described previously^65^. Briefly, each sample of human epithelial, nasal, human or air-liquid interface culture cells was subjected to sonication in the presence of detergent 3-[(3-Cholamidopropyl)dimethylammonio]-1-propanesulfonate hydrate (CHAPS), reduced in 4 M guanidine-HCl, carboxymethylated, and digested with trypsin (24 h, 37°C). The digested glycoproteins were then purified by Oasis plus short HLB Sep-Pak cartridges. N-linked glycans were released by peptide N-glycosidase F (E.C. 3.5.1.52) digestion (24 h, 37°C) and purified by classic short C18-Sep-Pak. O-linked glycans were released chemically with 400 μL of 55 mg potassium borohydride in 0.1 M KOH (24 h, 45°C). Released N- and O-linked glycans were permethylated using the sodium hydroxide procedure and purified by classic short C18-Sep-Pak cartridges. Permethylated N- and O-linked glycans were eluted at the 50% and 35% acetonitrile fractions respectively.

### Sialidase digestions

Cleavage of α2-3-linked sialic acids from each epithelial human sample was performed with sialidase-S (*Streptococcus pneumoniae*; E.C. 3.2.1.18) digestion as previously described^35,65^. N-linked glycans released by peptide N-glycosidase F were digested with 170 milliunits of the enzyme in 200 μl of 50 mM sodium acetate (24 h, 37 °C, pH 5.5). The digested N-linked glycans were lyophilized and purified on a classic short C18 Sep-Pak, followed by the same permethylation and purification procedures as described above.

### Mass spectrometry

MS and MS/MS data were acquired on a 4800 MALDI-TOF/TOF (Applied Biosystems, Darmstadt, Germany) mass spectrometer. Permethylated glycans were dissolved in 10 μl of methanol, and 1 μl of dissolved sample was premixed with 1 μl of matrix (10 mg/ml 3,4-diaminobenzophenone in 75% (v/v) aqueous acetonitrile), spotted onto a target plate, and dried under vacuum. For the MS/MS studies the collision energy was set to 1 kV, and argon was used as collision gas. The 4700 calibration standard kit, calmix (Applied Biosystems), was used as the external calibrant for the MS mode, and [Glu1] fibrinopeptide B human (Sigma) was used as an external calibrant for the MS/MS mode.

### MS and MS/MS analysis

The MS and MS/MS data were initially processed using Data Explorer 4.9 software (Applied Biosystems). The processed spectra were subjected to manual assignment and annotation with the aid of a glycobioinformatics tool, GlycoWorkBench^66^. The calculation of the relative abundance of each MS panel in the multi-panel spectra of the Supplementary Figures, or partial spectra in the main Figures, was based on the total ion current of each panel (TIC %).

For a more in-depth identification of structures in all the acquired spectra, the MS spectra were additionally subjected to a de-isotoping software^67^, automatically matching molecular ions [M+Na]^+^ with compositions corresponding to N-linked glycan structures (https://github.com/gw110/Glycomics-of-cervicovaginal-fluid-from-women-at-risk-of-preterm-birth.git). A proposed peak was considered for further evaluation based on the following two criteria: the composition matched a possible N-linked glycan structure according to the human N-linked glycan biosynthetic pathway (including blood groups) and the actual isotopic pattern distribution (found from the MS) correlated with the theoretical (calculated from the software) with a value equal or higher than 0.9 (R2>=0.9). The proposed compositions were then manually verified, and a structure was proposed. The de-isotoping was limited to a maximum m/z value of about 5000. This process verified the molecular ions [M+Na]^+^ manually assigned above, and identified new molecular ions of minor abundance or hidden within complex molecular ion clusters.

For more accurate relative quantitation of structures of interest, the relative abundance of each molecular ion [M+Na]^+^ was calculated normalising its de-isotoped intensity to the sum of intensities from all the de-isotoped peaks accepted as positive (expressed as %). This calculation excluded peaks corresponding to high mannose structures (Man_5_GlcNAc_2_ to Man_9_GlcNAc_2_) and structures containing blood-group epitopes (where possible).

All proposed assignments from the manual and automatic identified peaks were based on 12C isotopic composition together with knowledge of the biosynthetic pathways. The proposed structures (**Figure S5**) were then confirmed by data obtained from MS/MS analysis. In total, more than 500 MS/MS were acquired and manually analysed. Due to high spectra complexity, it was often impossible to isolate the molecular ion of interest found within a molecular ion cluster to perform MALDI-TOF/TOF MS/MS analysis. Therefore, the latter was performed on the molecular ion cluster containing the molecular ion of interest.

### Mass spectrometry analysis of sulfated N-linked glycans

Glycans were released and subsequently permethylated following a previously established protocol^68^. Briefly, 2 x 10^6^ airway epithelial cells (TTr02) were washed with PBS and resuspended in 300 µL of TX-100 extraction buffer (1% Triton X-100 in Tris-buffered saline). The supernatant, containing glycoproteins, was collected after sonication and centrifugation at 13,000 rpm for 10 min at 4°C. The glycoproteins were reduced and alkylated with 0.1 M dithiothreitol and 0.5 M iodoacetamide in 25 mM ammonium acetate. Subsequently, they were precipitated with 10% trichloroacetic acid at -20°C for 30 min and washed three times with cold acetone. The glycoproteins were digested with trypsin (1:20) at 37°C overnight, followed by chymotrypsin (1:20) at 37°C for an additional overnight digestion. N-glycans were released by PNGase F digestion at 37°C overnight. The released N-glycans were reduced using 1 M sodium borohydride in 0.05 M sodium hydroxide at 45°C for 16 h. The reaction was quenched with 30% ice-cold acetic acid (AcOH) and desalted using Dowex 50W-X8 beads. The N-glycans were then permethylated using a NaOH/DMSO slurry (50 mg NaOH powder, 200 µL dimethyl sulfoxide, and 200 µL iodomethane) at 4°C for 3 h. The reaction was quenched with 10% ice-cold AcOH and the permethylated N-glycans were fractionated into nonsulfated, monosulfated, and multiply sulfated species using an Oasis® Max solid-phase extraction (SPE) cartridge (Waters) by elution in 95% acetonitrile (ACN), 1 mM ammonium acetate in 80% ACN, and 100 mM ammonium acetate in 60% ACN/20% methanol, respectively. The permethylated N-glycans were further cleaned up using a ZipTip (Millipore) and re-dissolved in 10% ACN/0.1% formic acid (FA) for LC-MS/MS analysis.

The permethylated (sulfated) N-glycans were analyzed using an Ultimate 3000 HPLC system connected to an Orbitrap Fusion Tribrid mass spectrometer (Thermo Fisher Scientific). They were separated on a reversed-phase ReproSil-Pur 120 C18-AQ column (ESI Source Solution) at a flow rate of 500 nL/min at 50°C, with a gradient of 30%–80% solvent B (100% ACN with 0.1% FA) over 59 min. Solvent A was 0.1% FA in water. For data acquisition, an HCD-MS^2^ product– dependent MS^3^ data-dependent method was performed as described^69^. All data were manually annotated.

### Glycan microarray

The glycan microarray was manufactured as described previously^70,71^. Briefly, a library of synthetic glycans containing inkers with a terminal primary amine at the reducing end were used for printing on the N-Hydroxysuccinimide (NHS)-activated slides. The printed slides were blocked with free amine and kept in -20°C until the experiment. The full list of compounds is shown in **Table S1.**

### Analysis of HA binding to Glycan microarray

For analysis of influenza HA binding to the glycan array, recombinant HA trimers (50 μg/mL final) were pre-complexed with the anti-His mouse antibody (Biolegend) and the Alexa488-linked anti-mouse IgG (Invitrogen) at 4/2/1 (w/w/w) ratio for 15 min on ice in 100μL PBS-T. This complex was incubated on the array surface in a humidified chamber for 60 min before washing in PBS-T, PBS, and then water. Slides were dried before being scanned using an Innoscan 1100AL microarray scanner (Innopsys). Analyses were done using Mapix software (Innopsys) and Microsoft Excel (Microsoft). Full descriptions of the microarray experiment and datasets are presented in **Table S1** conforming to the MIRAGE consortium format.

### Bulk RNA seq of native and ALI-cultured human nasal epithelial cells

For bulk RNAseq, RNA was isolated with a QIAgen RNeasy mini kit. Nasal epithelial cells taken directly from tissue curettage (i.e. primary cells), or those grown in either submerged tissue cultures or in air-liquid interface (ALI) cultures, were washed/rinsed with cold PBS before lysis, in RLT lysis buffer supplemented with β-mercaptoethanol as recommended in the QIAgen protocol. RNA integrity (RIN) scores and RNA quantity were measured on an Agilent Bioanalyzer prior to sequencing. Although RIN was always higher than 8, no absolute lower limit of RIN was used as a cutoff for exclusion of samples, but rather individual sample quality was judged by their 28S/18S peak morphologies.

### RNA Sequencing library Preparation

Quality and quantity of RNA obtained as described in the previous section were measured using Agilent 4200 Tapestation using high Sensitivity RNA ScreenTape System (Agilent Technologies).

Briefly, mRNA was isolated from purified 200 ng of total RNA using oligo-dT beads (New England Biolabs, Inc). The NEBNext Ultra™ RNA kit was used for full-length cDNA synthesis and library preparation. Libraries were pooled, denatured and diluted, resulting in a 1.8 pM DNA solution. PhiX control was spiked at 1%. Libraries were sequenced on an Illumina NextSeq 500 instrument (Illumina Inc) using NextSeq 500 High Output reagent kit (Illumina Inc) (1x75 cycles) with a target read depth of approximate 10 million aligned reads per sample.

### scRNAseq and bulk RNA seq analysis

The normalized scRNAseq dataset^43^ was downloaded from UCSC Cell Browser and analyzed using R. Cells assigned as “proximal” positions (trachea) were extracted for the analysis. BulkRNAseq results were normalized and analyzed using DEseq2 package using R^72^.

### Immunofluorescent microscopy imaging and analysis

Slides with Formalin-Fixed Paraffin-Embedded (FFPE) samples of human trachea sections or ALI-cultured cells were deparaffinized in 100% xylene for 5 minutes, twice, then rehydrated by soaking in 100%, 95%, 70%, 50% ethanol-water for 3 minutes each. Slides were washed twice with PBS (5 mins each) then antigen-retrieval was performed in 10 mM citrate buffer (pH 6; 60°C for 2 hours), after which the slide-containing vessel was allowed to cool down at room temperature for 20 mins. Slides were then washed three times with PBS (5 mins each). Samples were permeabilized in PBS containing 0.1% Triton-X100 (Sigma) for 5 mins on ice, then washed three times in PBS (5 minutes each) before blocking with 5% goat serum for 1 hour. Slides were subsequently blocked with avidin/biotin blocking kit (Vector laboratories) following the manufacturer’s instructions. After briefly washing with PBS, slides were incubated with primary antibodies and SNA lectin or pre-complexed recombinant HA-anti-His antibody complex overnight at 4 °C. After washing three times with PBS, secondary antibodies were added and incubated at room temperature for 1 hour. Slides were washed two times with PBS containing 0.35 M NaCl for 5 mins each followed by sequential washing in PBS, PBS containing 0.1% Triton-X100, and then PBS three times, before mounting in Vectashield (Vector laboratories).

The primary antibodies used in this study were as follows: anti-acetyl-α-tubulin antibody (Sigma, 1:2000), anti-mucin-5AC antibody (Abcam, 1:150), anti-FoxJ1 antibody (RnD systems, 1:400), anti-His-tag antibody (Novus Biologicals, 25 μg/mL), biotinylated SNA lectin (Vector laboratories, 1:100). For detection, AlexaFluor-488, 555plus, 597plus, 647-congugated secondary antibodies (Life Technologies) or streptavidin-AlexaFluor-488 (Biolegend) were used.

Immunofluorescence images were obtained using Zeiss LSM780 confocal microscopy. Excitation wavelengths were 405 nm, 488 nm, 561 nm, and 633 nm with spectral scanning for emission range of 409 nm-696 nm (8.9 nm interval). A Plan-Apochromat 63x/1.40 Oil DIC M27 objective was used for image collection. The acquired spectral data were linearly unmixed using individual spectra defined by single-label samples. For the processing and analyses of confocal images, ZEN black (Zeiss), Imaris (Oxford Instruments) and ImageJ/FIJI software (National Institutes of Health) were used. Brightness/contrast and gamma were adjusted for clarity of presentation for figure preparation. Quantitative analyses were done on a z-stack of unprocessed images using Imaris software. HA puncta on the apical surfaces of epithelia were defined using the ‘Spots’ function. The surfaces of the FOXJ1-positive and MUC5AC-positive cells were defined using the ‘Surface’ function with manual refinements. The shortest distances between the HA puncta (Spots) and the cell surfaces were computed using the ‘shortest distance’ function of Imaris software for each punctum, and the difference was calculated using Microsoft Excel.

### Preparation of ^14^C-labeled N-glycans

Radiolabeled N-glycans were prepared from fetuin glycoprotein (Sigma) and purified N-glycans (GlyTech). To prepare radiolabeled fetuin N-glycan, 1 mg fetuin glycoprotein was desialylated with a cocktail of 2.5 μL and 1 μL of α2-3,6,8,9 Neuraminidase A and α2-3,6,8 Neuraminidase (New England Biolabs), respectively, then lyophilized overnight. The lyophilized powder was rehydrated in Ammonium Bicarbonate buffer (pH 8.0) and treated with trypsin (Sigma) for 8 hours then heat inactivated at 100°C for 5 min. Following an overnight lyophilization, digested glycopeptides were resuspended in metal-containing PBS (Gibco) and re-sialylated by hST6Gal1 for α2-6-silalylation or PmST3 for α2-3-sialylation using 1.8 nmol CMP-^14^C-NeuAc (American Radiolabeled Chemicals) at 37°C for 6 hours. Unreacted CMP-^14^C-NeuAc was removed using Amicon Ultra Centrifugal Filters (Sigma). Glycopeptides were then treated with PNGaseF (New England Biolabs) at 37°C overnight. Released ^14^C-labeled N-glycans were collected using Amicon Ultra Centrifugal Filters. Yield of the above reaction was approximately 250 μL with 7000 cpm/5 μL and 3500 cpm/5 μL for α2-6-linked and α2-3-linked samples, respectively.

To prepare a radiolabeled extended N-glycan, the precursor tri-LacNAc N-glycan, of which the reducing end was protected with a Cbz group, was prepared as described previously^70^. Three (3) mg of tri-LacNAc N-glycan was then sialylated with hST6Gal1 and 3eq of CMP-NeuAc, with or without a spike of 4.3 nmol (1x10^6^ cpm) CMP-^14^C-NeuAc. The unlabeled reaction was used to track the reaction and gave a spot-to-spot conversion to fully sialylated glycans, confirmed by LC-MS. Extended sialylated N-glycans were purified using a SepPak C_18_ column, lyophilized, and then kept at 20 mM in water, at approximately 3000 cpm/5 μL.

### SNA column testing with radiolabeled glycans

2 mL of 50% slurry of SNA-agarose (Vector laboratory) was placed on a column and washed with 10 mL ice-cold PBS.

To test the α2-3/α2-6 linkage separation, either 5 μL or a mixture of 2.5 μL each of ^14^C-labeled α2-3-linked and α2-6-linked fetuin-derived N-glycans were dissolved in 1 mL PBS and loaded onto the column. After collecting flow-through, the column was washed with 1 mL fractions of PBS then eluted with 1 mL fractions of PBS containing 400 mM Lactose (Sigma). Fractions were collected and ^14^C radioactivity was measured by Tri-Carb liquid scintillation counter (Perkin Elmer).

To test the retention and elution of α2-6-linked extended glycans, 5 μL of ^14^C-labeled α2-6-sialylated extended glycan was diluted in 1 mL of PBS and loaded onto the SNA column. After collecting flow-through, the column was then washed with 7 x 1 mL fractions of PBS, then eluted with 8 x 1 mL fractions of PBS containing 400 mM Lactose (Sigma), then further washed with PBS containing 500 mM Lactose and 200 mM acetic acid (Sigma). Fractions were collected and ^14^C radioactivity was measured by Tri-Carb liquid scintillation counter (Perkin Elmer).

### Enrichment of α2-6-sialylated tracheal N-glycans using SNA column

Tracheal epithelial cells (10^7^ cells total) were harvested as described above. N-glycans were prepared from the cells as described above, without the final permethylation step. Lyophilized N-glycans were dissolved in 3 mL water, and split into 0.75 mL and 2.25 mL aliquots, for total N-glycan and SNA column analysis, respectively, then lyophilized.

Lyophilized N-glycans were dissolved in 2 mL ice-cold PBS and loaded onto an SNA column prepared as described above. Flow-through and a subsequent 8 mL wash with ice-cold PBS were collected as an SNA-FT/wash fraction. Bound N-glycans were eluted with 10 mL of ice-cold PBS containing 400 mM Lactose and collected as an SNA-elute fraction.

For removal of salt and Lactose, both SNA-FT/wash and SNA-elute fractions were loaded onto HyperSep Hypercarb SPE Cartridges (Fisher Scientific) after conditioning with 10 mL 80% acetonitrile, 5% acetic acid, 15% water followed by 10 mL water. For the SNA-FT/wash fraction, after sample loading, the column was washed with 10 mL water, then 10 mL water containing 5% acetic acid. N-glycans were then eluted with water containing 50% acetonitrile and 5% acetic acid. For the SNA-elute fraction, after sample loading, the column was washed with 10 mL water, followed by 3 mL water containing 3% acetonitrile. N-glycans were eluted in 10 mL water containing 50% acetonitrile and 5% acetic acid. The collected eluates for both SNA-FT/wash and SNA-elute samples were concentrated with a rotary evaporator and then lyophilized. Lyophilized samples were permethylated as described above and subjected to MS analysis.

## QUANTIFICATION AND STATISTICAL ANALYSIS

Statistics were generated and calculated in GraphPad Prism software. All data are presented as the mean ± standard deviation (S.D.) of at least three technical replicates unless otherwise stated. Statistical details (e.g. P-values, sample sizes, analysis type) for individual experiments are listed in Figure legends. Not all statistical comparisons are shown.

## References

1. Krammer, F., Smith, G.J.D., Fouchier, R.A.M., Peiris, M., Kedzierska, K., Doherty, P.C., Palese, P., Shaw, M.L., Treanor, J., Webster, R.G., and Garcia-Sastre, A. (2018). Influenza. Nat Rev Dis Primers 4, 3. 10.1038/s41572-018-0002-y.

2. Thompson, A.J., and Paulson, J.C. (2021). Adaptation of influenza viruses to human airway receptors. J Biol Chem 296, 100017. 10.1074/jbc.REV120.013309.

3. Stevens, J., Blixt, O., Paulson, J.C., and Wilson, I.A. (2006). Glycan microarray technologies: tools to survey host specificity of influenza viruses. Nat Rev Microbiol 4, 857–864. 10.1038/nrmicro1530.

4. Gambaryan, A.S., and Matrosovich, M.N. (2015). What adaptive changes in hemagglutinin and neuraminidase are necessary for emergence of pandemic influenza virus from its avian precursor? Biochemistry (Mosc) 80, 872–880. 10.1134/S000629791507007X.

5. Connor, R.J., Kawaoka, Y., Webster, R.G., and Paulson, J.C. (1994). Receptor specificity in human, avian, and equine H2 and H3 influenza virus isolates. Virology 205, 17–23. 10.1006/viro.1994.1615.

6. Rogers, G.N., Paulson, J.C., Daniels, R.S., Skehel, J.J., Wilson, I.A., and Wiley, D.C. (1983). Single amino acid substitutions in influenza haemagglutinin change receptor binding specificity. Nature 304, 76–78. 10.1038/304076a0.

7. Stevens, J., Blixt, O., Glaser, L., Taubenberger, J.K., Palese, P., Paulson, J.C., and Wilson, I.A. (2006). Glycan microarray analysis of the hemagglutinins from modern and pandemic influenza viruses reveals different receptor specificities. J Mol Biol 355, 1143–1155. 10.1016/j.jmb.2005.11.002.

8. Shinya, K., Ebina, M., Yamada, S., Ono, M., Kasai, N., and Kawaoka, Y. (2006). Avian flu: influenza virus receptors in the human airway. Nature 440, 435–436. 10.1038/440435a.

9. Tumpey, T.M., Maines, T.R., Van Hoeven, N., Glaser, L., Solorzano, A., Pappas, C., Cox, N.J., Swayne, D.E., Palese, P., Katz, J.M., and Garcia-Sastre, A. (2007). A two-amino acid change in the hemagglutinin of the 1918 influenza virus abolishes transmission. Science 315, 655–659. 10.1126/science.1136212.

10. Cox, N.J., Trock, S.C., and Burke, S.A. (2014). Pandemic preparedness and the Influenza Risk Assessment Tool (IRAT). Curr Top Microbiol Immunol 385, 119–136. 10.1007/82_2014_419.

11. Taubenberger, J.K., and Morens, D.M. (2008). The pathology of influenza virus infections. Annu Rev Pathol 3, 499–522. 10.1146/annurev.pathmechdis.3.121806.154316.

12. Kuiken, T., and Taubenberger, J.K. (2008). Pathology of human influenza revisited. Vaccine 26 *Suppl 4*, D59–66. 10.1016/j.vaccine.2008.07.025.

13. Couceiro, J.N., Paulson, J.C., and Baum, L.G. (1993). Influenza virus strains selectively recognize sialyloligosaccharides on human respiratory epithelium; the role of the host cell in selection of hemagglutinin receptor specificity. Virus Res 29, 155–165. 10.1016/0168-1702(93)90056-s.

14. van Riel, D., den Bakker, M.A., Leijten, L.M., Chutinimitkul, S., Munster, V.J., de Wit, E., Rimmelzwaan, G.F., Fouchier, R.A., Osterhaus, A.D., and Kuiken, T. (2010). Seasonal and pandemic human influenza viruses attach better to human upper respiratory tract epithelium than avian influenza viruses. Am J Pathol 176, 1614–1618. 10.2353/ajpath.2010.090949.

15. van Riel, D., Munster, V.J., de Wit, E., Rimmelzwaan, G.F., Fouchier, R.A., Osterhaus, A.D., and Kuiken, T. (2007). Human and avian influenza viruses target different cells in the lower respiratory tract of humans and other mammals. Am J Pathol 171, 1215–1223. 10.2353/ajpath.2007.070248.

16. Matrosovich, M.N., Matrosovich, T.Y., Gray, T., Roberts, N.A., and Klenk, H.D. (2004). Human and avian influenza viruses target different cell types in cultures of human airway epithelium. Proc Natl Acad Sci U S A 101, 4620–4624. 10.1073/pnas.0308001101.

17. Wan, H., and Perez, D.R. (2007). Amino acid 226 in the hemagglutinin of H9N2 influenza viruses determines cell tropism and replication in human airway epithelial cells. J Virol 81, 5181–5191. 10.1128/JVI.02827-06.

18. Liu, M., van Kuppeveld, F.J., de Haan, C.A., and de Vries, E. (2023). Gradual adaptation of animal influenza A viruses to human-type sialic acid receptors. Curr Opin Virol 60, 101314. 10.1016/j.coviro.2023.101314.

19. Yang, H., Carney, P.J., Chang, J.C., Guo, Z., Villanueva, J.M., and Stevens, J. (2015). Structure and receptor binding preferences of recombinant human A(H3N2) virus hemagglutinins. Virology 477, 18–31. 10.1016/j.virol.2014.12.024.

20. Chandrasekaran, A., Srinivasan, A., Raman, R., Viswanathan, K., Raguram, S., Tumpey, T.M., Sasisekharan, V., and Sasisekharan, R. (2008). Glycan topology determines human adaptation of avian H5N1 virus hemagglutinin. Nat Biotechnol 26, 107–113. 10.1038/nbt1375.

21. Peng, W., de Vries, R.P., Grant, O.C., Thompson, A.J., McBride, R., Tsogtbaatar, B., Lee, P.S., Razi, N., Wilson, I.A., Woods, R.J., and Paulson, J.C. (2017). Recent H3N2 Viruses Have Evolved Specificity for Extended, Branched Human-type Receptors, Conferring Potential for Increased Avidity. Cell Host Microbe 21, 23–34. 10.1016/j.chom.2016.11.004.

22. Xu, R., Zhu, X., McBride, R., Nycholat, C.M., Yu, W., Paulson, J.C., and Wilson, I.A. (2012). Functional balance of the hemagglutinin and neuraminidase activities accompanies the emergence of the 2009 H1N1 influenza pandemic. J Virol 86, 9221–9232. 10.1128/JVI.00697-12.

23. Srinivasan, A., Viswanathan, K., Raman, R., Chandrasekaran, A., Raguram, S., Tumpey, T.M., Sasisekharan, V., and Sasisekharan, R. (2008). Quantitative biochemical rationale for differences in transmissibility of 1918 pandemic influenza A viruses. Proc Natl Acad Sci U S A 105, 2800–2805. 10.1073/pnas.0711963105.

24. Thompson, A.J., Wu, N.C., Canales, A., Kikuchi, C., Zhu, X., de Toro, B.F., Canada, F.J., Worth, C., Wang, S., McBride, R., et al. (2024). Evolution of human H3N2 influenza virus receptor specificity has substantially expanded the receptor-binding domain site. Cell Host Microbe 32, 261–275 e264. 10.1016/j.chom.2024.01.003.

25. Broszeit, F., van Beek, R.J., Unione, L., Bestebroer, T.M., Chapla, D., Yang, J.Y., Moremen, K.W., Herfst, S., Fouchier, R.A.M., de Vries, R.P., and Boons, G.J. (2021). Glycan remodeled erythrocytes facilitate antigenic characterization of recent A/H3N2 influenza viruses. Nat Commun 12, 5449. 10.1038/s41467-021-25713-1.

26. Nobusawa, E., Ishihara, H., Morishita, T., Sato, K., and Nakajima, K. (2000). Change in receptor-binding specificity of recent human influenza A viruses (H3N2): a single amino acid change in hemagglutinin altered its recognition of sialyloligosaccharides. Virology 278, 587–596. 10.1006/viro.2000.0679.

27. Mögling, R., Richard, M.J., van der Vliet, S., van Beek, R., Schrauwen, E.J.A., Spronken, M.I., Rimmelzwaan, G.F., and Fouchier, R.A.M. (2017). Neuraminidase-mediated haemagglutination of recent human influenza A(H3N2) viruses is determined by arginine 150 flanking the neuraminidase catalytic site. Journal of General Virology 98, 1274–1281. 10.1099/jgv.0.000809.

28. Chambers, B.S., Li, Y., Hodinka, R.L., and Hensley, S.E. (2014). Recent H3N2 Influenza Virus Clinical Isolates Rapidly Acquire Hemagglutinin or Neuraminidase Mutations When Propagated for Antigenic Analyses. Journal of Virology 88, 10986 LP-10989. 10.1128/JVI.01077-14.

29. Kikuchi, C., Antonopoulos, A., Wang, S., Maemura, T., Karamanska, R., Lee, C., Thompson, A.J., Dell, A., Kawaoka, Y., Haslam, S.M., and Paulson, J.C. (2023). Glyco-engineered MDCK cells display preferred receptors of H3N2 influenza absent in eggs used for vaccines. Nature Communications 14, 6178. 10.1038/s41467-023-41908-0.

30. Powell, H., Liu, H., and Pekosz, A. (2021). Changes in sialic acid binding associated with egg adaptation decrease live attenuated influenza virus replication in human nasal epithelial cell cultures. Vaccine 39, 3225–3235. 10.1016/j.vaccine.2021.04.057.

31. Zost, S.J., Parkhouse, K., Gumina, M.E., Kim, K., Diaz Perez, S., Wilson, P.C., Treanor, J.J., Sant, A.J., Cobey, S., and Hensley, S.E. (2017). Contemporary H3N2 influenza viruses have a glycosylation site that alters binding of antibodies elicited by egg-adapted vaccine strains. Proc Natl Acad Sci U S A 114, 12578–12583. 10.1073/pnas.1712377114.

32. Jia, N., Byrd-Leotis, L., Matsumoto, Y., Gao, C., Wein, A.N., Lobby, J.L., Kohlmeier, J.E., Steinhauer, D.A., and Cummings, R.D. (2020). The Human Lung Glycome Reveals Novel Glycan Ligands for Influenza A Virus. 10, 1–14. 10.1038/s41598-020-62074-z.

33. Walther, T., Karamanska, R., Chan, R.W.Y.Y., Chan, M.C.W.W., Jia, N., Air, G., Hopton, C., Wong, M.P., Dell, A., Peiris, J.S.M., et al. (2013). Glycomic Analysis of Human Respiratory Tract Tissues and Correlation with Influenza Virus Infection. 9, e1003223–e1003223.

34. Jang - Lee, J., North, S.J., Sutton - Smith, M., Goldberg, D., Panico, M., Morris, H., Haslam, S., and Dell, A. (2006). Glycomic Profiling of Cells and Tissues by Mass Spectrometry: Fingerprinting and Sequencing Methodologies. Methods in Enzymology 415, 59–86. 10.1016/S0076-6879(06)15005-3.

35. Bateman, A.C., Karamanska, R., Busch, M.G., Dell, A., Olsen, C.W., and Haslam, S.M. (2010). Glycan analysis and influenza A virus infection of primary swine respiratory epithelial cells: The importance of NeuAcα2-6 glycans. Journal of Biological Chemistry 285, 34016–34026. 10.1074/jbc.M110.115998.

36. Joziasse, D.H., Schiphorst, W.E., Van den Eijnden, D.H., Van Kuik, J.A., Van Halbeek, H., and Vliegenthart, J.F. (1987). Branch specificity of bovine colostrum CMP-sialic acid: Gal beta 1----4GlcNAc-R alpha 2 6-sialyltransferase. Sialylation of bi-, tri-, and tetraantennary oligosaccharides and glycopeptides of the N-acetyllactosamine type. J Biol Chem 262, 2025–2033.

37. Beyer, T.A., Rearick, J.I., Paulson, J.C., Prieels, J.P., Sadler, J.E., and Hill, R.L. (1979). Biosynthesis of mammalian glycoproteins. Glycosylation pathways in the synthesis of the nonreducing terminal sequences. J Biol Chem 254, 12531–12534.

38. Zhao, C., and Pu, J. (2022). Influence of Host Sialic Acid Receptors Structure on the Host Specificity of Influenza Viruses. Viruses 14 10.3390/v14102141.

39. Kikutani, Y., Okamatsu, M., Nishihara, S., Takase-Yoden, S., Hiono, T., de Vries, R.P., McBride, R., Matsuno, K., Kida, H., and Sakoda, Y. (2020). E190V substitution of H6 hemagglutinin is one of key factors for binding to sulfated sialylated glycan receptor and infection to chickens. Microbiol Immunol 64, 304–312. 10.1111/1348-0421.12773.

40. Gambaryan, A.S., Tuzikov, A.B., Pazynina, G.V., Webster, R.G., Matrosovich, M.N., and Bovin, N.V. (2004). H5N1 chicken influenza viruses display a high binding affinity for Neu5Acalpha2-3Galbeta1-4(6-HSO3)GlcNAc-containing receptors. Virology 326, 310–316. 10.1016/j.virol.2004.06.002.

41. Wang, S., Lin, T.-H., Antonopoulos, A., Kikuchi, C., Chapla, D.G., Sanchez, P.V., Moremen, K.W., Haslam, S.M., Wilson, I.A., and Paulson, J.C. (2025). Chemoenzymatic synthesis of N-linked glycan receptors of H1N1 influenza virus on human airway epithelial cells. ChemRxiv. doi:10.26434/chemrxiv-2025-w085x.

42. Wu, G., Grassi, P., Molina, B.G., MacIntyre, D.A., Sykes, L., Bennett, P.R., Dell, A., and Haslam, S.M. (2024). Glycomics of cervicovaginal fluid from women at risk of preterm birth reveals immuno-regulatory epitopes that are hallmarks of cancer and viral glycosylation. Sci Rep 14, 20813. 10.1038/s41598-024-71950-x.

43. Deprez, M., Zaragosi, L.E., Truchi, M., Becavin, C., García, S.R., Arguel, M.J., Plaisant, M., Magnone, V., Lebrigand, K., Abelanet, S., et al. (2020). A single-cell atlas of the human healthy airways. American Journal of Respiratory and Critical Care Medicine 202, 1636–1645. 10.1164/rccm.201911-2199OC.

44. Varki, A., Cummings, R.D., Esko, J.D., Stanley, P., Hart, G.W., Aebi, M., Mohnen, D., Kinoshita, T., Packer, N.H., Prestegard, J.H., et al. (2022). Essentials of Glycobiology (Cold Spring Harbor Laboratory Press). 10.1101/9781621824213.

45. Rillahan, C.D., Antonopoulos, A., Lefort, C.T., Sonon, R., Azadi, P., Ley, K., Dell, A., Haslam, S.M., and Paulson, J.C. (2012). Global metabolic inhibitors of sialyl- and fucosyltransferases remodel the glycome. Nat Chem Biol 8, 661–668. 10.1038/nchembio.999.

46. Ishida, H., Togayachi, A., Sakai, T., Iwai, T., Hiruma, T., Sato, T., Okubo, R., Inaba, N., Kudo, T., Gotoh, M., et al. (2005). A novel beta1,3-N-acetylglucosaminyltransferase (beta3Gn-T8), which synthesizes poly-N-acetyllactosamine, is dramatically upregulated in colon cancer. FEBS Lett 579, 71–78. 10.1016/j.febslet.2004.11.037.

47. Seko, A., and Yamashita, K. (2004). beta1,3-N-Acetylglucosaminyltransferase-7 (beta3Gn-T7) acts efficiently on keratan sulfate-related glycans. FEBS Lett 556, 216–220. 10.1016/s0014-5793(03)01440-6.

48. Togayachi, A. (2021). Enzyme assay of beta1,3-glycosyltransferase family. In Glycoscience Protocols (GlycoPODv2), S. Nishihara, K. Angata, K.F. Aoki-Kinoshita, and J. Hirabayashi, eds.

49. Fang, R., Min Jou, W., Huylebroeck, D., Devos, R., and Fiers, W. (1981). Complete structure of A/duck/Ukraine/63 influenza hemagglutinin gene: animal virus as progenitor of human H3 Hong Kong 1968 influenza hemagglutinin. Cell 25, 315–323. 10.1016/0092-8674(81)90049-0.

50. Garten, R.J., Davis, C.T., Russell, C.A., Shu, B., Lindstrom, S., Balish, A., Sessions, W.M., Xu, X., Skepner, E., Deyde, V., et al. (2009). Antigenic and genetic characteristics of swine-origin 2009 A(H1N1) influenza viruses circulating in humans. Science 325, 197–201. 10.1126/science.1176225.

51. Schafer, J.R., Kawaoka, Y., Bean, W.J., Suss, J., Senne, D., and Webster, R.G. (1993). Origin of the pandemic 1957 H2 influenza A virus and the persistence of its possible progenitors in the avian reservoir. Virology 194, 781–788. 10.1006/viro.1993.1319.

52. Trifonov, V., Khiabanian, H., and Rabadan, R. (2009). Geographic dependence, surveillance, and origins of the 2009 influenza A (H1N1) virus. N Engl J Med 361, 115–119. 10.1056/NEJMp0904572.

53. Worobey, M., Han, G.Z., and Rambaut, A. (2014). A synchronized global sweep of the internal genes of modern avian influenza virus. Nature 508, 254–257. 10.1038/nature13016.

54. Unione, L., Ammerlaan, A.N.A., Bosman, G.P., Uslu, E., Liang, R., Broszeit, F., van der Woude, R., Liu, Y., Ma, S., Liu, L., et al. (2024). Probing altered receptor specificities of antigenically drifting human H3N2 viruses by chemoenzymatic synthesis, NMR, and modeling. Nat Commun 15, 2979. 10.1038/s41467-024-47344-y.

55. Baum, L.G., and Paulson, J.C. (1990). Sialyloligosaccharides of the respiratory epithelium in the selection of human influenza virus receptor specificity. Acta Histochem Suppl 40, 35–38.

56. Ibricevic, A., Pekosz, A., Walter, M.J., Newby, C., Battaile, J.T., Brown, E.G., Holtzman, M.J., and Brody, S.L. (2006). Influenza virus receptor specificity and cell tropism in mouse and human airway epithelial cells. J Virol 80, 7469–7480. 10.1128/JVI.02677-05.

57. Alford, R.H., Kasel, J.A., Gerone, P.J., and Knight, V. (1966). Human influenza resulting from aerosol inhalation. Proc Soc Exp Biol Med 122, 800–804. 10.3181/00379727-122-31255.

58. Nikitin, N., Petrova, E., Trifonova, E., and Karpova, O. (2014). Influenza virus aerosols in the air and their infectiousness. Adv Virol 2014, 859090. 10.1155/2014/859090.

59. Tellier, R. (2006). Review of aerosol transmission of influenza A virus. Emerg Infect Dis 12, 1657–1662. 10.3201/eid1211.060426.

60. Little, J.W., Douglas, R.G., Jr., Hall, W.J., and Roth, F.K. (1979). Attenuated influenza produced by experimental intranasal inoculation. J Med Virol 3, 177–188. 10.1002/jmv.1890030303.

61. Hayden, F.G., Treanor, J.J., Betts, R.F., Lobo, M., Esinhart, J.D., and Hussey, E.K. (1996). Safety and efficacy of the neuraminidase inhibitor GG167 in experimental human influenza. JAMA 275, 295–299.

62. Yang, W., Elankumaran, S., and Marr, L.C. (2011). Concentrations and size distributions of airborne influenza A viruses measured indoors at a health centre, a day-care centre and on aeroplanes. J R Soc Interface 8, 1176–1184. 10.1098/rsif.2010.0686.

63. Massey, C.J., Diaz Del Valle, F., Abuzeid, W.M., Levy, J.M., Mueller, S., Levine, C.G., Smith, S.S., Bleier, B.S., and Ramakrishnan, V.R. (2020). Sample collection for laboratory-based study of the nasal airway and sinuses: a research compendium. International Forum of Allergy and Rhinology 10, 303–313. 10.1002/alr.22510.

64. Damian, S., Klarmann, G., Smithhisler, M., and Lonza Walkersville, I. (2010). B-ALI Bronchial Air Liquid Interface Media Kit, a Guaranteed 3D in vitro Model for Respiratory Research. Lonza Research Notes, 13–16.

65. Kikuchi, C., Antonopoulos, A., Wang, S., Maemura, T., Karamanska, R., Lee, C., Thompson, A.J., Dell, A., Kawaoka, Y., Haslam, S.M., and Paulson, J.C. (2023). Glyco-engineered MDCK cells display preferred receptors of H3N2 influenza absent in eggs used for vaccines. Nature Communications 14, 6178. 10.1038/s41467-023-41908-0.

66. Ceroni, A., Maass, K., Geyer, H., Geyer, R., Dell, A., and Haslam, S.M. (2008). GlycoWorkbench: A tool for the computer-assisted annotation of mass spectra of glycans. Journal of Proteome Research 7, 1650–1659.

67. Wu, G., Grassi, P., Molina, B.G., MacIntyre, D.A., Sykes, L., Bennett, P.R., Dell, A., and Haslam, S.M. (2024). Glycomics of cervicovaginal fluid from women at risk of preterm birth reveals immuno-regulatory epitopes that are hallmarks of cancer and viral glycosylation. Scientific Reports 14, 20813. 10.1038/s41598-024-71950-x.

68. Yu, S.Y., Snovida, S., and Khoo, K.H. (2020). Permethylation and Microfractionation of Sulfated Glycans for MS Analysis. Bio Protoc 10, e3617. 10.21769/BioProtoc.3617.

69. Hsiao, C.T., Wang, P.W., Chang, H.C., Chen, Y.Y., Wang, S.H., Chern, Y., and Khoo, K.H. (2017). Advancing a High Throughput Glycotope-centric Glycomics Workflow Based on nanoLC-MS(2)-product Dependent-MS(3) Analysis of Permethylated Glycans. Mol Cell Proteomics 16, 2268–2280. 10.1074/mcp.TIR117.000156.

70. Peng, W., de Vries, R.P., Grant, O.C., Thompson, A.J., McBride, R., Tsogtbaatar, B., Lee, P.S., Razi, N., Wilson, I.A., Woods, R.J., and Paulson, J.C. (2017). Recent H3N2 Viruses Have Evolved Specificity for Extended, Branched Human-type Receptors, Conferring Potential for Increased Avidity. Cell host & microbe 21, 23–34. 10.1016/j.chom.2016.11.004.

71. Blixt, O., Head, S., Mondala, T., Scanlan, C., Huflejt, M.E., Alvarez, R., Bryan, M.C., Fazio, F., Calarese, D., Stevens, J., et al. (2004). Printed covalent glycan array for ligand profiling of diverse glycan binding proteins. Proceedings of the National Academy of Sciences of the United States of America 101, 17033–17038. 10.1073/pnas.0407902101.

72. Love, M.I., Huber, W., and Anders, S. (2014). Moderated estimation of fold change and dispersion for RNA-seq data with DESeq2. Genome Biology 15, 1–21. 10.1186/s13059-014-0550-8.

