## Supplemental Figures for "H3N2 influenza virus tropism shifts to glycan receptors on tracheal ciliated cells"

#### Supplemental figure titles

**Figure S1** MALDI-TOF mass spectra of permethylated N-glycans derived from primary tracheal epithelial cells from human donors.

**Figure S2** MALDI-TOF mass spectra of permethylated N-glycans derived from primary tracheal epithelial cells from human donors after Sial-S digestion.

**Figure S3** MALDI TOF/TOF MS/MS analysis of permethylated N-glycans derived from primary tracheal epithelial cells from human donors.

**Figure S4** MALDI TOF/TOF MS/MS analysis of permethylated N-glycans derived from primary tracheal epithelial cells from human donors before or after Sial-S digestion.

**Figure S5** N-glycan structure library derived from primary tracheal and nasal epithelial cells from human donors.

**Figure S6** MALDI-TOF mass spectra of permethylated N-glycans derived from primary nasal epithelial cells from human donors.

**Figure S7** MALDI-TOF/TOF MS/MS analysis of permethylated N-linked glycans from primary nasal epithelial cells from human donors.

**Figure S8** Human influenza H3 receptor N-linked glycan has low relative abundance on primary tracheal epithelial cells from human donors and almost absent in nasal epithelial cells.

**Figure S9** MALDI-TOF mass spectra of N-linked glycans enriched with an SNA column.

**Figure S10** MALDI-TOF/TOF MS/MS analysis of permethylated N-linked glycans from SNA column.

**Figure S11** O-linked glycans analysis.

**Figure S12** O-linked glycans structures library derived from primary tracheal and nasal epithelial cells from human donors.

**Figure S13** Sulfated glycan analyses.

**Figure S14** Glycan ELISA.

**Figure S15** Glycan array.

**Figure S16** MALDI-TOF mass spectra of permethylated N-linked glycans of primary tracheal epithelial cells from human donors cultured in air-liquid interface.

**Figure S17** MALDI-TOF/TOF MS/MS analysis of permethylated N-linked glycans of primary tracheal epithelial cells from human donors or after culture in air-liquid interface.

**Figure S18** Comparison of the relative abundances of the N-linked glycans of primary tracheal epithelial cells from human donors against the same cells cultured in air-liquid interface.

**Figure S19** IF of human trachea sections.

**Figure S20** Additional scRNAseq analyses.

#### Figure S1

**A**

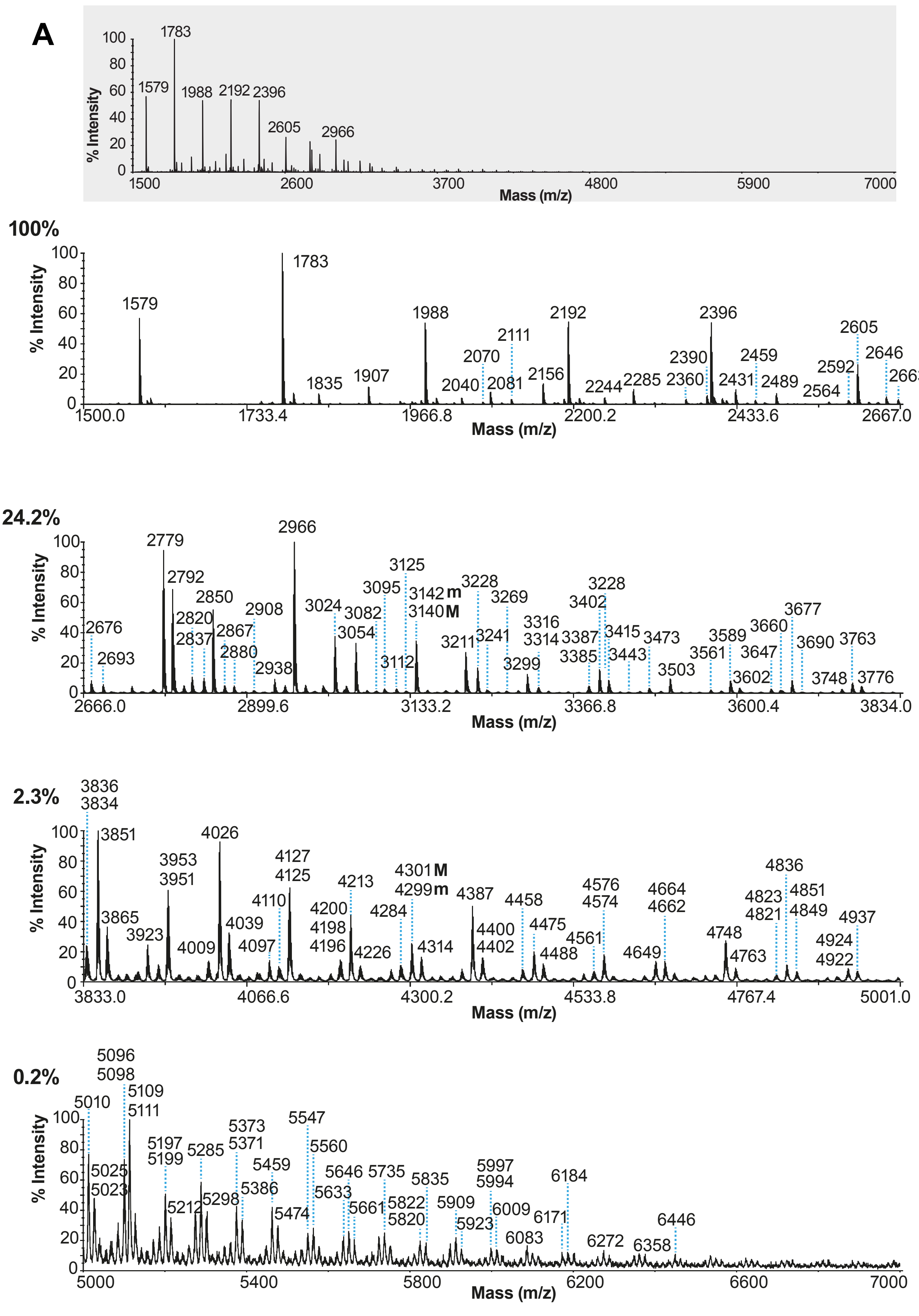

% Intensity
Mass (m/z)
1783
1579
1988
2192
2396
2605
2966
100%
1579
1783
1835
1907
1988
2070
2040
2081
2111
2156
2192
2244
2285
2360
2390
2396
2431
2459
2489
2564
2592
2605
2646
2663
24.2%
2676
2693
2779
2792
2820
2837
2850
2867
2880
2908
2938
2966
3024
3054
3082
3095
3112
3125
3142 m
3140 M
3211
3228
3241
3269
3299
3316
3314
3387
3385
3402
3228
3415
3443
3473
3503
3561
3589
3602
3647
3660
3677
3690
3748
3763
3776
2.3%
3836
3834
3851
3865
3923
3953
3951
4009
4026
4039
4097
4110
4127
4125
4200
4198
4196
4213
4226
4284
4301 M
4299 m
4314
4387
4400
4402
4458
4475
4488
4561
4576
4574
4649
4664
4662
4748
4763
4823
4821
4836
4851
4849
4924
4922
4937
0.2%
5010
5025
5023
5096
5098
5109
5111
5197
5199
5212
5285
5298
5373
5371
5386
5459
5474
5547
5560
5633
5646
5661
5735
5822
5820
5835
5909
5923
5997
5994
6009
6083
6171
6184
6272
6358
6446

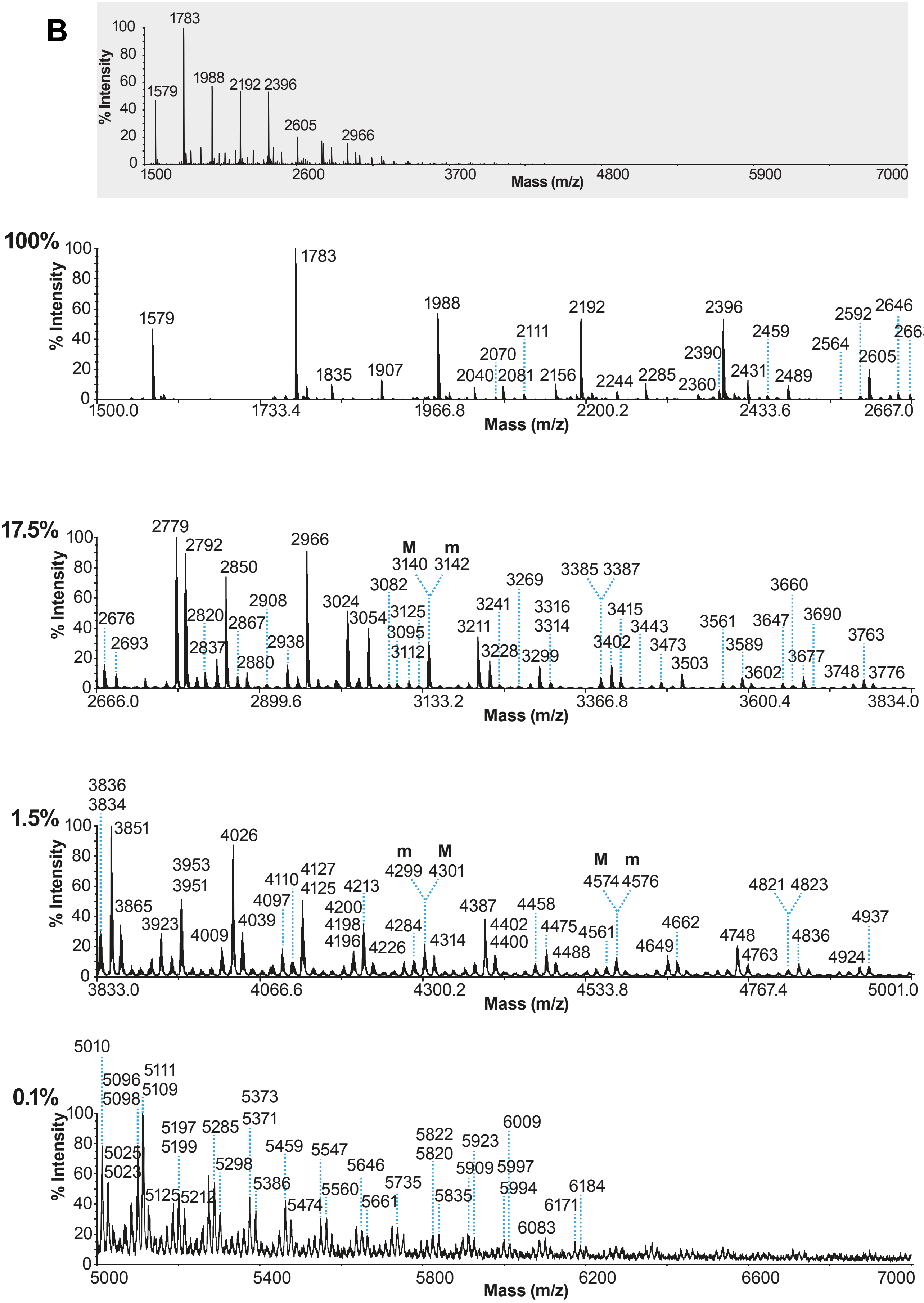
B
% Intensity
Mass (m/z)
100%
17.5%
1.5%
0.1%
M
m

C

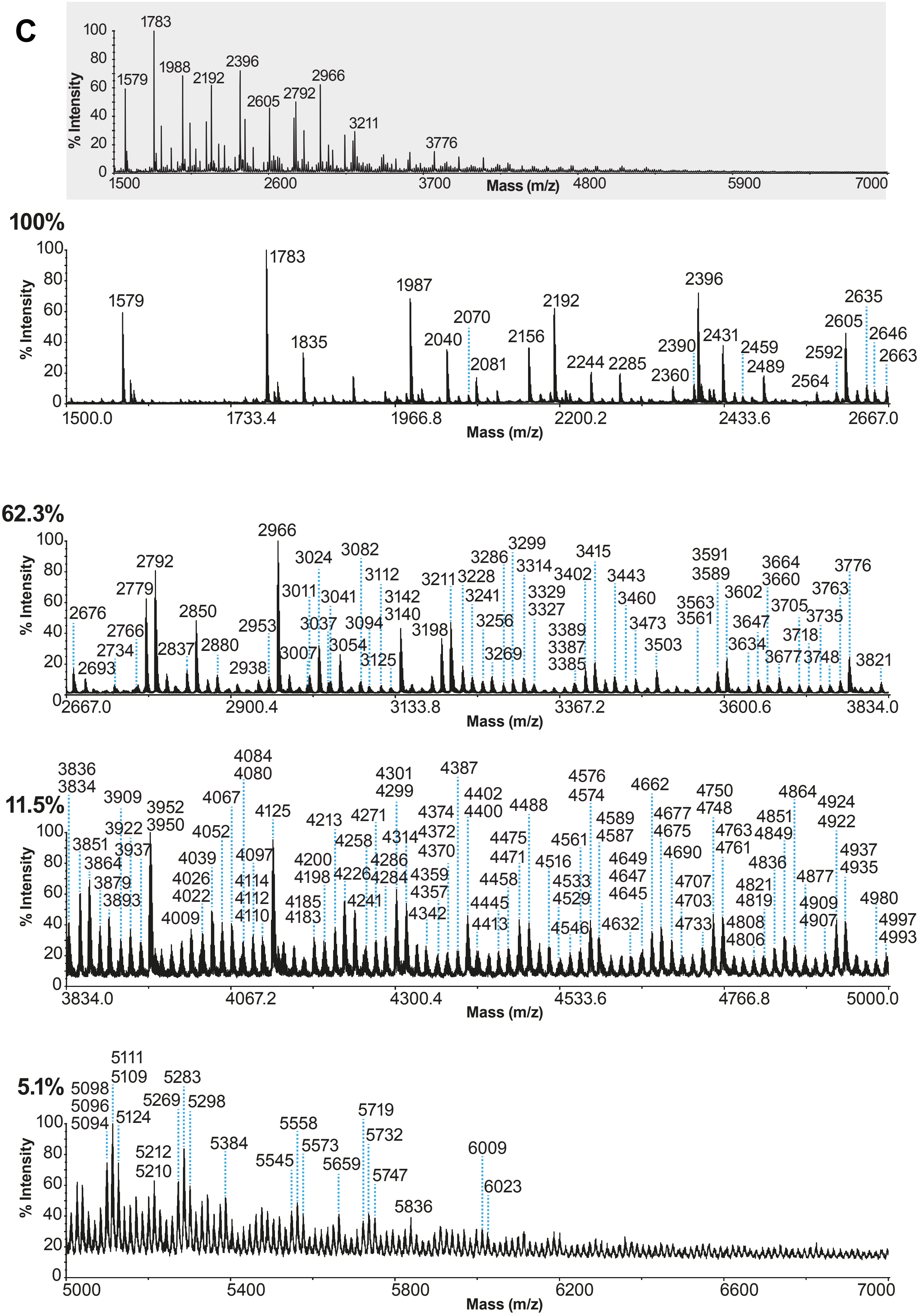

% Intensity
Mass (m/z)
1500 2600 3700 4800 5900 7000
1579 1783 1988 2192 2396 2605 2792 2966 3211 3776
100%
1500.0 1733.4 1966.8 2200.2 2433.6 2667.0
1579 1783 1835 1987 2040 2070 2081 2156 2192 2244 2285 2360 2390 2396 2431 2459 2489 2564 2592 2605 2635 2646 2663
62.3%
2667.0 2900.4 3133.8 3367.2 3600.6 3834.0
2676 2693 2734 2766 2779 2792 2837 2850 2880 2938 2953 2966 3007 3011 3024 3037 3041 3054 3082 3094 3112 3125 3140 3142 3198 3211 3228 3241 3256 3269 3286 3299 3314 3327 3329 3385 3387 3389 3402 3415 3443 3460 3473 3503 3561 3563 3589 3591 3602 3634 3647 3660 3664 3677 3705 3718 3735 3748 3763 3776 3821
11.5%
3834.0 4067.2 4300.4 4533.6 4766.8 5000.0
3834 3836 3851 3864 3879 3893 3909 3922 3937 3950 3952 4009 4022 4026 4039 4052 4067 4080 4084 4097 4110 4112 4114 4125 4183 4185 4198 4200 4213 4226 4241 4258 4271 4284 4286 4299 4301 4314 4342 4357 4359 4370 4372 4374 4387 4400 4402 4413 4445 4458 4471 4475 4488 4516 4529 4533 4546 4561 4574 4576 4587 4589 4632 4645 4647 4649 4662 4675 4677 4690 4703 4707 4733 4748 4750 4761 4763 4806 4808 4819 4821 4836 4849 4851 4864 4877 4907 4909 4922 4924 4935 4937 4980 4993 4997
5.1%
5000 5400 5800 6200 6600 7000
5094 5096 5098 5109 5111 5124 5210 5212 5269 5283 5298 5384 5545 5558 5573 5659 5719 5732 5747 5836 6009 6023

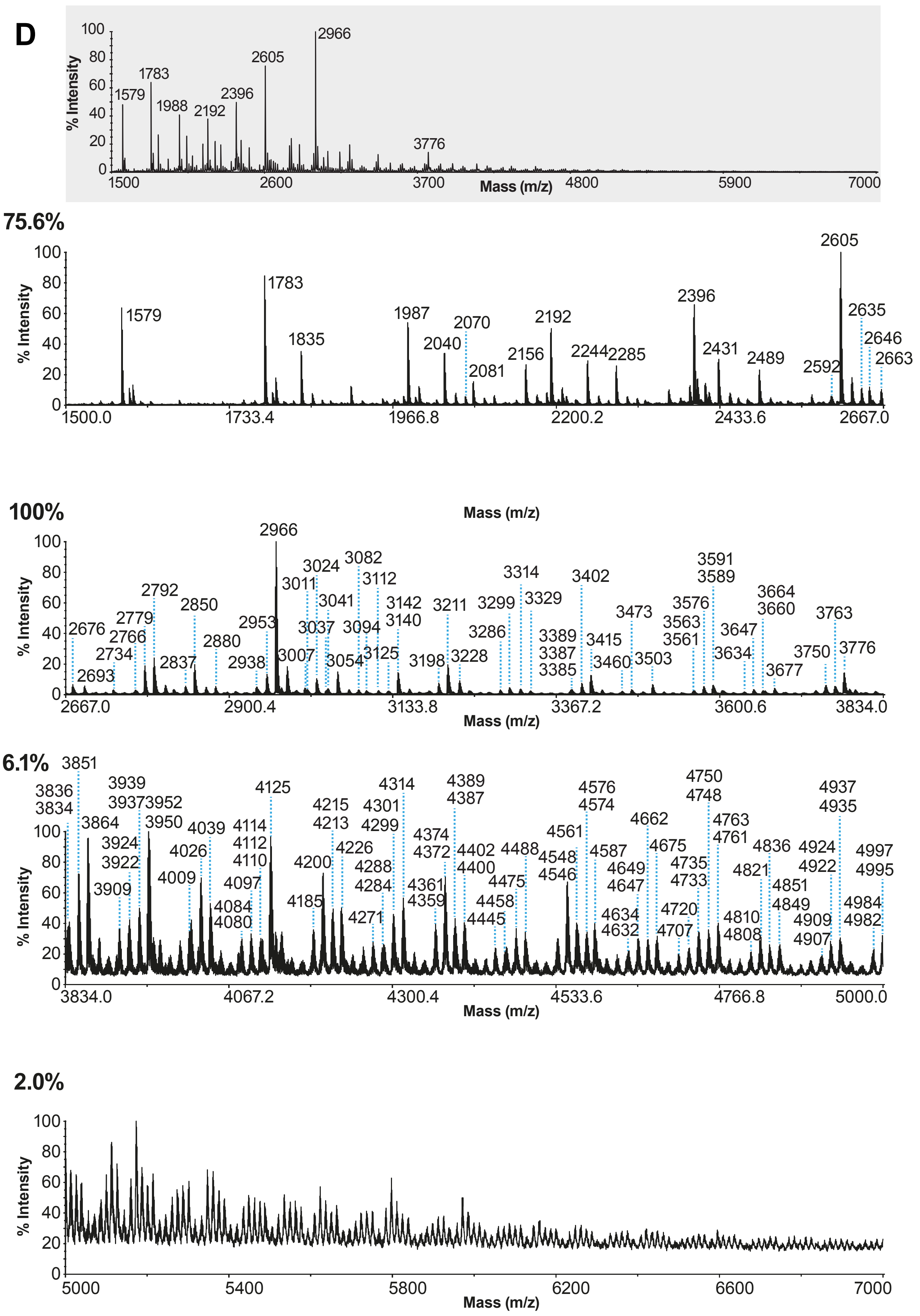
D
% Intensity
Mass (m/z)
2966
2605
1783
1579
1988
2192
2396
3776
1500
2600
3700
4800
5900
7000
75.6%
1579
1783
1835
1987
2040
2070
2081
2156
2192
2244
2285
2396
2431
2489
2592
2605
2635
2646
2663
1500.0
1733.4
1966.8
2200.2
2433.6
2667.0
100%
2676
2693
2734
2766
2779
2792
2837
2850
2880
2938
2953
2966
3007
3011
3024
3037
3041
3054
3082
3094
3112
3125
3140
3142
3198
3211
3228
3286
3299
3314
3329
3385
3387
3389
3402
3415
3460
3473
3503
3561
3563
3576
3589
3591
3634
3647
3660
3664
3677
3750
3763
3776
2667.0
2900.4
3133.8
3367.2
3600.6
3834.0
6.1%
3834
3836
3851
3864
3909
3922
3924
3937
3939
3950
3952
4009
4026
4039
4080
4084
4097
4110
4112
4114
4125
4185
4200
4213
4215
4226
4271
4284
4288
4299
4301
4314
4359
4361
4372
4374
4387
4389
4400
4402
4445
4458
4475
4488
4546
4548
4561
4574
4576
4587
4632
4634
4647
4649
4662
4675
4707
4720
4733
4735
4748
4750
4761
4763
4808
4810
4821
4836
4849
4851
4907
4909
4922
4924
4935
4937
4982
4984
4995
4997
3834.0
4067.2
4300.4
4533.6
4766.8
5000.0
2.0%
5000
5400
5800
6200
6600
7000

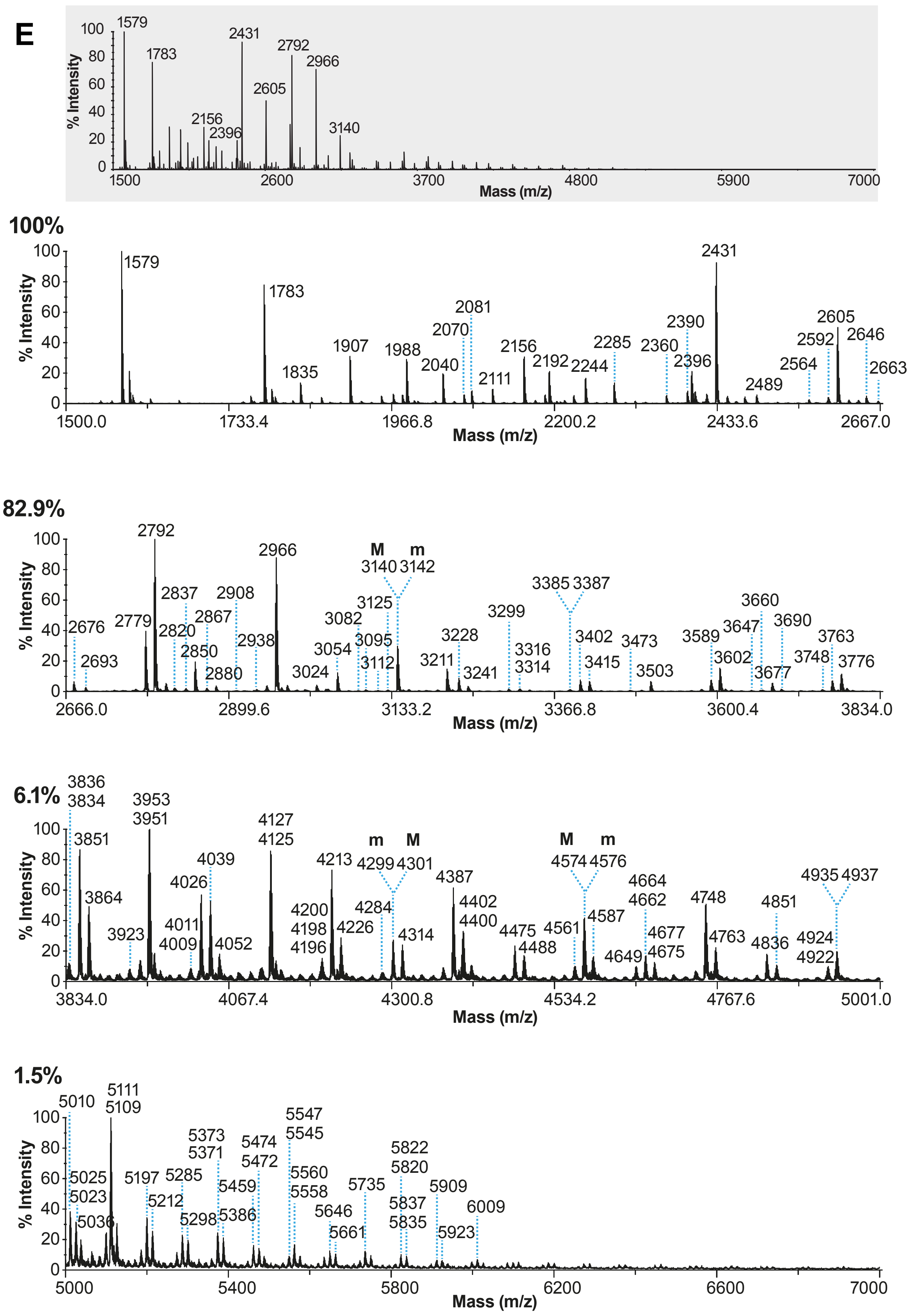
E
% Intensity
Mass (m/z)
1579
1783
2156
2396
2431
2605
2792
2966
3140
1500
2600
3700
4800
5900
7000
100%
1579
1783
1835
1907
1988
2040
2070
2081
2111
2156
2192
2244
2285
2360
2390
2396
2431
2489
2564
2592
2605
2646
2663
1500.0
1733.4
1966.8
2200.2
2433.6
2667.0
82.9%
2676
2693
2779
2792
2820
2837
2850
2867
2880
2908
2938
2966
3024
3054
3082
3095
3112
3125
M m
3140 3142
3211
3228
3241
3299
3316
3314
3385 3387
3402
3415
3473
3503
3589
3602
3647
3660
3677
3690
3748
3763
3776
2666.0
2899.6
3133.2
3366.8
3600.4
3834.0
6.1%
3836
3834
3851
3864
3923
3953
3951
4011
4009
4026
4039
4052
4127
4125
4200
4198
4196
4213
4226
4284
m M
4299 4301
4314
4387
4402
4400
4475
4488
4561
M m
4574 4576
4587
4649
4664
4662
4677
4675
4748
4763
4836
4851
4924
4922
4935 4937
3834.0
4067.4
4300.8
4534.2
4767.6
5001.0
1.5%
5010
5025
5023
5036
5111
5109
5197
5212
5285
5298
5373
5371
5386
5459
5474
5472
5547
5545
5560
5558
5646
5661
5735
5822
5820
5837
5835
5909
5923
6009
5000
5400
5800
6200
6600
7000
0
20
40
60
80
100

F

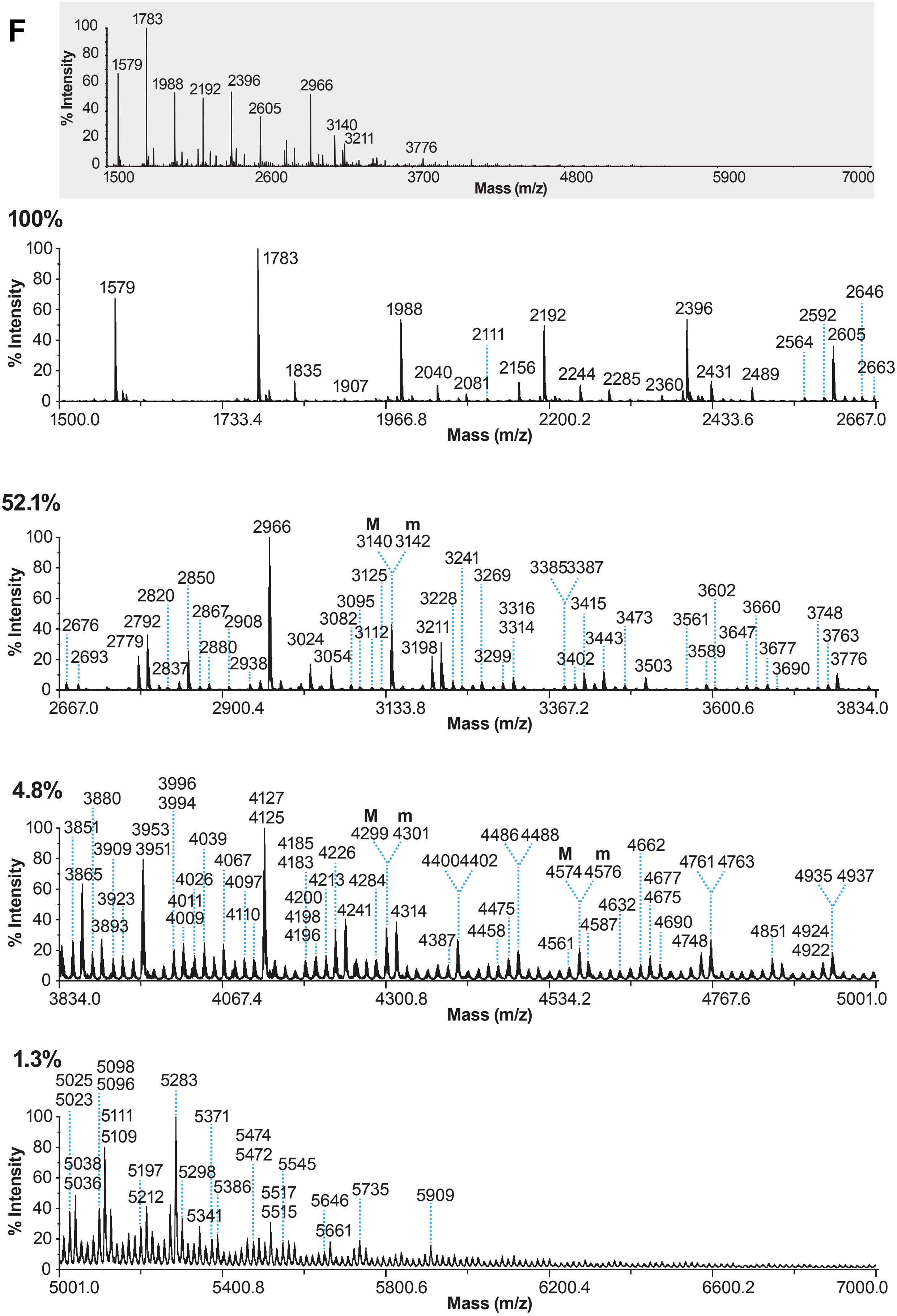
% Intensity
Mass (m/z)
1500 2600 3700 4800 5900 7000
1579 1783 1988 2192 2396 2605 2966 3140 3211 3776
100%
1579 1783 1835 1907 1988 2040 2081 2111 2156 2192 2244 2285 2360 2396 2431 2489 2564 2592 2605 2646 2663
1500.0 1733.4 1966.8 2200.2 2433.6 2667.0
52.1%
M m
2676 2693 2779 2792 2820 2837 2850 2867 2880 2908 2938 2966 3024 3054 3082 3095 3112 3125 3140 3142 3198 3211 3228 3241 3269 3299 3316 3314 3385 3387 3402 3415 3443 3473 3503 3561 3589 3602 3647 3660 3677 3690 3748 3763 3776
2667.0 2900.4 3133.8 3367.2 3600.6 3834.0
4.8%
M m
3851 3865 3880 3893 3909 3923 3953 3951 3996 3994 4011 4009 4026 4039 4067 4097 4110 4127 4125 4185 4183 4200 4198 4196 4213 4226 4241 4284 4299 4301 4314 4387 4400 4402 4458 4475 4486 4488 4561 4574 4576 4587 4632 4662 4677 4675 4690 4748 4761 4763 4851 4924 4922 4935 4937
3834.0 4067.4 4300.8 4534.2 4767.6 5001.0
1.3%
5025 5023 5038 5036 5098 5096 5111 5109 5197 5212 5283 5298 5341 5371 5386 5474 5472 5517 5515 5545 5646 5661 5735 5909
5001.0 5400.8 5800.6 6200.4 6600.2 7000.0
0 20 40 60 80 100

G

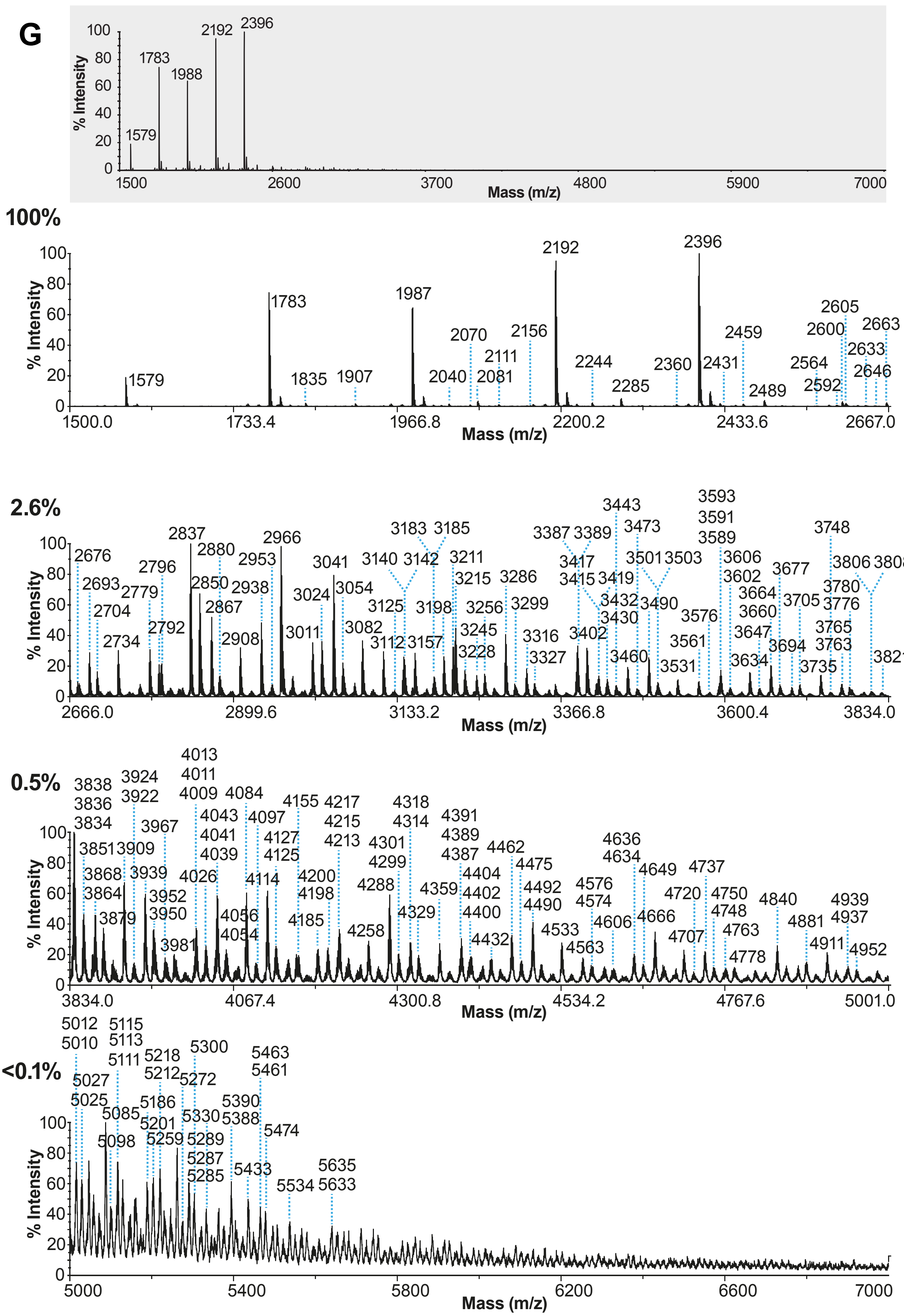
100
80
60
40
20
0
% Intensity
1579
1783
1988
2192
2396
1500
2600
3700
4800
5900
7000
Mass (m/z)
100%
% Intensity
1579
1783
1835
1907
1987
2040
2070
2081
2111
2156
2192
2244
2285
2360
2396
2431
2459
2489
2564
2592
2600
2605
2633
2646
2663
1500.0
1733.4
1966.8
2200.2
2433.6
2667.0
Mass (m/z)
2.6%
% Intensity
2676 2693 2704 2734 2779 2792 2796 2837 2850 2867 2880 2908 2938 2953 2966
3011 3024 3041 3054 3082 3112 3125 3140 3142 3157 3183 3185 3198 3211 3215 3228 3245 3256 3286 3299
3316 3327 3387 3389 3402 3415 3417 3419 3430 3432 3443 3460 3473 3490 3501 3503 3531 3561 3576 3589 3591 3593
3602 3606 3634 3647 3660 3664 3677 3694 3705 3735 3748 3763 3765 3776 3780 3806 3808 3821
2666.0
2899.6
3133.2
3366.8
3600.4
3834.0
Mass (m/z)
0.5%
% Intensity
3834 3836 3838 3851 3864 3868 3879 3909 3922 3924 3939 3950 3952 3967 3981
4009 4011 4013 4026 4039 4041 4043 4054 4056 4084 4097 4114 4125 4127 4155 4185 4198 4200 4213 4215 4217 4258 4288
4299 4301 4314 4318 4329 4359 4387 4389 4391 4400 4402 4404 4432 4462 4475 4490 4492 4533 4563 4574 4576
4606 4634 4636 4649 4666 4707 4720 4737 4748 4750 4763 4778 4840 4881 4911 4937 4939 4952
3834.0
4067.4
4300.8
4534.2
4767.6
5001.0
Mass (m/z)
<0.1%
% Intensity
5010 5012 5025 5027 5085 5098 5111 5113 5115 5186 5201 5212 5218 5259 5272 5285 5287 5289 5300 5330
5388 5390 5433 5461 5463 5474 5534 5633 5635
5000
5400
5800
6200
6600
7000
Mass (m/z)

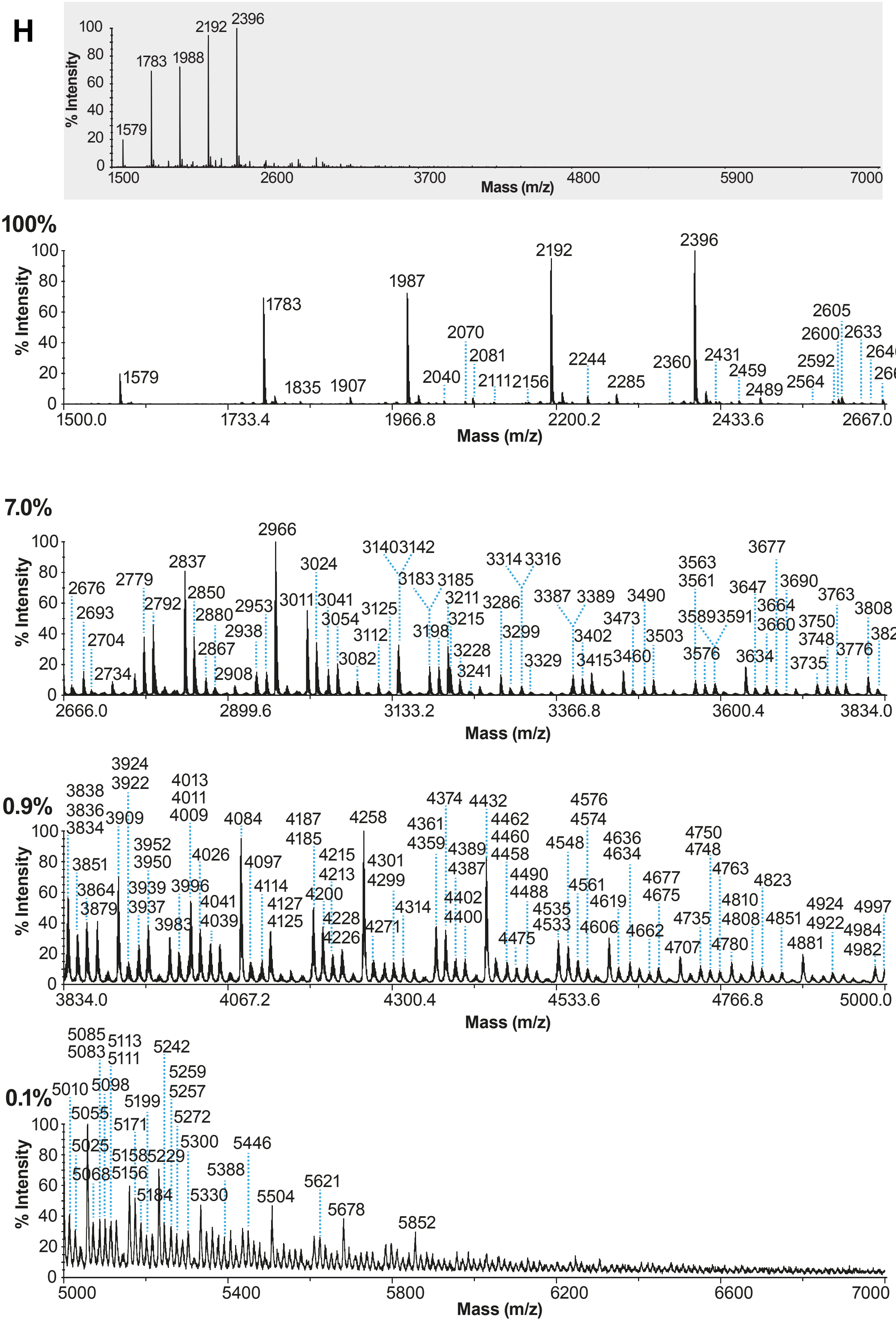
H
% Intensity
Mass (m/z)
1579
1783
1988
2192
2396
100%
1579
1783
1835
1907
1987
2040
2070
2081
2111
2156
2192
2244
2285
2360
2396
2431
2459
2489
2564
2592
2600
2605
2633
2646
2663
7.0%
2676
2693
2704
2734
2779
2792
2837
2850
2867
2880
2908
2938
2953
2966
3011
3024
3041
3054
3082
3112
3125
3140
3142
3183
3185
3198
3211
3215
3228
3241
3286
3299
3314
3316
3329
3387
3389
3402
3415
3460
3473
3490
3503
3561
3563
3576
3589
3591
3634
3647
3660
3664
3677
3690
3735
3748
3750
3763
3776
3808
3821
0.9%
3834
3836
3838
3851
3864
3879
3909
3922
3924
3937
3939
3950
3952
3983
3996
4009
4011
4013
4026
4039
4041
4084
4097
4114
4125
4127
4185
4187
4200
4213
4215
4226
4228
4258
4271
4299
4301
4314
4359
4361
4374
4387
4389
4400
4402
4432
4458
4460
4462
4475
4488
4490
4533
4535
4548
4561
4574
4576
4606
4619
4634
4636
4662
4675
4677
4707
4735
4748
4750
4763
4780
4808
4810
4823
4851
4881
4922
4924
4982
4984
4997
0.1%
5010
5025
5055
5068
5083
5085
5098
5111
5113
5156
5158
5171
5184
5199
5229
5242
5257
5259
5272
5300
5330
5388
5446
5504
5621
5678
5852

I

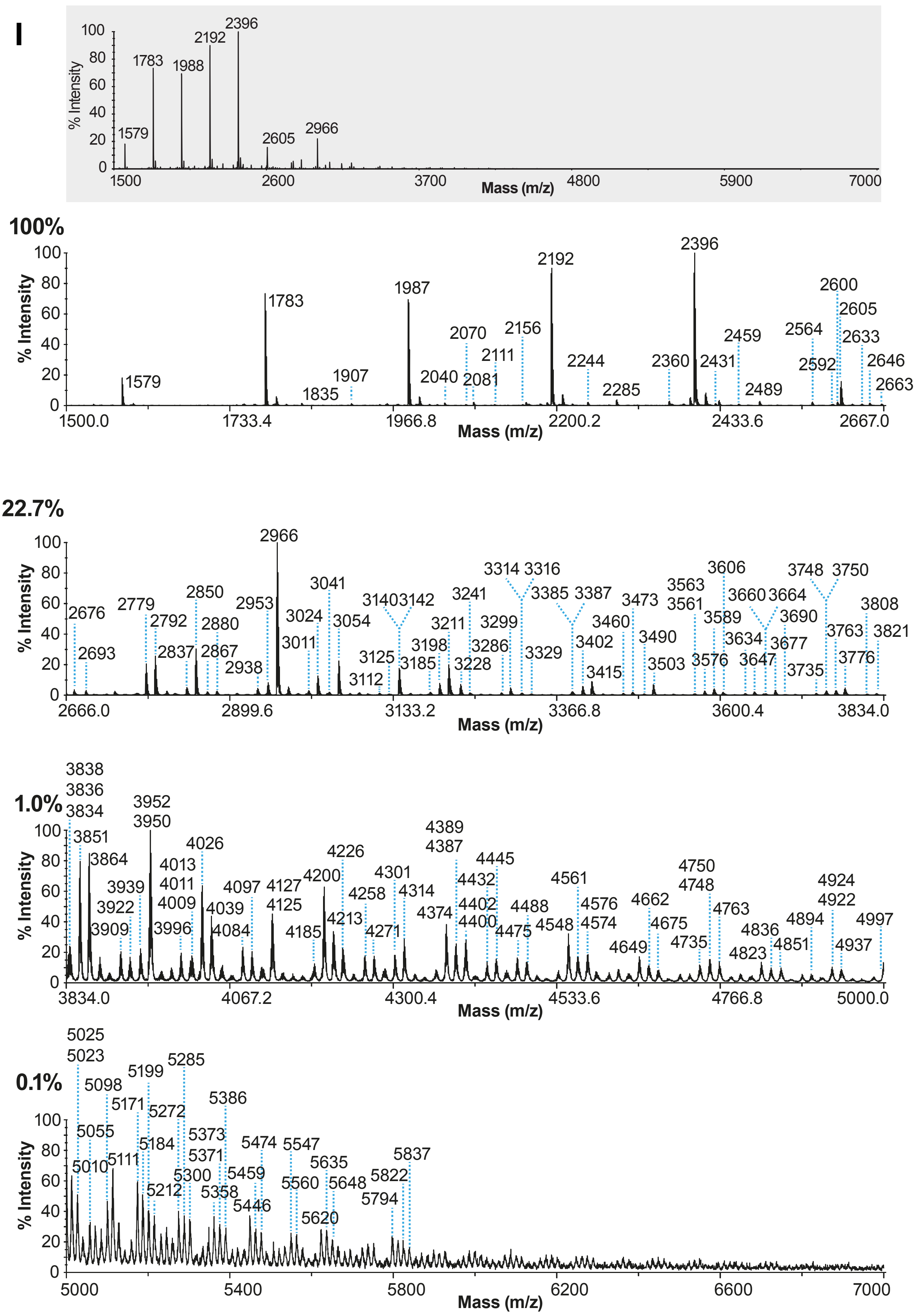

% Intensity
Mass (m/z)
1500
2600
3700
4800
5900
7000
1579
1783
1988
2192
2396
2605
2966
100%
1500.0
1733.4
1966.8
2200.2
2433.6
2667.0
1579
1783
1835
1907
1987
2040
2070
2081
2111
2156
2192
2244
2285
2360
2396
2431
2459
2489
2564
2592
2600
2605
2633
2646
2663
22.7%
2666.0
2899.6
3133.2
3366.8
3600.4
3834.0
2676
2693
2779
2792
2837
2850
2867
2880
2938
2953
2966
3011
3024
3041
3054
3112
3125
3140 3142
3185
3198
3211
3228
3241
3286
3299
3314 3316
3329
3385 3387
3402
3415
3460
3473
3490
3503
3561
3563
3576
3589
3606
3634
3647
3660 3664
3677
3690
3735
3748 3750
3763
3776
3808
3821
1.0%
3834.0
4067.2
4300.4
4533.6
4766.8
5000.0
3838
3836
3834
3851
3864
3909
3922
3939
3950
3952
3996
4009
4011
4013
4026
4039
4084
4097
4125
4127
4185
4200
4213
4226
4258
4271
4301
4314
4374
4387
4389
4400
4402
4432
4445
4475
4488
4548
4561
4574
4576
4649
4662
4675
4735
4748
4750
4763
4823
4836
4851
4894
4922
4924
4937
4997
0.1%
5000
5400
5800
6200
6600
7000
5025
5023
5010
5055
5098
5111
5171
5184
5199
5212
5272
5285
5300
5358
5371
5373
5386
5446
5459
5474
5547
5560
5620
5635
5648
5794
5822
5837

**Figure S1**

**MALDI-TOF mass spectra of permethylated N-glycans derived from primary tracheal epithelial cells from human donors.** MALDI-TOF MS analysis of permethylated N-linked glycans from epithelial cells of (A) TTr01a, (B) TTr01b, (C) TTr02, (D) TTr03, (E) TTr05, (F) TTr06, (G) TTr08, (H) TTr09 and (I) TTr10 human trachea donors. All molecular ions are $[M+Na]^+$. Mass spectra in grey shaded areas correspond to full range spectra in single panels (m/z 1500-7000). Numbers above peaks correspond to the molecular ions identified based on manual annotation. These values were verified by the de-isotoping algorithm (see Methods details). The percentages on top left of each panel correspond to the relative percentage of the maximum peak of the corresponding panel compared to the relative intensity of the maximum peak of the top panel (based on the total ion current of each panel, without de-isotoping). “M” or “m” correspond to major or minor relative abundance of the corresponding molecular ion relative to the molecular ion found in the same molecular ion cluster. Full structure annotations can be found in **Figure S5**.

The molecular ion at m/z 4763 has been detected on all human trachea samples analyzed in various relative abundances.

**Figure S2**

# A

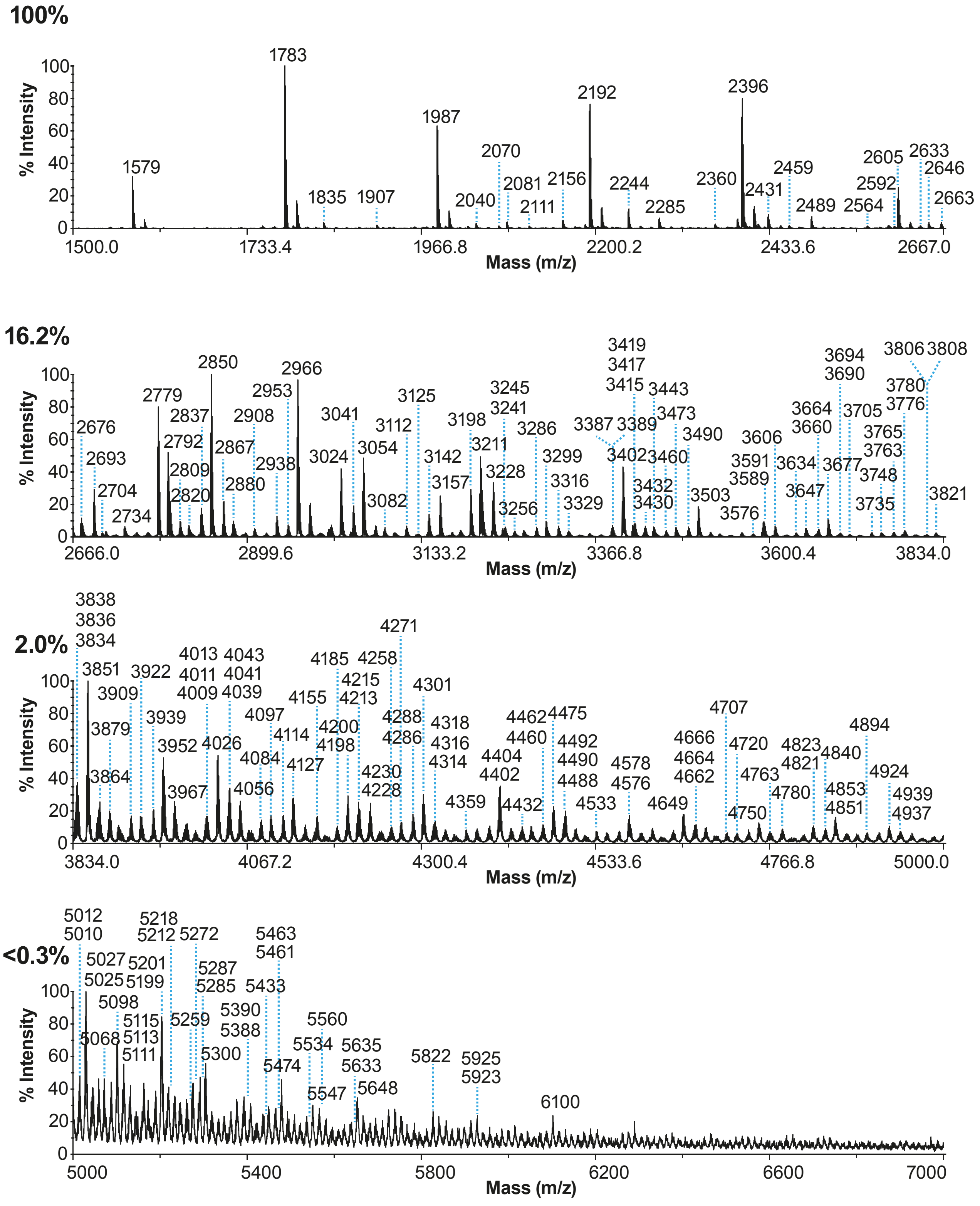
100%
% Intensity
Mass (m/z)
1500.0 1733.4 1966.8 2200.2 2433.6 2667.0
1579 1783 1835 1907 1987 2040 2070 2081 2111 2156 2192 2244 2285 2360 2396 2431 2459 2489 2564 2592 2605 2633 2646 2663
16.2%
2666.0 2899.6 3133.2 3366.8 3600.4 3834.0
2676 2693 2704 2734 2779 2792 2809 2820 2837 2850 2867 2880 2908 2938 2953 2966 3024 3041 3054 3082 3112 3125 3142 3157 3198 3211 3228 3241 3245 3256 3286 3299 3316 3329 3387 3389 3402 3415 3417 3419 3430 3432 3443 3460 3473 3490 3503 3576 3589 3591 3606 3634 3647 3660 3664 3677 3690 3694 3705 3735 3748 3763 3765 3776 3780 3806 3808 3821
2.0%
3834.0 4067.2 4300.4 4533.6 4766.8 5000.0
3834 3836 3838 3851 3864 3879 3909 3922 3939 3952 3967 4009 4011 4013 4026 4039 4041 4043 4056 4084 4097 4114 4127 4155 4185 4198 4200 4213 4215 4228 4230 4258 4271 4286 4288 4301 4314 4316 4318 4359 4402 4404 4432 4460 4462 4475 4488 4490 4492 4533 4576 4578 4649 4662 4664 4666 4707 4720 4750 4763 4780 4821 4823 4840 4851 4853 4894 4924 4937 4939
<0.3%
5000 5400 5800 6200 6600 7000
5010 5012 5025 5027 5068 5098 5111 5113 5115 5199 5201 5212 5218 5259 5272 5285 5287 5300 5388 5390 5433 5461 5463 5474 5534 5547 5560 5633 5635 5648 5822 5923 5925 6100

B

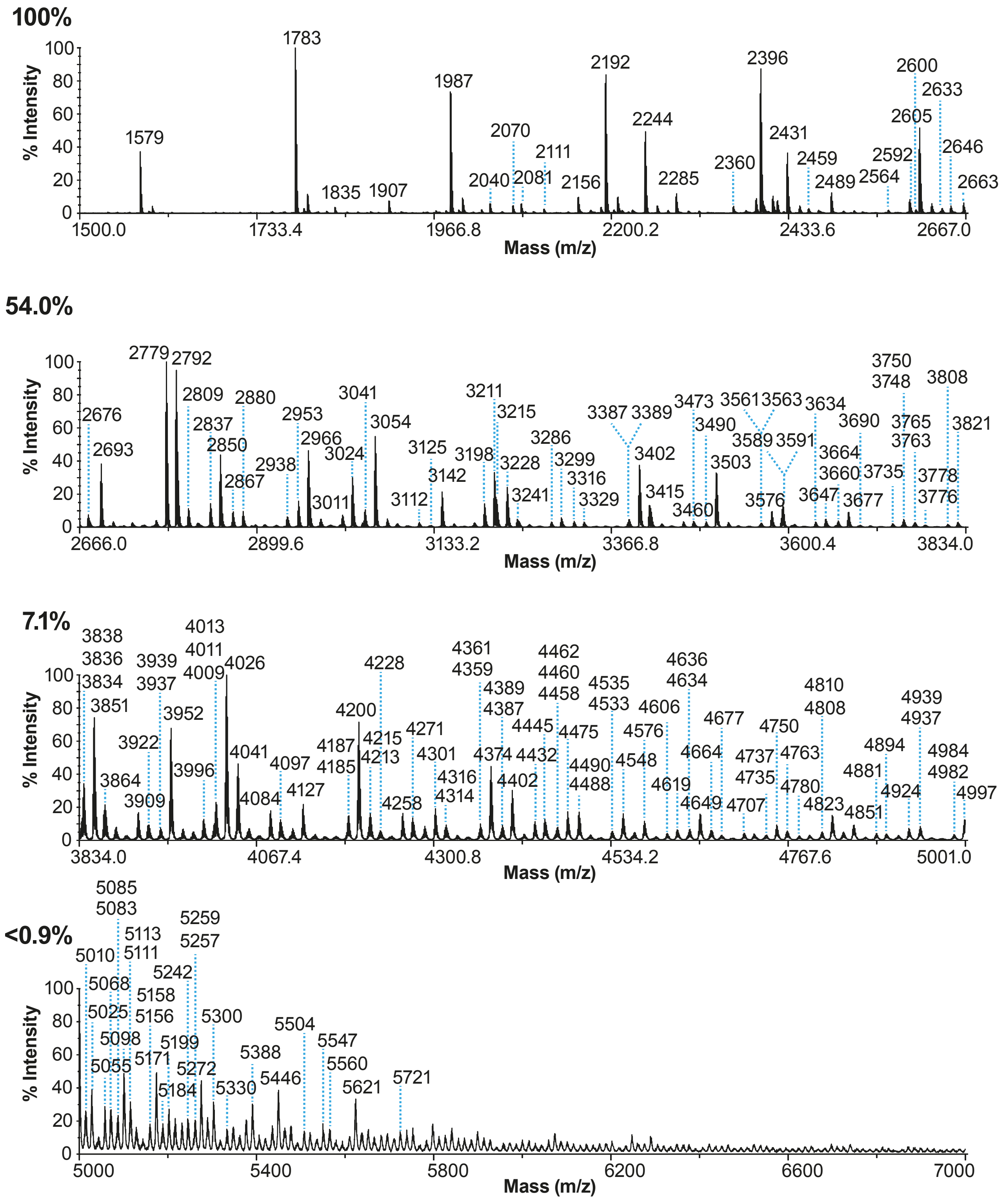

100%
% Intensity
100
80
60
40
20
0
1579
1783
1835
1907
1987
2040
2070
2081
2111
2156
2192
2244
2285
2360
2396
2431
2459
2489
2564
2592
2600
2605
2633
2646
2663
1500.0
1733.4
1966.8
2200.2
2433.6
2667.0
Mass (m/z)
54.0%
% Intensity
100
80
60
40
20
0
2676
2693
2779
2792
2809
2837
2850
2867
2880
2938
2953
2966
3011
3024
3041
3054
3112
3125
3142
3198
3211
3215
3228
3241
3286
3299
3316
3329
3387
3389
3402
3415
3460
3473
3490
3503
3561
3563
3576
3589
3591
3634
3647
3660
3664
3677
3690
3735
3748
3750
3763
3765
3776
3778
3808
3821
2666.0
2899.6
3133.2
3366.8
3600.4
3834.0
Mass (m/z)
7.1%
% Intensity
100
80
60
40
20
0
3838
3836
3834
3851
3864
3909
3922
3939
3937
3952
3996
4013
4011
4009
4026
4041
4084
4097
4127
4187
4185
4200
4215
4213
4228
4258
4271
4301
4316
4314
4361
4359
4374
4389
4387
4402
4445
4432
4462
4460
4458
4475
4490
4488
4535
4533
4548
4576
4606
4619
4636
4634
4649
4664
4677
4707
4737
4735
4750
4763
4780
4810
4808
4823
4851
4881
4894
4924
4939
4937
4984
4982
4997
3834.0
4067.4
4300.8
4534.2
4767.6
5001.0
Mass (m/z)
<0.9%
% Intensity
100
80
60
40
20
0
5010
5025
5055
5068
5085
5083
5098
5113
5111
5158
5156
5171
5184
5199
5242
5259
5257
5272
5300
5330
5388
5446
5504
5547
5560
5621
5721
5000
5400
5800
6200
6600
7000
Mass (m/z)

C

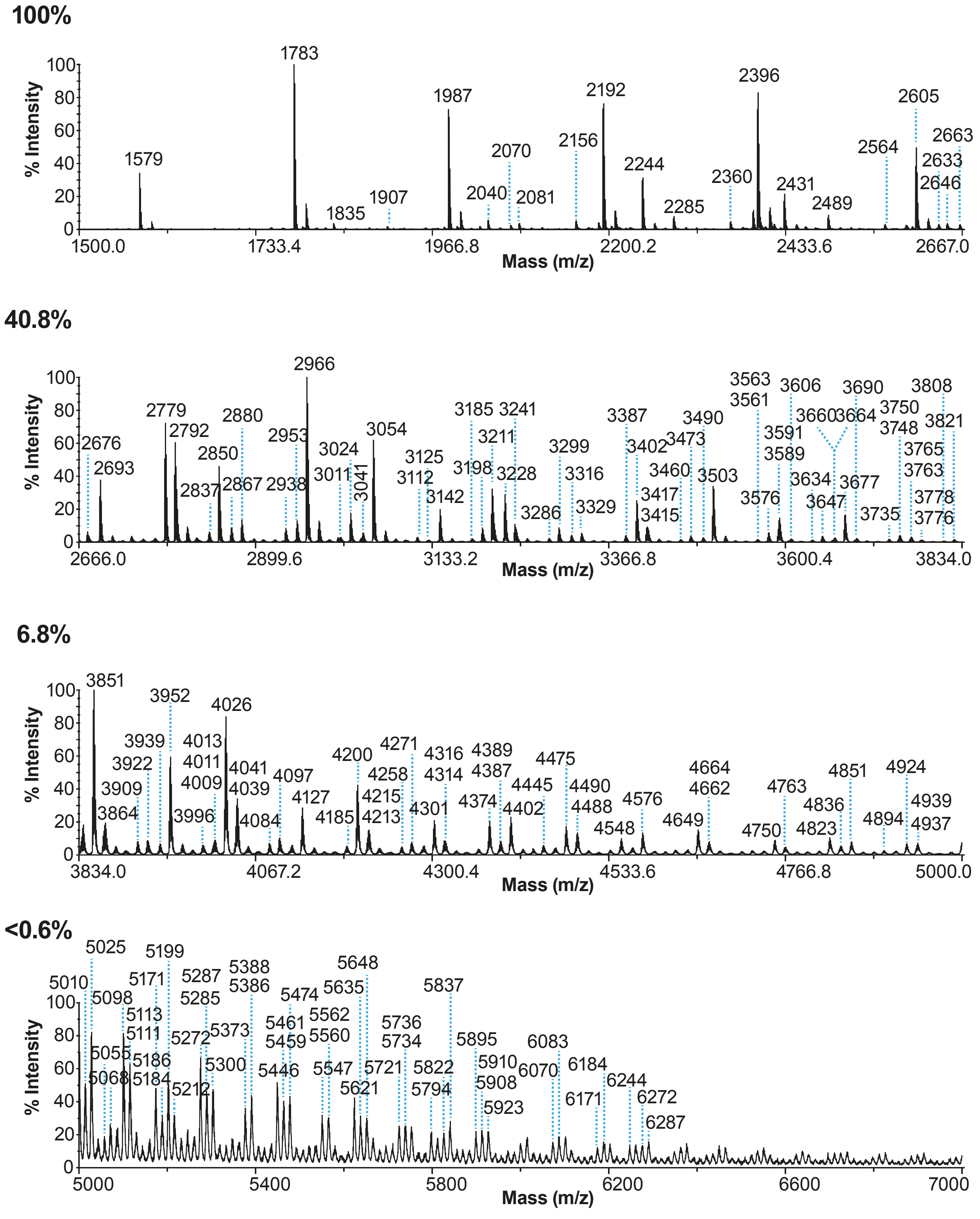
100%
% Intensity
Mass (m/z)
1500.0
1733.4
1966.8
2200.2
2433.6
2667.0
1579
1783
1835
1907
1987
2040
2070
2081
2156
2192
2244
2285
2360
2396
2431
2489
2564
2605
2633
2646
2663
40.8%
2666.0
2899.6
3133.2
3366.8
3600.4
3834.0
2676
2693
2779
2792
2837
2850
2867
2880
2938
2953
2966
3011
3024
3041
3054
3112
3125
3142
3185
3198
3211
3228
3241
3286
3299
3316
3329
3387
3402
3415
3417
3460
3473
3490
3503
3561
3563
3576
3589
3591
3606
3634
3647
3660
3664
3677
3690
3735
3748
3750
3763
3765
3776
3778
3808
3821
6.8%
3834.0
4067.2
4300.4
4533.6
4766.8
5000.0
3851
3864
3909
3922
3939
3952
3996
4009
4011
4013
4026
4039
4041
4084
4097
4127
4185
4200
4213
4215
4258
4271
4301
4314
4316
4374
4387
4389
4402
4445
4475
4488
4490
4548
4576
4649
4662
4664
4750
4763
4823
4836
4851
4894
4924
4937
4939
<0.6%
5000
5400
5800
6200
6600
7000
5010
5025
5055
5068
5098
5111
5113
5171
5184
5186
5199
5212
5272
5285
5287
5300
5373
5386
5388
5446
5459
5461
5474
5547
5560
5562
5621
5635
5648
5721
5734
5736
5794
5822
5837
5895
5908
5910
5923
6070
6083
6171
6184
6244
6272
6287
0
20
40
60
80
100

**Figure S2**

**MALDI-TOF mass spectra of permethylated N-glycans derived from primary tracheal epithelial cells from human donors after Sial-S digestion.** MALDI-TOF MS analysis of permethylated N-linked glycans from epithelial cells of (A) TTr8; (B) TTr9 and (C) TTr10 human trachea samples (found in **Figure S1**) after Sial-S digestion. All molecular ions are $[M+Na]^+$. Numbers above peaks correspond to the molecular ions identified based on manual annotation. These values were verified by the de-isotoping algorithm (see Methods details). Percentages on top left of each panel correspond to the relative percentage of the maximum peak of the corresponding panel compared to the relative intensity of the maximum peak of the top panel (based on the total ion current of each panel, without de-isotoping). Full structure annotations can be found in **Figure S5**.

The molecular ion at m/z 4763 has been detected on all human trachea samples analyzed in various relative abundances.

**Figure S3**

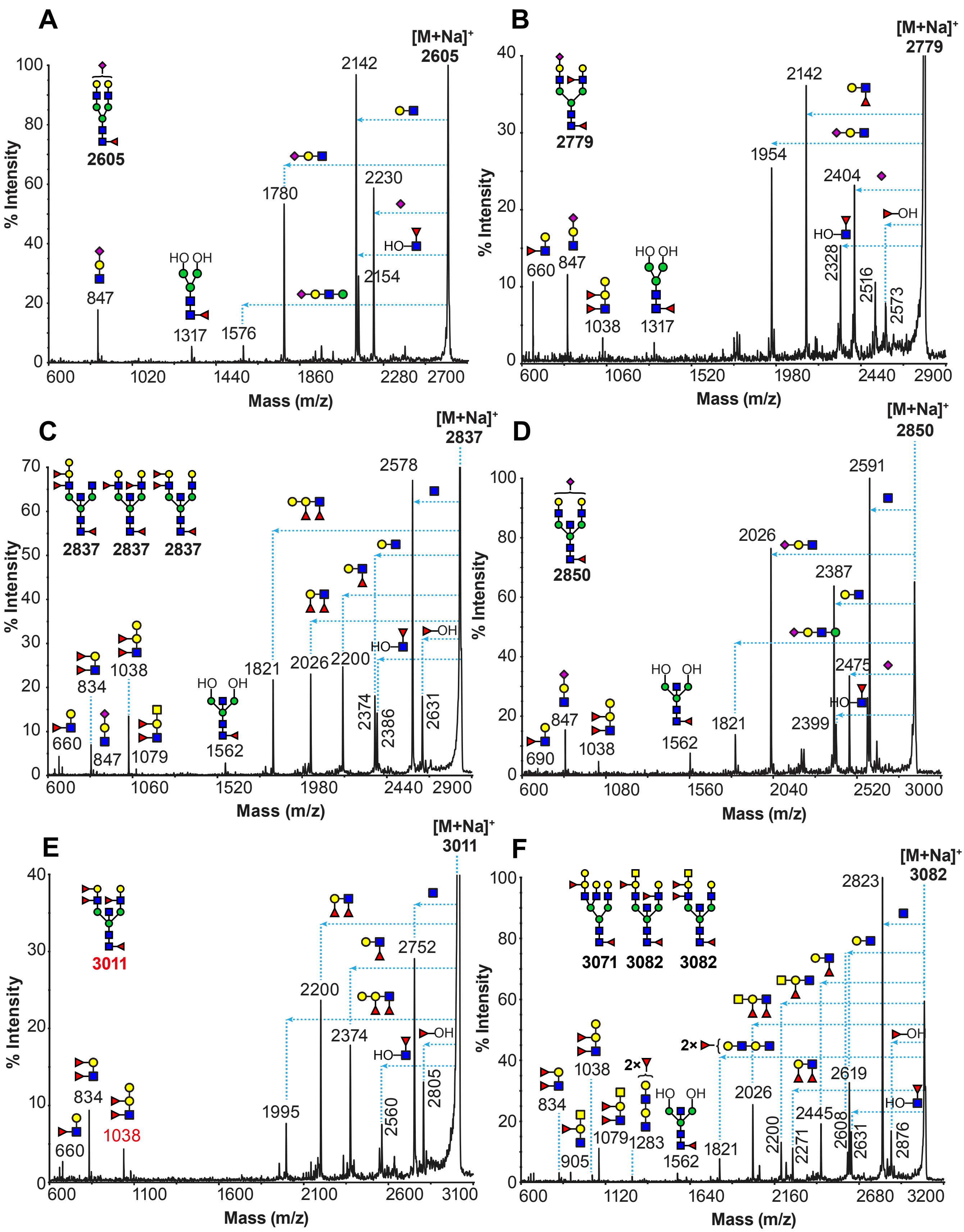


Fuc Man Gal GlcNAc GlcNAc NeuAc

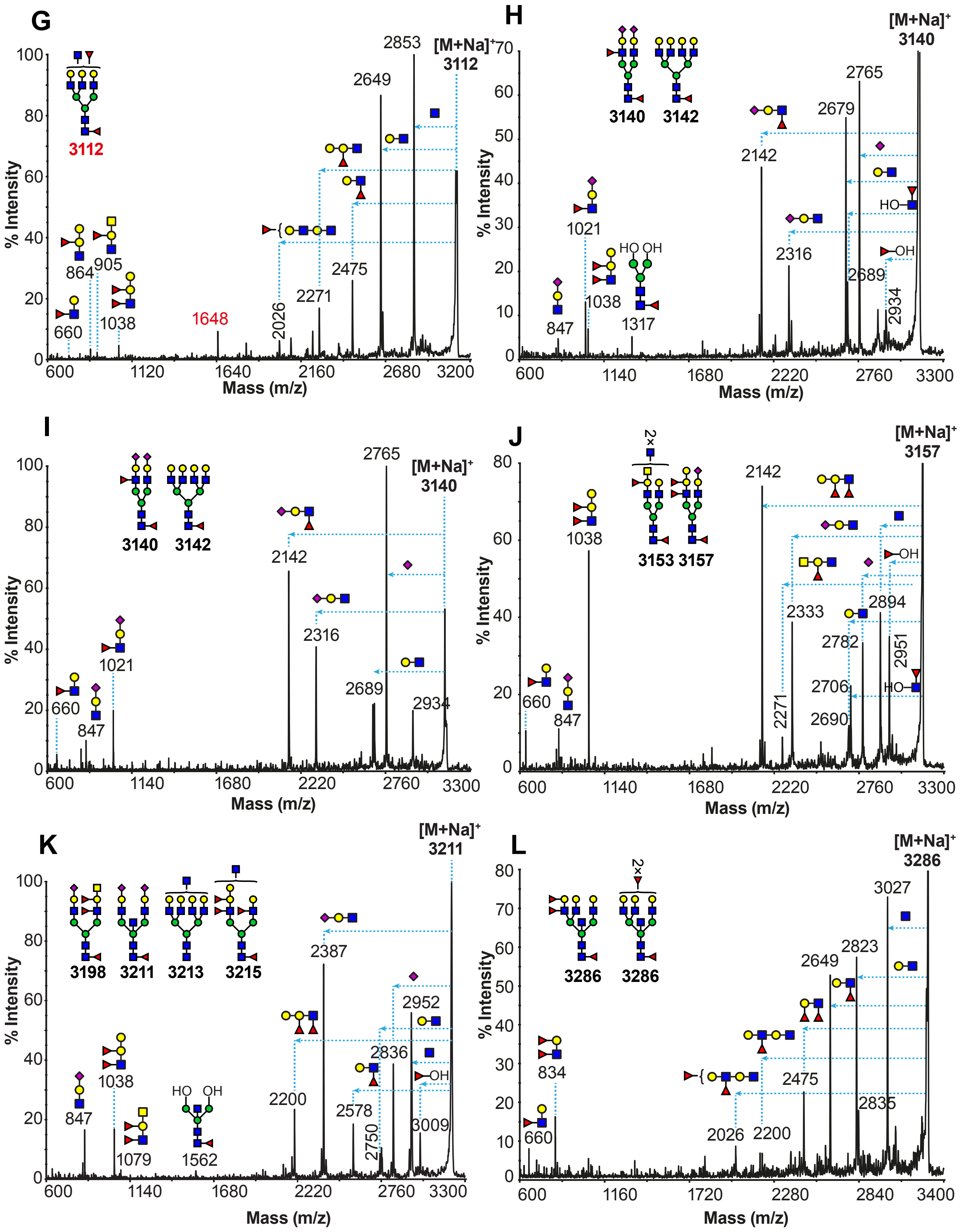
G
% Intensity
Mass (m/z)
[M+Na]+
3112
2853
2649
2475
2271
2026
1648
1038
905
864
660
H
[M+Na]+
3140
2765
2679
2689
2934
2316
2142
1317
1038
1021
847
3140
3142
HO
OH
I
[M+Na]+
3140
2765
2934
2689
2316
2142
1021
847
660
3140
3142
J
[M+Na]+
3157
2142
2333
2271
2894
2951
2782
2706
2690
1038
847
660
3153 3157
2×
HO
OH
K
[M+Na]+
3211
2387
2952
2836
2750
2578
3009
2200
1562
1079
1038
847
3198
3211
3213
3215
HO
OH
L
[M+Na]+
3286
3027
2823
2649
2475
2835
2200
2026
834
660
3286
3286
2×

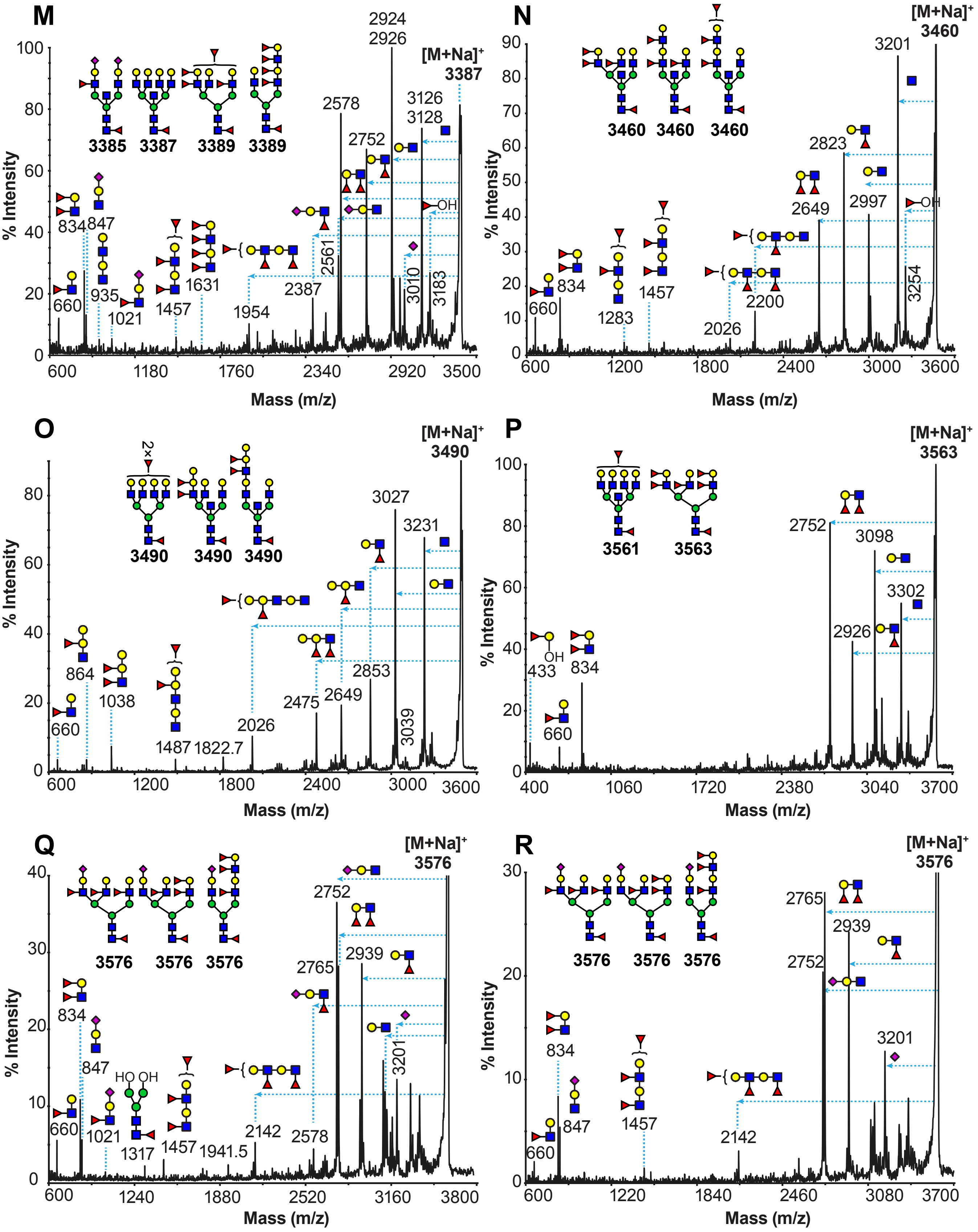

M
3385 3387 3389 3389
[M+Na]+ 3387
% Intensity
Mass (m/z)
660 834 847 935 1021 1457 1631 1954 2387 2561 2578 2752 2924 2926 3010 3126 3128 3183
OH
N
3460 3460 3460
[M+Na]+ 3460
% Intensity
Mass (m/z)
660 834 1283 1457 2026 2200 2649 2823 2997 3201 3254
OH
O
2×
3490 3490 3490
[M+Na]+ 3490
% Intensity
Mass (m/z)
660 864 1038 1487 1822.7 2026 2475 2649 2853 3027 3039 3231
P
3561 3563
[M+Na]+ 3563
% Intensity
Mass (m/z)
OH
433 660 834 2752 2926 3098 3302
Q
3576 3576 3576
[M+Na]+ 3576
% Intensity
Mass (m/z)
HO OH
660 834 847 1021 1317 1457 1941.5 2142 2578 2752 2765 2939 3201
R
3576 3576 3576
[M+Na]+ 3576
% Intensity
Mass (m/z)
660 834 847 1457 2142 2752 2765 2939 3201

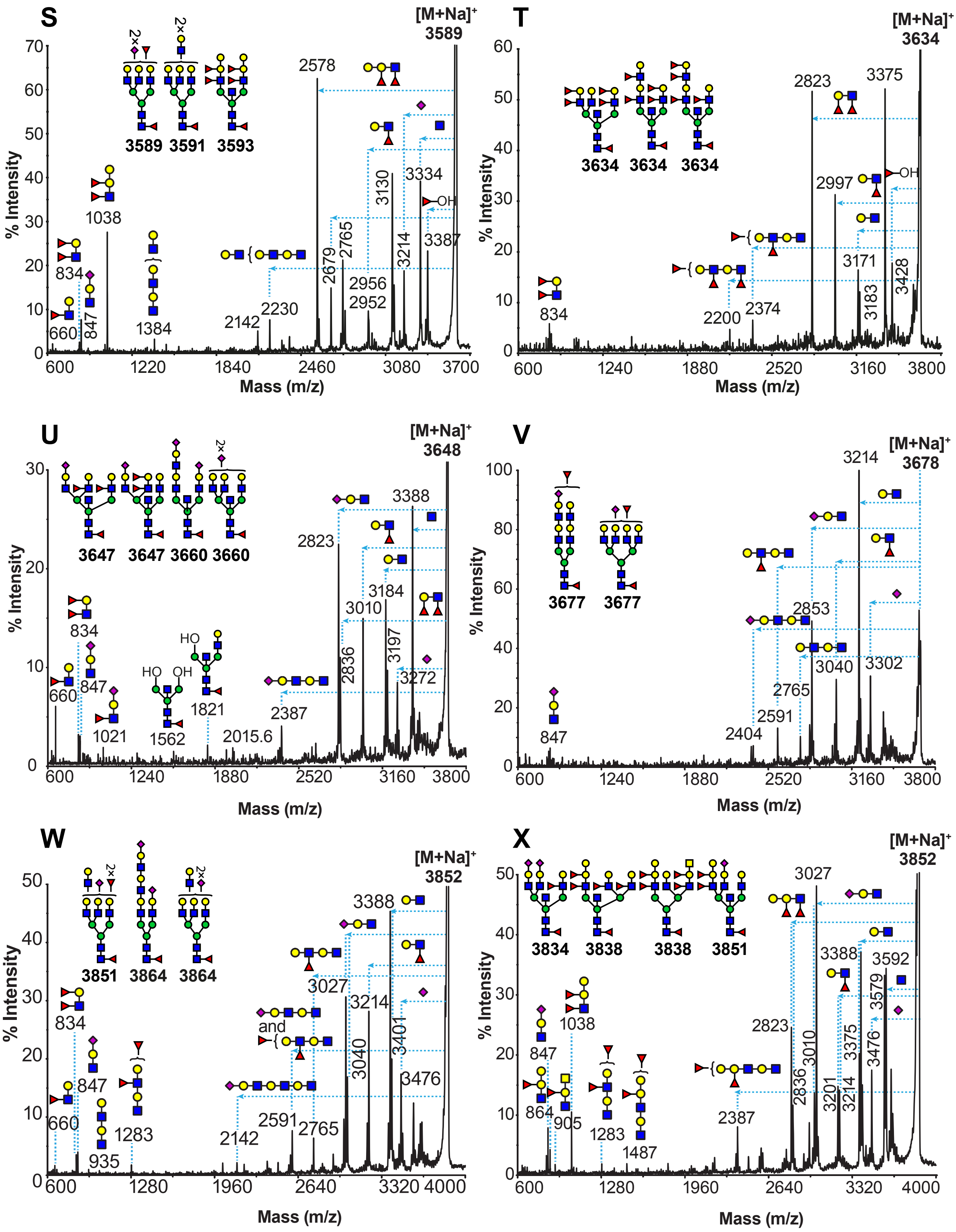

S
[M+Na]+
3589
% Intensity
Mass (m/z)
3589 3591 3593
2×
2578
1038
834
660
847
1384
2142
2230
2679
2765
2956
2952
3130
3214
3334
3387
OH
T
[M+Na]+
3634
3634 3634 3634
2823
3375
2997
OH
834
2200
2374
3171
3183
3428
U
[M+Na]+
3648
3647 3647 3660 3660
2×
2823
3388
3184
3010
834
660
847
1021
HO
OH
1562
1821
2015.6
2387
2836
3197
3272
V
[M+Na]+
3678
3214
3677 3677
847
2853
3040
3302
2765
2591
2404
W
[M+Na]+
3852
3851 3864 3864
2×
3388
3027
3214
834
847
660
935
1283
and
2142
2591
2765
3040
3401
3476
X
[M+Na]+
3852
3834 3838 3838 3851
3027
3388
3592
2823
3010
2836
3201
3214
3375
3579
3476
847
1038
864
905
1283
1487
2387

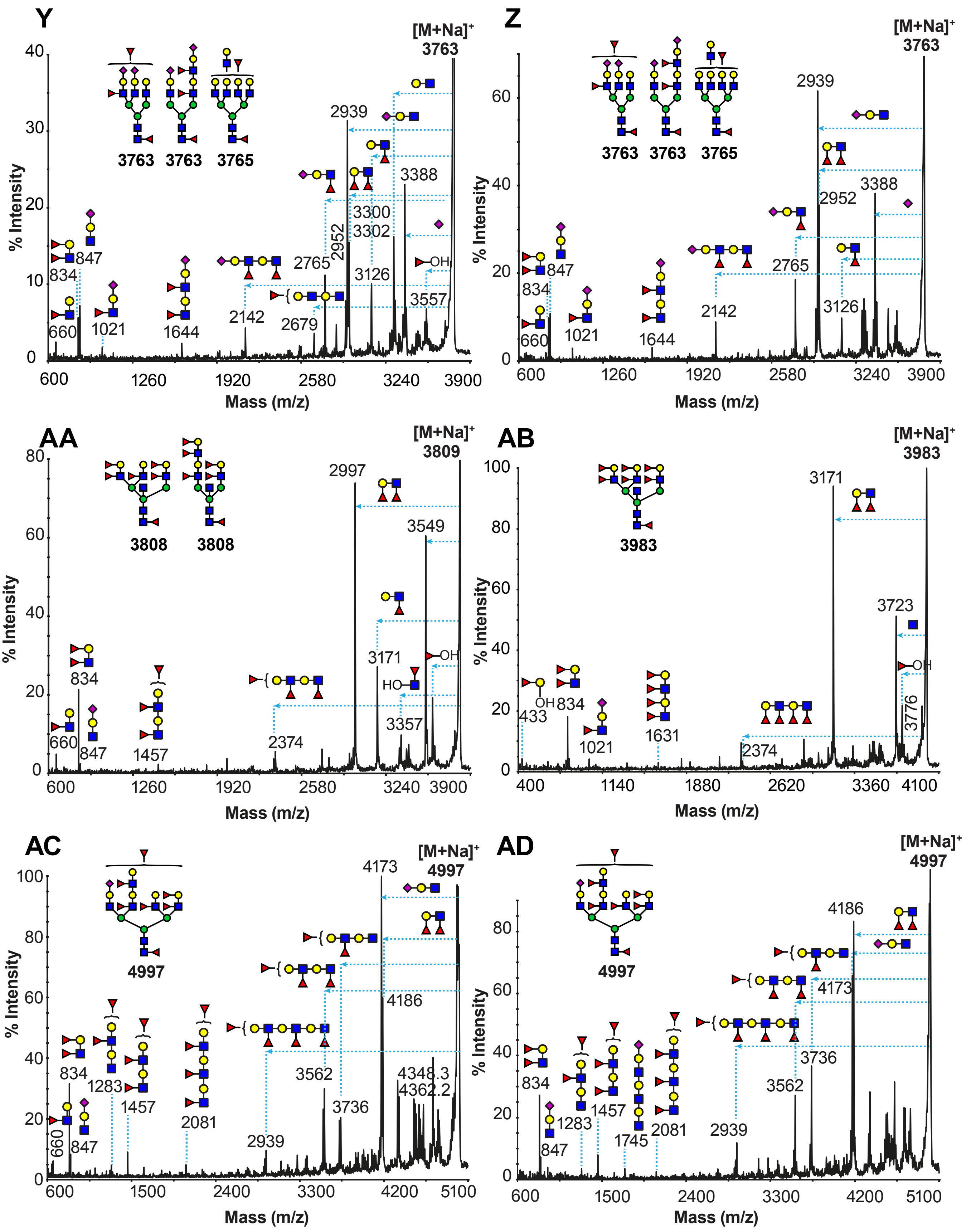

Y
[M+Na]+ 3763
3763 3763 3765
% Intensity
Mass (m/z)
660
834
847
1021
1644
2142
2679
2765
2952
2939
3300
3302
3126
3388
3557
OH
Z
[M+Na]+ 3763
3763 3763 3765
2939
2952
3388
2765
2142
3126
660
834
847
1021
1644
AA
[M+Na]+ 3809
3808 3808
2997
3549
3171
3357
2374
1457
660
834
847
HO
OH
AB
[M+Na]+ 3983
3983
3171
3723
3776
2374
1631
1021
433
834
OH
AC
[M+Na]+ 4997
4997
4173
4186
4348.3
4362.2
3562
3736
2939
2081
1457
1283
834
847
660
AD
[M+Na]+ 4997
4997
4186
4173
3736
3562
2939
2081
1745
1457
1283
834
847

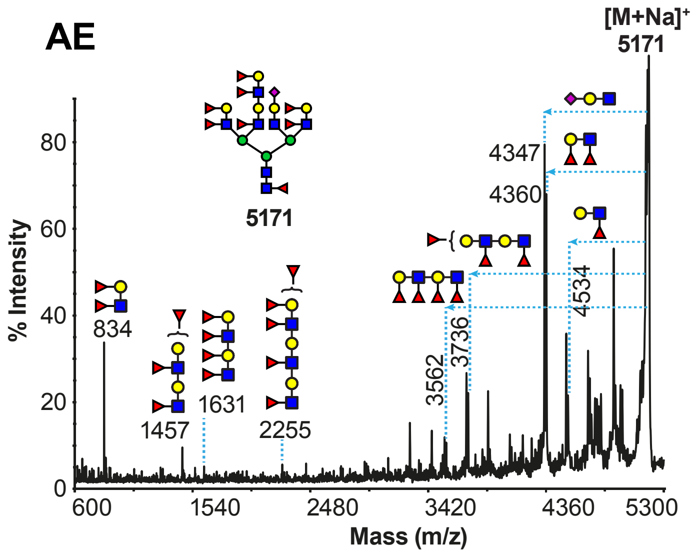

AE
[M+Na]+
5171
5171
80
60
40
20
0
% Intensity
834
1457
1631
2255
3562
3736
4347
4360
4534
600
1540
2480
3420
4360
5300
Mass (m/z)

**Figure S3**

**MALDI TOF/TOF MS/MS analysis of permethylated N-glycans derived from primary tracheal epithelial cells from human donors.** Molecular ions $[M+Na]^+$ present in **Figure S1** (TTr08, TTr09 and TTr10) were selected for MALDI-TOF/TOF MS/MS analysis. (A) m/z 2605 from TTr10, (B) m/z 2779 from TTr08, (C) m/z 2837 from TTr08, (D) m/z 2850 from TTr08, (E) m/z 3011 from TTr08, (F) m/z 3082 from TTr08, (G) m/z 3112 from TTr08, (H) m/z 3140 from TTR08, (I) m/z 3140 from TTr10, (J) m/z 3157 from TTr08, (K) m/z 3211 from TTr08 (L) m/z 3286 from TTr09, (M) m/z 3387 from TTr09, (N) m/z 3460 from TTr09, (O) m/z 3490 from TTr08, (P) m/z 3563 from TTr09, (Q) m/z 3576 from TTr09, (R) m/z 3576 from TTr10, (S) m/z 3589 from TTr08, (T) m/z 3634 from TTr09, (U) m/z 3648 from TTr10, (V) m/z 3678 from TTr10, (W) m/z 3852 from TTr10, (X) m/z 3852 from TTr08, (Y) m/z 3763 from TTr09, (Z) m/z 3763 from TTr10, (AA) m/z 3809 from TTr09, (AB) m/z 3983 from TTr09, (AC) m/z 4997 from TTr09, (AD) m/z 4997 from TTr10, (AE) m/z 5171 from TTr09. Cartoon structures in the corresponding spectra do not represent all possible structural isomers, for clarity. Horizontal dashed lines correspond to the indicated fragment losses from each of the corresponding molecular ion $[M+Na]^+$. Vertical dashed lines indicate the m/z fragment ion peak that corresponds to the fragment loss as shown from the corresponding horizontal dashed line. Structures outside the bracket have not unequivocally been defined. For clarity, not all fragments were annotated.

The detection of fragment ions corresponding to an elimination of a fucose residue are indicative of being linked to the C-3 position of a GlcNAc residue corresponding to LeX and LeY epitopes. Examples are as follows: the fragment ions at m/z 2573

(B), 2631 (C), 2805 (E), 2876 (F), 2951 (J), 3183 (M), 3254 (N). This elimination is also indicative of sLeX epitope. Examples are as follows: the fragment ions at m/z 2934 (H and I).

The detection of fragment ions corresponding to an elimination of a terminal fucose-galactose structure is indicative of being linked to the C-3 position of a GlcNAc residue corresponding to LeB epitopes. Example is as follows: the fragment ion at m/z 433 (P and AB).

#### Figure S4

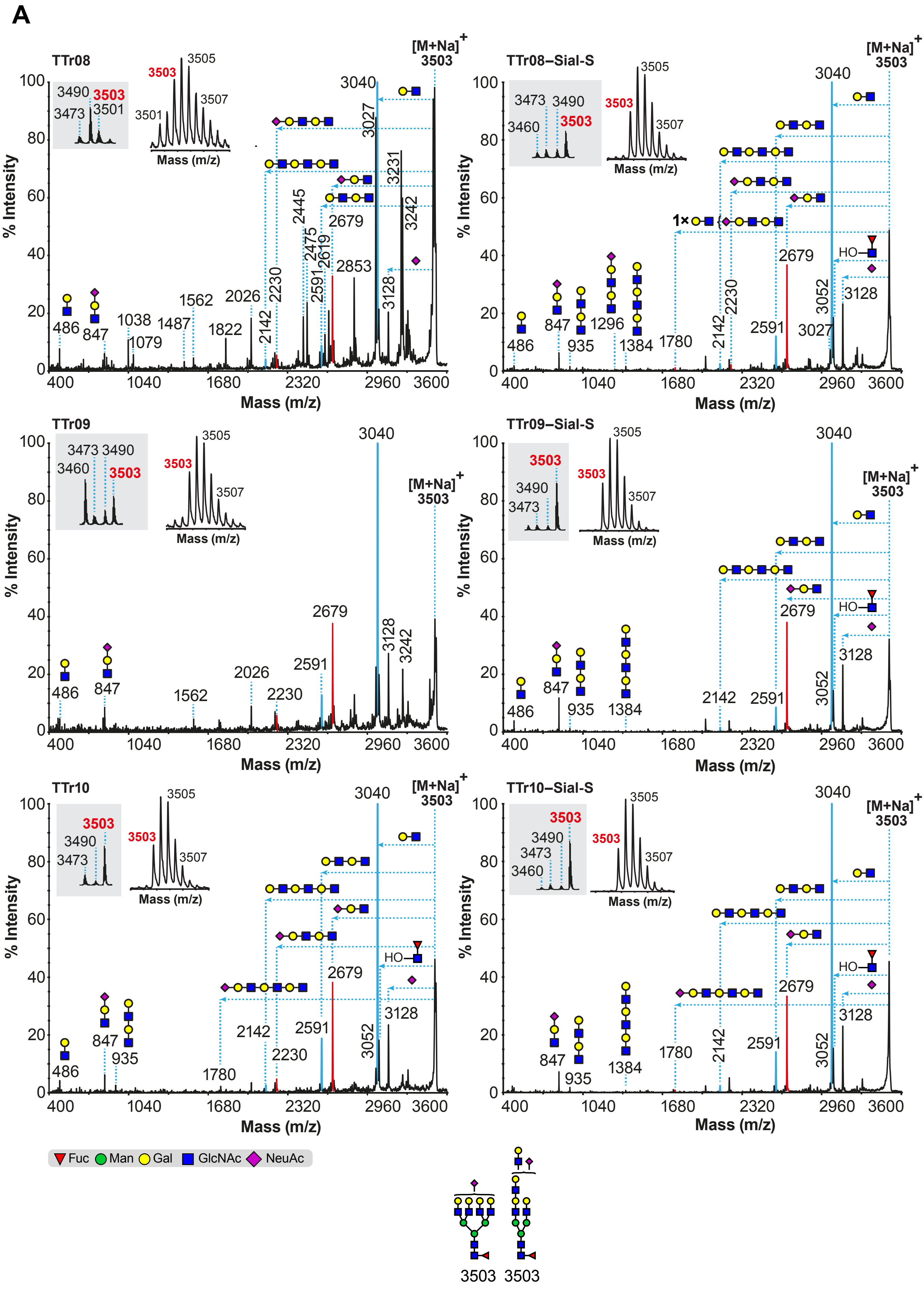
A
TTr08
TTr08–Sial-S
TTr09
TTr09–Sial-S
TTr10
TTr10–Sial-S
[M+Na]+
3503
3040
% Intensity
Mass (m/z)
2679
3128
3052
2591
2142
2230
1780
1384
935
847
486
Fuc Man Gal GlcNAc NeuAc
3503 3503

B

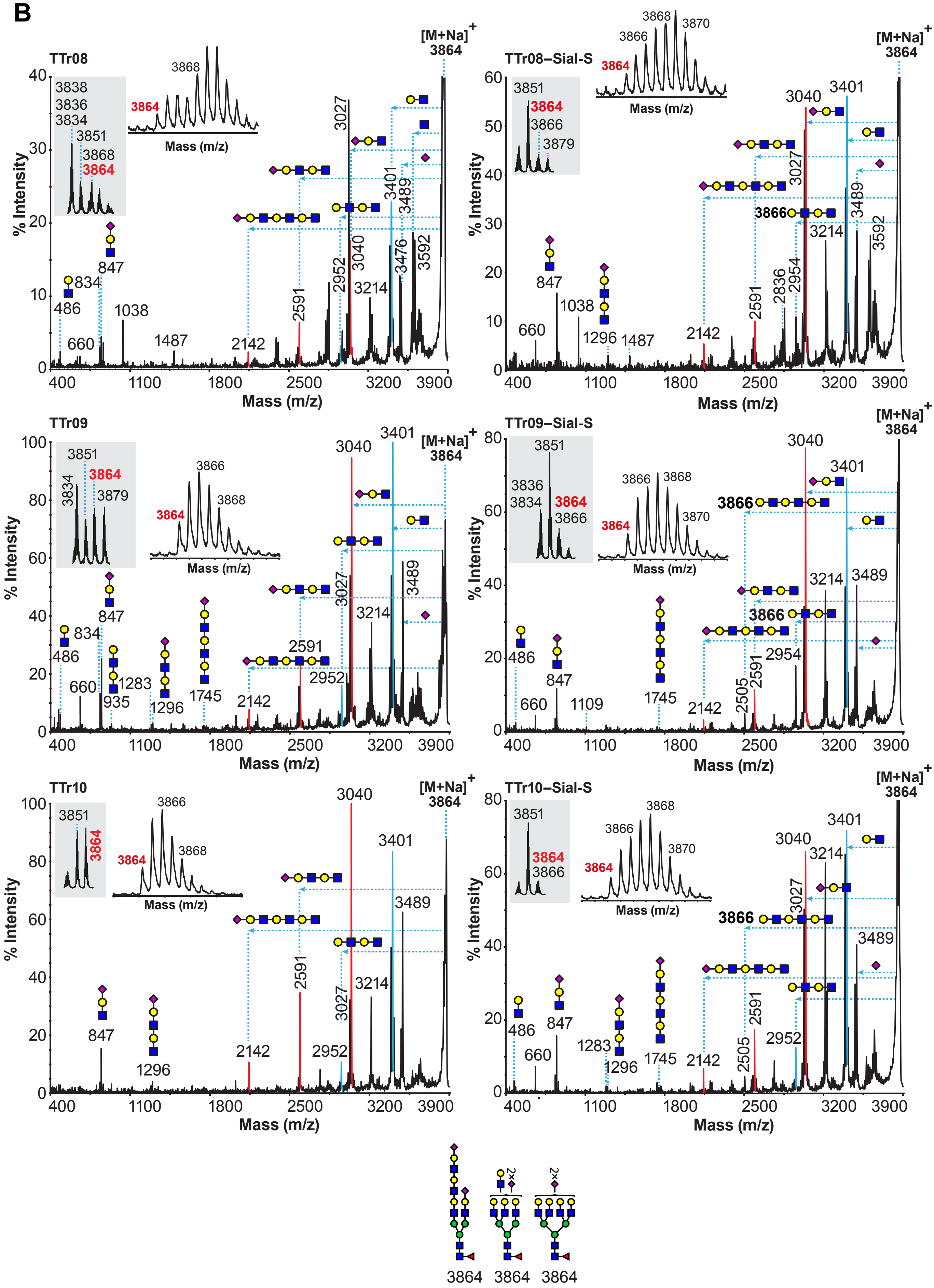
TTr08
TTr08–Sial-S
TTr09
TTr09–Sial-S
TTr10
TTr10–Sial-S
[M+Na]+
3864
% Intensity
Mass (m/z)
3838
3836
3834
3851
3868
3866
3870
3879
486
660
834
847
935
1038
1109
1283
1296
1487
1745
2142
2505
2591
2836
2952
2954
3027
3040
3214
3401
3476
3489
3592
2×
3864 3864 3864

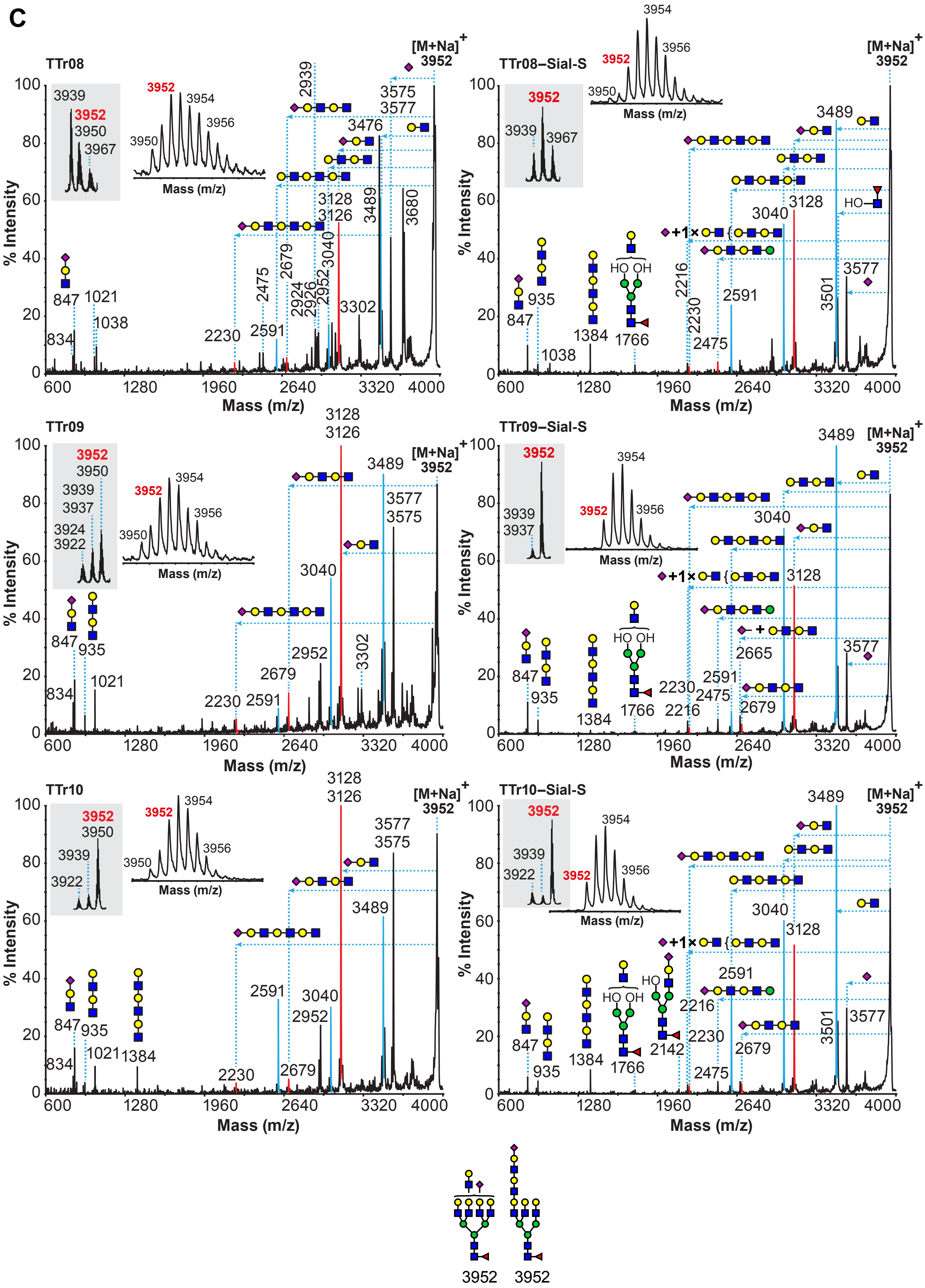

C
TTr08
TTr08–Sial-S
TTr09
TTr09–Sial-S
TTr10
TTr10–Sial-S
[M+Na]+
3952
% Intensity
Mass (m/z)
3939
3950
3954
3956
3967
3937
3924
3922
847
834
935
1021
1038
1384
1766
2142
2216
2230
2475
2591
2665
2679
2924
2926
2939
2952
3040
3126
3128
3302
3476
3489
3501
3575
3577
3680
HO OH
HO
+1×

D

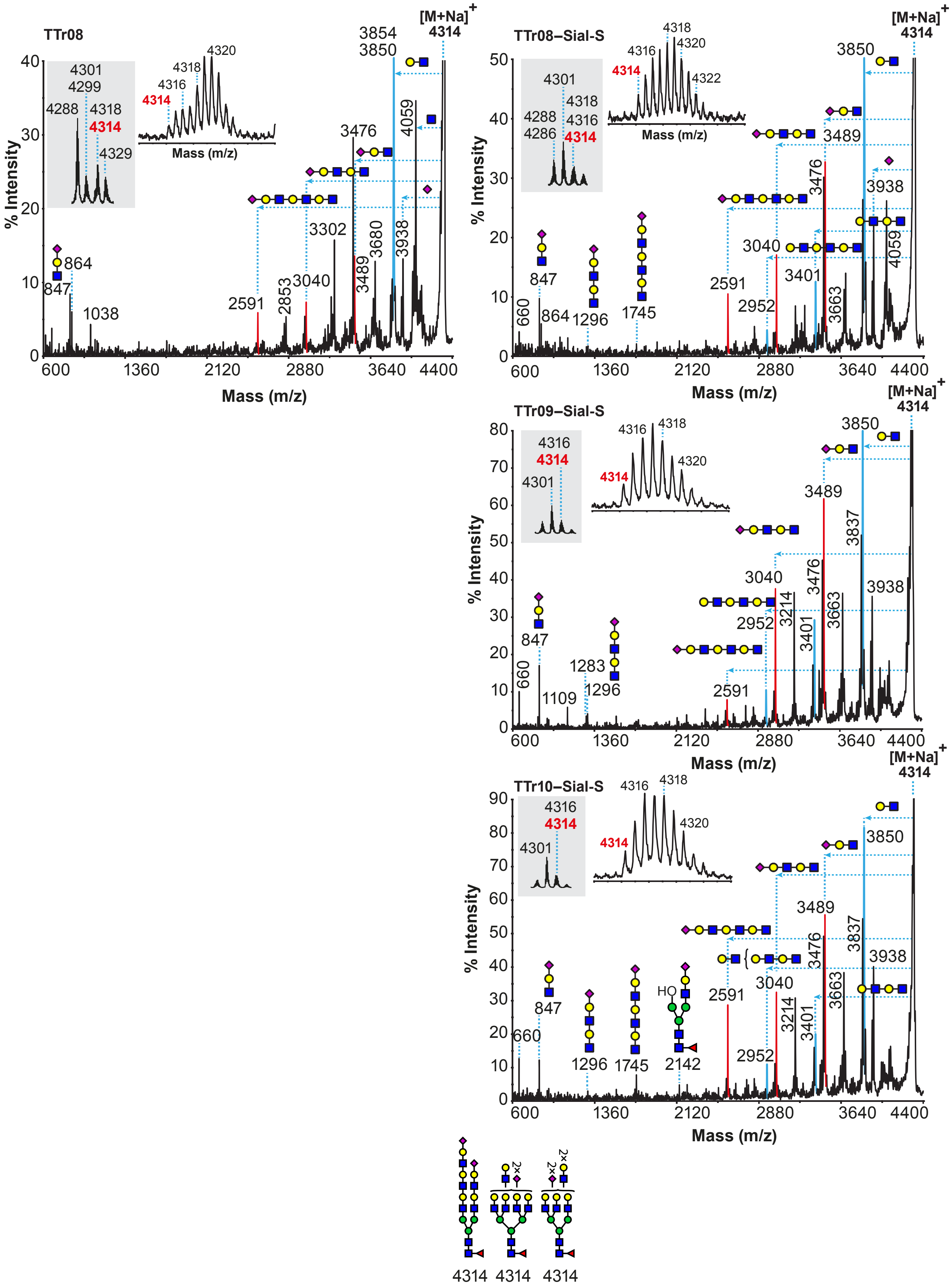

TTr08
TTr08–Sial-S
TTr09–Sial-S
TTr10–Sial-S
[M+Na]+
4314
% Intensity
Mass (m/z)
3850
3854
3476
3489
3040
2591
3302
3680
3938
4059
847
864
1038
1296
1745
2952
3401
3663
3214
3837
660
1109
1283
2142
4301
4299
4288
4286
4316
4318
4320
4322
4329
HO
2×
4314 4314 4314

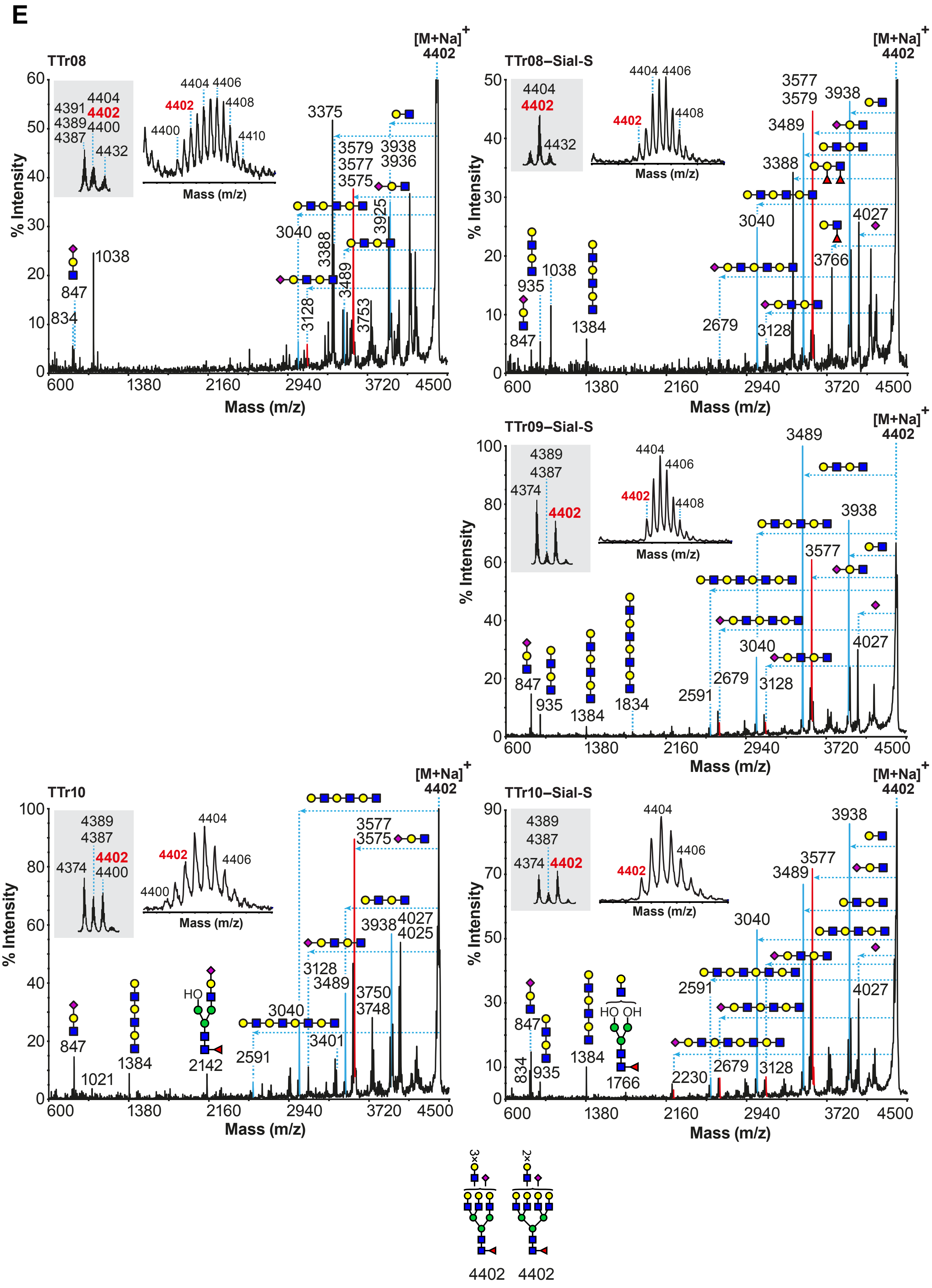
E
TTr08
[M+Na]+
4402
% Intensity
Mass (m/z)
4391
4389
4387
4404
4402
4400
4432
4404 4406
4408
4400
4410
3375
3579 3938
3577 3936
3575
3040
3388
3925
3128
3489
3753
847
834
1038
600
1380
2160
2940
3720
4500
TTr08–Sial-S
3577
3579
3938
3489
3388
3040
4027
3766
2679
3128
935
1038
847
1384
TTr09–Sial-S
4389
4387
4374
4402
4404
4406
4408
3489
3938
3577
3040
4027
2679
3128
2591
847
935
1384
1834
TTr10
4389
4387
4374
4402
4400
4404
4406
3577
3575
3938
4027
4025
3128
3489
3750
3748
3040
3401
2591
847
1021
1384
HO
2142
TTr10–Sial-S
3938
3577
3489
3040
2591
4027
2230
2679
3128
834
847
935
1384
HO OH
1766
3×
2×
4402
4402

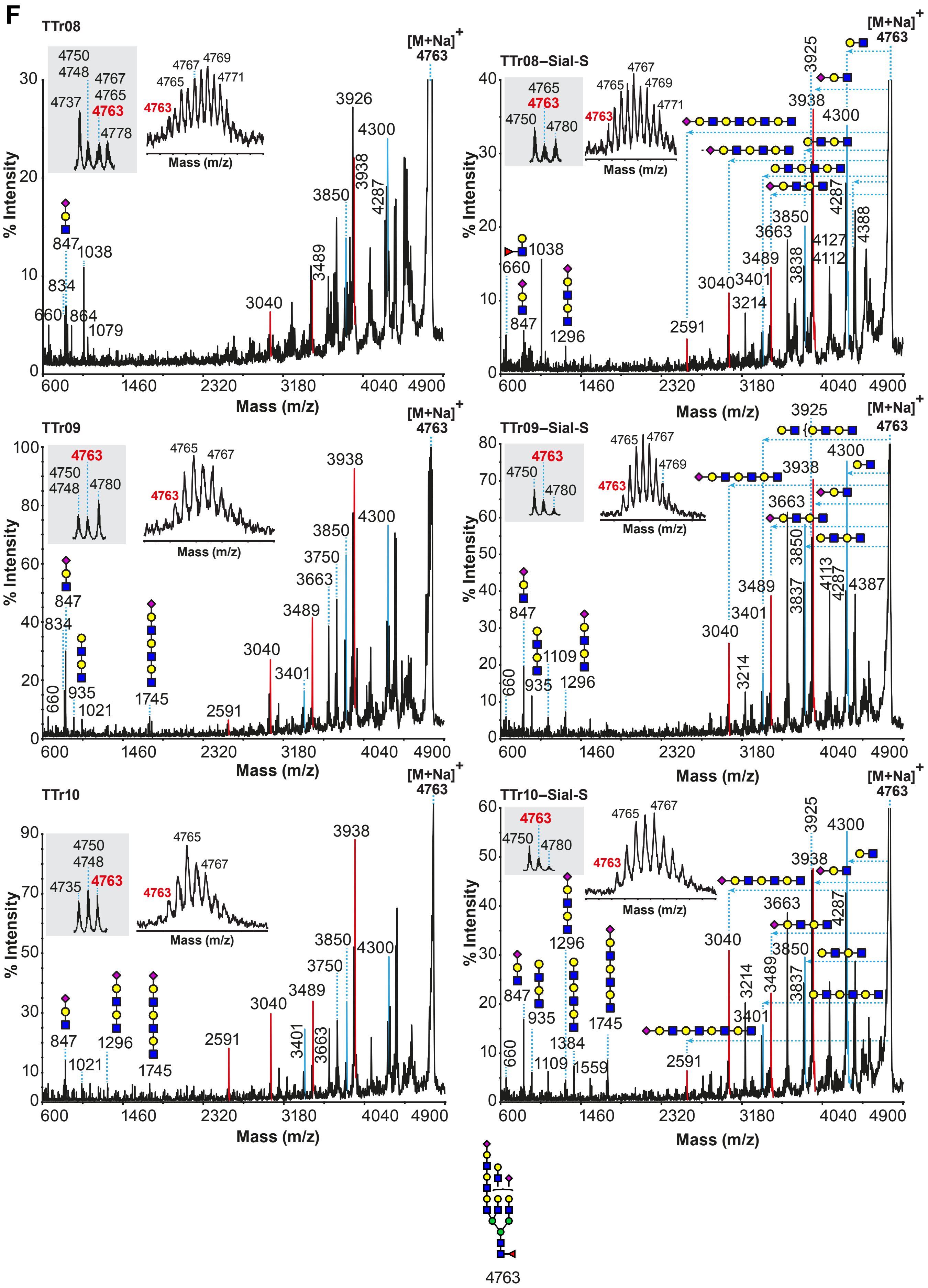
F
TTr08
TTr08–Sial-S
TTr09
TTr09–Sial-S
TTr10
TTr10–Sial-S
[M+Na]+
4763
% Intensity
Mass (m/z)
600
1460
2320
3180
4040
4900
3926
3938
4300
3850
4287
3489
3040
3925
3663
3750
3401
3214
2591
847
834
660
864
1038
1079
935
1021
1109
1296
1384
1559
1745
4750
4748
4737
4767
4765
4769
4771
4778
4780
4735
3838
3837
4127
4112
4113
4388
4387

**Figure S4**

**MALDI TOF/TOF MS/MS analysis of permethylated N-glycans derived from primary tracheal epithelial cells from human donors before or after Sial-S digestion.** Molecular ions $[M+Na]^+$ present in **Figure S1G, H** and **I** (TTr08, TTr09, and TTr10, respectively) were selected for MALDI-TOF/TOF MS/MS analysis. Only the fragment ions specific to molecular ions selected for fragmentation are annotated with structures for presentation. (A) m/z 3503, (B) m/z 3864, (C) m/z 3952, (D) m/z 4314, (E) m/z 4402, (F) m/z 4763. Left and right panels correspond to mass spectra before and after Sial-S digestion respectively. Red numbers in insets correspond to the molecular ion selected for MALDI TOF/TOF MS/MS analysis corresponding to N-linked glycans structures that contained sialylated LacNAcs extensions. Horizontal dashed lines indicate fragment losses from the corresponding molecular ion $[M+Na]^+$. Vertical dashed lines indicate the m/z fragment ion peak that corresponded to the fragment loss as shown from the corresponding horizontal dashed line. Solid blue and red colored peaks are fragment ion peaks corresponding to losses of $(LacNAc)_n$ and NeuAc-$(LacNAc)_n$ (where n>=1) type structures respectively which have derived from the selected molecular ions. Structures outside the bracket have not unequivocally been defined. For clarity, cartoon structures in the corresponding spectra do not represent all possible structural isomers, and not all fragment ions have been annotated. Insets on each panel depict zoomed areas and isotopic ion clusters of the corresponding molecular ion derived from **Figure S1G**, **H** and **I**. Note the high complexity of the MSMS data deriving from multiple structural isomers of the molecular ion selected for fragmentation, but also, in certain cases, because of the existence of fragment ions deriving from other molecular ions in proximity to the selected molecular ion.

(A, C, E and F, right panels) In the Sial-S samples note the presence of the fragment ion at m/z 1384 (together with the loss from each molecular ion) corresponding to a 3×LacNAc structure. These fragment ions were detected in higher relative abundance in the samples after Sial-S digestions (right panels) than the samples before Sial-S digestion (left panels). This suggested the existence of poly-LacNAc structures capped with NeuAcα2,3-linked residues that were cleaved off after Sial-S digestion (cleaves α2,3-linked NeuAc residues) meaning that N-linked glycan structures exhibited poly-LacNAc chain covered with NeuAcα2,3-linked residues were also present.

(C) Note the changes occurring to the relative abundances of the molecular ion clusters at m/z 3950-3952 (insets). Before Sial-S digestion (insets in left panels) we could detect the presence of the molecular ion at m/z 3950 corresponding to a N-linked glycan containing a sLeX/sLeA epitope which indicated the presence of NeuAcα2,3-linked residue (**Figure S5** and this Figure below). This was verified by the presence of fragment ions at m/z 1021. However, after Sial-S digestion (insets in right panels) the molecular ion at m/z 3950 was absent, as expected from the Sial-S digestion. Also, this was verified by the absence of the fragment ion at m/z 1021.

#### Figure S5

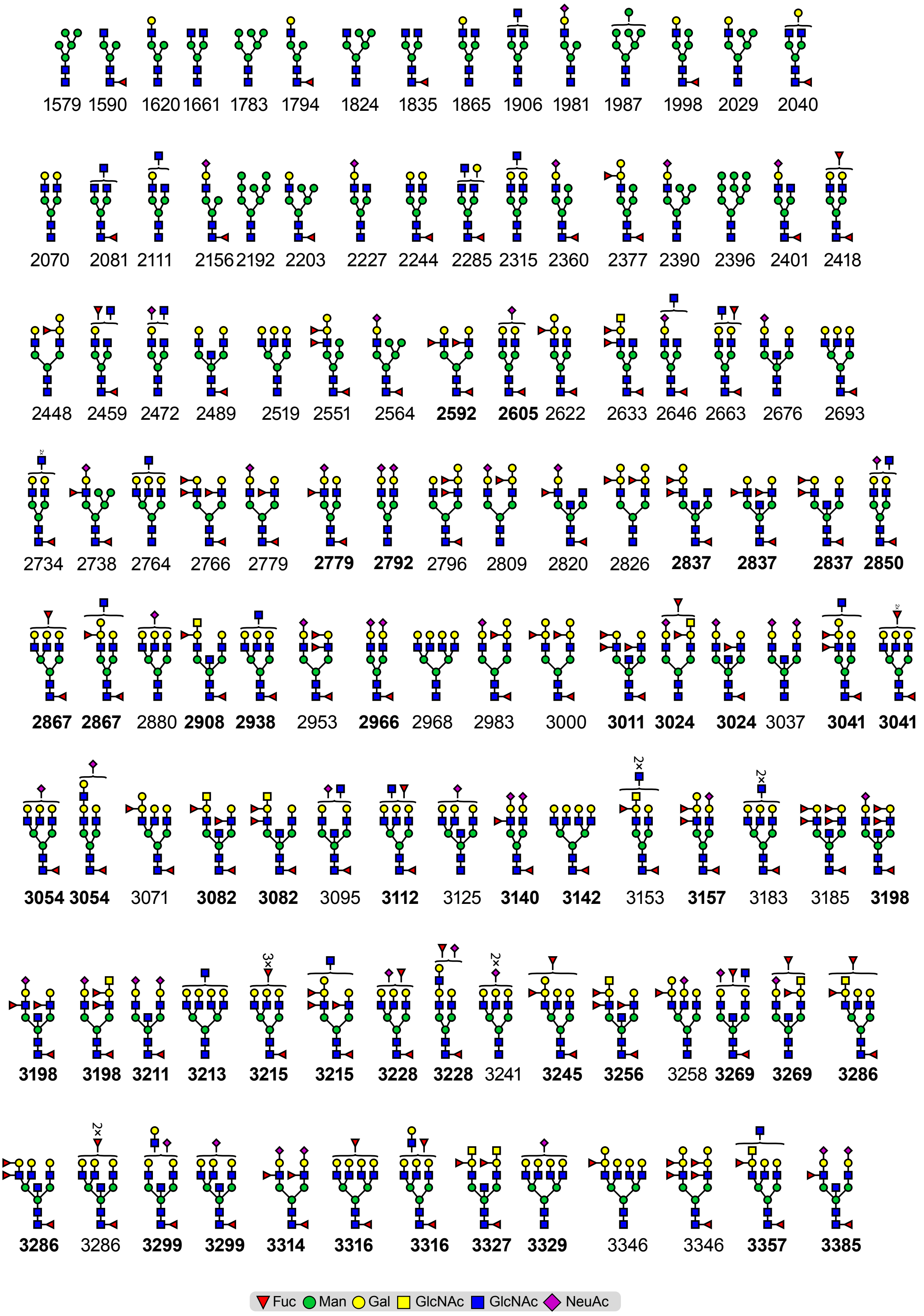
1579 1590 1620 1661 1783 1794 1824 1835 1865 1906 1981 1987 1998 2029 2040
2070 2081 2111 2156 2192 2203 2227 2244 2285 2315 2360 2377 2390 2396 2401 2418
2448 2459 2472 2489 2519 2551 2564 2592 2605 2622 2633 2646 2663 2676 2693
2734 2738 2764 2766 2779 2779 2792 2796 2809 2820 2826 2837 2837 2837 2850
2867 2867 2880 2908 2938 2953 2966 2968 2983 3000 3011 3024 3024 3037 3041 3041
3054 3054 3071 3082 3082 3095 3112 3125 3140 3142 3153 3157 3183 3185 3198
3198 3198 3211 3213 3215 3215 3228 3228 3241 3245 3256 3258 3269 3269 3286
3286 3286 3299 3299 3314 3316 3316 3327 3329 3346 3346 3357 3385
Fuc Man Gal GlcNAc GlcNAc NeuAc

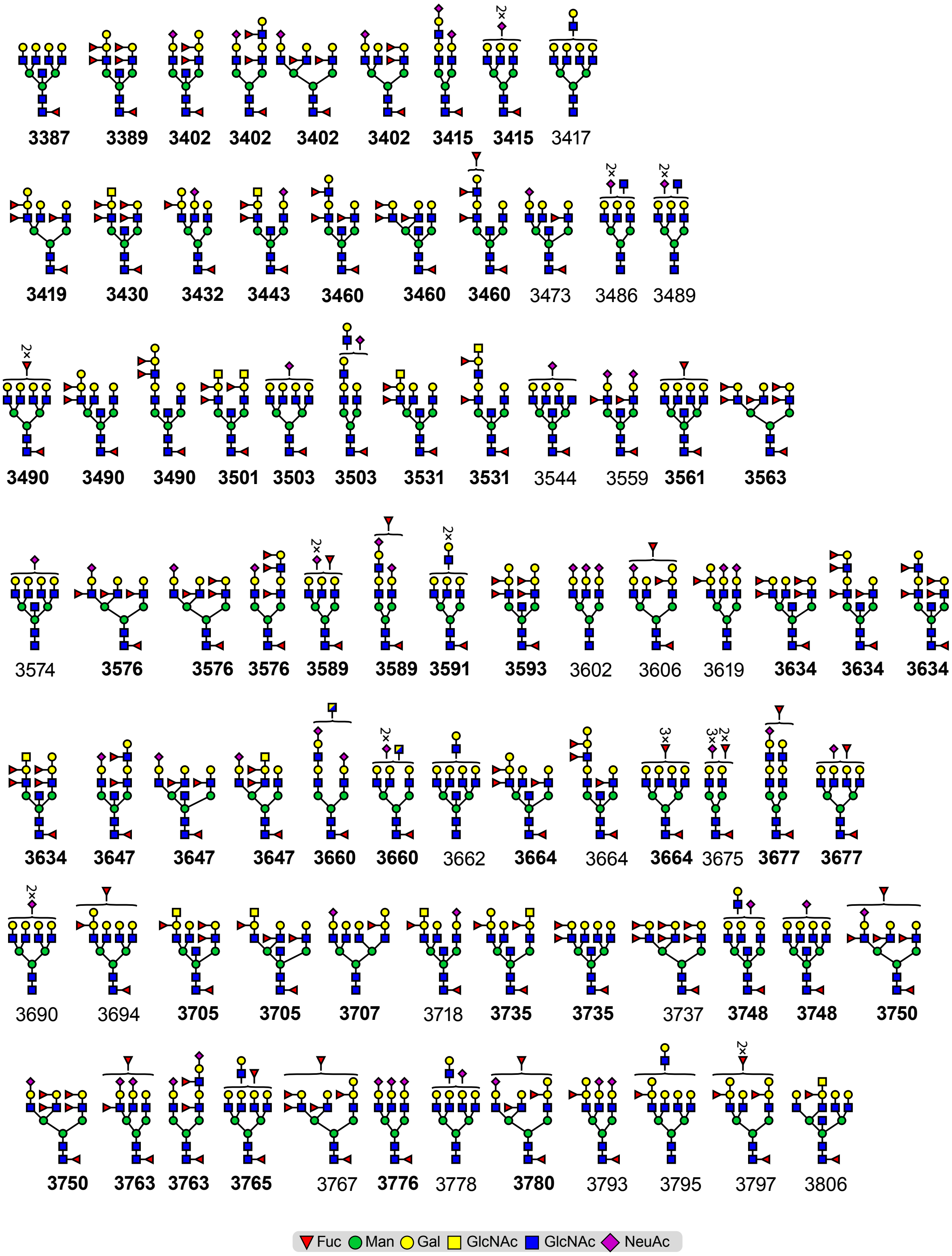
3387 3389 3402 3402 3402 3402 3415 3415 3417
3419 3430 3432 3443 3460 3460 3460 3473 3486 3489
3490 3490 3490 3501 3503 3503 3531 3531 3544 3559 3561 3563
3574 3576 3576 3576 3589 3589 3591 3593 3602 3606 3619 3634 3634 3634
3634 3647 3647 3647 3660 3660 3662 3664 3664 3664 3675 3677 3677
3690 3694 3705 3705 3707 3718 3735 3735 3737 3748 3748 3750
3750 3763 3763 3765 3767 3776 3778 3780 3793 3795 3797 3806
Fuc Man Gal GlcNAc GlcNAc NeuAc

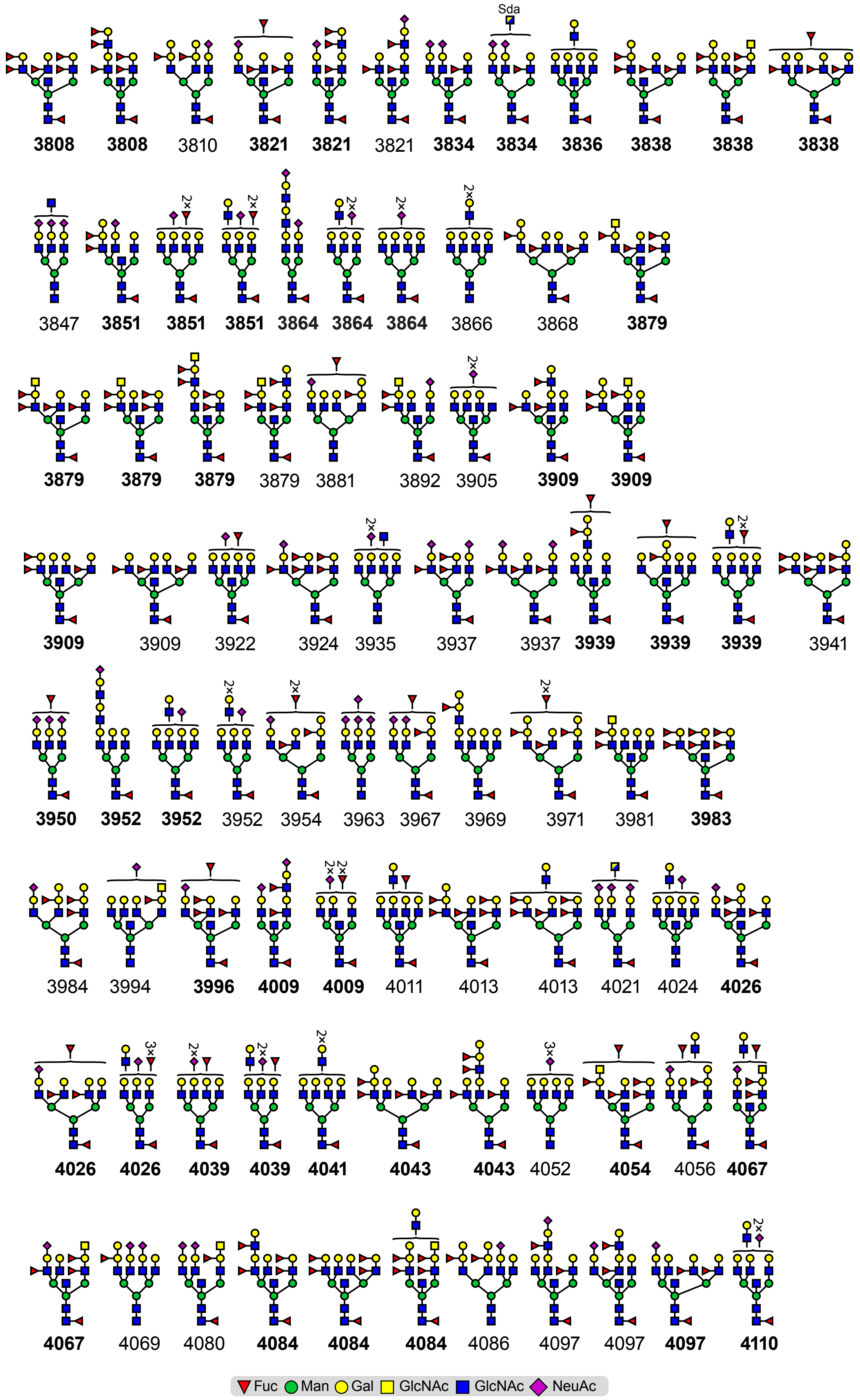
3808 3808 3810 3821 3821 3821 3834 3834 3836 3838 3838 3838
3847 3851 3851 3851 3864 3864 3864 3866 3868 3879
3879 3879 3879 3879 3881 3892 3905 3909 3909
3909 3909 3922 3924 3935 3937 3937 3939 3939 3939 3941
3950 3952 3952 3952 3954 3963 3967 3969 3971 3981 3983
3984 3994 3996 4009 4009 4011 4013 4013 4021 4024 4026
4026 4026 4039 4039 4041 4043 4043 4052 4054 4056 4067
4067 4069 4080 4084 4084 4084 4086 4097 4097 4097 4110
Fuc Man Gal GlcNAc GlcNAc NeuAc

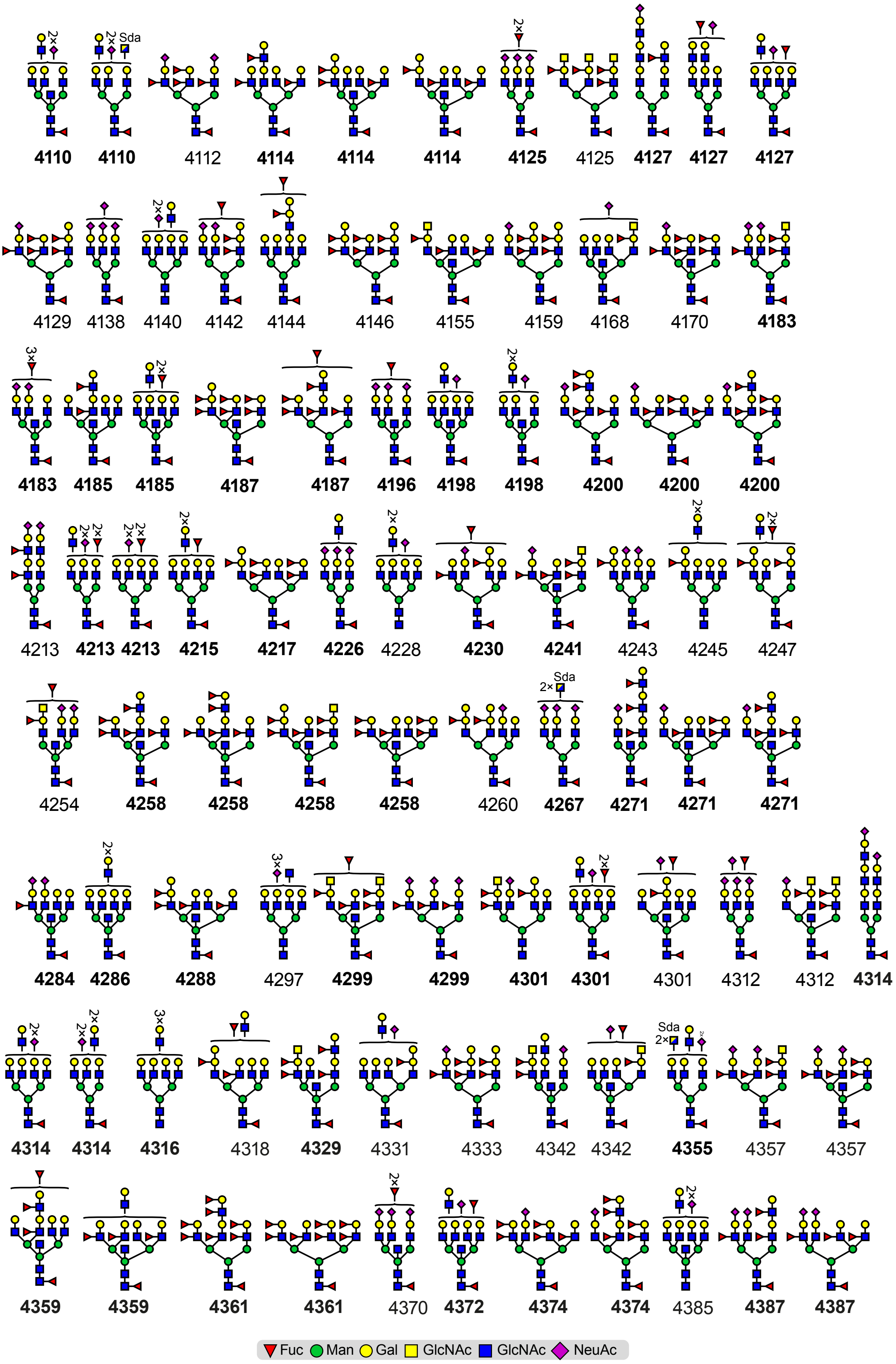

4110 4110 4112 4114 4114 4114 4125 4125 4127 4127 4127
4129 4138 4140 4142 4144 4146 4155 4159 4168 4170 4183
4183 4185 4185 4187 4187 4196 4198 4198 4200 4200 4200
4213 4213 4213 4215 4217 4226 4228 4230 4241 4243 4245 4247
4254 4258 4258 4258 4258 4260 4267 4271 4271 4271
4284 4286 4288 4297 4299 4299 4301 4301 4301 4312 4312 4314
4314 4314 4316 4318 4329 4331 4333 4342 4342 4355 4357 4357
4359 4359 4361 4361 4370 4372 4374 4374 4385 4387 4387
Fuc Man Gal GlcNAc GlcNAc NeuAc

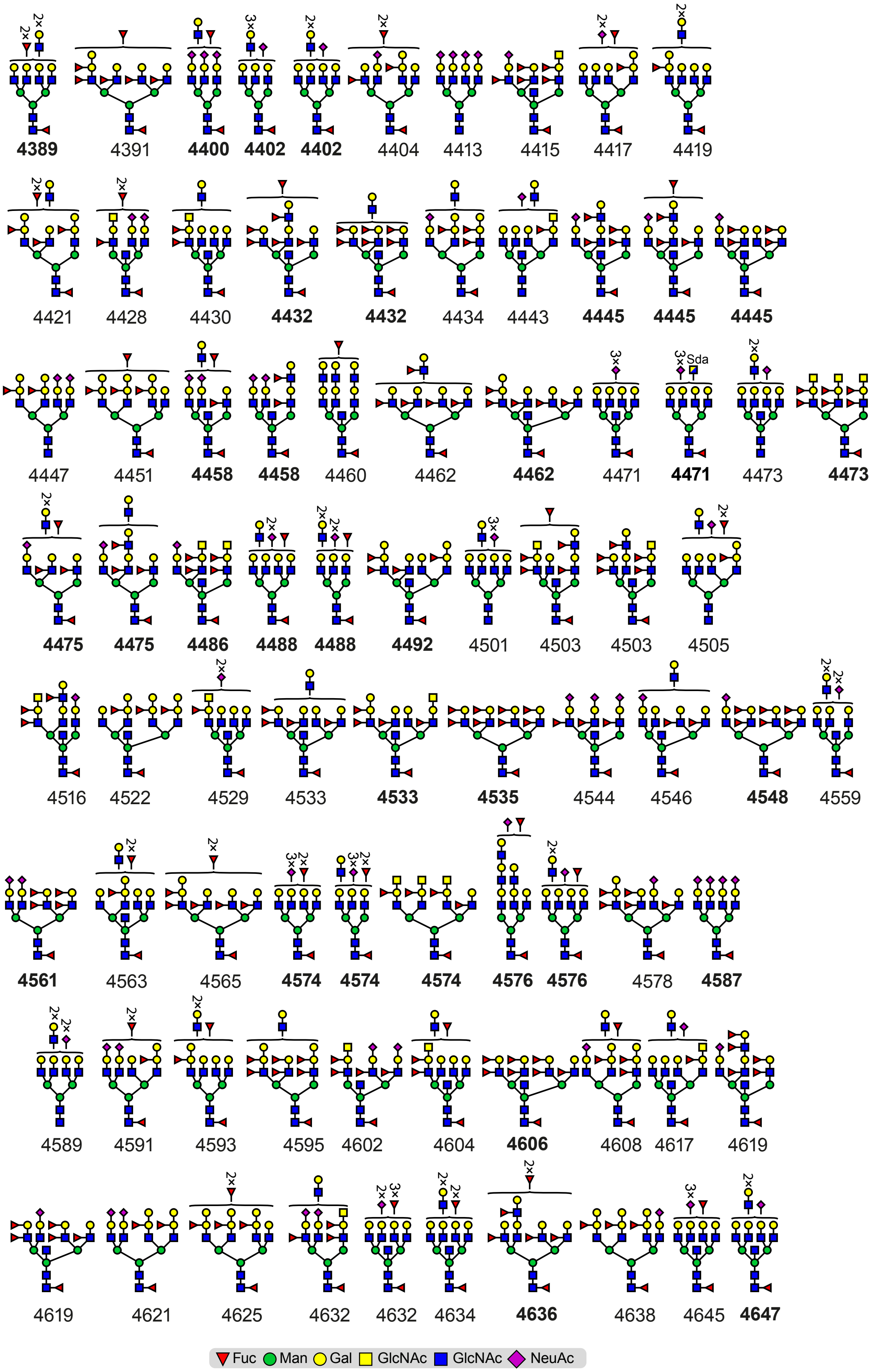
4389 4391 4400 4402 4402 4404 4413 4415 4417 4419
4421 4428 4430 4432 4432 4434 4443 4445 4445 4445
Sda
4447 4451 4458 4458 4460 4462 4462 4471 4471 4473 4473
4475 4475 4486 4488 4488 4492 4501 4503 4503 4505
4516 4522 4529 4533 4533 4535 4544 4546 4548 4559
4561 4563 4565 4574 4574 4574 4576 4576 4578 4587
4589 4591 4593 4595 4602 4604 4606 4608 4617 4619
4619 4621 4625 4632 4632 4634 4636 4638 4645 4647
Fuc
Man
Gal
GlcNAc
GlcNAc
NeuAc

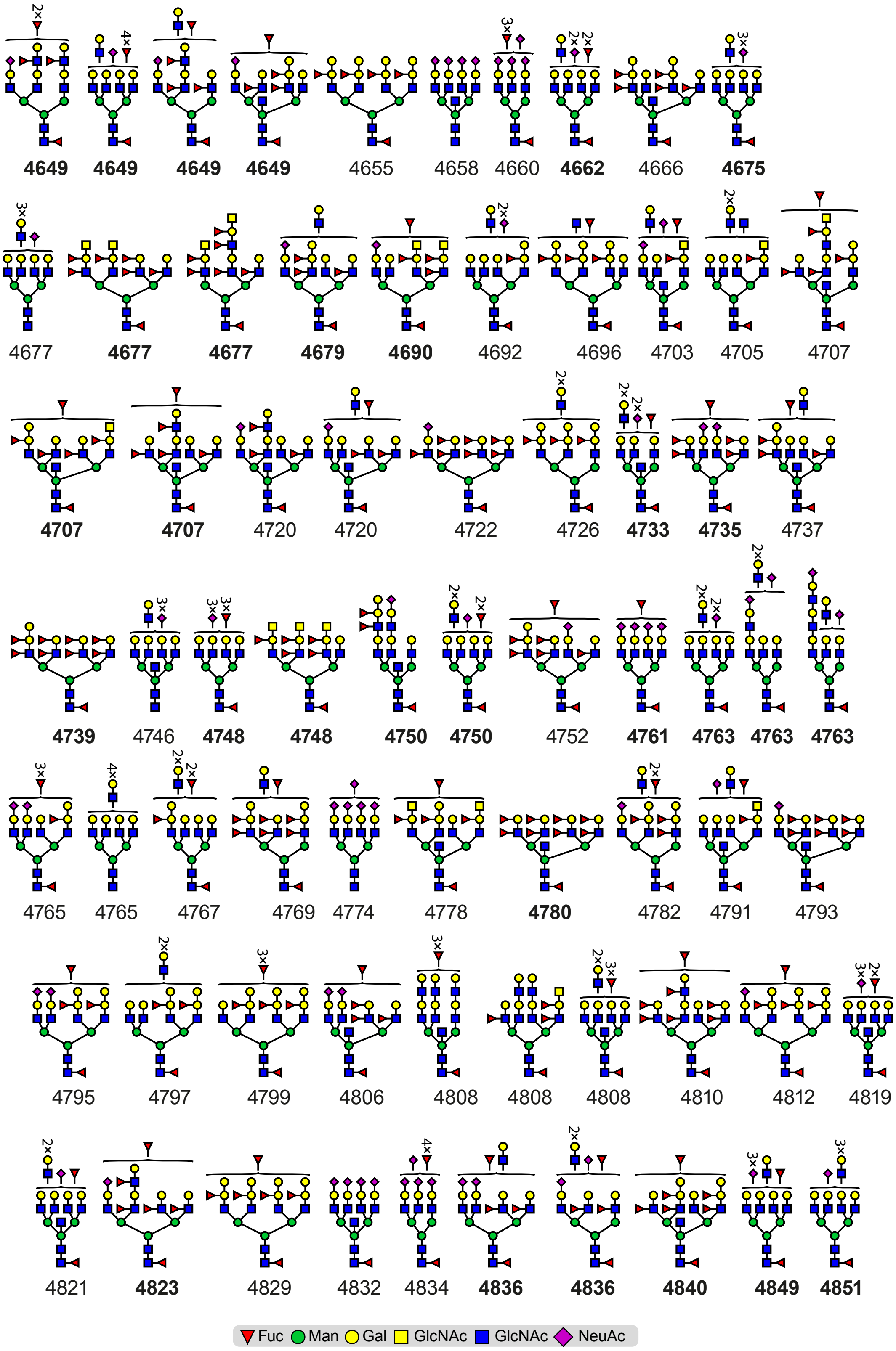
4649 4649 4649 4649 4655 4658 4660 4662 4666 4675
4677 4677 4677 4679 4690 4692 4696 4703 4705 4707
4707 4707 4720 4720 4722 4726 4733 4735 4737
4739 4746 4748 4748 4750 4750 4752 4761 4763 4763 4763
4765 4765 4767 4769 4774 4778 4780 4782 4791 4793
4795 4797 4799 4806 4808 4808 4808 4810 4812 4819
4821 4823 4829 4832 4834 4836 4836 4840 4849 4851
Fuc Man Gal GlcNAc GlcNAc NeuAc

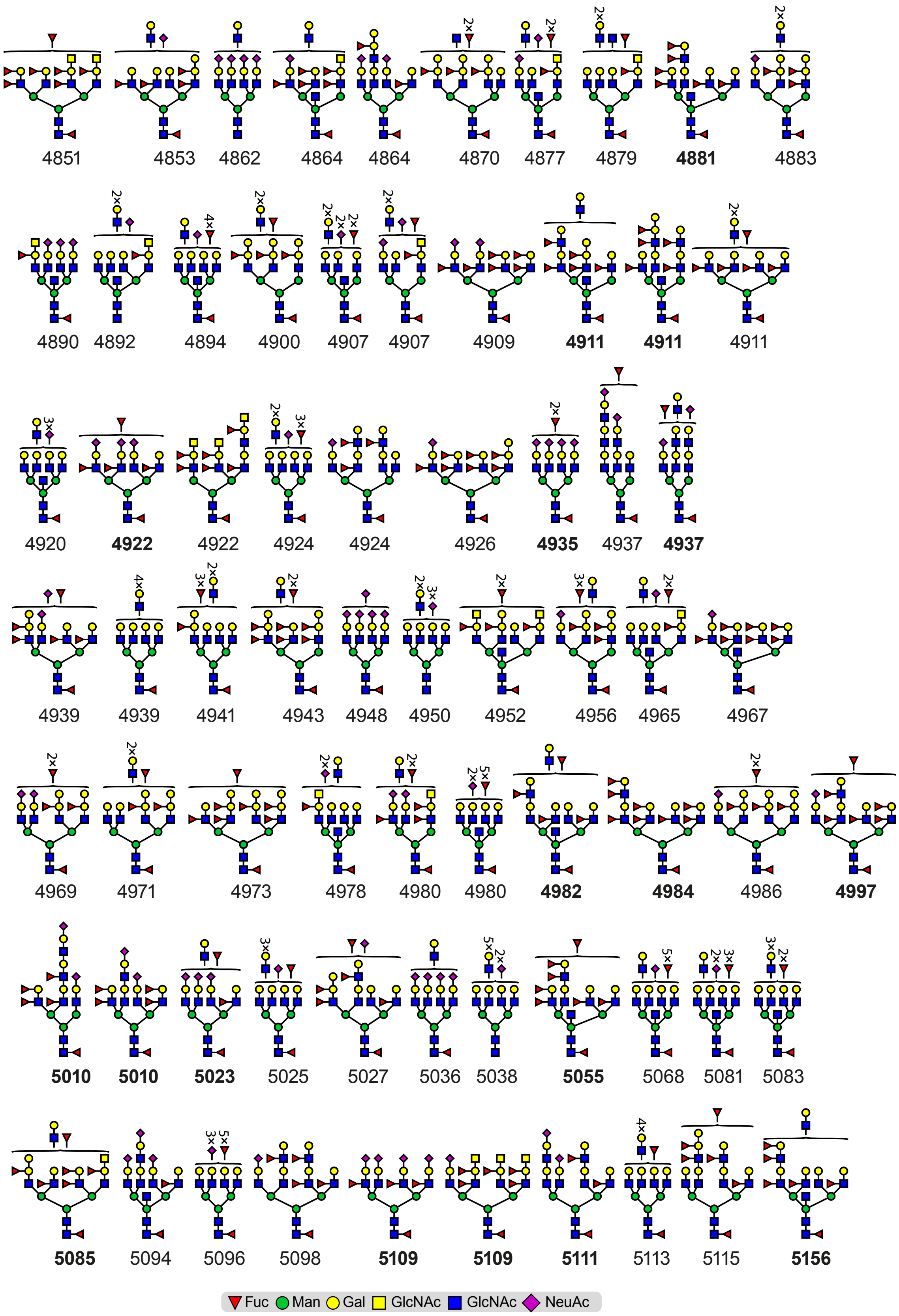
4851 4853 4862 4864 4864 4870 4877 4879 4881 4883
4890 4892 4894 4900 4907 4907 4909 4911 4911 4911
4920 4922 4922 4924 4924 4926 4935 4937 4937
4939 4939 4941 4943 4948 4950 4952 4956 4965 4967
4969 4971 4973 4978 4980 4980 4982 4984 4986 4997
5010 5010 5023 5025 5027 5036 5038 5055 5068 5081 5083
5085 5094 5096 5098 5109 5109 5111 5113 5115 5156
Fuc Man Gal GlcNAc GlcNAc NeuAc

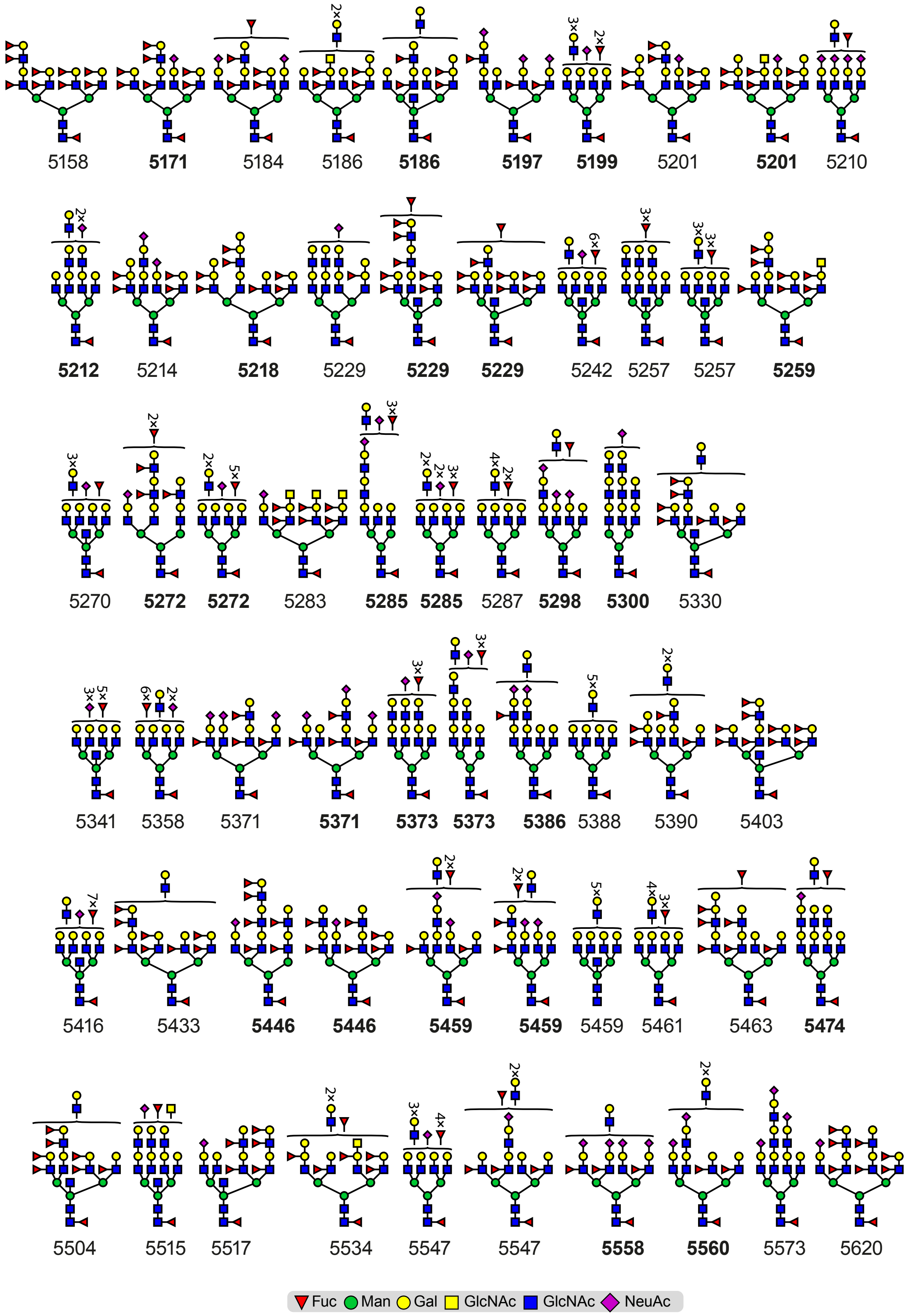

5158 5171 5184 5186 5186 5197 5199 5201 5201 5210
5212 5214 5218 5229 5229 5229 5242 5257 5257 5259
5270 5272 5272 5283 5285 5285 5287 5298 5300 5330
5341 5358 5371 5371 5373 5373 5386 5388 5390 5403
5416 5433 5446 5446 5459 5459 5459 5461 5463 5474
5504 5515 5517 5534 5547 5547 5558 5560 5573 5620
Fuc Man Gal GlcNAc GlcNAc NeuAc

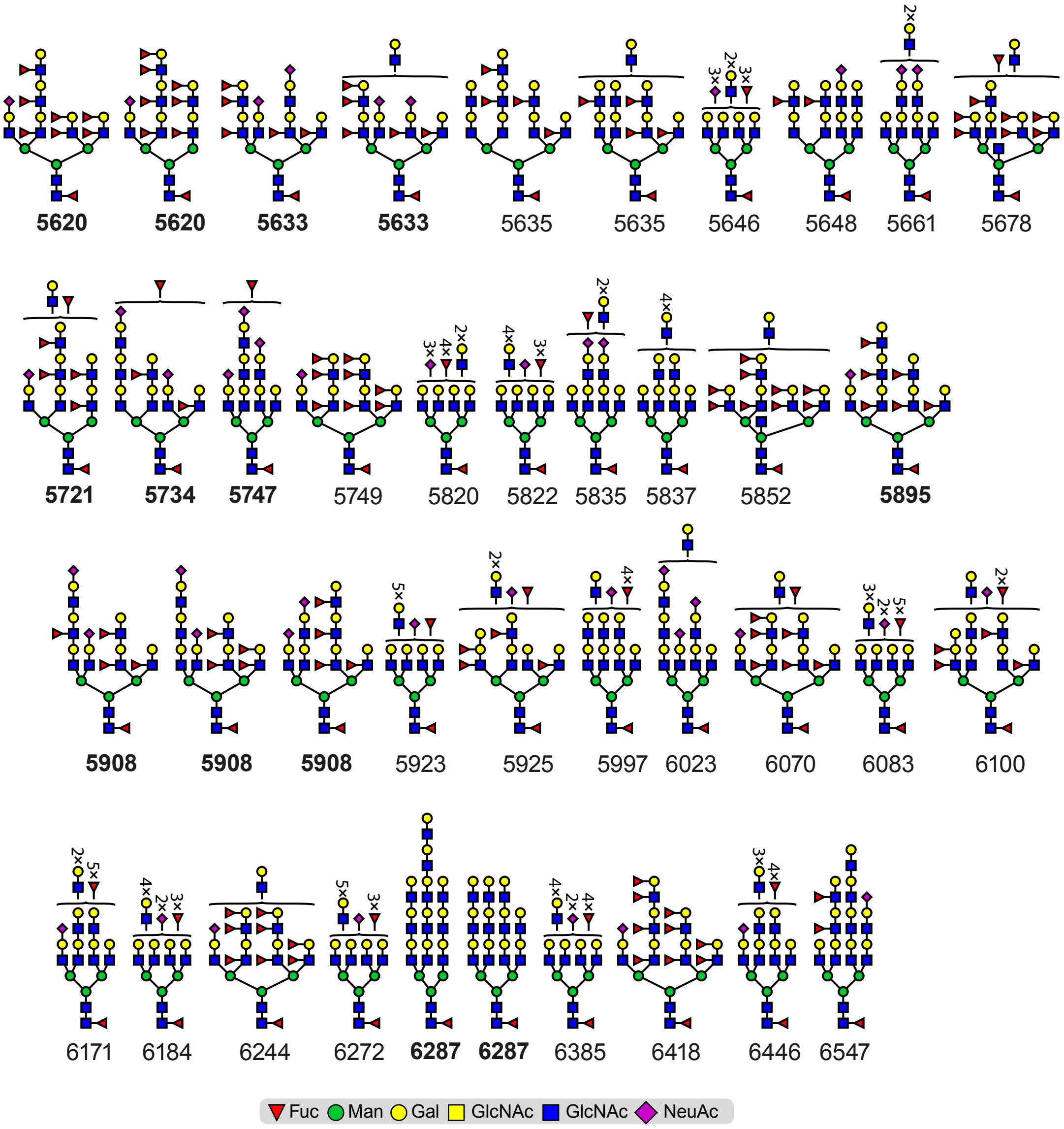
5620 5620 5633 5633 5635 5635 5646 5648 5661 5678
5721 5734 5747 5749 5820 5822 5835 5837 5852 5895
5908 5908 5908 5923 5925 5997 6023 6070 6083 6100
6171 6184 6244 6272 6287 6287 6385 6418 6446 6547
Fuc Man Gal GlcNAc GlcNAc NeuAc

**Figure S5**

**N-glycan structure library derived from primary tracheal and nasal epithelial cells from human donors.** Putative structures are based on composition, biosynthetic knowledge, and tandem mass spectrometry where available. Cartoon structures were drawn according to the Symbol Nomenclature for Glycans (SNFG) (Neelamegham et al., 2019 Glycobiology 29, 620-624) guidelines. Value under each putative structure corresponds to the molecular ion $[M+Na]^+$ m/z value detected in the corresponding Figures S1, S2, S8, S10, and S13. Values in bold correspond to a detected structure subjected to MALDI-TOF/TOF MS/MS analysis. The antenna position of the LacNAcs has not been determined. Because of the high complexity of the trachea N-glycome, this library does not include all possible structures. It is possible that molecular ions with a certain composition to correspond to either structural isomers containing blood groups A and B, or structural isomers without blood groups but with an extra LAcNAc unit.

Examples of structures corresponding to N-linked glycans with a total number of four (4) LacNAcs and various fucose residues (values in parenthesis correspond to number of fucose residues): Molecular ions at m/z 2968 (0), 3142 (1), 3316 (2), 3490 (3), 3664 (4), 3838 (5), 4013 (6), 4187 (7), 4361 (8) and 4535 (9). Examples of structures corresponding to N-linked glycans with a total number of five (5) LacNAcs and various fucose residues: Molecular ions at m/z 3417 (0), 3591 (1), 3765 (2), 3939 (3), 4288 (5), 4462 (6), 4636 (7), 4810 (8), 4984 (9). Examples of structures corresponding to N-linked glycans with a total number of six (6) LacNAcs and various fucose residues: Molecular ions at m/z 3866 (0), 4041 (1), 4215 (2), 4389 (3), 4563 (4), 4737 (5), 4911 (6), 5085 (7), 5259 (8) and 5433 (9). Examples of structures containing LeX and/or LeY epitopes: Molecular ions at m/z 3402, 3460,

3563, 3634, 3737, 3808, 3838, 3983, 4084, 4114, 4187, 4200, 4258, 4359 and 4361. Examples of structures containing both sialylated and fucosylated N-linked glycans: Molecular ions at 3750, 3763, 3937, 4097, 4213, 4271, 4299, 4387. Examples of structures containing blood groups: Molecular ions at m/z 3198, 3256, 3327, 3402, 3490 and 3593. For MALDI-TOF/TOF MS/MS analysis of structures containing the above epitopes, see **Figure S3**.

**Figure S6**

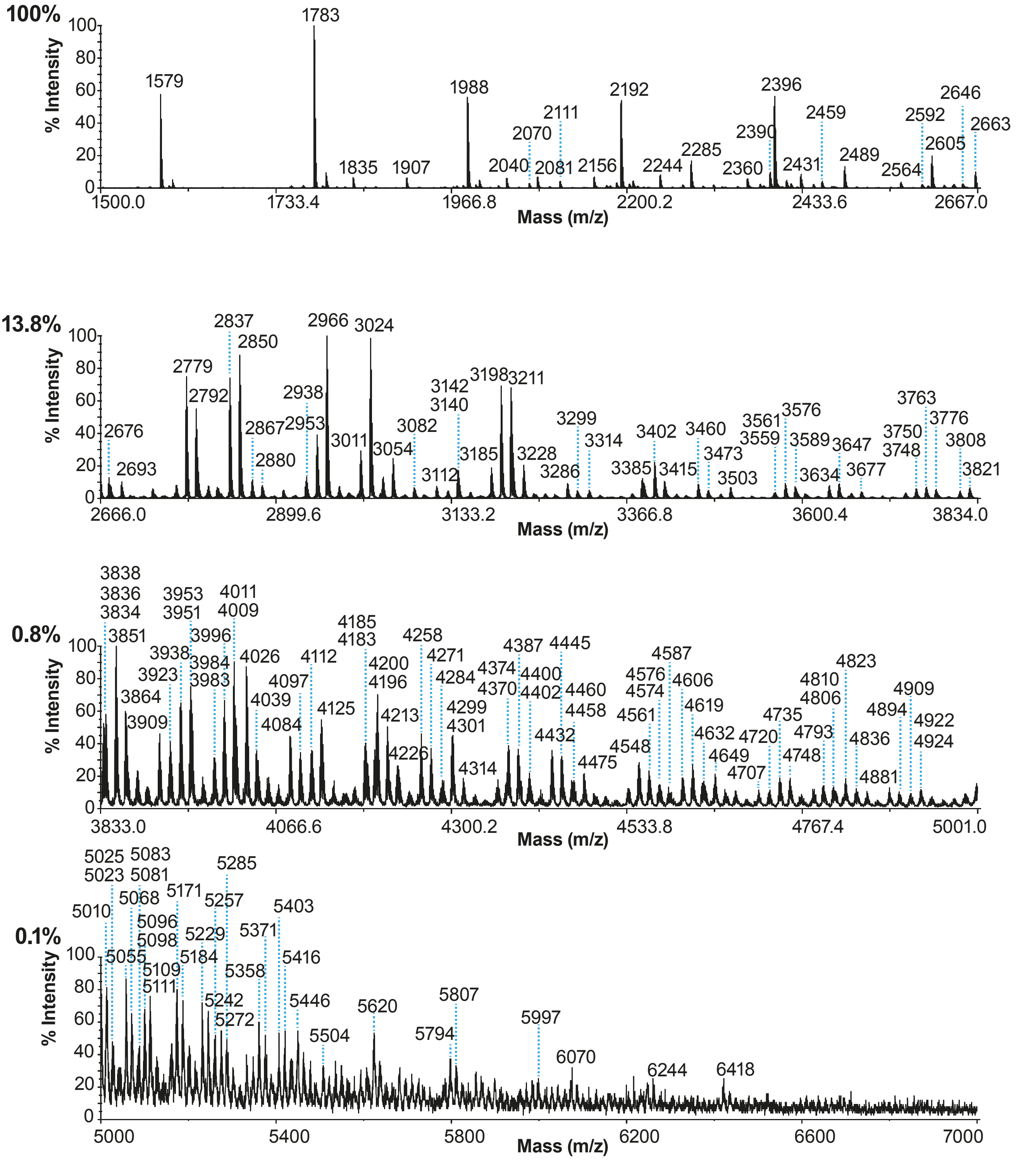
100%
% Intensity
Mass (m/z)
1500.0
1733.4
1966.8
2200.2
2433.6
2667.0
1579
1783
1835
1907
1988
2040
2070
2081
2111
2156
2192
2244
2285
2360
2390
2396
2431
2459
2489
2564
2592
2605
2646
2663
13.8%
2666.0
2899.6
3133.2
3366.8
3600.4
3834.0
0.8%
3833.0
4066.6
4300.2
4533.8
4767.4
5001.0
0.1%
5000
5400
5800
6200
6600
7000

**Figure S6**

**MALDI-TOF mass spectra of permethylated N-glycans derived from primary nasal epithelial cells from human donors.** MALDI-TOF MS analysis of permethylated N-linked glycans from epithelial cells of human nasal samples pooled from healthy donors. All molecular ions are $[M+Na]^+$. Numbers above peaks correspond to the molecular ions identified based on manual annotation (see Methods details). “M” or “m” correspond to major or minor relative abundance of the corresponding molecular ion relative to the molecular ion found in the same molecular ion cluster. Percentages on top left of each panel correspond to the relative percentage of the maximum peak of the corresponding panel compared to the relative intensity of the maximum peak of the top panel (based on the total ion current of each panel, without de-isotoping). Full structure annotations can be found in **Figure S5**.

Figure S7

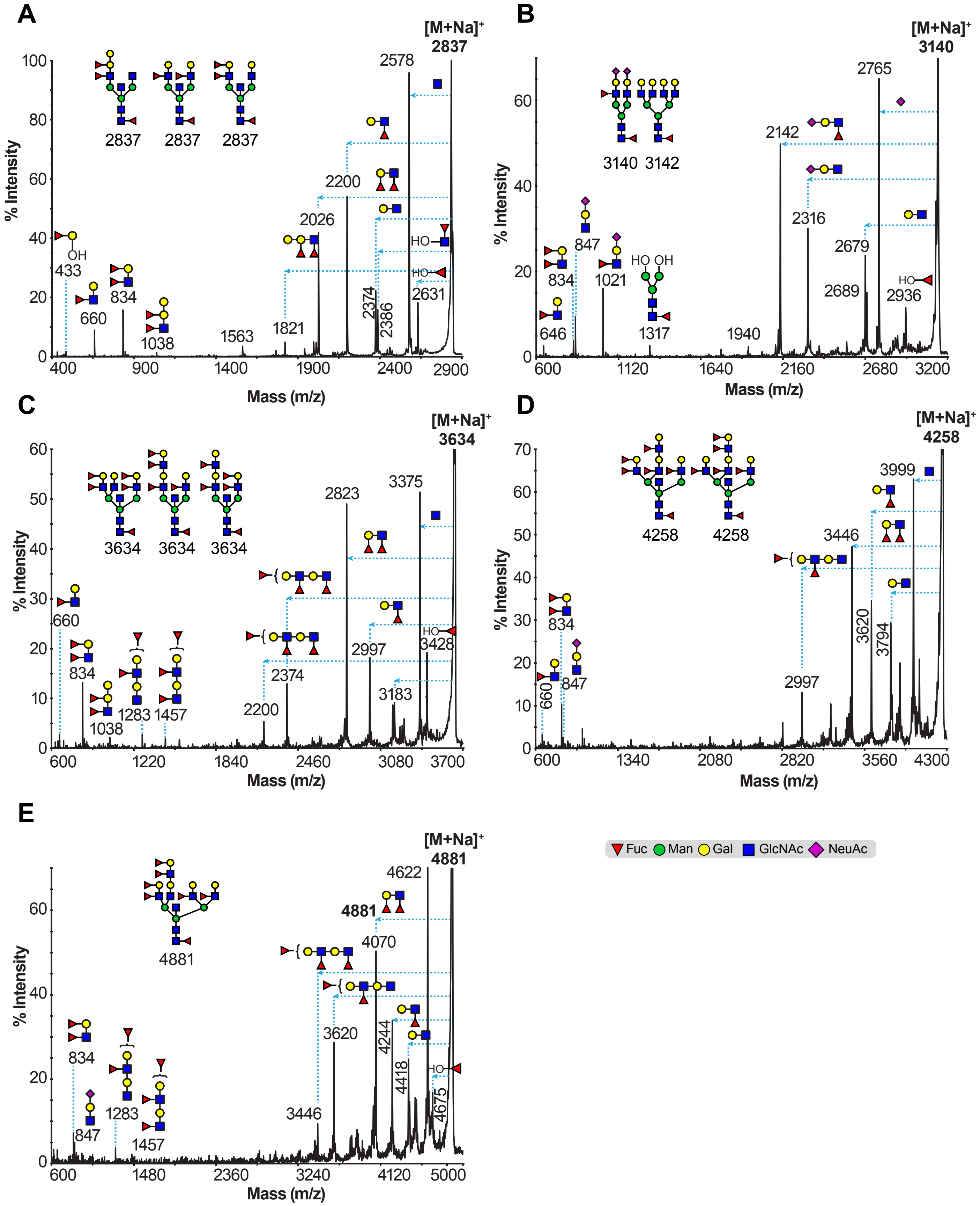

A
B
C
D
E
[M+Na]+ 2837
[M+Na]+ 3140
[M+Na]+ 3634
[M+Na]+ 4258
[M+Na]+ 4881
% Intensity
Mass (m/z)
2837
2578
2200
2026
1821
1563
1038
834
660
433
2374
2386
2631
3140
3142
2765
2142
2316
2679
2689
2936
1940
1317
1021
847
646
3375
2823
2997
3428
3183
2200
1283
1457
3999
3446
3620
3794
2997
4622
4070
3620
4244
4418
4675
3446
HO
OH
Fuc
Man
Gal
GlcNAc
NeuAc

**Figure S7**

**MALDI-TOF/TOF MS/MS analysis of permethylated N-linked glycans from primary nasal epithelial cells from human donors.** Molecular ions $[M+Na]^+$ present in **Figure S6** were selected for MALDI-TOF/TOF MS/MS analysis. (A) m/z 2837, (B) m/z 3140, (C) m/z 3634, (D) m/z 4258, (E) m/z 4881. Cartoon structures in the corresponding spectra do not represent all possible structural isomers, for clarity. Horizontal dashed lines correspond to the indicated fragment loss from the corresponding molecular ion $[M+Na]^+$. Vertical dashed lines indicate the m/z fragment peak that corresponds to the fragment loss as shown from the corresponding horizontal dashed line. Structures outside the bracket have not unequivocally been defined. For clarity, not all fragments were annotated.

### Figure S8

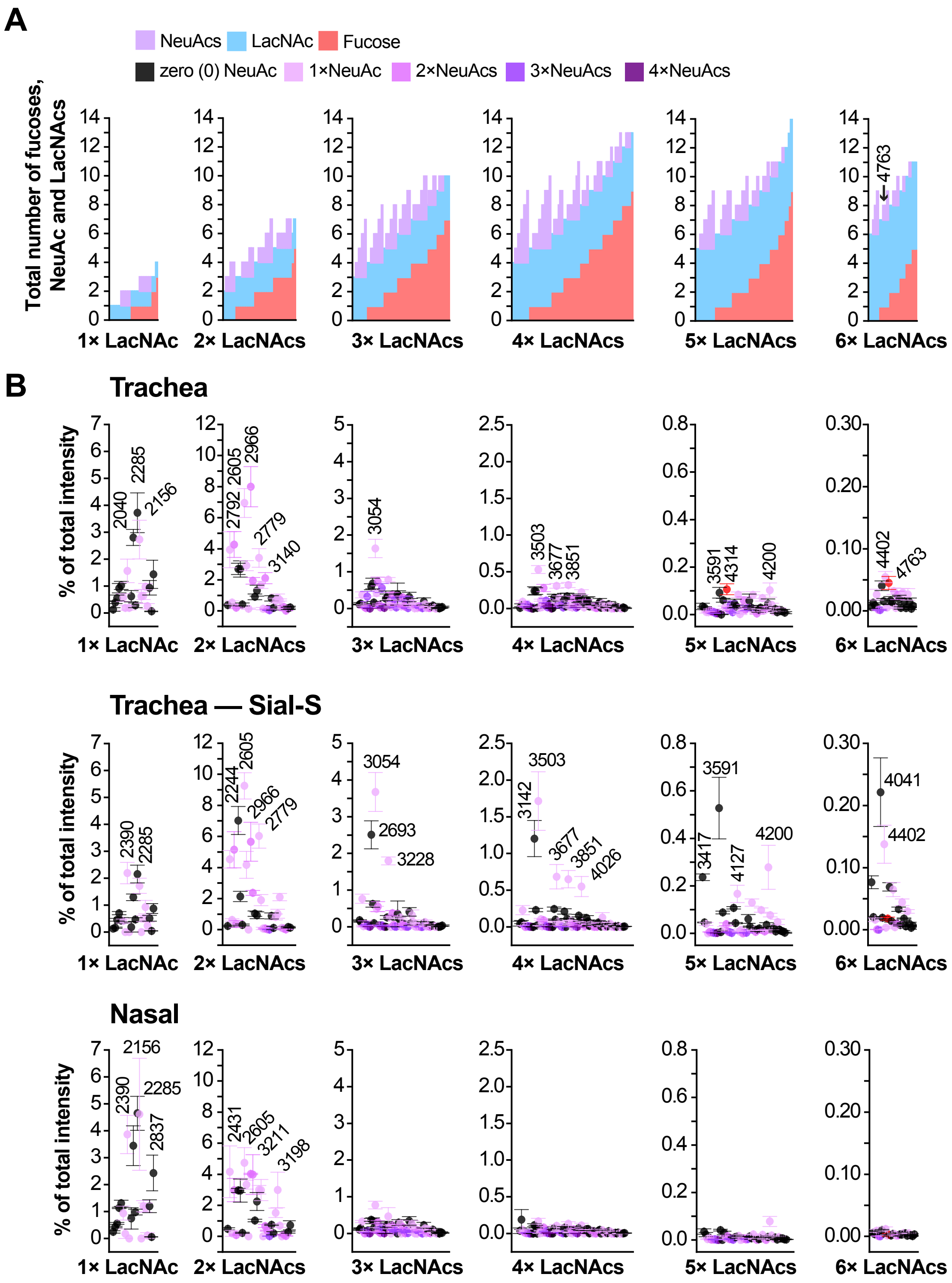
A
NeuAcs
LacNAc
Fucose
zero (0) NeuAc
1×NeuAc
2×NeuAcs
3×NeuAcs
4×NeuAcs
Total number of fucoses, NeuAc and LacNAcs
4763
1× LacNAc
2× LacNAcs
3× LacNAcs
4× LacNAcs
5× LacNAcs
6× LacNAcs
B
Trachea
% of total intensity
2040
2285
2156
2792
2605
2966
2779
3140
3054
3503
3677
3851
3591
4314
4200
4402
4763
Trachea — Sial-S
2390
2285
2244
2605
2966
2779
3054
2693
3228
3142
3503
3677
3851
4026
3591
3417
4127
4200
4041
4402
Nasal
2156
2390
2285
2837
2431
2605
3211
3198

**Figure S8**

**Human influenza H3 receptor N-linked glycan has low relative abundance on primary tracheal epithelial cells from human donors and almost absent in nasal epithelial cells.** (A) Stacked bar columns depicting the compositions of N-linked glycans of trachea samples ranging from a single to six LacNAc repeating units (blue columns); fucose (red columns) and NeuAc (purple columns) residues. Compositions are sorted by increasing number of fucose residues and then by increasing number of NeuAcs residues. (B) Relative abundance of N-linked glycans as percentage (%) of the total relative abundance after de-isotoping (see Methods details) of trachea samples before or after Sial-S digestion, and nasal epithelial cells respectively. Note in (A) that as the number of antennas fucosylation is increasing, the possible number of NeuAc residues is decreasing. For instance, for N-linked glycans with five (5) total LacNAcs, the total number of antennas fucosylation (minus the core fucose) can reach even of eight (8) residues, limiting the number of putative influenza receptors in the epithelial cells of trachea and nasal human samples. Numbers next to the error bars correspond to m/z values found in **Figure S5**. Note that the molecular ion at m/z 4763, a putative Influenza H3 receptor, has a mean relative abundance of 0.045% in the trachea samples, 0.018% in the trachea samples after Sial-S digestion and 0.003% in nasal samples. Error bars correspond to mean value ± standard error of the mean. Values next to the error bars correspond to m/z values shown in **Figure S5**. Results are from biological replicates of trachea (n=9), trachea Sial-S (n=3) and nasal samples (n=3).

Concerning the relation of antenna fucosylation and NeuAc residues depending on the number of total LacNAc units (A): For N-linked glycans with a total number of three (3), four (4) and five (5) LacNAc on their antennas, when the total number of fucosylation was restricted to zero (0) or one (1) residue, then the number of NeuAc residues could reach up to five (5) residues. Reversely, when the total number of fucosylation was five (5) residues or higher, then the NeuAc residues were restricted to up to two (2) NeuAc residues.

**Figure S9**

**A**

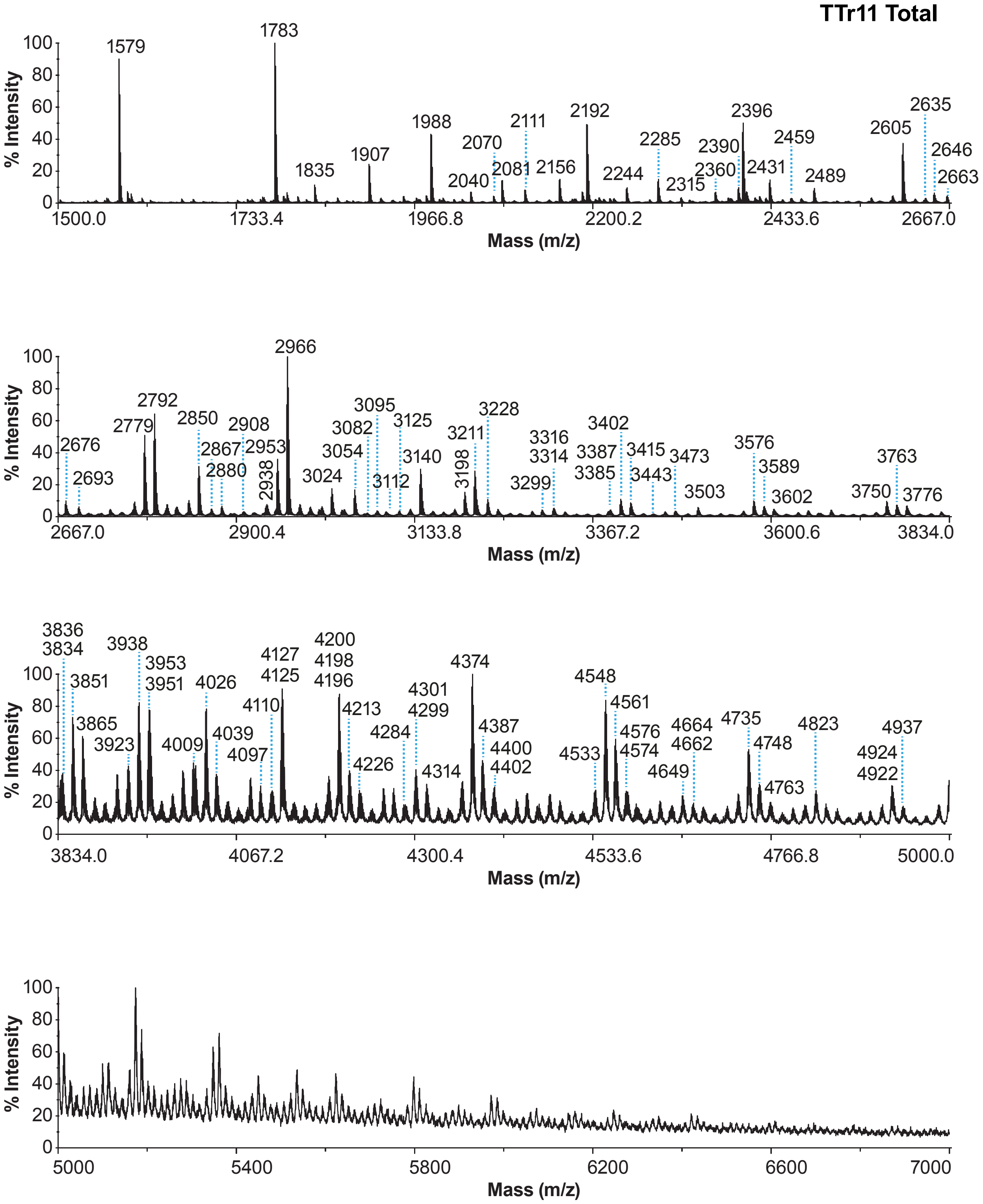

TTr11 Total
% Intensity
Mass (m/z)
100
80
60
40
20
0
1500.0
1733.4
1966.8
2200.2
2433.6
2667.0
1579
1783
1835
1907
1988
2040
2070
2081
2111
2156
2192
2244
2285
2315
2360
2390
2396
2431
2459
2489
2605
2635
2646
2663
2667.0
2900.4
3133.8
3367.2
3600.6
3834.0
2676
2693
2779
2792
2850
2867
2880
2908
2938
2953
2966
3024
3054
3082
3095
3112
3125
3140
3198
3211
3228
3299
3316
3314
3387
3385
3402
3415
3443
3473
3503
3576
3589
3602
3750
3763
3776
3834.0
4067.2
4300.4
4533.6
4766.8
5000.0
3836
3834
3851
3865
3923
3938
3953
3951
4009
4026
4039
4097
4110
4127
4125
4200
4198
4196
4213
4226
4284
4301
4299
4314
4374
4387
4400
4402
4533
4548
4561
4576
4574
4649
4664
4662
4735
4748
4763
4823
4924
4922
4937
5000
5400
5800
6200
6600
7000

B

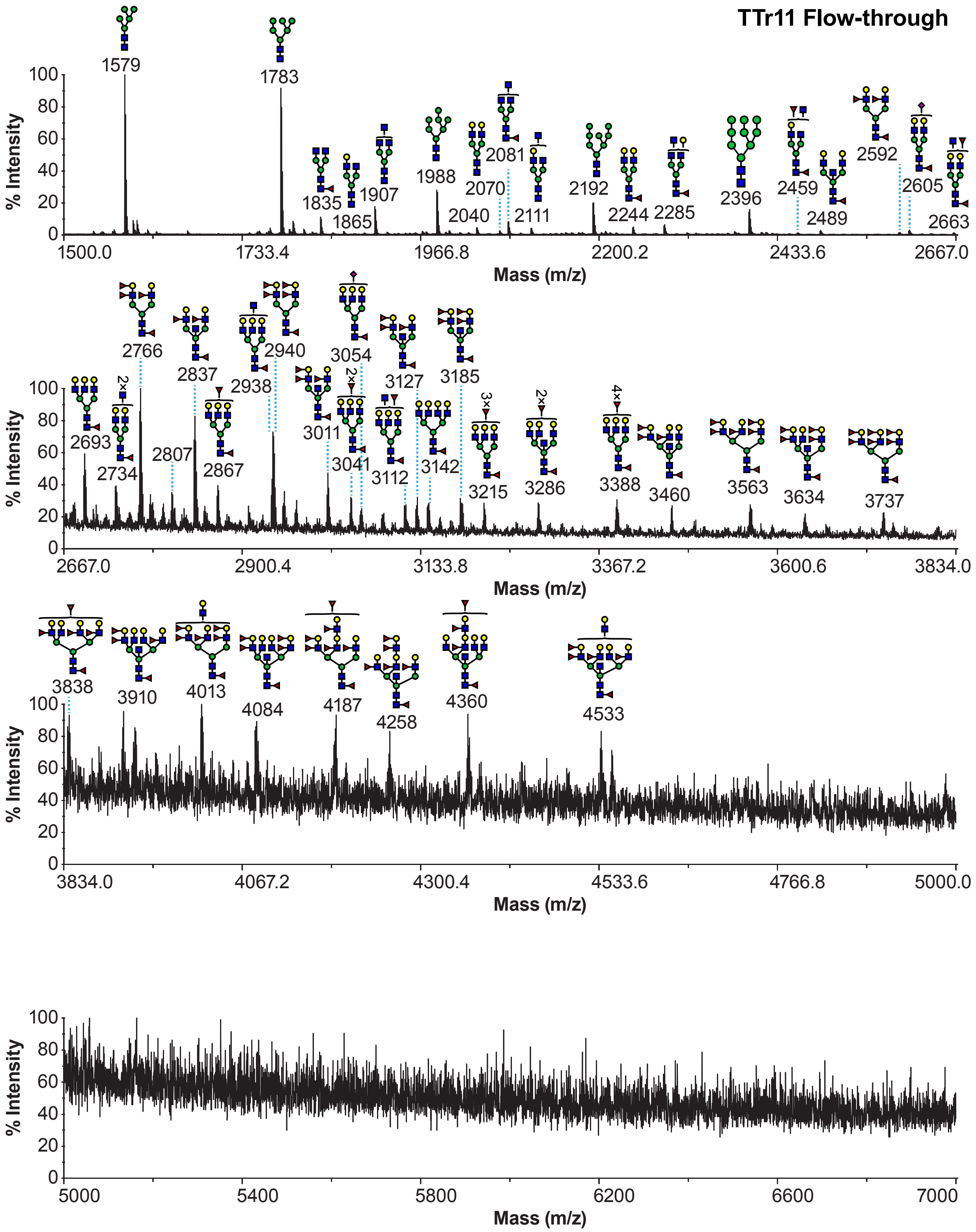
TTr11 Flow-through
1579
1783
1835
1865
1907
1988
2040
2070
2081
2111
2192
2244
2285
2396
2459
2489
2592
2605
2663
% Intensity
Mass (m/z)
1500.0
1733.4
1966.8
2200.2
2433.6
2667.0
2693
2734
2766
2807
2837
2867
2938
2940
3011
3041
3054
3112
3127
3142
3185
3215
3286
3388
3460
3563
3634
3737
2×
3×
4×
2667.0
2900.4
3133.8
3367.2
3600.6
3834.0
3838
3910
4013
4084
4187
4258
4360
4533
3834.0
4067.2
4300.4
4533.6
4766.8
5000.0
5000
5400
5800
6200
6600
7000

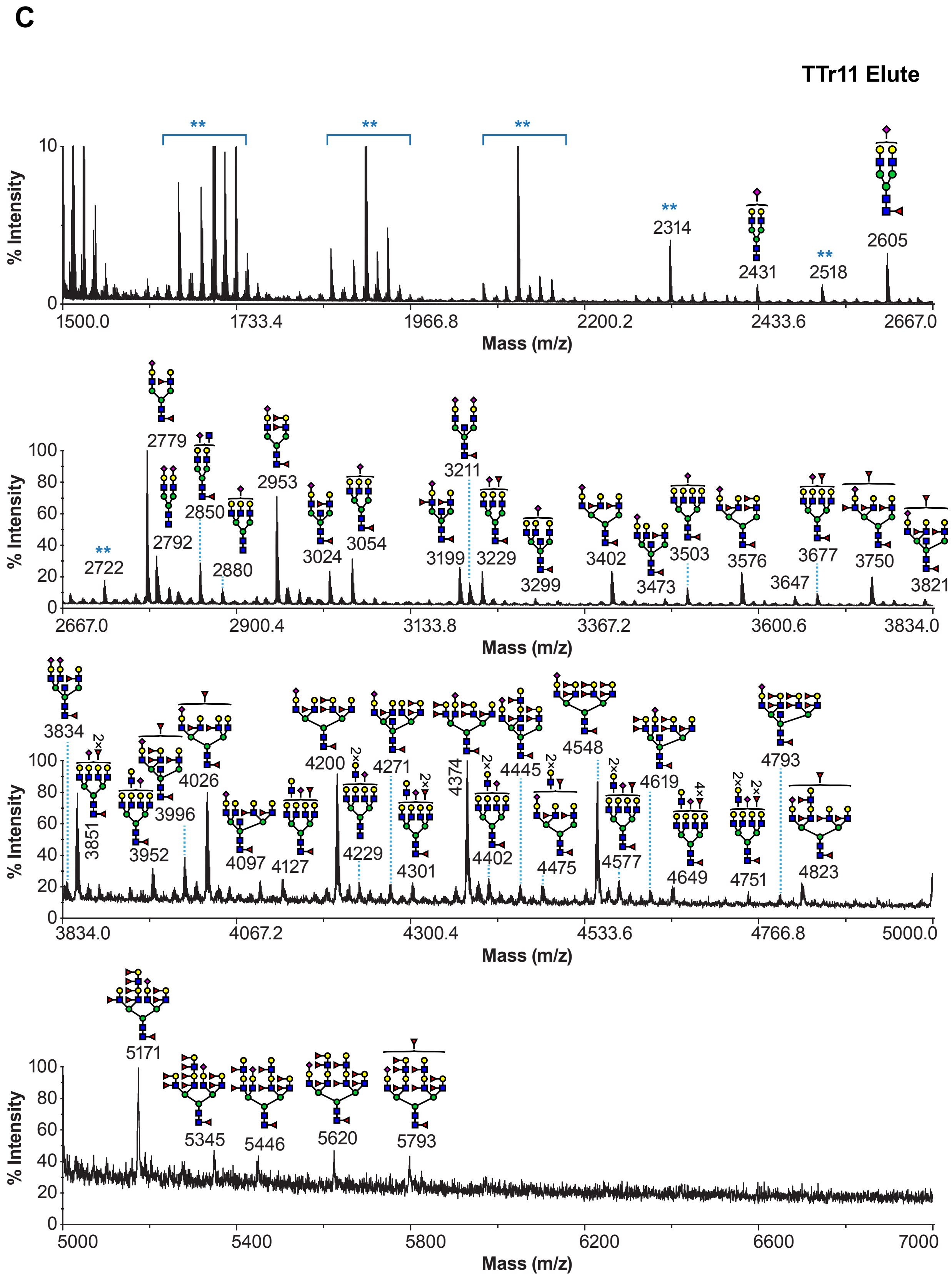

C
TTr11 Elute
% Intensity
Mass (m/z)
**
2314
2431
2518
2605
1500.0
1733.4
1966.8
2200.2
2433.6
2667.0
2722
2779
2792
2850
2880
2953
3024
3054
3199
3211
3229
3299
3402
3473
3503
3576
3647
3677
3750
3821
2900.4
3133.8
3367.2
3600.6
3834.0
3834
3851
3952
3996
4026
4097
4127
4200
4229
4271
4301
4374
4402
4445
4475
4548
4577
4619
4649
4751
4793
4823
4067.2
4300.4
4533.6
4766.8
5000.0
5171
5345
5446
5620
5793
5000
5400
5800
6200
6600
7000

**Figure S9**

**MALDI-TOF mass spectra of N-linked glycans enriched with an SNA column.**

MALDI-TOF MS of permethylated N-linked glycans derived from primary tracheal epithelial cells from human donors (TTr11) (A) before SNA column (total), (B) flow-through and (C) eluted fraction. All molecular ions are $[M+Na]^+$. (A) Full structure annotations can be found in Figure S3. (B) Structures outside the bracket have not unequivocally been defined. (C) Peaks with blue asterisks correspond to hexose contamination. Cartoon structures were drawn according to the Symbol Nomenclature for Glycans (SNFG) (Neelamegham et al., 2019 Glycobiology 29, 620-624) guidelines. Numbers above peaks correspond to the molecular ions identified based on manual annotation. These values were verified by the de-isotoping algorithm (see Methods details). Y-axis in top panel has been scaled to 10% of maximum relative intensity.

#### Figure S10

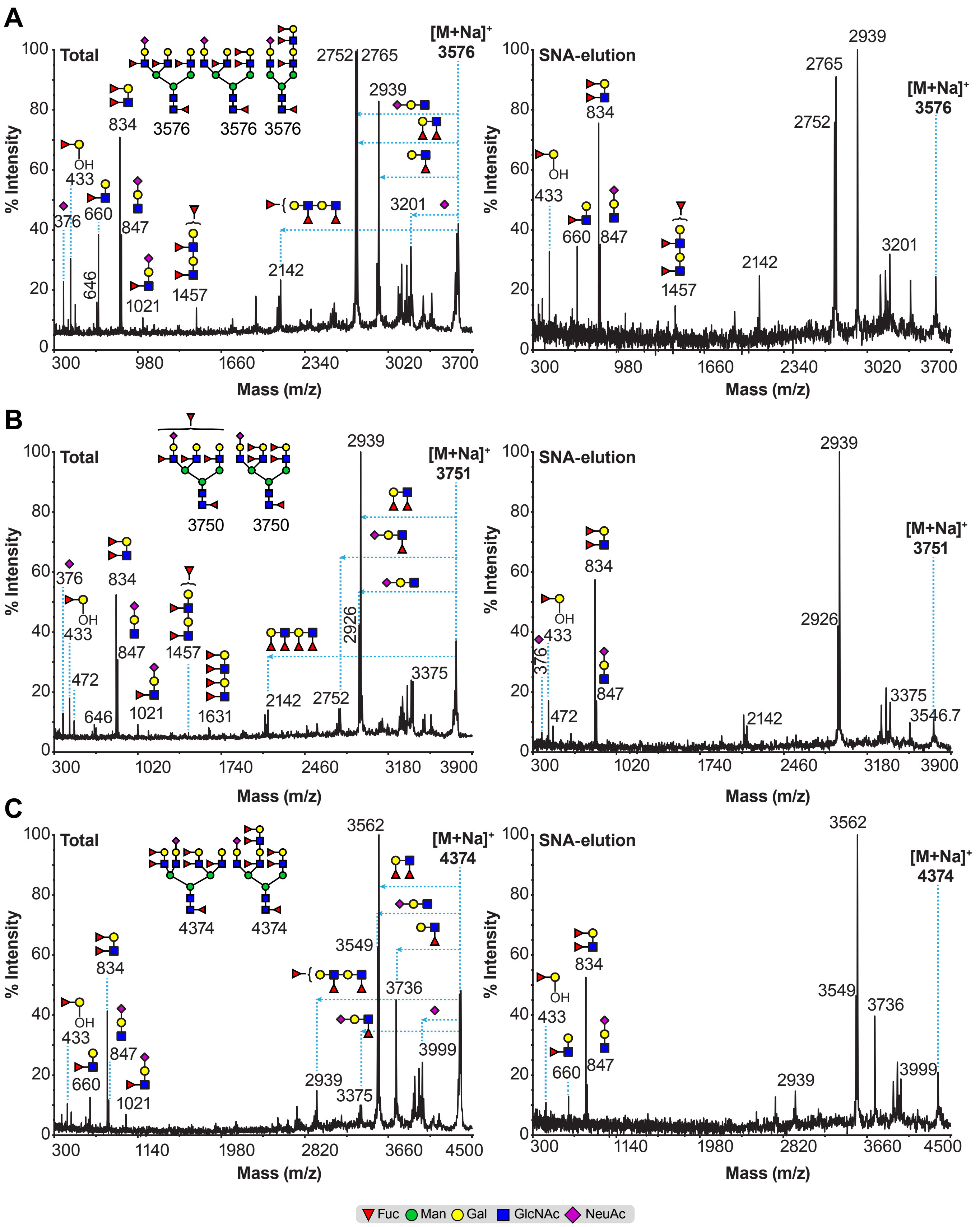
A
Total
SNA-elution
[M+Na]+ 3576
% Intensity
Mass (m/z)
B
Total
SNA-elution
[M+Na]+ 3751
C
Total
SNA-elution
[M+Na]+ 4374
Fuc Man Gal GlcNAc NeuAc

**Figure S10**

**MALDI-TOF/TOF MS/MS analysis of permethylated N-linked glycans from SNA column.** Molecular ions $[M+Na]^+$ detected in TTr11 human tracheal (**Figure S9**) sample before (left panels) or after SNA-elution (right panel) were selected for MALDI-TOF/TOF MS/MS analysis. (A) m/z 3576, (B) m/z 3751 and (C) m/z 4374. Horizontal dashed lines correspond to the indicated fragment losses from each of the corresponding molecular ion $[M+Na]^+$. Vertical dashed lines indicate the m/z fragment peak that corresponds to the fragment loss as shown from the corresponding horizontal dashed line. Structures outside the bracket have not unequivocally been defined. For clarity, not all fragments were annotated.

Note that all MALDI-TOF/TOF MS/MS spectra derived from a N-linked glycan containing a single NeuAc residue. Left panels (before SNA column): The presence of the fragment ions at m/z 1021 on the MALDI-TOF/TOF MS/MS spectra corresponds to sLeX and/or sLeA epitopes, an indication of NeuAcα2,3-linked residue. Therefore, the molecular ions at m/z 3676, 3751 and 4374 contain various structural isomers some of which contain NeuAcα2,3-linked residue. Right panels (elution SNA column): Note the absence of the fragment ion at m/z 1021 on the MALDI-TOF/TOF MS/MS spectra. This indicated that the structural isomers that contained the NeuAcα2,3-linked residue were not captured by the SNA column. The presence of fragment ion at m/z 433 is indicative of LeB epitopes (elimination Fucose-galactose structure β1,3-linked a GlcNAc residue).

**Figure S11**

**A**

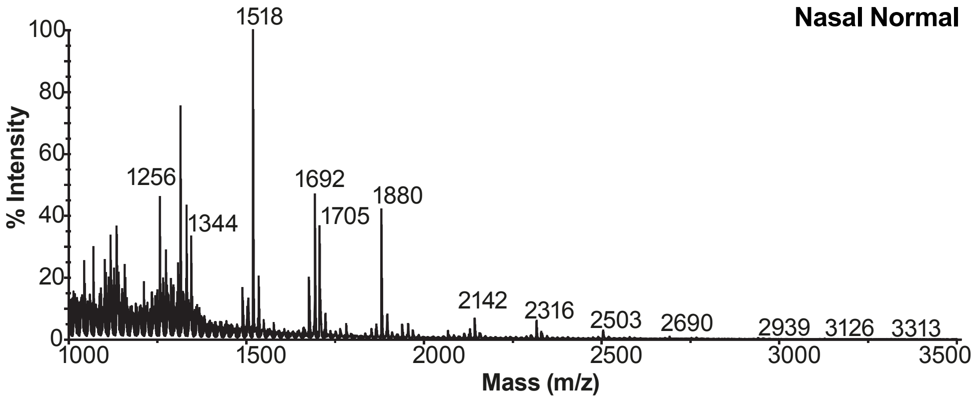


**B**

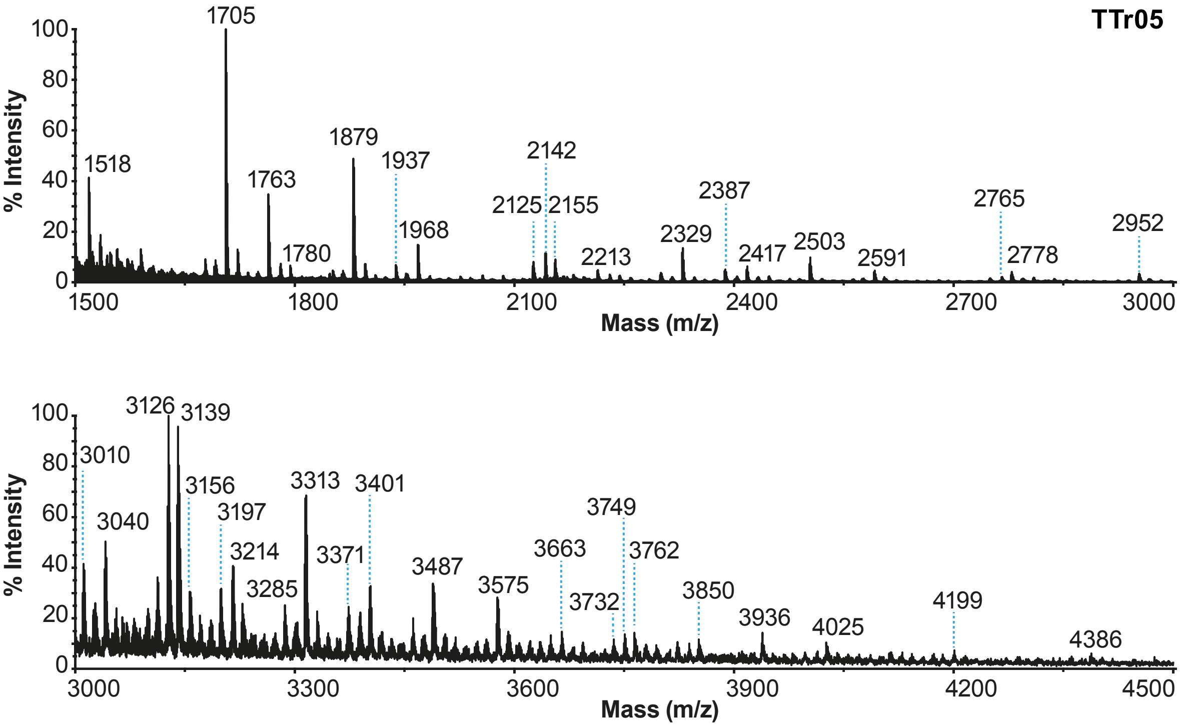

C

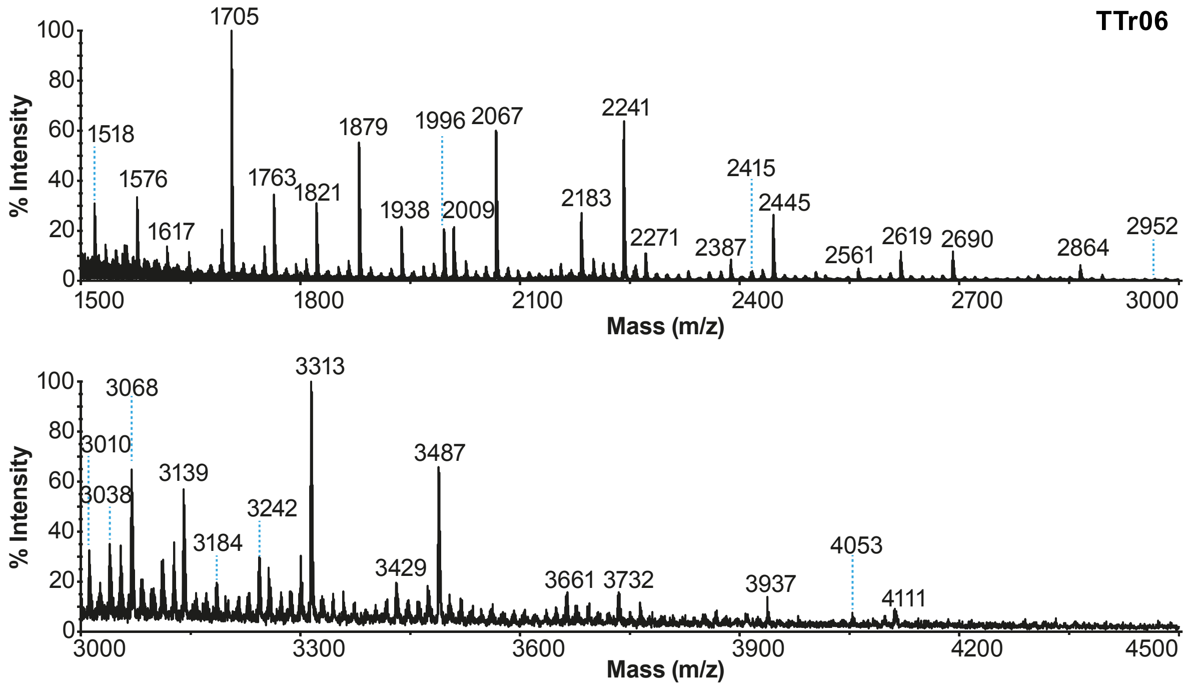
TTr06
% Intensity
Mass (m/z)
1518
1576
1617
1705
1763
1821
1879
1938
1996
2009
2067
2183
2241
2271
2387
2415
2445
2561
2619
2690
2864
2952
3010
3038
3068
3139
3184
3242
3313
3429
3487
3661
3732
3937
4053
4111

D

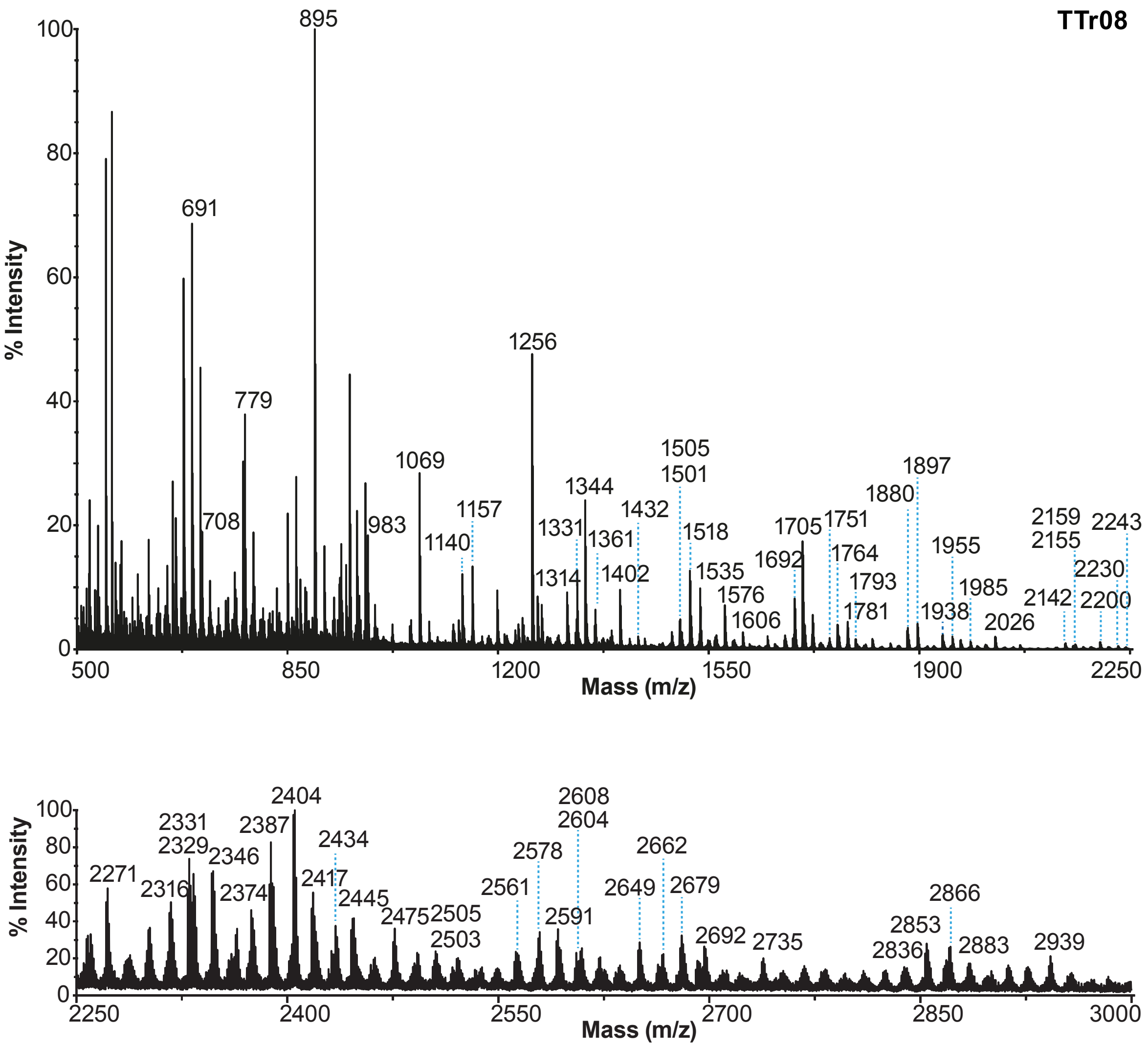
TTr08
% Intensity
Mass (m/z)
895
691
779
708
983
1069
1140
1157
1256
1314
1331
1344
1361
1402
1432
1505
1501
1518
1535
1576
1606
1692
1705
1751
1764
1781
1793
1880
1897
1938
1955
1985
2026
2142
2159
2155
2200
2230
2243
2271
2316
2329
2331
2346
2374
2387
2404
2417
2434
2445
2475
2503
2505
2561
2578
2591
2604
2608
2649
2662
2679
2692
2735
2836
2853
2866
2883
2939

E

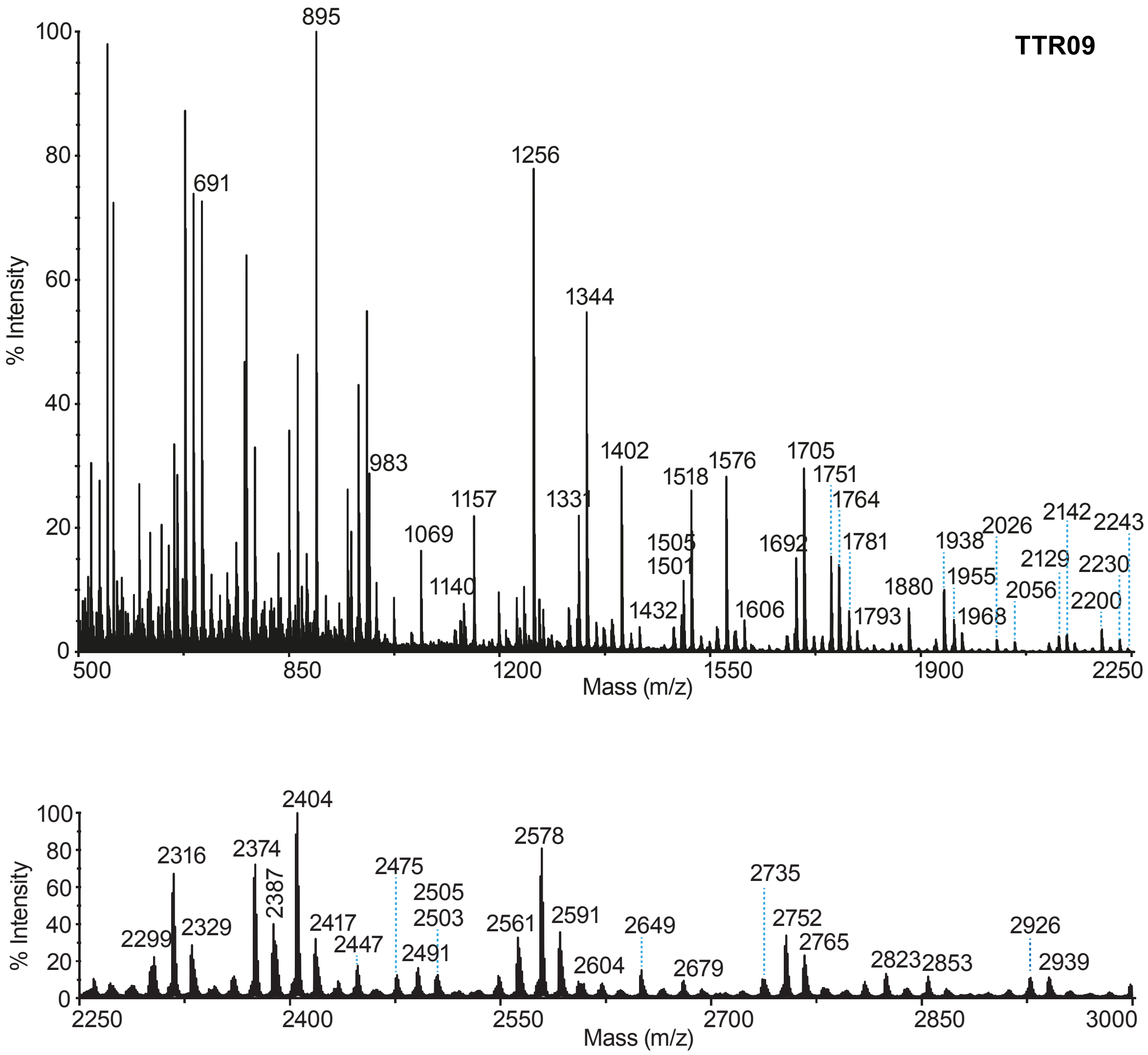

TTR09
% Intensity
Mass (m/z)
895
691
1256
1344
983
1157
1069
1140
1331
1402
1518
1576
1505
1501
1432
1606
1692
1705
1751
1764
1781
1793
1880
1938
1955
1968
2026
2056
2129
2142
2200
2230
2243
0
20
40
60
80
100
500
850
1200
1550
1900
2250
% Intensity
Mass (m/z)
2404
2316
2374
2387
2299
2329
2417
2447
2475
2505
2503
2491
2561
2578
2591
2604
2649
2679
2735
2752
2765
2823
2853
2926
2939
2400
2550
2700
2850
3000

F

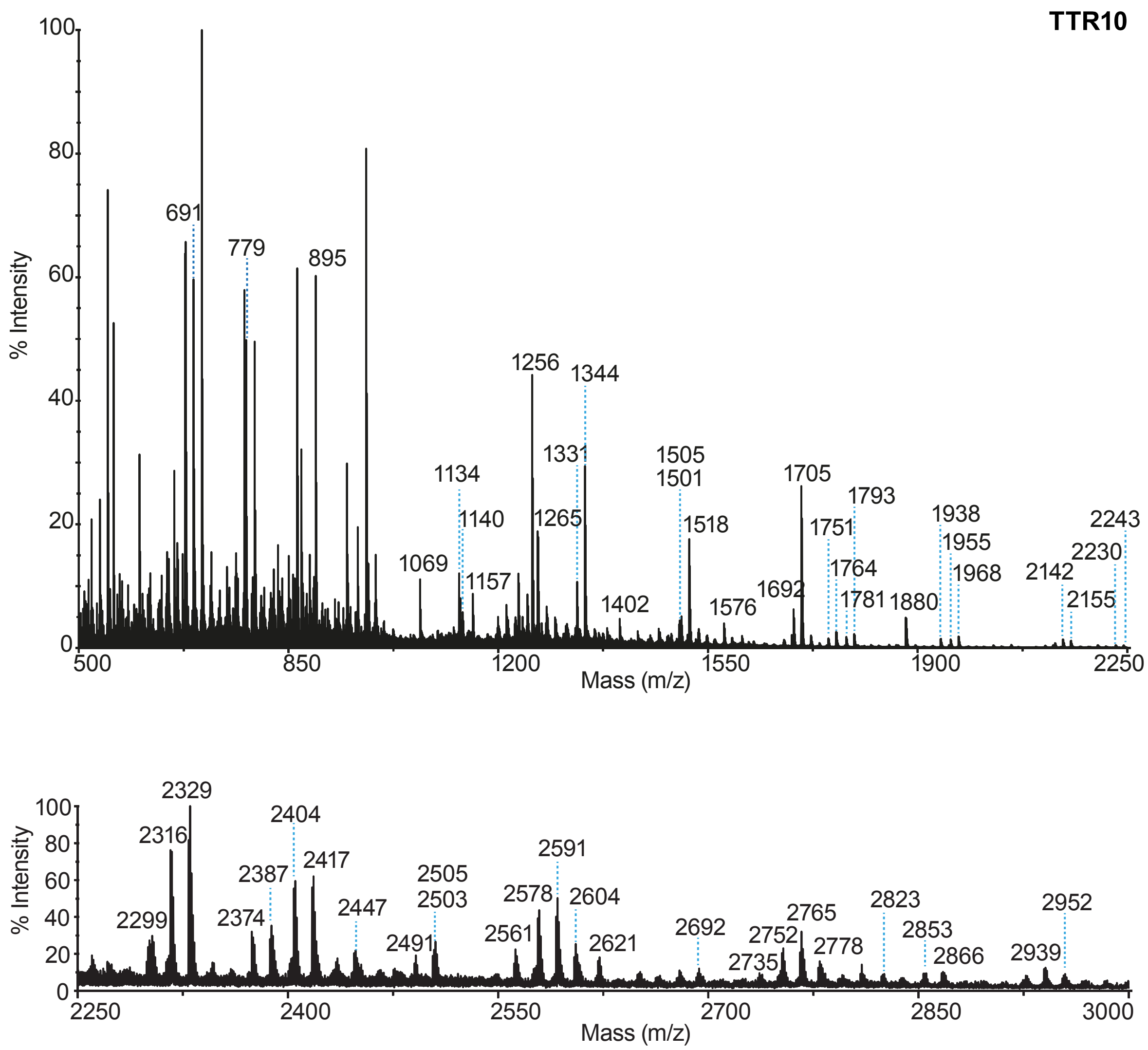
TTR10
% Intensity
Mass (m/z)
691
779
895
1069
1134
1140
1157
1256
1265
1331
1344
1402
1505
1501
1518
1576
1692
1705
1751
1764
1781
1793
1880
1938
1955
1968
2142
2155
2230
2243
2299
2316
2329
2374
2387
2404
2417
2447
2491
2503
2505
2561
2578
2591
2604
2621
2692
2735
2752
2765
2778
2823
2853
2866
2939
2952

**Figure S11**

**O-linked glycans analysis.** MALDI-TOF mass spectra of pooled primary nasal epithelial cells from human donors (A), primary tracheal epithelial cells from human donors (B, TTr05, C, TTr06, D, TTr08, E TTr09 and F, TTr10). Numbers above peaks correspond to the molecular ions identified based on manual annotation.

#### Figure S12

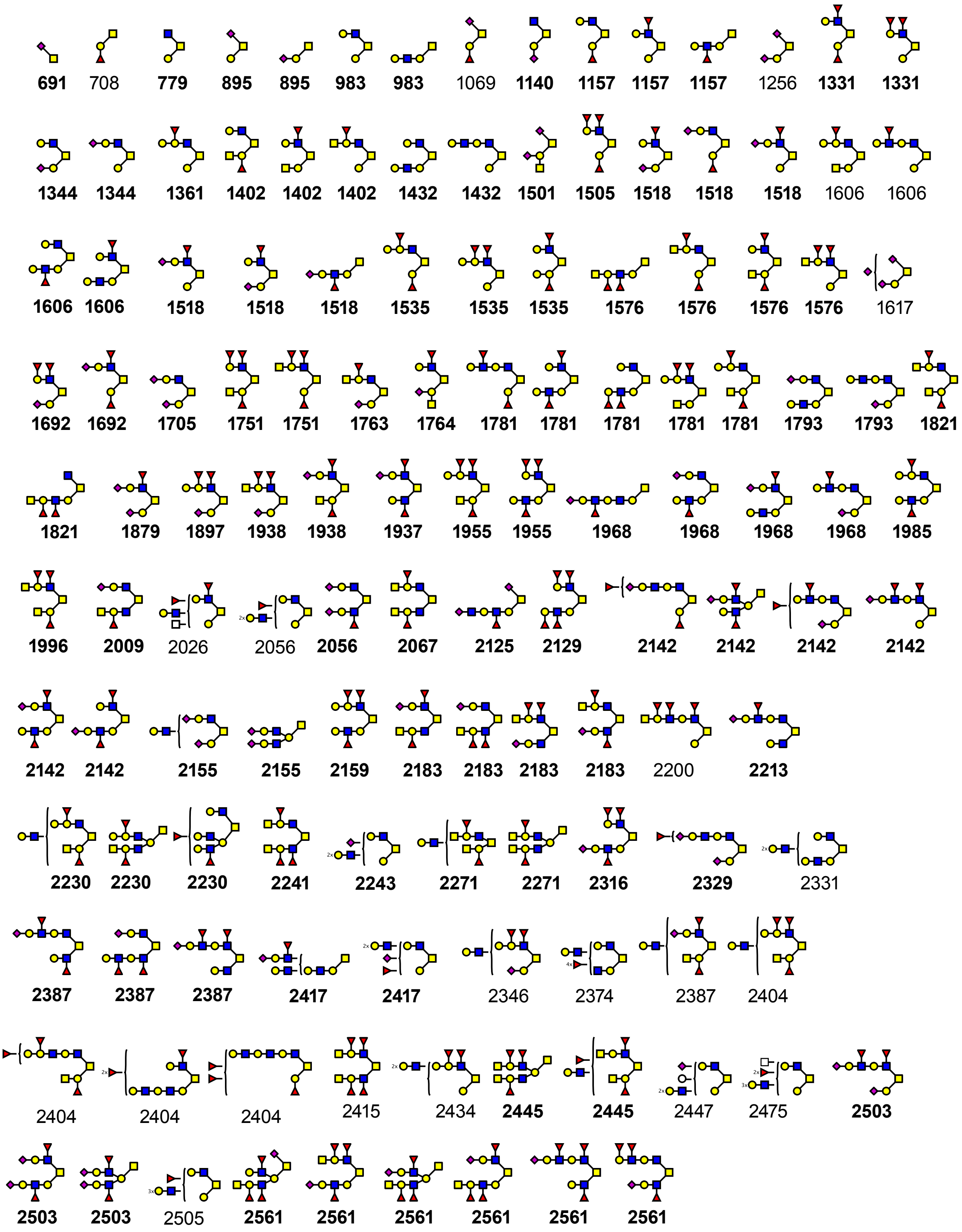
691 708 779 895 895 983 983 1069 1140 1157 1157 1157 1256 1331 1331
1344 1344 1361 1402 1402 1402 1432 1432 1501 1505 1518 1518 1518 1606 1606
1606 1606 1518 1518 1518 1535 1535 1535 1576 1576 1576 1576 1617
1692 1692 1705 1751 1751 1763 1764 1781 1781 1781 1781 1781 1793 1793 1821
1821 1879 1897 1938 1938 1937 1955 1955 1968 1968 1968 1968 1985
1996 2009 2026 2056 2056 2067 2125 2129 2142 2142 2142 2142
2142 2142 2155 2155 2159 2183 2183 2183 2183 2200 2213
2230 2230 2230 2241 2243 2271 2271 2316 2329 2331
2387 2387 2387 2417 2417 2346 2374 2387 2404
2404 2404 2404 2415 2434 2445 2445 2447 2475 2503
2503 2503 2505 2561 2561 2561 2561 2561 2561

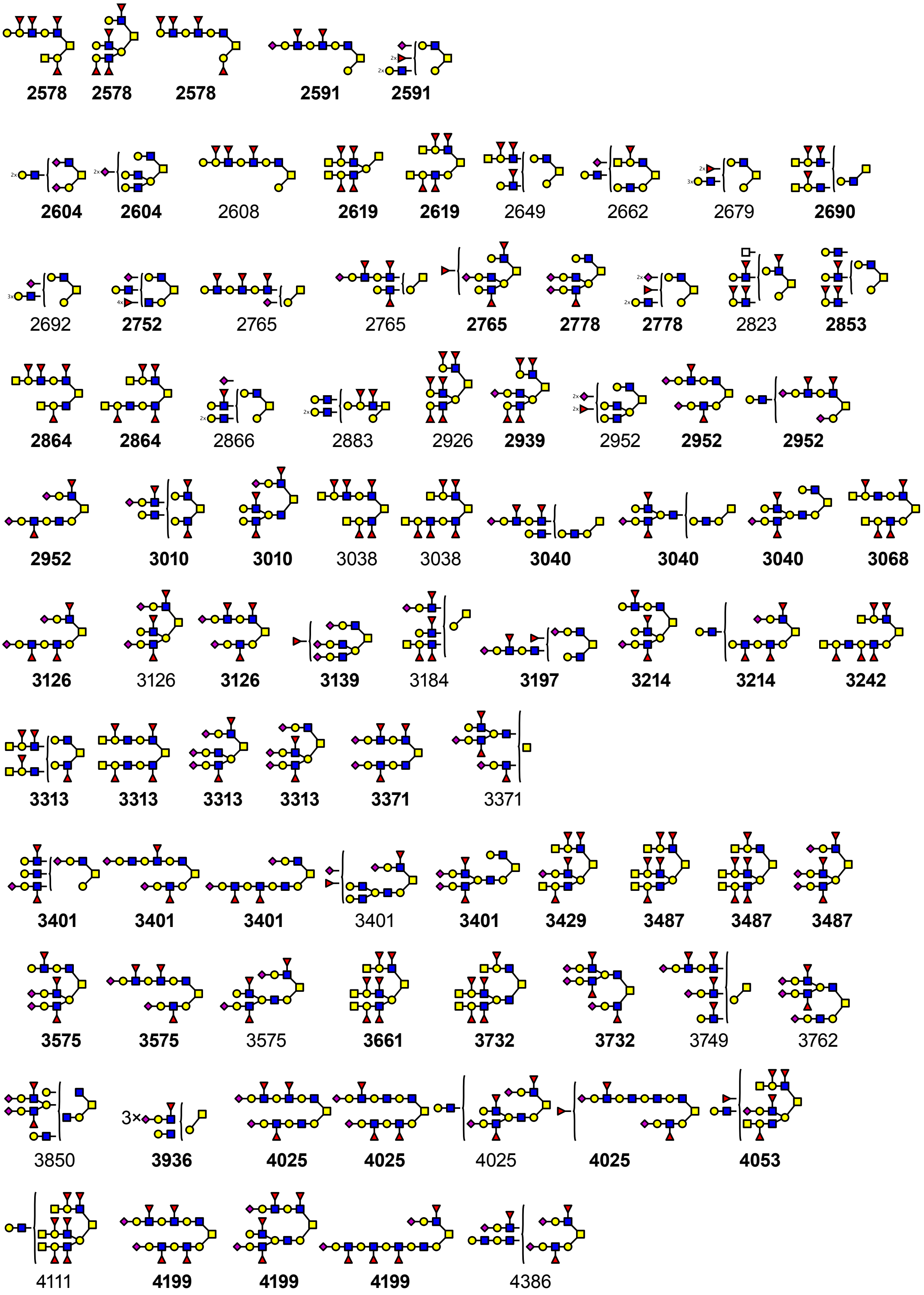
2578 2578 2578 2591 2591
2604 2604 2608 2619 2619 2649 2662 2679 2690
2692 2752 2765 2765 2765 2778 2778 2823 2853
2864 2864 2866 2883 2926 2939 2952 2952 2952
2952 3010 3010 3038 3038 3040 3040 3040 3068
3126 3126 3126 3139 3184 3197 3214 3214 3242
3313 3313 3313 3313 3371 3371
3401 3401 3401 3401 3401 3429 3487 3487 3487
3575 3575 3575 3661 3732 3732 3749 3762
3850 3936 4025 4025 4025 4025 4053
4111 4199 4199 4199 4386

**Figure S12**

**O-linked glycans structures library derived from primary tracheal and nasal epithelial cells from human donors.** Putative structures are based on composition, biosynthetic knowledge, and tandem mass spectrometry where available. Cartoon structures were drawn according to the Symbol Nomenclature for Glycans (SNFG) (Neelamegham et al., 2019 Glycobiology 29, 620-624) guidelines. Value under each putative structure corresponds to the molecular ion [M+Na]+ m/z value detected in the corresponding **Figure S11**. Values in bold correspond to a detected structure subjected to MALDI-TOF/TOF MS/MS analysis.

**Figure S13**

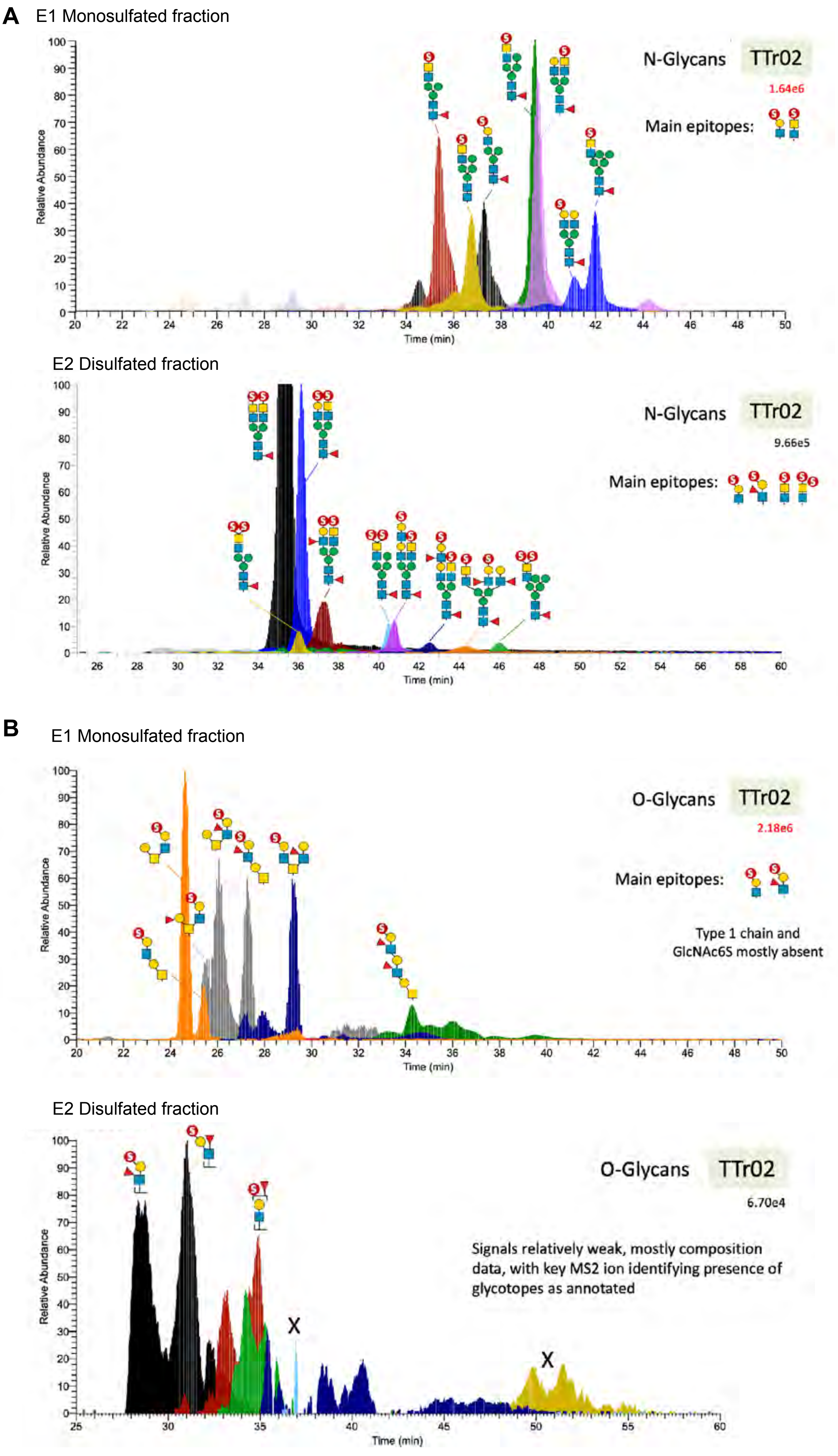
A
E1 Monosulfated fraction
N-Glycans
TTr02
1.64e6
Main epitopes:
Relative Abundance
Time (min)
E2 Disulfated fraction
N-Glycans
TTr02
9.66e5
Main epitopes:
Relative Abundance
Time (min)
B
E1 Monosulfated fraction
O-Glycans
TTr02
2.18e6
Main epitopes:
Type 1 chain and
GlcNAc6S mostly absent
Relative Abundance
Time (min)
E2 Disulfated fraction
O-Glycans
TTr02
6.70e4
Signals relatively weak, mostly composition
data, with key MS2 ion identifying presence of
glycotopes as annotated
X
X
Relative Abundance
Time (min)

**Figure S13**

**Sulfated glycan analyses.** Partial LC-MS spectra of (A) N-linked sulfated glycans and (B) O-linked glycans derived from tracheal epithelial cells from the human donor (TTr2). The panels show the dominant species detected in the analyses. The putative structures shown in the figure were manually annotated.

### Figure S14

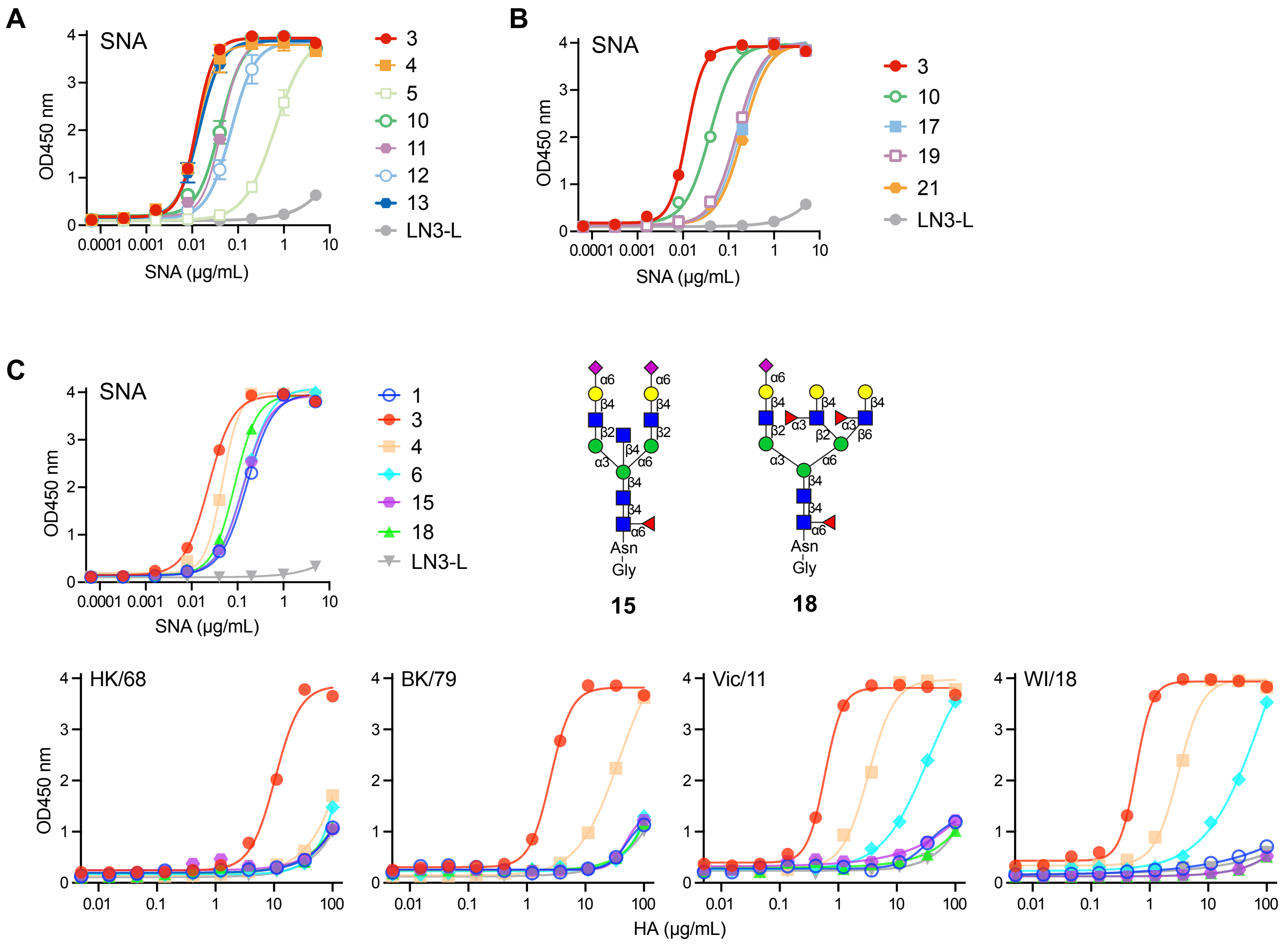
A
SNA
OD450 nm
SNA (μg/mL)
3
4
5
10
11
12
13
LN3-L
B
SNA
OD450 nm
SNA (μg/mL)
3
10
17
19
21
LN3-L
C
SNA
OD450 nm
SNA (μg/mL)
1
3
4
6
15
18
LN3-L
Asn
Gly
15
18
HK/68
BK/79
Vic/11
WI/18
OD450 nm
HA (μg/mL)

**Figure S14**

**Glycan ELISA.** Glycan structures corresponding to the shown compound numbers are shown in **Figure 3**. (**A,B**) The loading of glycans was assessed using the SNA lectin for the glycan ELISA analyses shown in **Figure 3**. (**C**) Binding recombinant H3 HAs to bisected (compound 15) and short tri-antennary (compound 18) glycans. The loading of glycans was assessed by SNA lectin binding.

**Figure S15**

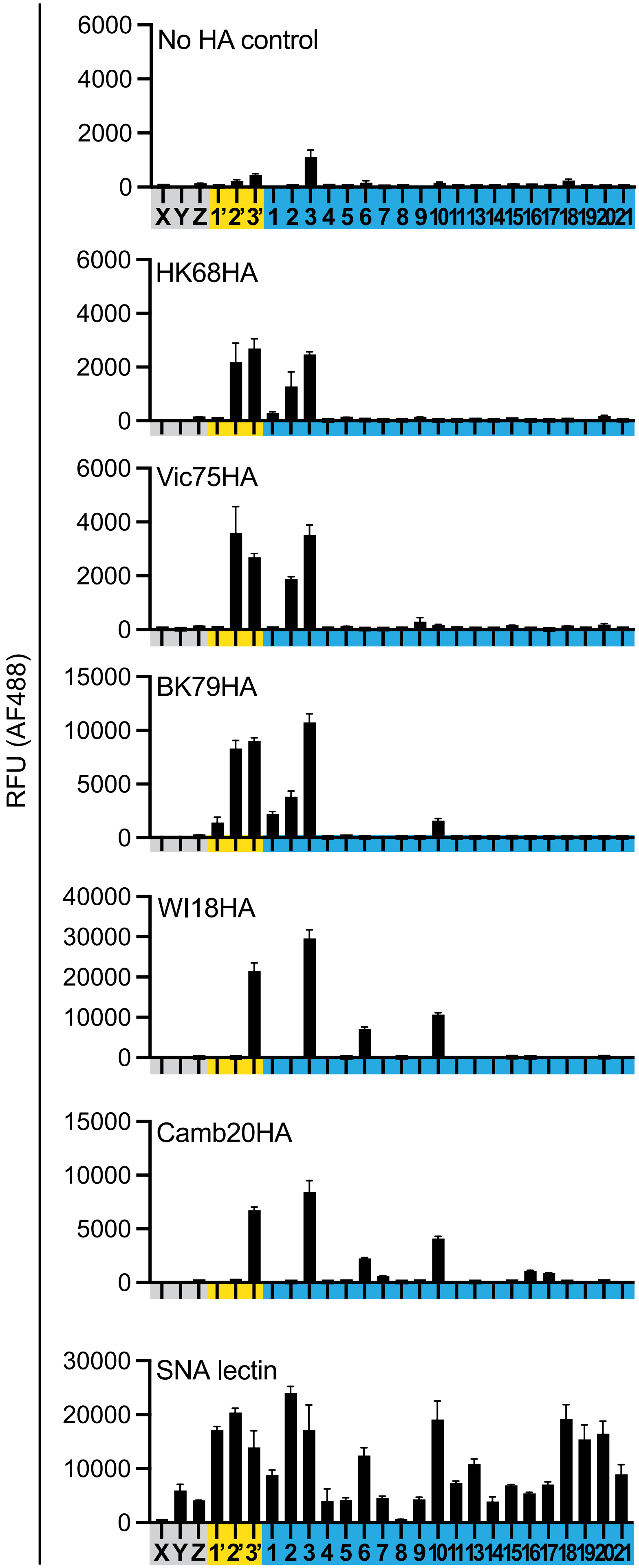


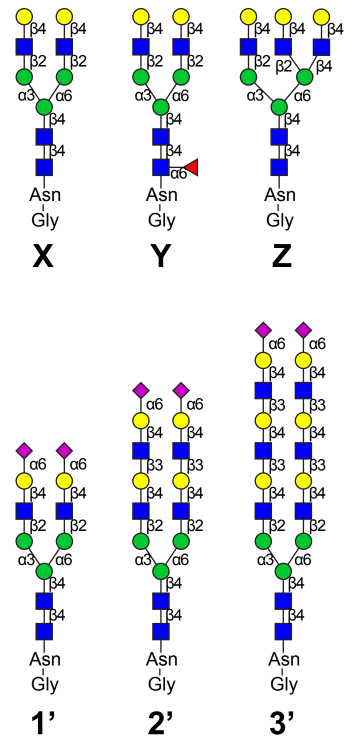


**1-21:** Shown in Fig 3

**Figure S15**

**Glycan array.** A panel of recombinant H3 HAs were tested for the binding to a custom glycan microarray with control glycans (compounds X-Z and 1'-3'; structures shown in the figure) and human airway glycan library (**Figure 3**, compounds 1-21). On the left, graphs show the binding of different samples, from the top, no HA control, HK68, Vic75, BK79, WI18, and Camb20 HAs. The printing efficiency was confirmed by the binding of SNA lectin (bottom graph). Bars depict mean intensity minus mean background of the four medians out of six total replicate observations with error bars showing standard error.

**Figure S16**

**A**

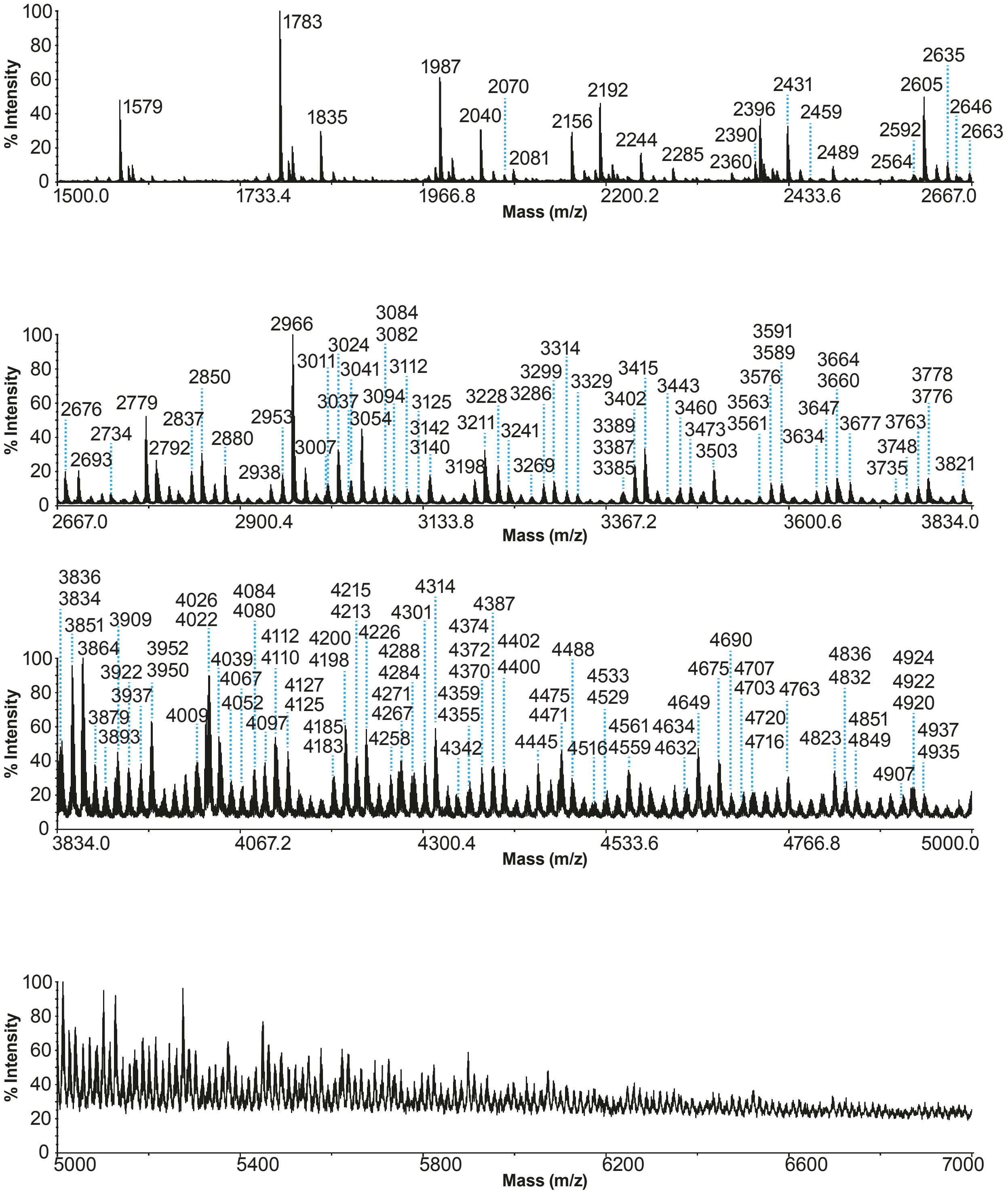

% Intensity
Mass (m/z)
1500.0
1733.4
1966.8
2200.2
2433.6
2667.0
1579
1783
1835
1987
2040
2070
2081
2156
2192
2244
2285
2360
2390
2396
2431
2459
2489
2564
2592
2605
2635
2646
2663
2676
2693
2734
2779
2792
2837
2850
2880
2938
2953
2966
3007
3011
3024
3037
3041
3054
3082
3084
3094
3112
3125
3140
3142
3198
3211
3228
3241
3269
3286
3299
3314
3329
3385
3387
3389
3402
3415
3443
3460
3473
3503
3561
3563
3576
3589
3591
3600.6
3634
3647
3660
3664
3677
3735
3748
3763
3776
3778
3821
3834.0
3834
3836
3851
3864
3879
3893
3909
3922
3937
3950
3952
4009
4022
4026
4039
4052
4067
4067.2
4080
4084
4097
4110
4112
4125
4127
4183
4185
4198
4200
4213
4215
4226
4258
4267
4271
4284
4288
4300.4
4301
4314
4342
4355
4359
4370
4372
4374
4387
4400
4402
4445
4471
4475
4488
4516
4529
4533
4533.6
4559
4561
4632
4634
4649
4675
4690
4703
4707
4716
4720
4763
4766.8
4823
4832
4836
4849
4851
4907
4920
4922
4924
4935
4937
5000.0
5000
5400
5800
6200
6600
7000
0
20
40
60
80
100

**B**

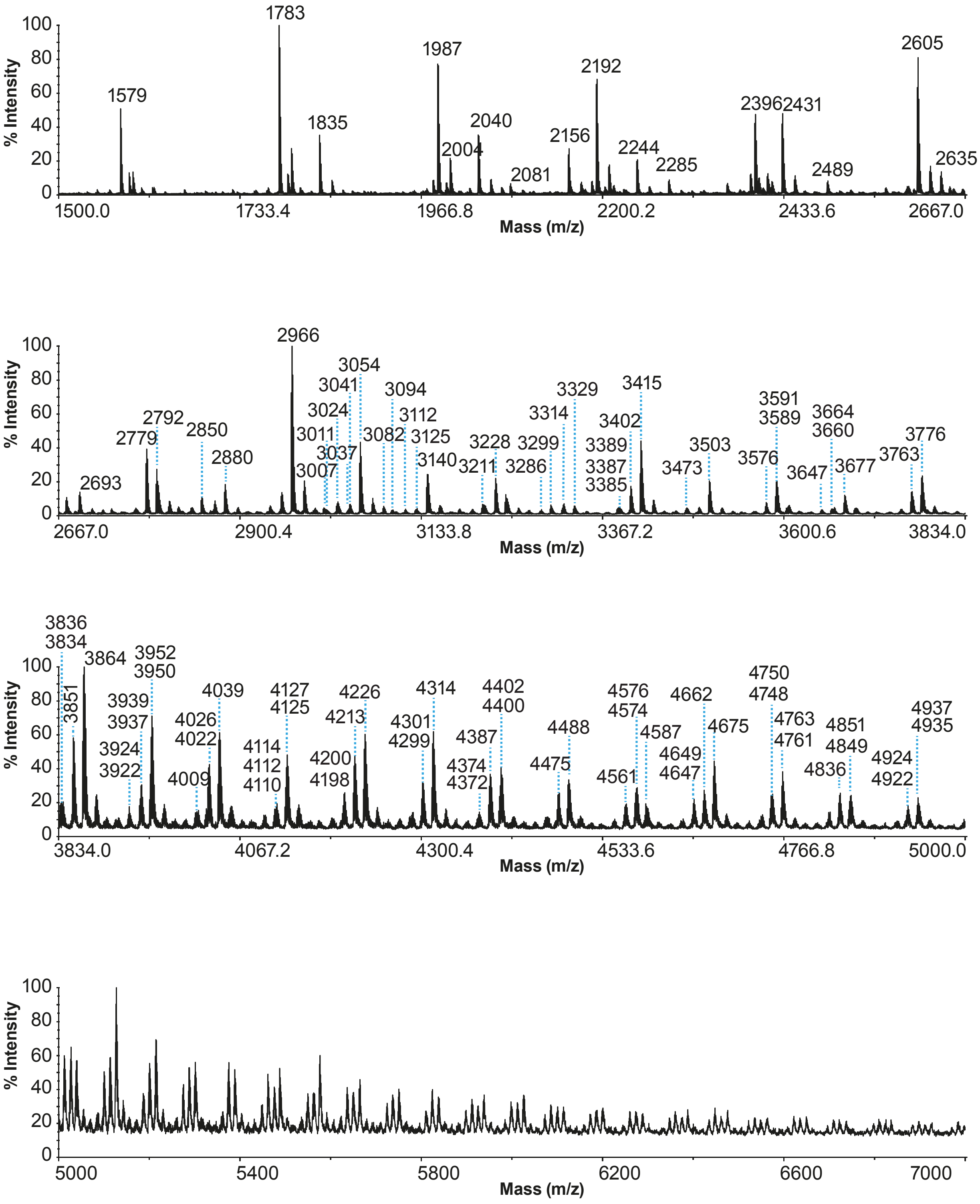
% Intensity
Mass (m/z)
100
80
60
40
20
0
1500.0
1733.4
1966.8
2200.2
2433.6
2667.0
1579
1783
1835
1987
2004
2040
2081
2156
2192
2244
2285
2396
2431
2489
2605
2635
2900.4
3133.8
3367.2
3600.6
3834.0
2693
2779
2792
2850
2880
2966
3007
3011
3024
3037
3041
3054
3082
3094
3112
3125
3140
3211
3228
3286
3299
3314
3329
3385
3387
3389
3402
3415
3473
3503
3576
3589
3591
3647
3660
3664
3677
3763
3776
4067.2
4300.4
4533.6
4766.8
5000.0
3834
3836
3851
3864
3922
3924
3937
3939
3950
3952
4009
4022
4026
4039
4110
4112
4114
4125
4127
4198
4200
4213
4226
4299
4301
4314
4372
4374
4387
4400
4402
4475
4488
4561
4574
4576
4587
4647
4649
4662
4675
4748
4750
4761
4763
4836
4849
4851
4922
4924
4935
4937
5000
5400
5800
6200
6600
7000

**Figure S16**

**MALDI-TOF mass spectra of permethylated N-linked glycans of primary tracheal epithelial cells from human donors cultured in air-liquid interface.**

MALDI-TOF MS analysis of permethylated N-linked glycans from (A) TTr02-ALI and (B) TTr03-ALI epithelial cells of human trachea samples cultured in air-liquid interface (ALI). All molecular ions are $[M+Na]^+$. Putative structures are based on composition, tandem mass spectrometry, and biosynthetic knowledge. Percentages on top left of each panel correspond to the relative percentage of the maximum peak of the corresponding panel compared to the relative intensity of the maximum peak of the top panel (based on the total ion current of each panel, without de-isotoping). Full structure annotations can be found in **Figure S5**.

**Figure S17**

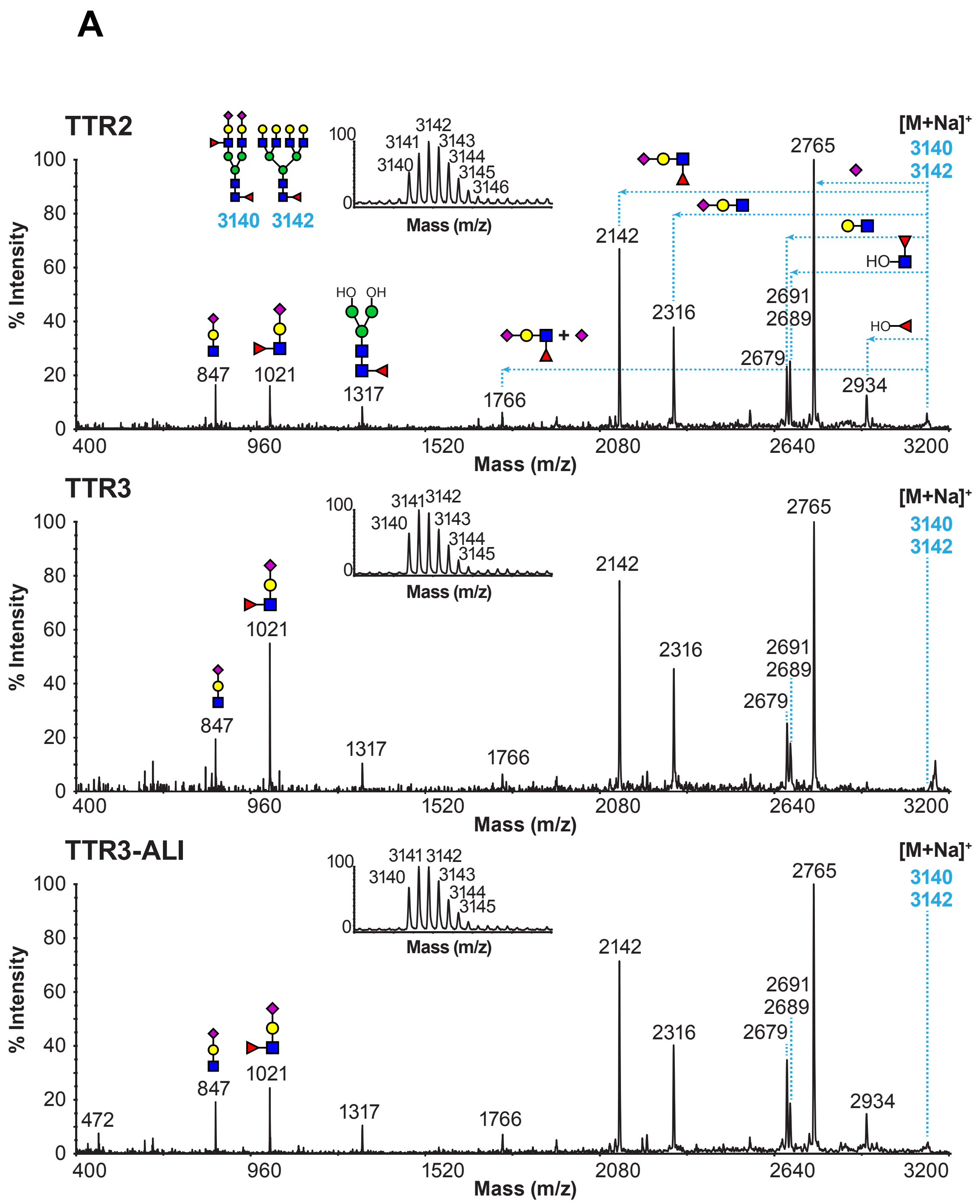
A
TTR2
% Intensity
Mass (m/z)
3140
3142
3141
3143
3144
3145
3146
[M+Na]+
2765
2142
2316
2691
2689
2679
2934
1766
1317
1021
847
HO
OH
TTR3
TTR3-ALI
472

**B**

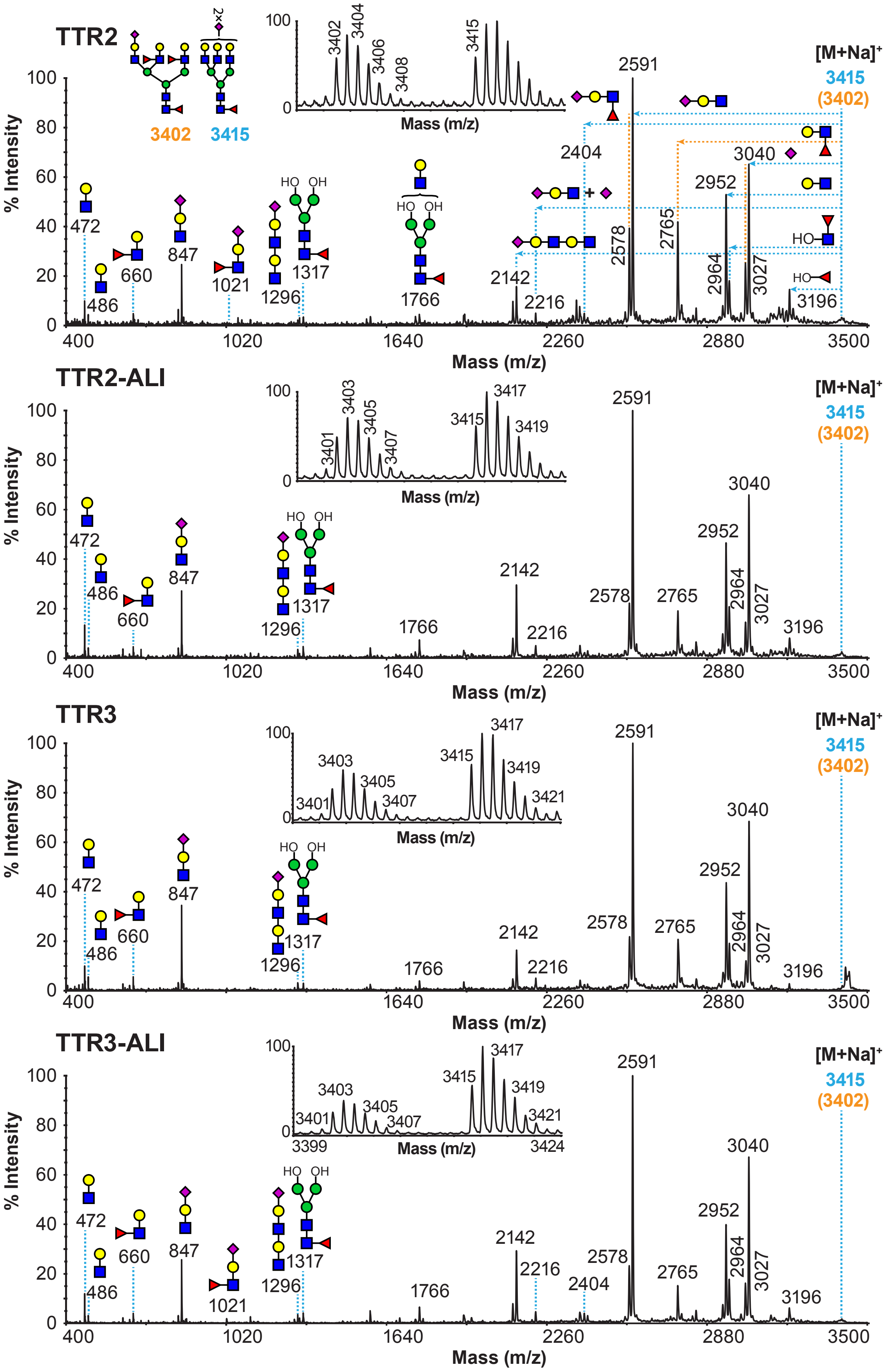

TTR2
3402
3415
[M+Na]+
3415
(3402)
% Intensity
Mass (m/z)
472
486
660
847
1021
1296
1317
1766
2142
2216
2404
2578
2591
2765
2952
2964
3027
3040
3196
3402
3404
3406
3408
3415
TTR2-ALI
3401
3403
3405
3407
3417
3419
TTR3
3421
TTR3-ALI
3399
3424

**C**

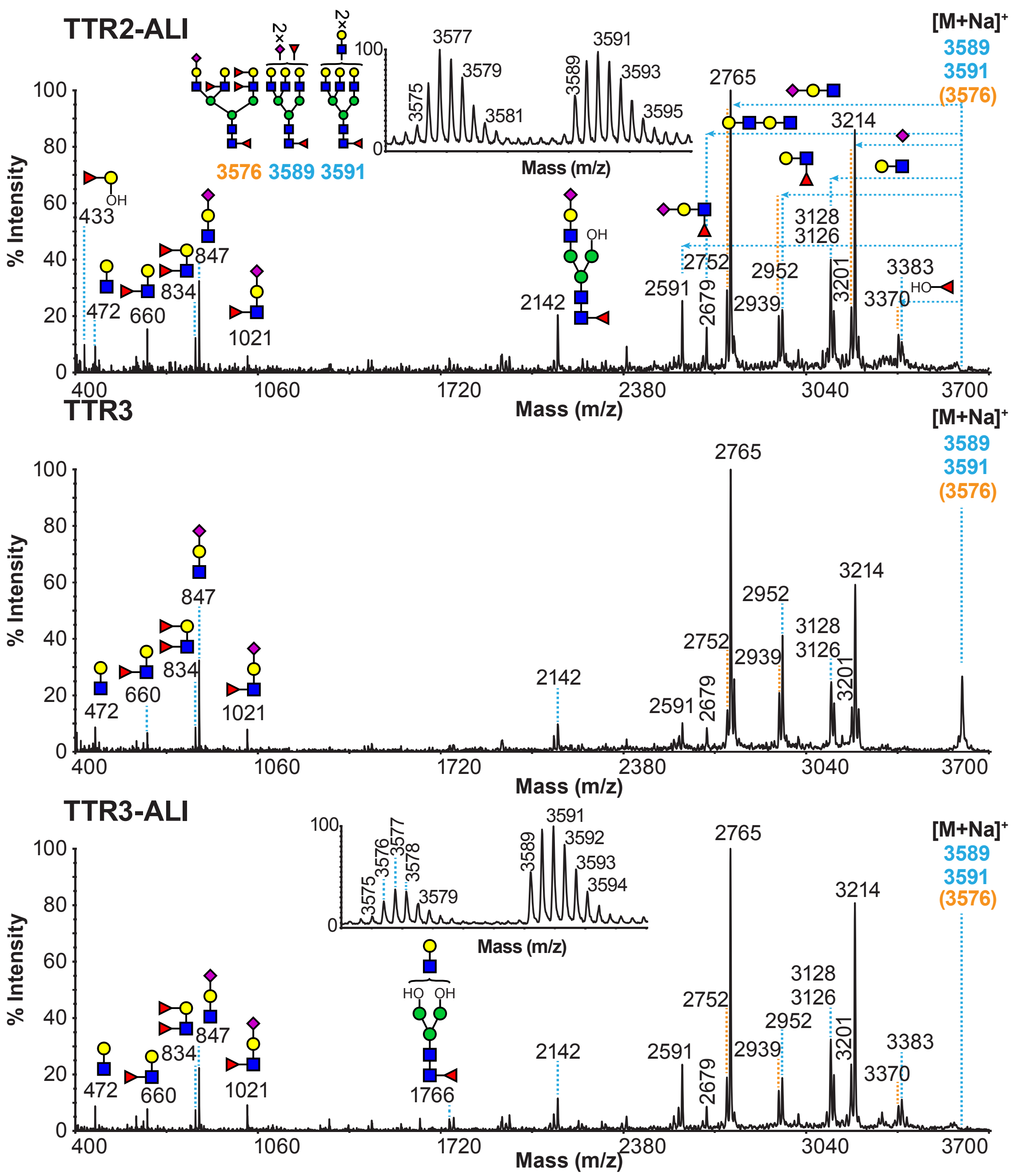
TTR2-ALI
TTR3
TTR3-ALI
[M+Na]+
3589
3591
(3576)
3576 3589 3591
% Intensity
Mass (m/z)
2765
3214
3128
3126
2952
2939
2752
2679
2591
2142
3201
3383
3370
1766
1021
847
834
660
472
433
3577
3579
3581
3575
3591
3589
3593
3595
3592
3594
3576
3578
400
1060
1720
2380
3040
3700

## D

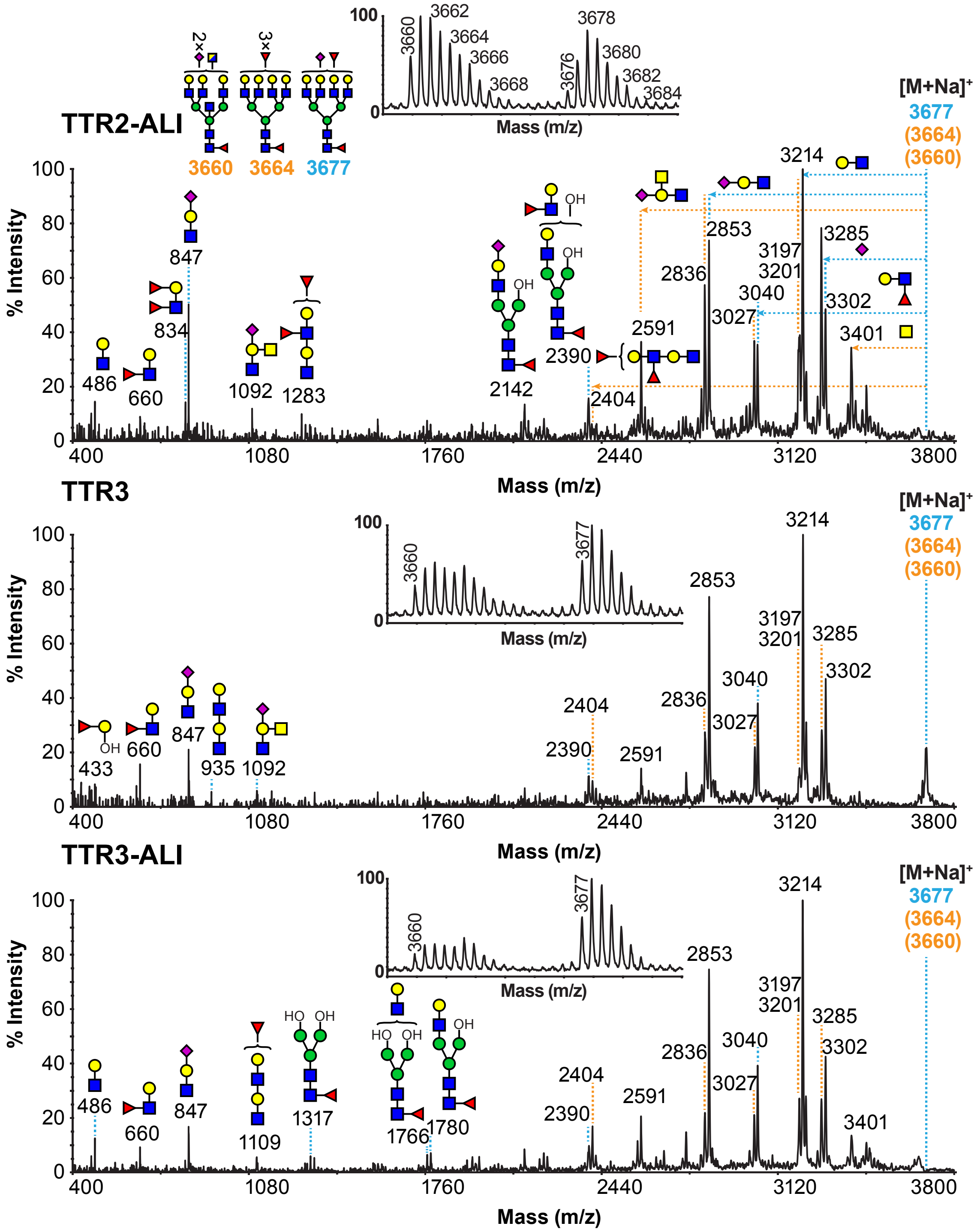
TTR2-ALI
3660 3664 3677
[M+Na]+
3677
(3664)
(3660)
% Intensity
Mass (m/z)
486 660 834 847 1092 1283 2142 2390 2404 2591 2836 2853 3027 3040 3197 3201 3214 3285 3302 3401
3660 3662 3664 3666 3668 3676 3678 3680 3682 3684
TTR3
433 660 847 935 1092 2390 2404 2591 2836 2853 3027 3040 3197 3201 3214 3285 3302
TTR3-ALI
486 660 847 1109 1317 1766 1780 2390 2404 2591 2836 2853 3027 3040 3197 3201 3214 3285 3302 3401

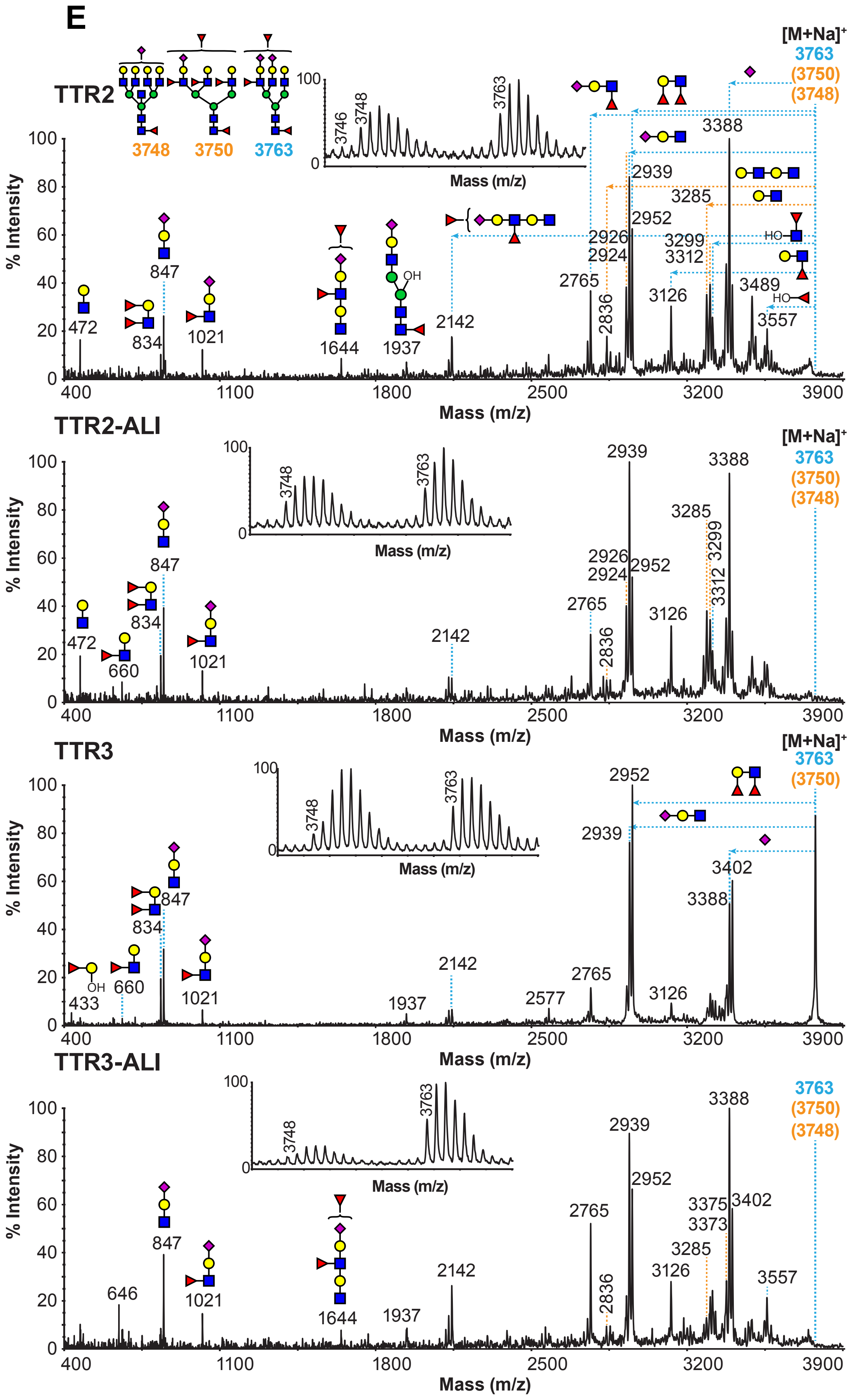

E
TTR2
3748
3750
3763
[M+Na]+
3763
(3750)
(3748)
TTR2-ALI
TTR3
TTR3-ALI
% Intensity
Mass (m/z)

F

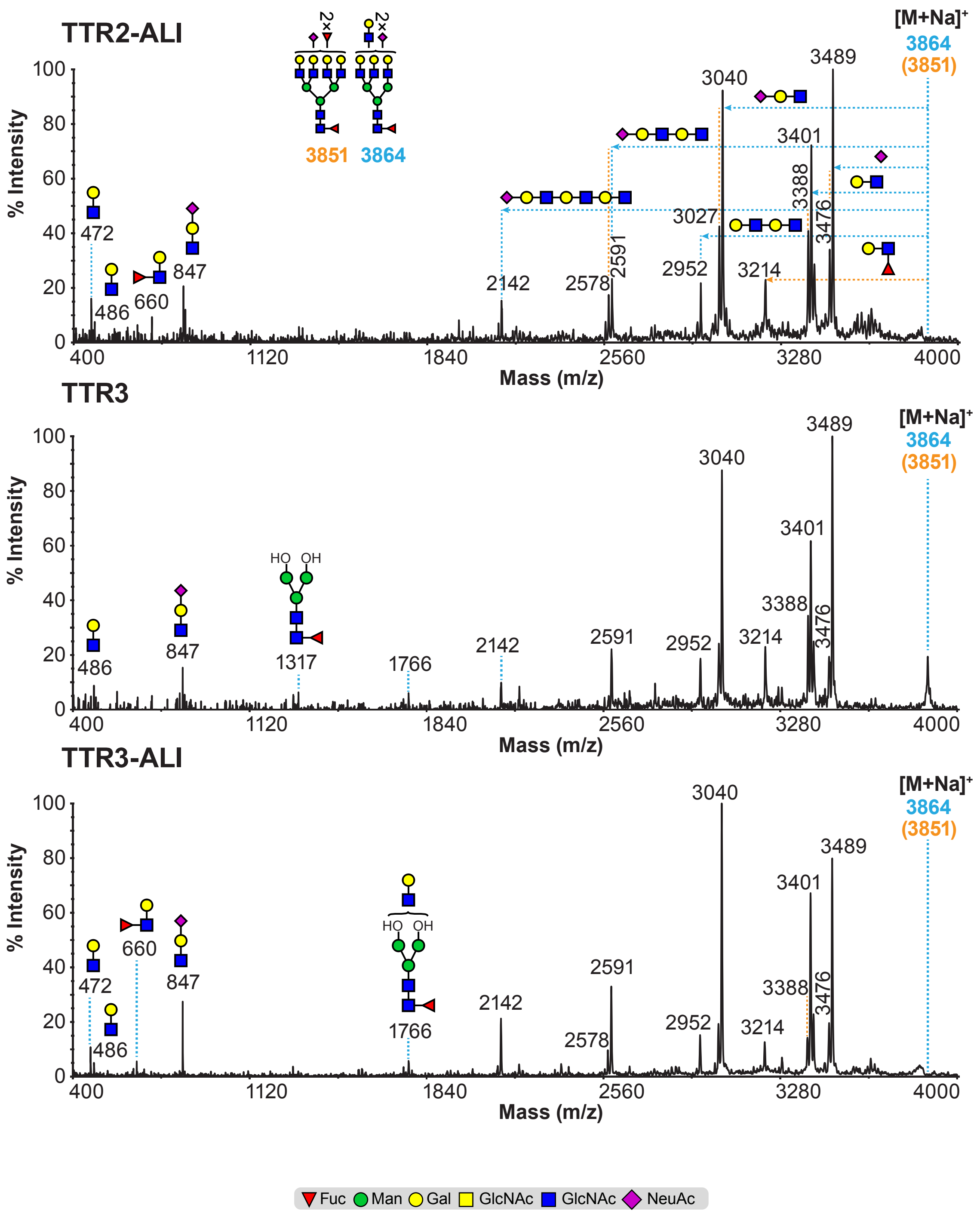
TTR2-ALI
TTR3
TTR3-ALI
% Intensity
Mass (m/z)
[M+Na]+
3864
(3851)
3851 3864
2×
472
486
660
847
1317
1766
2142
2578
2591
2952
3027
3040
3214
3388
3401
3476
3489
HO OH
Fuc Man Gal GlcNAc GlcNAc NeuAc

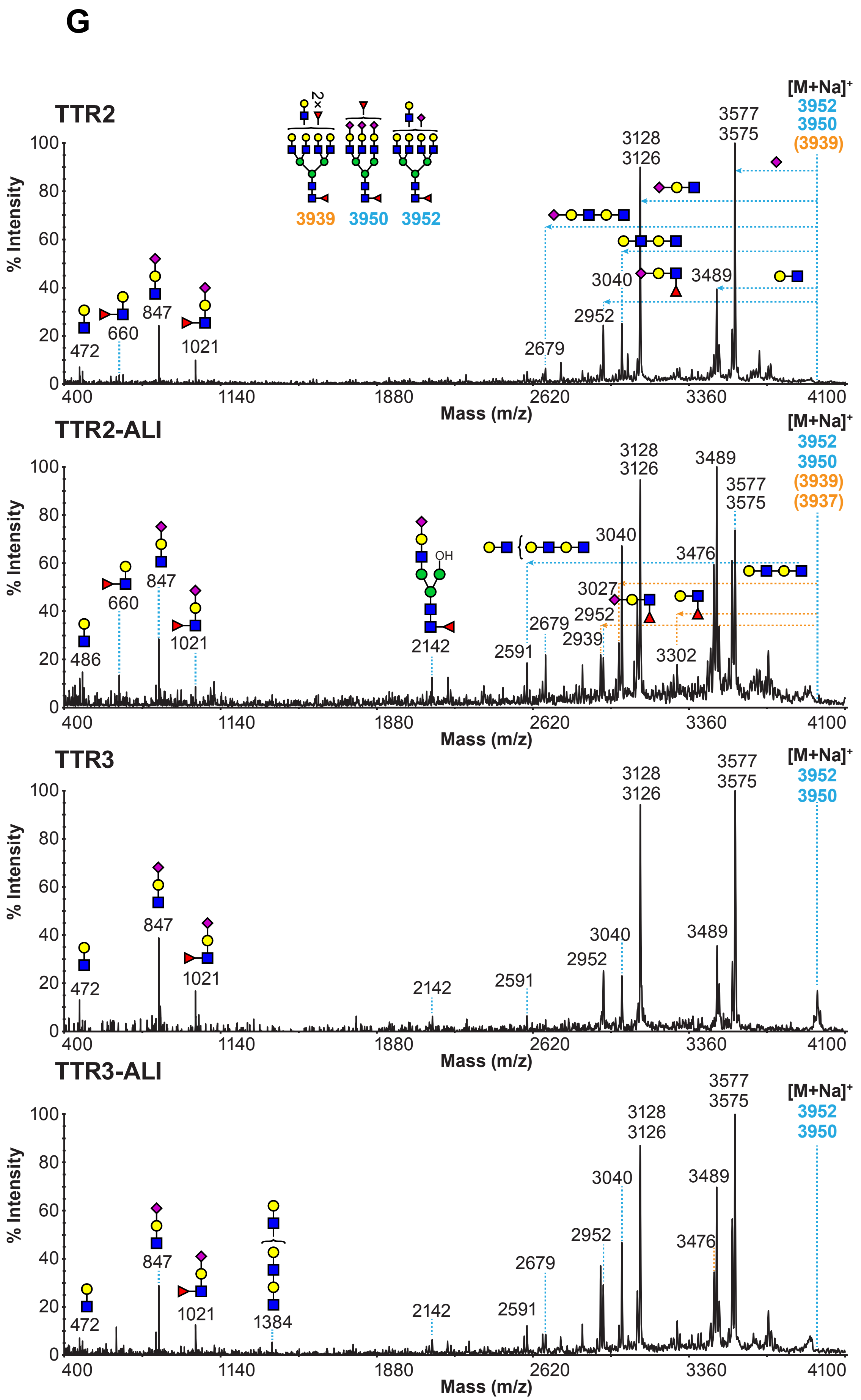
G
TTR2
[M+Na]+
3952
3950
(3939)
3939 3950 3952
2×
3577
3575
3128
3126
3040
3489
2952
2679
472
660
847
1021
% Intensity
Mass (m/z)
TTR2-ALI
[M+Na]+
3952
3950
(3939)
(3937)
3128
3126
3489
3577
3575
3040
3476
3027
2952
2939
2679
2591
3302
2142
OH
486
660
847
1021
TTR3
[M+Na]+
3952
3950
3128
3126
3577
3575
3040
3489
2952
2591
2142
472
847
1021
TTR3-ALI
[M+Na]+
3952
3950
3577
3575
3128
3126
3040
3489
2952
3476
2679
2591
2142
1384
1021
847
472
400 1140 1880 2620 3360 4100
0 20 40 60 80 100

**H**

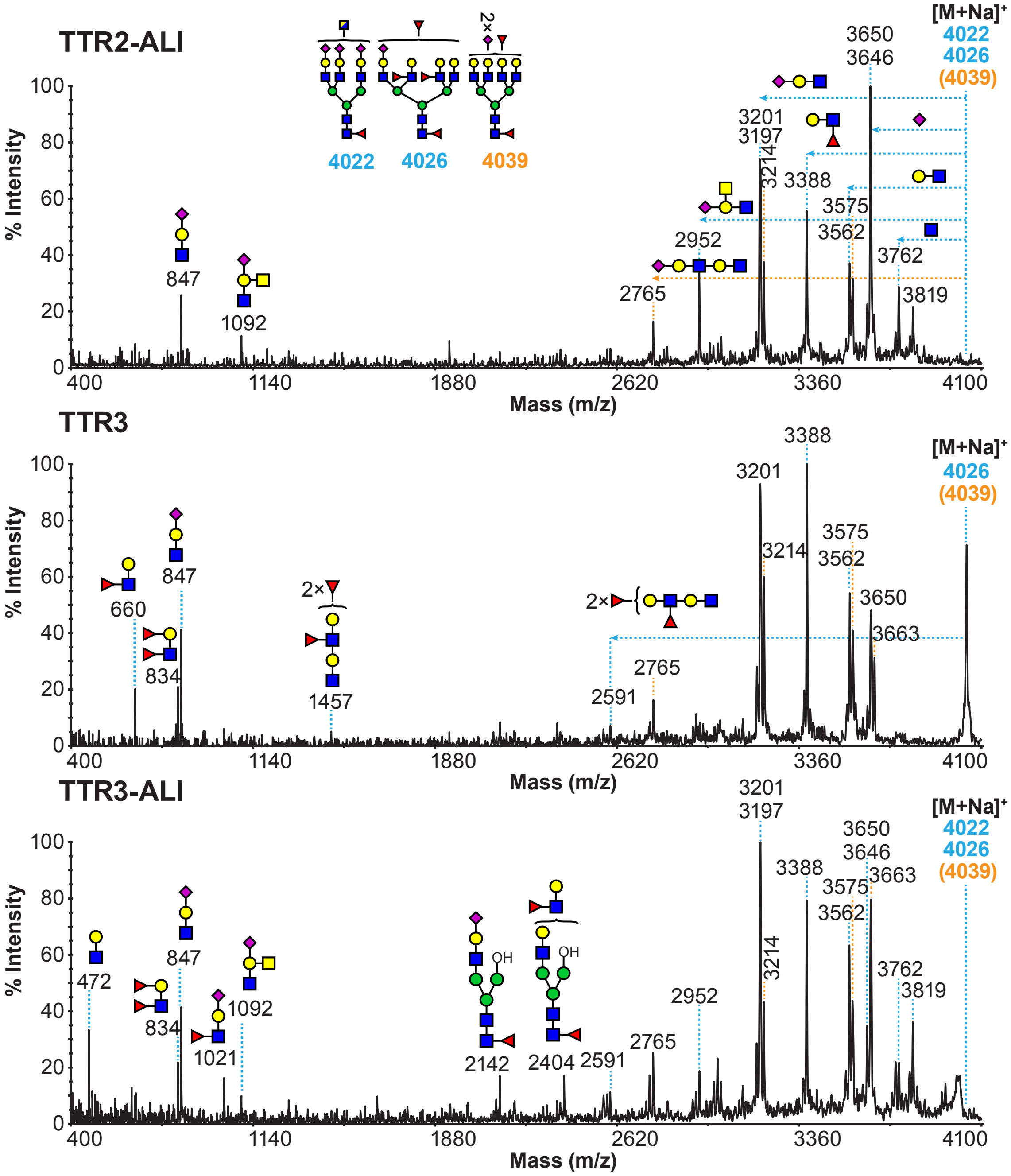

TTR2-ALI
[M+Na]+
4022
4026
(4039)
4022
4026
4039
2×
% Intensity
Mass (m/z)
847
1092
2765
2952
3201
3197
3214
3388
3575
3562
3650
3646
3762
3819
TTR3
[M+Na]+
4026
(4039)
660
834
847
2×
1457
2×
2591
2765
3201
3214
3388
3575
3562
3650
3663
TTR3-ALI
[M+Na]+
4022
4026
(4039)
472
834
847
1021
1092
OH
2142
2404
2591
2765
2952
3201
3197
3214
3388
3575
3562
3650
3646
3663
3762
3819
400
1140
1880
2620
3360
4100

**I**

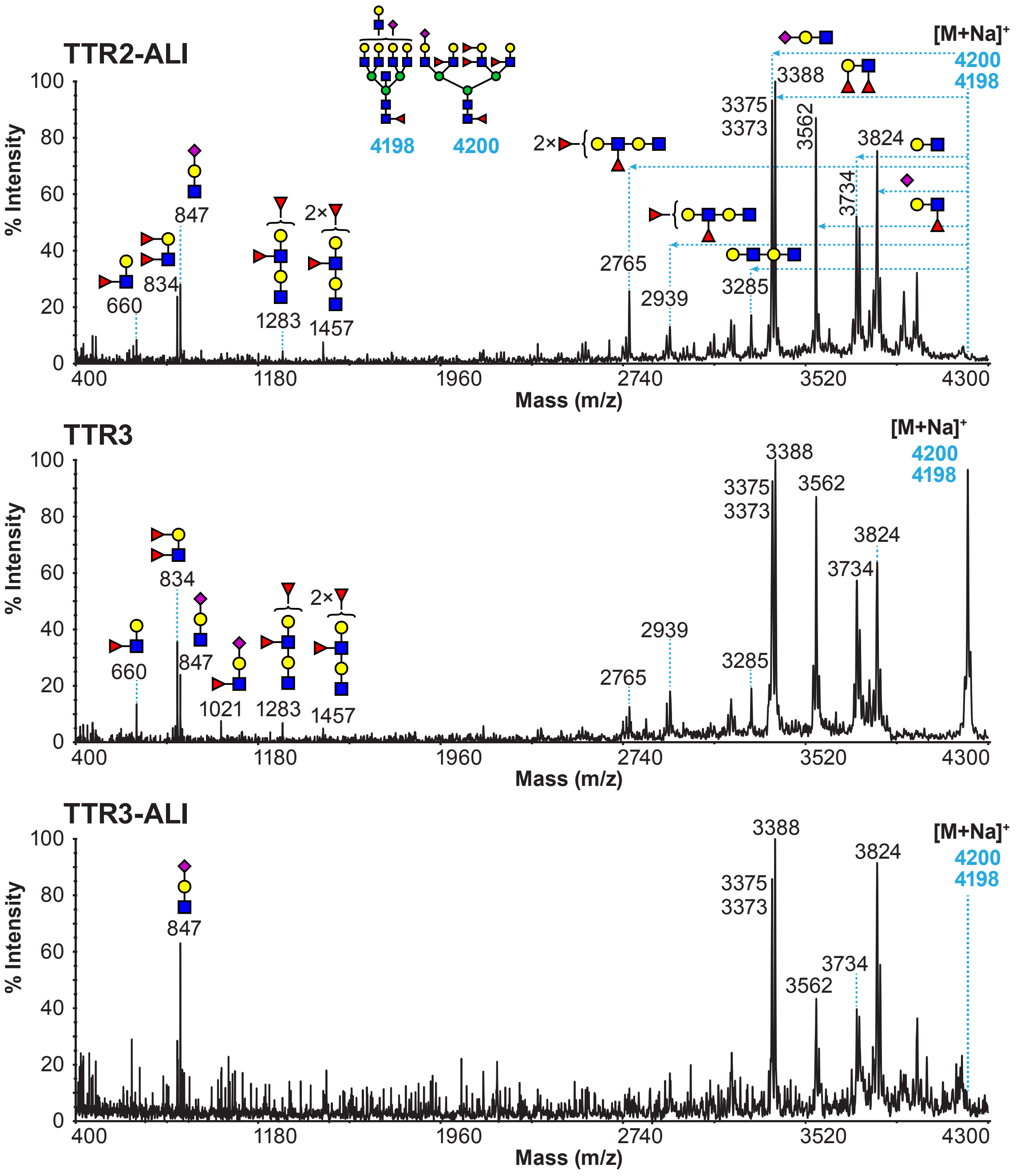
TTR2-ALI
% Intensity
Mass (m/z)
[M+Na]+
4200
4198
660
834
847
1283
1457
2765
2939
3285
3375
3373
3388
3562
3734
3824
2×
TTR3
1021
TTR3-ALI

J

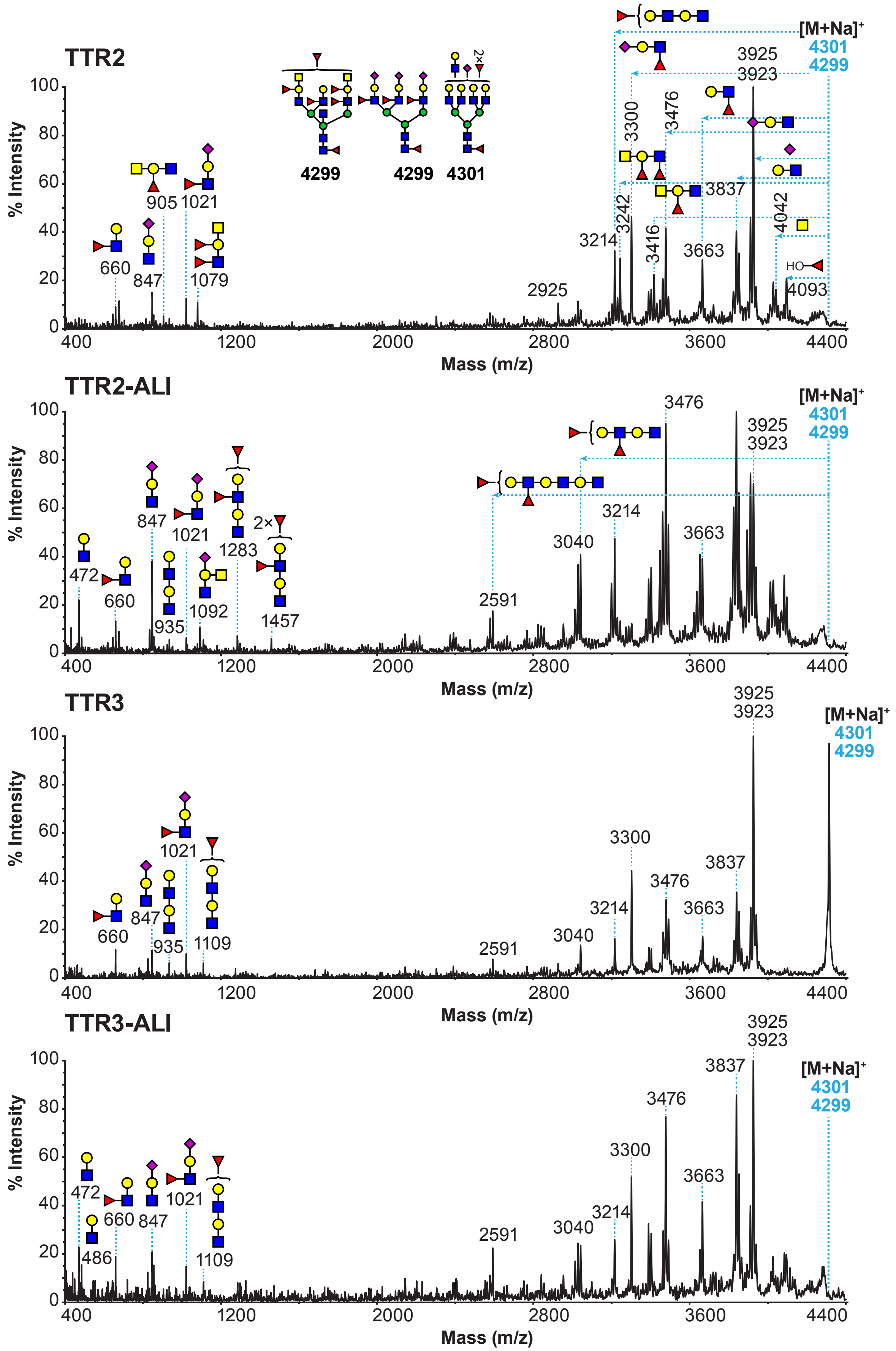
TTR2
TTR2-ALI
TTR3
TTR3-ALI
% Intensity
Mass (m/z)
[M+Na]+
4301
4299
3925
3923
3300
3476
3242
3214
3416
3663
3837
4042
4093
2925
660
847
905
1021
1079
472
935
1092
1283
1457
2591
3040
486
1109
HO
2×

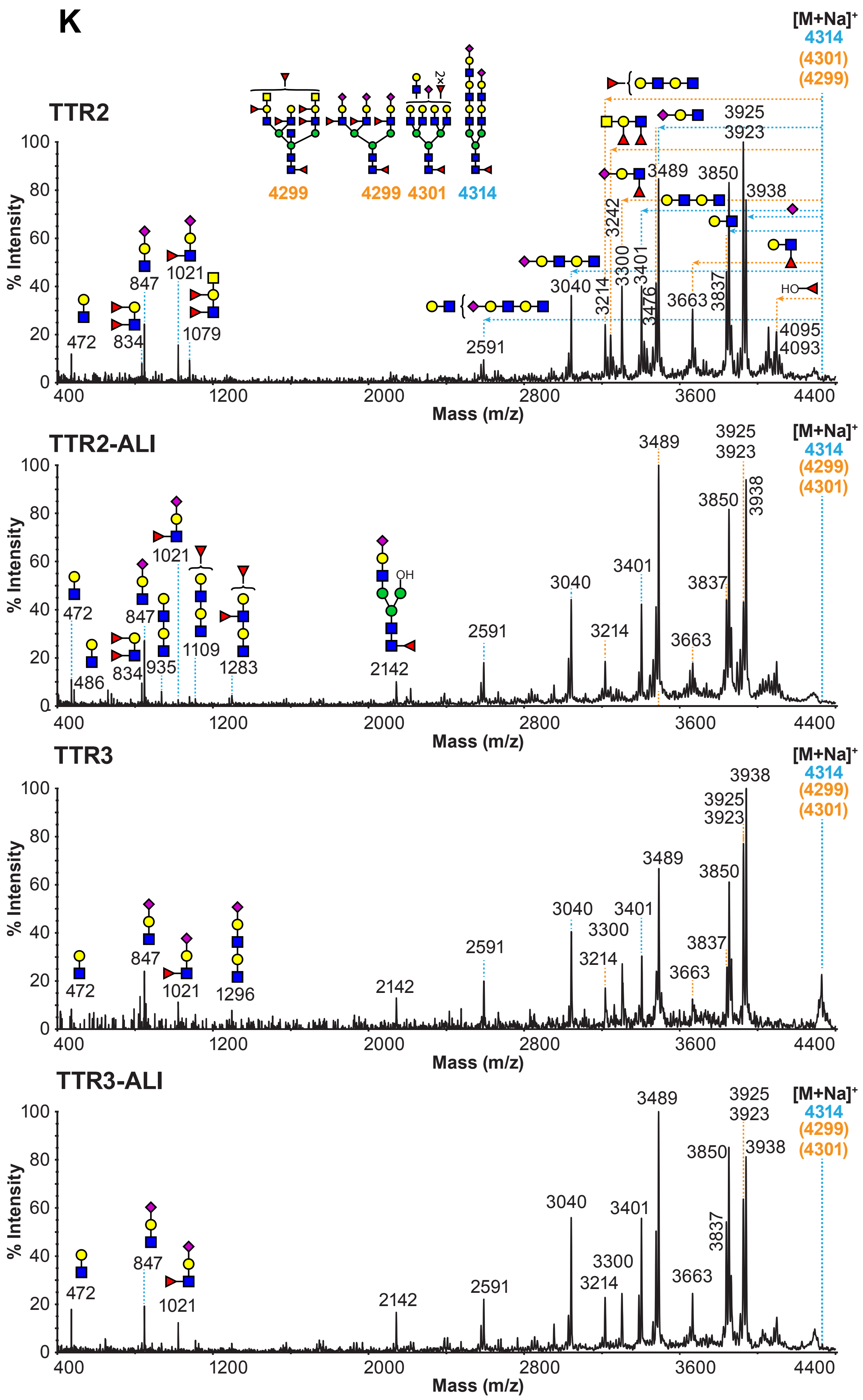
K
TTR2
TTR2-ALI
TTR3
TTR3-ALI
[M+Na]+
4314
(4301)
(4299)
4299
4299 4301 4314
% Intensity
Mass (m/z)
472
834
847
1021
1079
2591
3040
3214
3242
3300
3401
3476
3489
3663
3837
3850
3925
3923
3938
4095
4093
486
935
1109
1283
2142
1296
HO
OH

L

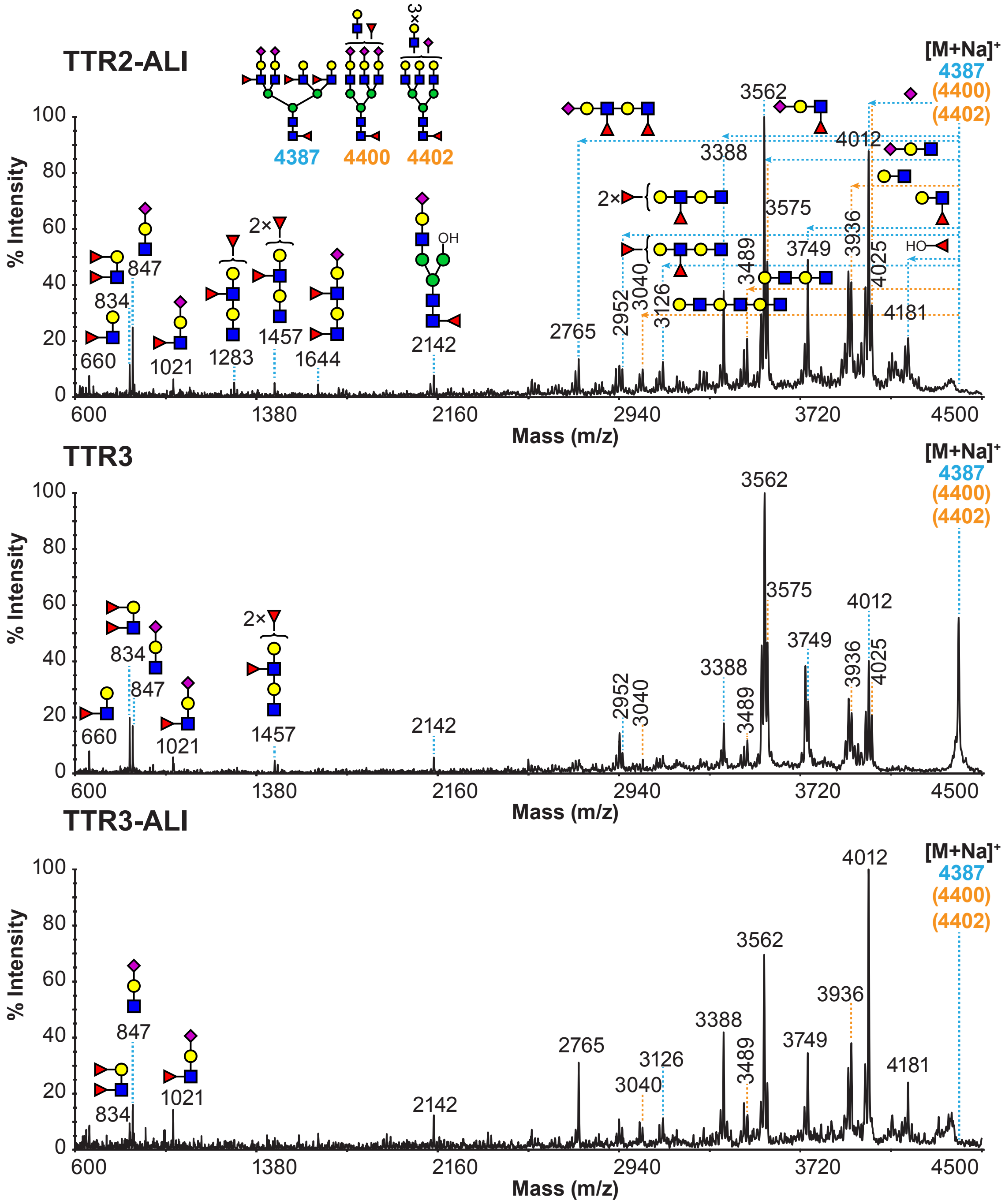

TTR2-ALI
TTR3
TTR3-ALI
[M+Na]+
4387
(4400)
(4402)
% Intensity
Mass (m/z)
660
834
847
1021
1283
1457
1644
2142
2765
2952
3040
3126
3388
3489
3562
3575
3749
3936
4012
4025
4181
4387
4400
4402

M

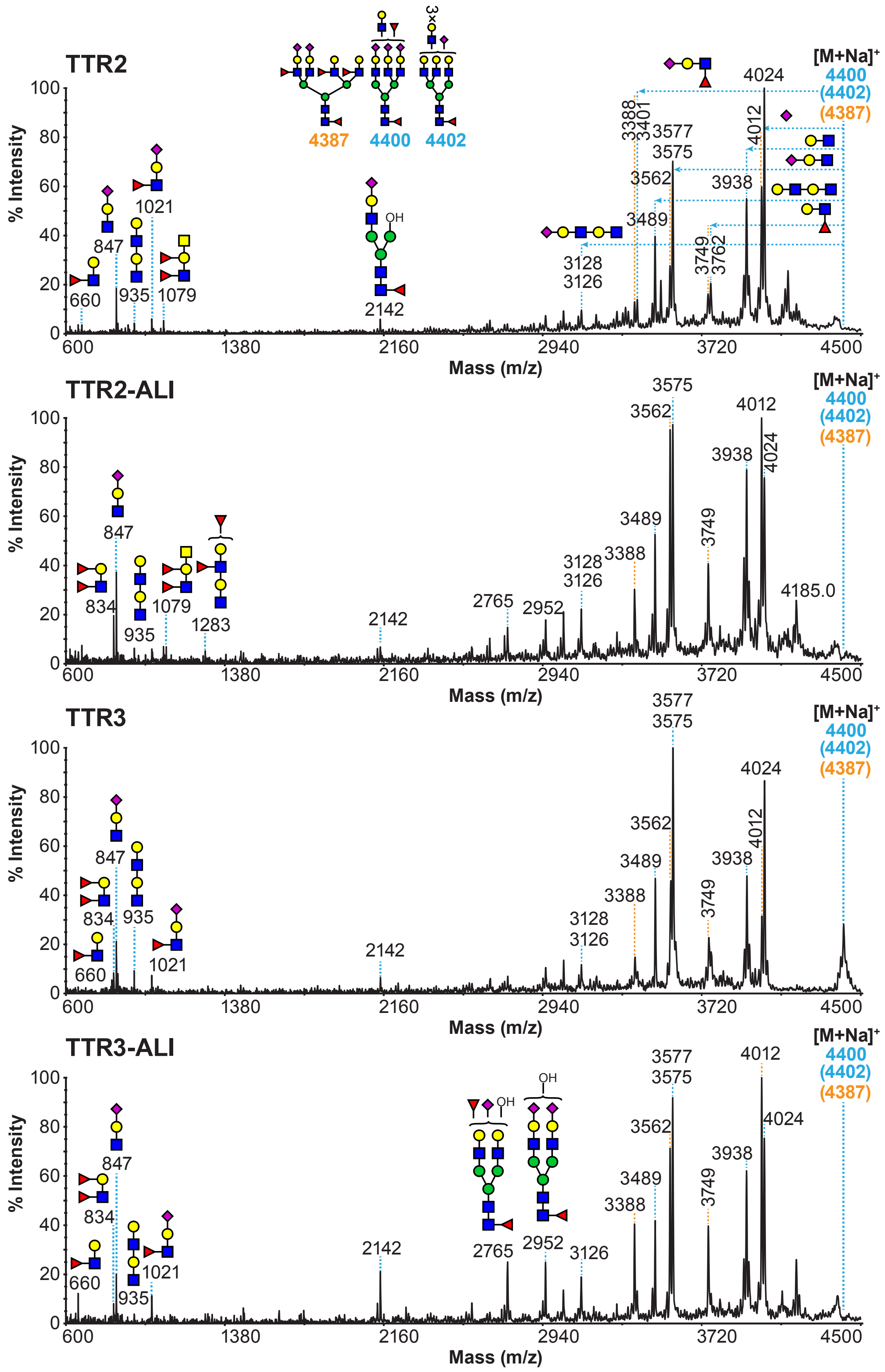

TTR2
TTR2-ALI
TTR3
TTR3-ALI
% Intensity
Mass (m/z)
[M+Na]+
4400
(4402)
(4387)
4387
4400
4402
3×
660
847
935
1021
1079
2142
3128
3126
3388
3401
3489
3562
3575
3577
3749
3762
3938
4012
4024
834
1283
2765
2952
4185.0
OH
600
1380
2160
2940
3720
4500

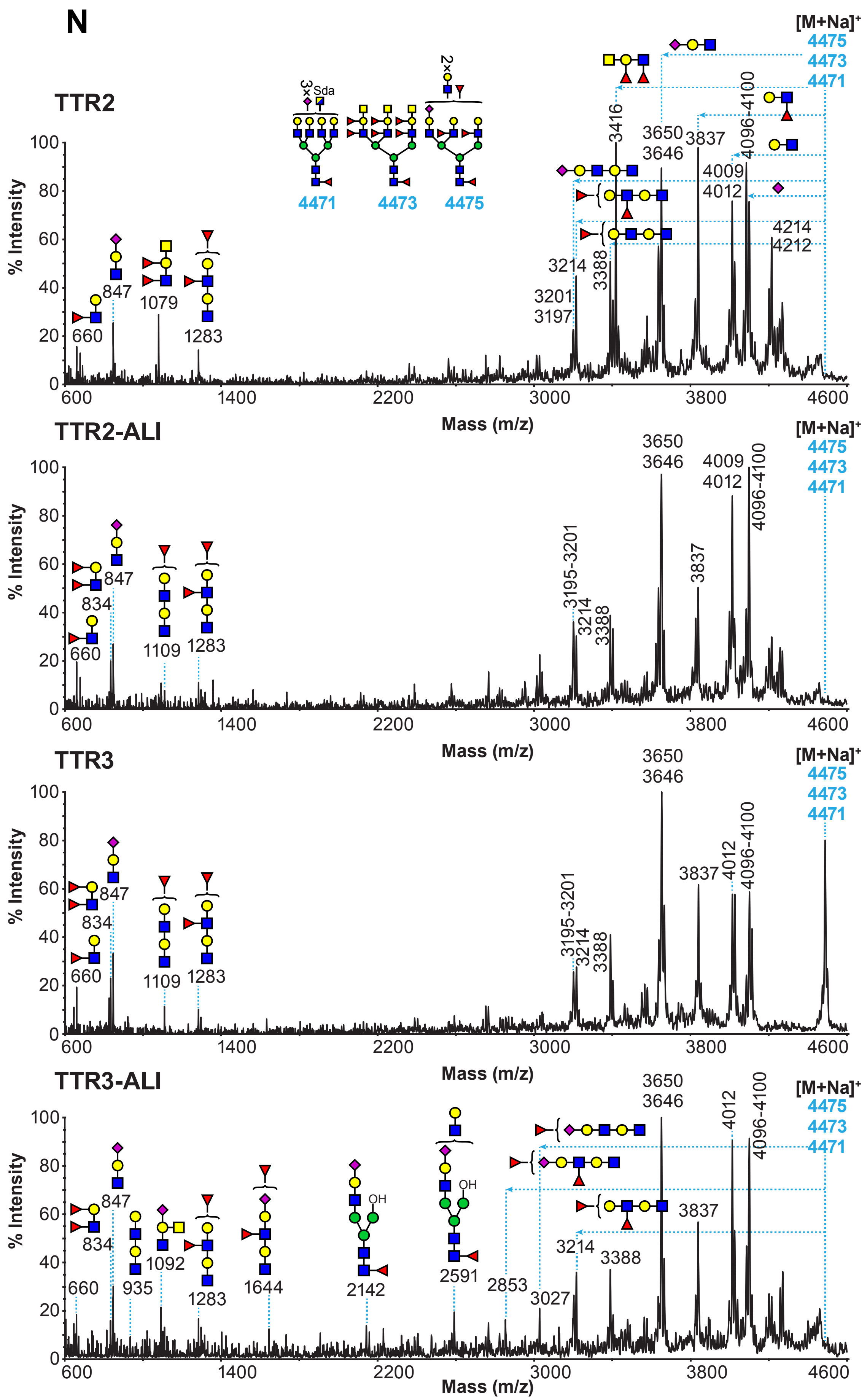

N
TTR2
TTR2-ALI
TTR3
TTR3-ALI
[M+Na]+
4475
4473
4471
3×Sda
2×
% Intensity
Mass (m/z)
660
847
1079
1283
3197
3201
3214
3388
3416
3650
3646
3837
4009
4012
4096-4100
4214
4212
834
1109
3195-3201
935
1092
1644
2142
2591
2853
3027
OH

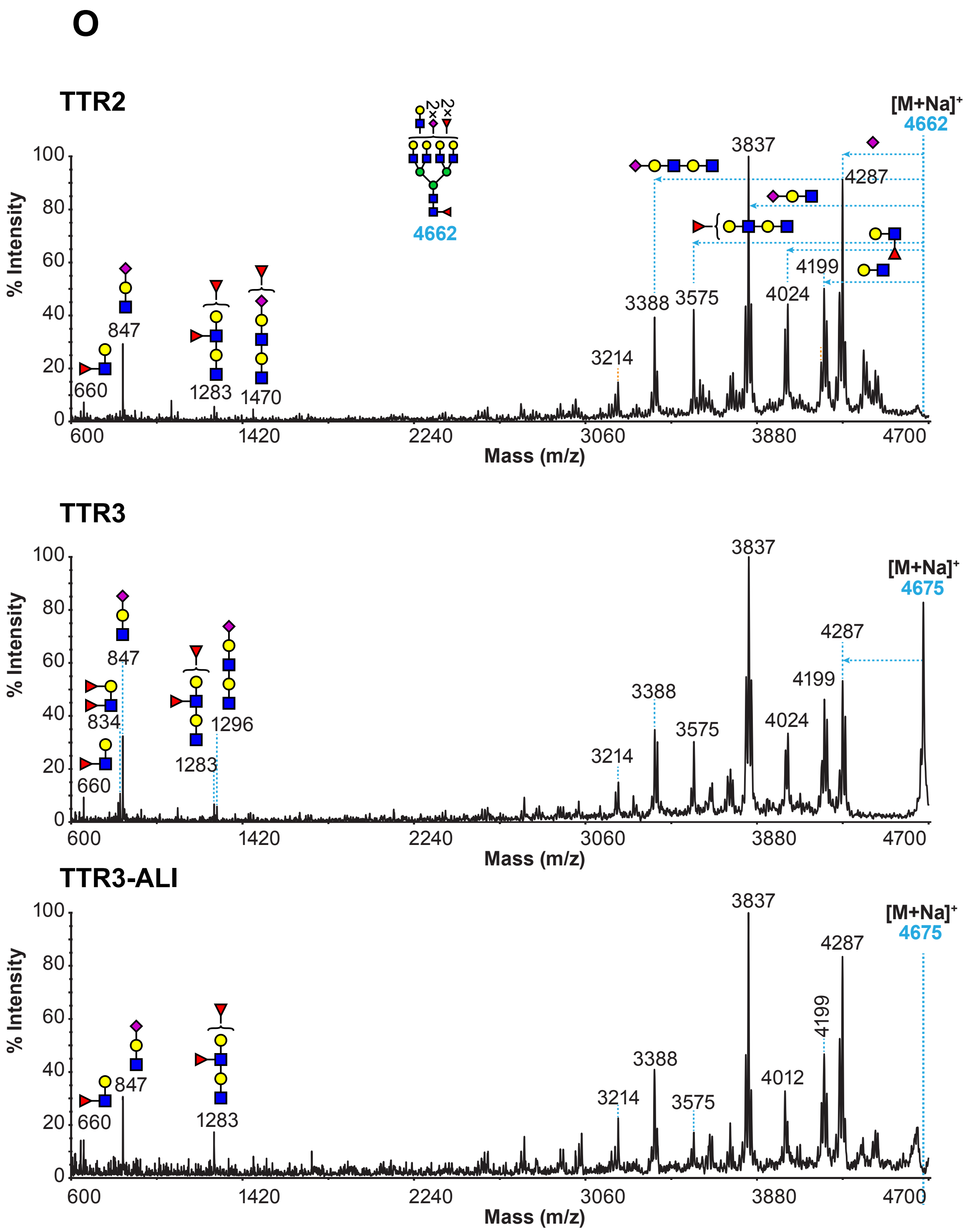
O
TTR2
[M+Na]+
4662
100
80
60
40
20
0
% Intensity
660
847
1283
1470
3214
3388
3575
3837
4024
4199
4287
600
1420
2240
3060
3880
4700
Mass (m/z)
TTR3
[M+Na]+
4675
660
834
847
1283
1296
3214
3388
3575
3837
4024
4199
4287
TTR3-ALI
[M+Na]+
4675
660
847
1283
3214
3388
3575
3837
4012
4199
4287

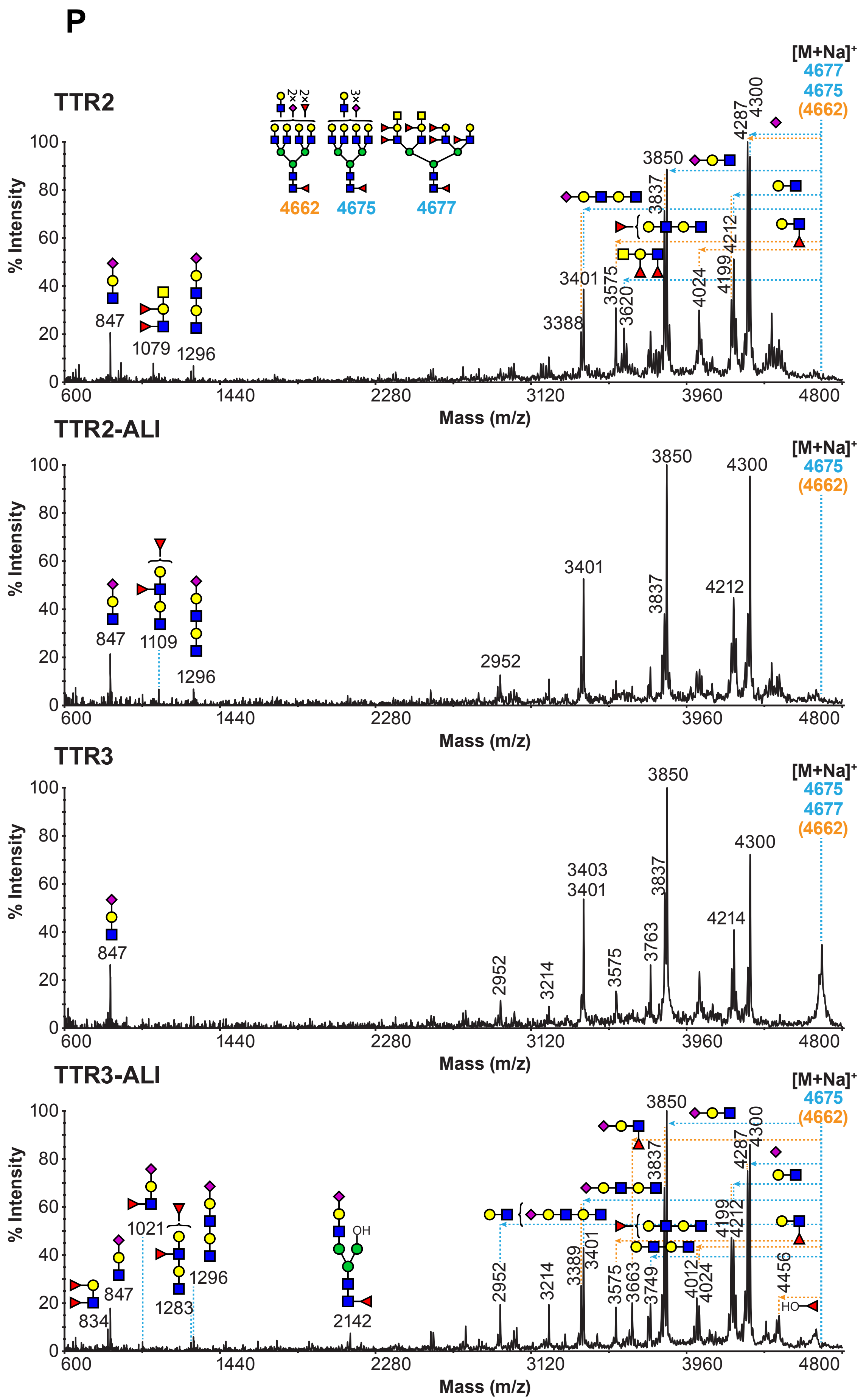

P
TTR2
TTR2-ALI
TTR3
TTR3-ALI
[M+Na]+
4677
4675
(4662)
4662
% Intensity
Mass (m/z)
847
1079
1296
1109
834
1021
1283
2142
2952
3214
3388
3389
3401
3403
3575
3620
3663
3749
3763
3837
3850
4012
4024
4199
4212
4214
4287
4300
4456

**Figure S17**

**MALDI-TOF/TOF MS/MS analysis of permethylated N-linked glycans of primary tracheal epithelial cells from human donors or after culture in air-liquid interface.** Molecular ions [M+Na]+ present in **Figures S1C** and **D** (TTr02, TTr03) and **Figure S16** (TTr02-ALI and TTr03-ALI) were selected for MALDI-TOF/TOF MS/MS analysis. (A) m/z 3140-3142 ion cluster, (B) m/z 3415, (C) m/z 3589-3591 ion cluster, (D) m/z 3677, (E) m/z 3763, (F) m/z 3864, (G) m/z 3950-3952 ion cluster, (H) m/z 4022-4026 ion cluster, (I) m/z 4198-4200 ion cluster, (J) m/z 4299-4301 ion cluster, (K) m/z 4314 ion, (L) m/z 4387, (M) m/z 4400-4402 ion cluster, (N) m/z 4471-4475 ion cluster, (O) m/z 4662, (P) m/z 4675-4677 ion cluster. Cartoon structures in the corresponding spectra do not represent all possible structural isomers, for clarity. Horizontal dashed lines correspond to the indicated fragment loss from the corresponding molecular ion $[M+Na]^+$. Vertical dashed lines indicate the m/z fragment peak that corresponds to the fragment loss as shown from the corresponding horizontal dashed line. Structures outside the bracket have not unequivocally been defined. For clarity, not all fragments were annotated. Given the high complexity of the MALDI-TOF MS, in terms of multiple and overlapping isotopes clustering, MALDI-TOF/TOF MS/MS molecular ion selection was on the corresponding isotope cluster. However, due to the very low relative abundance of the selected molecular ions, in certain cases adjacent molecular ions were also fragmented. The molecular ions selected for fragmentation are blue-coded color (with blue horizontal and vertical lines), while the adjacent molecular ions were orange-coded color (with orange horizontal and vertical lines). For clarity, not all horizontal yellow lines are shown and not all fragments are annotated.

(B) Major ions detected in the spectrum correspond to fragments deriving from the molecular ion at m/z 3415. The fragment at m/z 3027 corresponding to a loss of NeuAc residue from the molecular ion at m/z 3402 was less abundant than the corresponding fragment from the molecular ion at m/z 3415.

(C) The ion at m/z 2765 corresponded either to a fragment of NeuAc-LacNAc or $LacNAcFuc_2$ structure which have derived from the molecular ion at m/z 3589 or 3576 respectively. However, a characteristic fragment ion at m/z 3201 which corresponded to a loss of NeuAc was in minimum relative abundance. Therefore, the fragments presented in the spectrum corresponded mainly to the fragments deriving from the molecular ion at 3589. (D, H, J, N) The fragment ion at m/z 1092 corresponded to the Sda epitope.

**Figure S18**

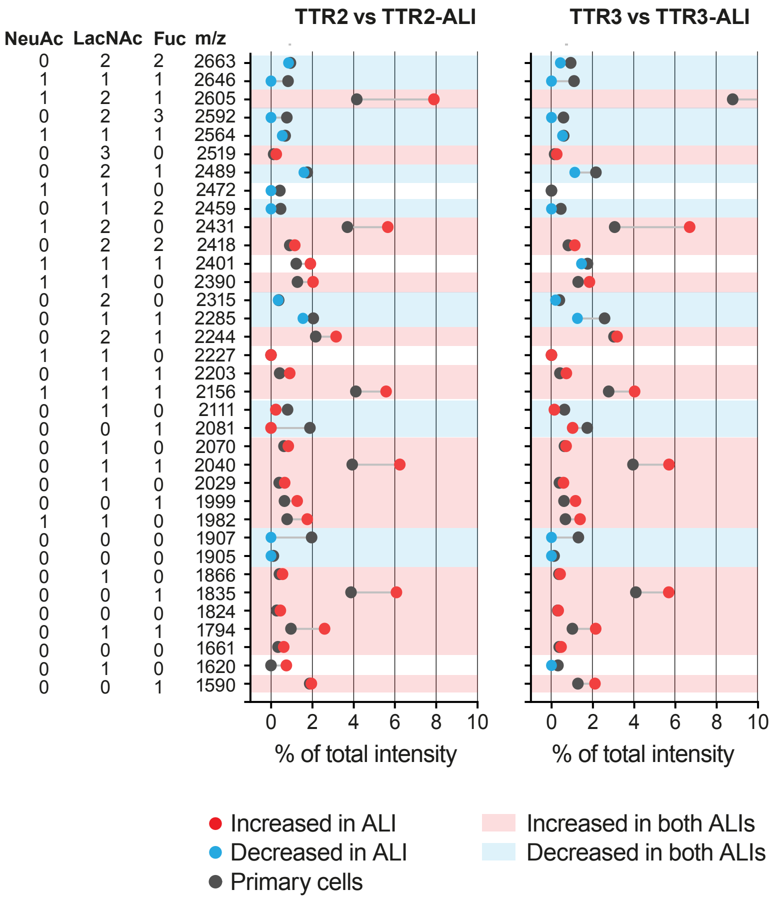

TTR2 vs TTR2-ALI
TTR3 vs TTR3-ALI
NeuAc LacNAc Fuc m/z
0 2 2 2663
1 1 1 2646
1 2 1 2605
0 2 3 2592
1 1 1 2564
0 3 0 2519
0 2 1 2489
1 1 0 2472
0 1 2 2459
1 2 0 2431
0 2 2 2418
1 1 1 2401
1 1 0 2390
0 2 0 2315
0 1 1 2285
0 2 1 2244
1 1 0 2227
0 1 1 2203
1 1 1 2156
0 1 0 2111
0 0 1 2081
0 1 0 2070
0 1 1 2040
0 1 0 2029
0 0 1 1999
1 1 0 1982
0 0 0 1907
0 0 0 1905
0 1 0 1866
0 0 1 1835
0 0 0 1824
0 1 1 1794
0 0 0 1661
0 1 0 1620
0 0 1 1590
0 2 4 6 8 10
% of total intensity
Increased in ALI
Decreased in ALI
Primary cells
Increased in both ALIs
Decreased in both ALIs

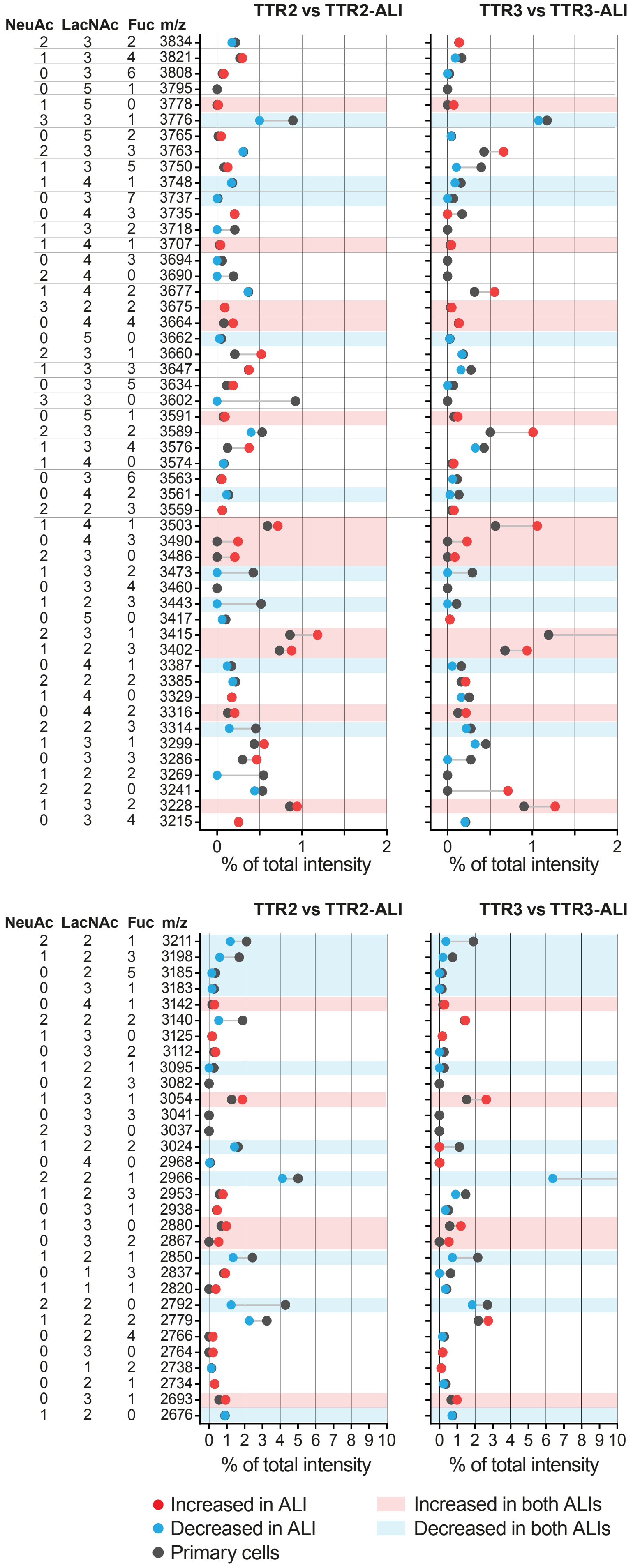
TTR2 vs TTR2-ALI
TTR3 vs TTR3-ALI
NeuAc LacNAc Fuc m/z
2 3 2 3834
1 3 4 3821
0 3 6 3808
0 5 1 3795
1 5 0 3778
3 3 1 3776
0 5 2 3765
2 3 3 3763
1 3 5 3750
1 4 1 3748
0 3 7 3737
0 4 3 3735
1 3 2 3718
1 4 1 3707
0 4 3 3694
2 4 0 3690
1 4 2 3677
3 2 2 3675
0 4 4 3664
0 5 0 3662
2 3 1 3660
1 3 3 3647
0 3 5 3634
3 3 0 3602
0 5 1 3591
2 3 2 3589
1 3 4 3576
1 4 0 3574
0 3 6 3563
0 4 2 3561
2 2 3 3559
1 4 1 3503
0 4 3 3490
2 3 0 3486
1 3 2 3473
0 3 4 3460
1 2 3 3443
0 5 0 3417
2 3 1 3415
1 2 3 3402
0 4 1 3387
2 2 2 3385
1 4 0 3329
0 4 2 3316
2 2 3 3314
1 3 1 3299
0 3 3 3286
1 2 2 3269
2 2 0 3241
1 3 2 3228
0 3 4 3215
0 1 2
% of total intensity
TTR2 vs TTR2-ALI
TTR3 vs TTR3-ALI
NeuAc LacNAc Fuc m/z
2 2 1 3211
1 2 3 3198
0 2 5 3185
0 3 1 3183
0 4 1 3142
2 2 2 3140
1 3 0 3125
0 3 2 3112
1 2 1 3095
0 2 3 3082
1 3 1 3054
0 3 3 3041
2 3 0 3037
1 2 2 3024
0 4 0 2968
2 2 1 2966
1 2 3 2953
0 3 1 2938
1 3 0 2880
0 3 2 2867
1 2 1 2850
0 1 3 2837
1 1 1 2820
2 2 0 2792
1 2 2 2779
0 2 4 2766
0 3 0 2764
0 1 2 2738
0 2 1 2734
0 3 1 2693
1 2 0 2676
0 1 2 3 4 5 6 7 8 9 10
% of total intensity
Increased in ALI
Decreased in ALI
Primary cells
Increased in both ALIs
Decreased in both ALIs

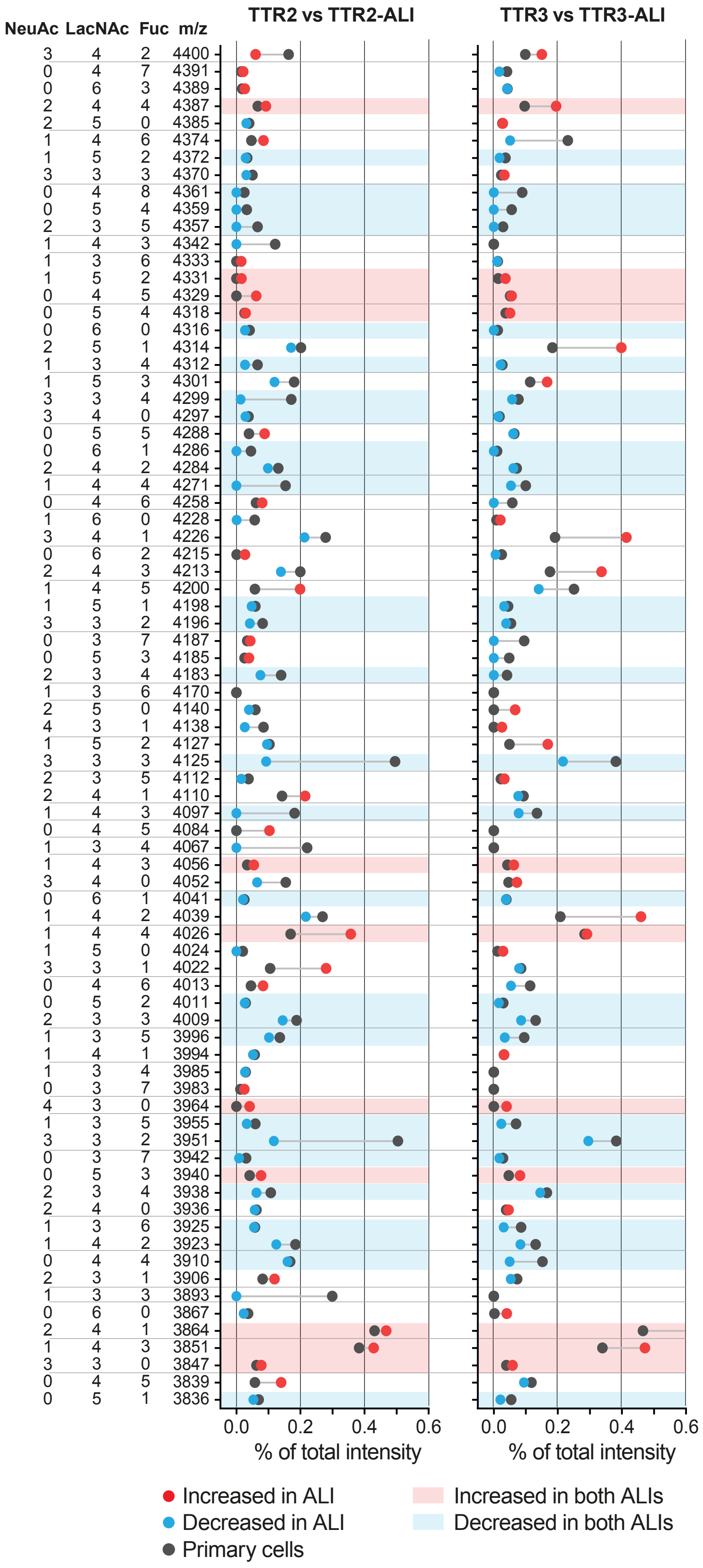
TTR2 vs TTR2-ALI
TTR3 vs TTR3-ALI
NeuAc LacNAc Fuc m/z
3 4 2 4400
0 4 7 4391
0 6 3 4389
2 4 4 4387
2 5 0 4385
1 4 6 4374
1 5 2 4372
3 3 3 4370
0 4 8 4361
0 5 4 4359
2 3 5 4357
1 4 3 4342
1 3 6 4333
1 5 2 4331
0 4 5 4329
0 5 4 4318
0 6 0 4316
2 5 1 4314
1 3 4 4312
1 5 3 4301
3 3 4 4299
3 4 0 4297
0 5 5 4288
0 6 1 4286
2 4 2 4284
1 4 4 4271
0 4 6 4258
1 6 0 4228
3 4 1 4226
0 6 2 4215
2 4 3 4213
1 4 5 4200
1 5 1 4198
3 3 2 4196
0 3 7 4187
0 5 3 4185
2 3 4 4183
1 3 6 4170
2 5 0 4140
4 3 1 4138
1 5 2 4127
3 3 3 4125
2 3 5 4112
2 4 1 4110
1 4 3 4097
0 4 5 4084
1 3 4 4067
1 4 3 4056
3 4 0 4052
0 6 1 4041
1 4 2 4039
1 4 4 4026
1 5 0 4024
3 3 1 4022
0 4 6 4013
0 5 2 4011
2 3 3 4009
1 3 5 3996
1 4 1 3994
1 3 4 3985
0 3 7 3983
4 3 0 3964
1 3 5 3955
3 3 2 3951
0 3 7 3942
0 5 3 3940
2 3 4 3938
2 4 0 3936
1 3 6 3925
1 4 2 3923
0 4 4 3910
2 3 1 3906
1 3 3 3893
0 6 0 3867
2 4 1 3864
1 4 3 3851
3 3 0 3847
0 4 5 3839
0 5 1 3836
0.0 0.2 0.4 0.6
% of total intensity
0.0 0.2 0.4 0.6
% of total intensity
Increased in ALI
Decreased in ALI
Primary cells
Increased in both ALIs
Decreased in both ALIs

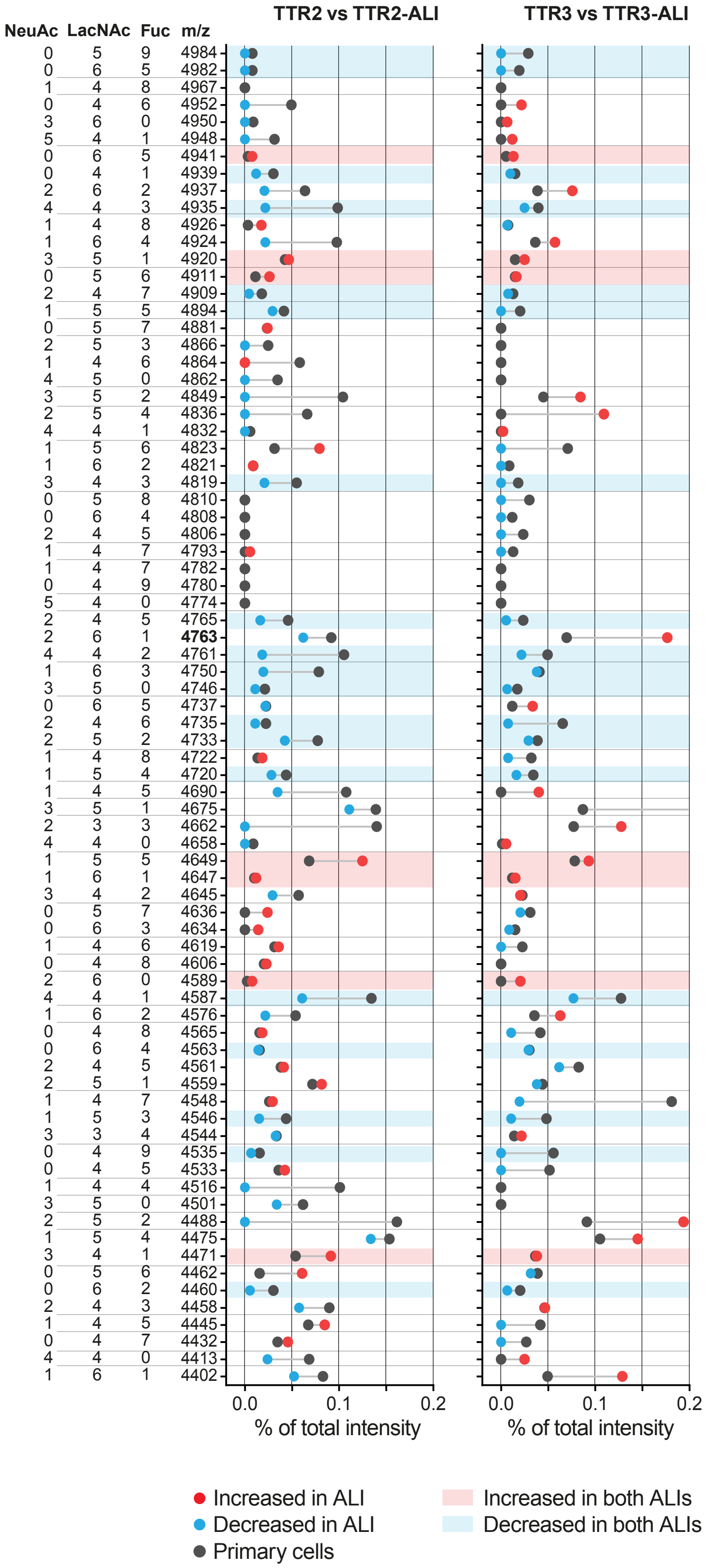
TTR2 vs TTR2-ALI
TTR3 vs TTR3-ALI
NeuAc LacNAc Fuc m/z
0 5 9 4984
0 6 5 4982
1 4 8 4967
0 4 6 4952
3 6 0 4950
5 4 1 4948
0 6 5 4941
0 4 1 4939
2 6 2 4937
4 4 3 4935
1 4 8 4926
1 6 4 4924
3 5 1 4920
0 5 6 4911
2 4 7 4909
1 5 5 4894
0 5 7 4881
2 5 3 4866
1 4 6 4864
4 5 0 4862
3 5 2 4849
2 5 4 4836
4 4 1 4832
1 5 6 4823
1 6 2 4821
3 4 3 4819
0 5 8 4810
0 6 4 4808
2 4 5 4806
1 4 7 4793
1 4 7 4782
0 4 9 4780
5 4 0 4774
2 4 5 4765
2 6 1 **4763**
4 4 2 4761
1 6 3 4750
3 5 0 4746
0 6 5 4737
2 4 6 4735
2 5 2 4733
1 4 8 4722
1 5 4 4720
1 4 5 4690
3 5 1 4675
2 3 3 4662
4 4 0 4658
1 5 5 4649
1 6 1 4647
3 4 2 4645
0 5 7 4636
0 6 3 4634
1 4 6 4619
0 4 8 4606
2 6 0 4589
4 4 1 4587
1 6 2 4576
0 4 8 4565
0 6 4 4563
2 4 5 4561
2 5 1 4559
1 4 7 4548
1 5 3 4546
3 3 4 4544
0 4 9 4535
0 4 5 4533
1 4 4 4516
3 5 0 4501
2 5 2 4488
1 5 4 4475
3 4 1 4471
0 5 6 4462
0 6 2 4460
2 4 3 4458
1 4 5 4445
0 4 7 4432
4 4 0 4413
1 6 1 4402
0.0 0.1 0.2
% of total intensity
0.0 0.1 0.2
% of total intensity
Increased in ALI
Decreased in ALI
Primary cells
Increased in both ALIs
Decreased in both ALIs

**Figure S18**

**Comparison of the relative abundances of the N-linked glycans of primary tracheal epithelial cells from human donors against the same cells cultured in air-liquid interface.** Lollipop graphs depicting the increase or decrease of the relative abundance of N-linked glycans of epithelial cells cultured in air-liquid interface (ALI) deriving from human trachea donors against the same primary epithelial cells (non-cultured). Left graph shows the comparison of primary TTr02 vs cultured TTr02-ALI, right graph shows the comparison of primary TTr03 vs cultured TTr03-ALI. For a given m/z value, on both graphs, a red or blue dot correspond to an increased or decreased relative abundance of a N-linked glycan deriving from the cultured ALI cells compared to the same N-linked glycan derived from the primary cells (black dot). The NeuAcs, LacNAcs and fucose (Fuc) columns correspond to the number of residues/units that consist the N-linked glycan detected at m/z value. Red or blue shaded areas across both graphs correspond to N-linked glycans (m/z) that increased or decreased respectively on both graphs (TTr02-TTr02-ALI and TTr03-TTr03-ALI). For clarity the differences of the molecular ions at m/z 2605, 2966, 3415 and 3864 were outside the range of X-axis. Horizontal black lines across both graphs separate the various molecular ion clusters detected in the MALDI-TOF MS (**Figure S16**).

### Figure S19

## A

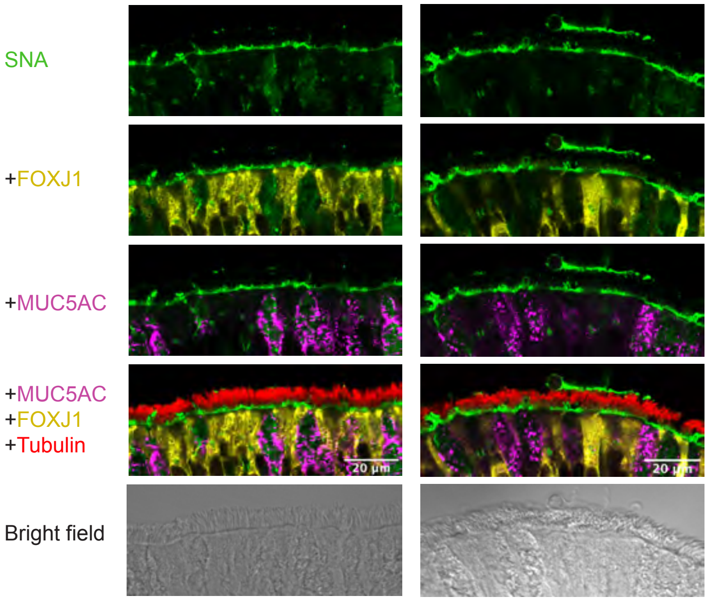


## B

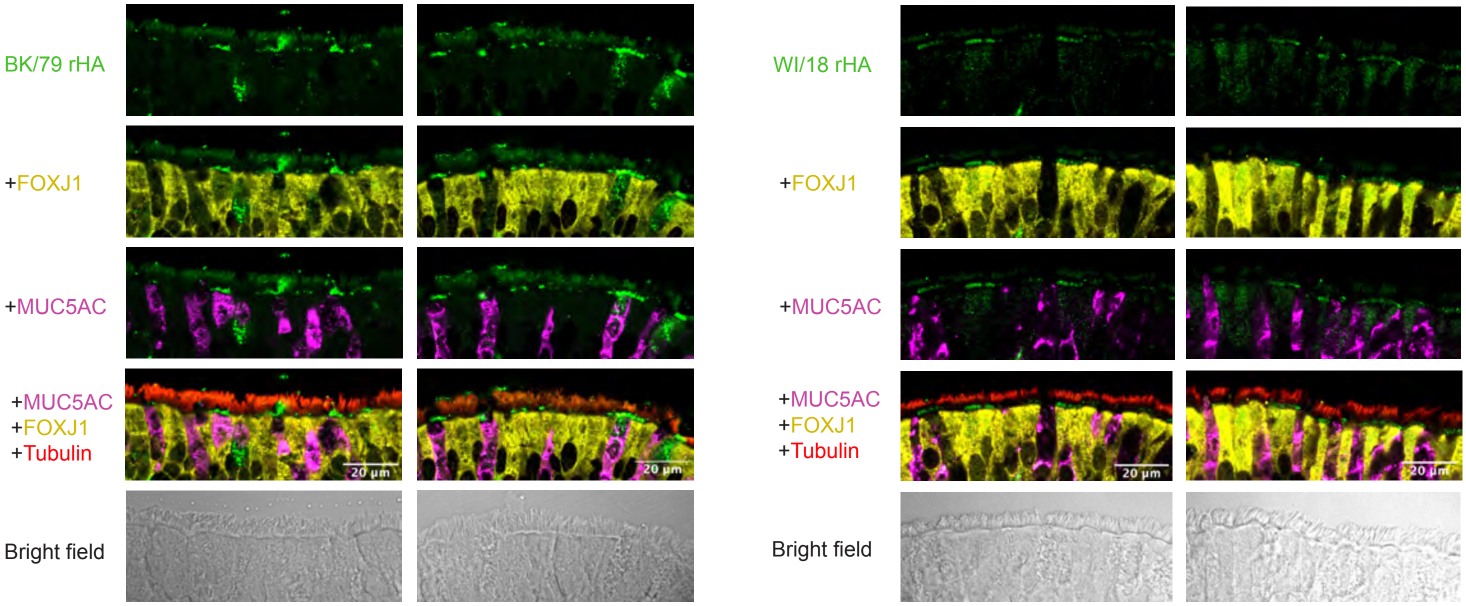

**Figure S19**

**IF of human trachea sections.** Additional IF images of human trachea sections shown in **Figure 5**. Sections were stained with cell type markers and either SNA lectin (**A**), or BK/79 and WI/18 recombinant HA (green) (**B**). Bright field images of each section are also shown.

**Figure S20**

**A**

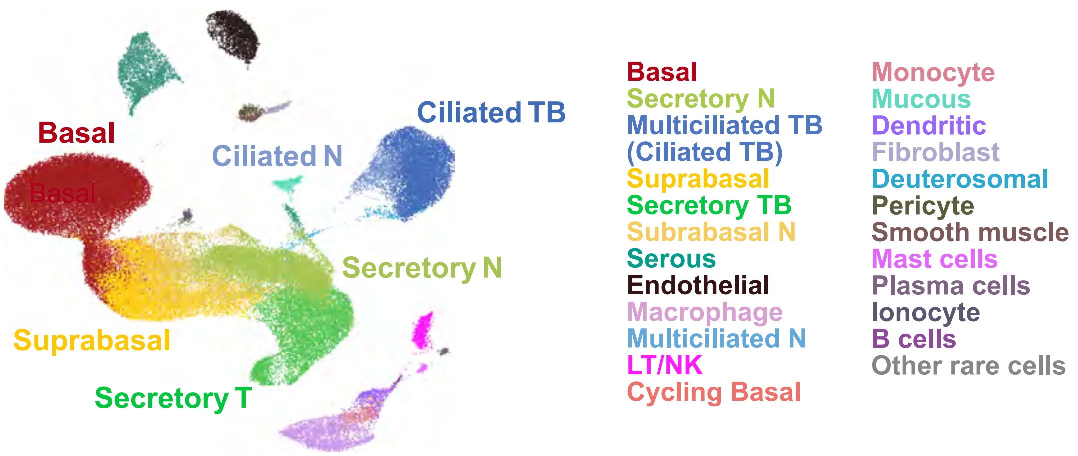


**B**

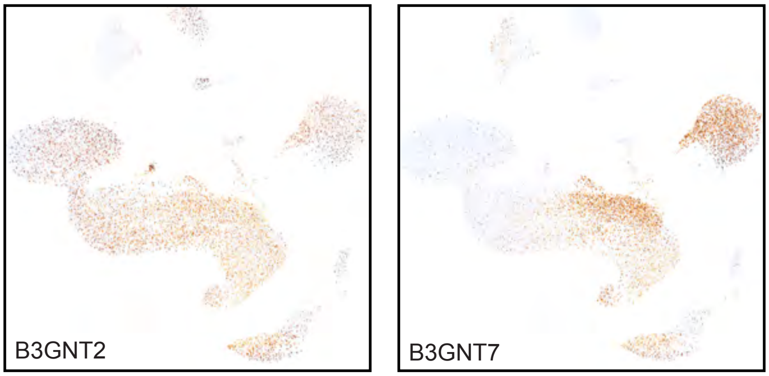

**Figure S20**

**Additional scRNAseq analyses.** Related to **Figure 5**. (**A**) UMAP representation of the dataset with a complete annotation of populations (**Figure 5F**). (**B**) Expression of *B3GNT2* (*left*) and *B3GNT7* (*right*) genes.
